# Long non-coding RNA-3′UTR interactions coordinate GABA signaling in the brain

**DOI:** 10.1101/2025.09.05.674572

**Authors:** R. Loker, C. Desplan

## Abstract

How functionally related genes are coordinately expressed within a given cell type remains poorly understood. Here we reveal that the 3′UTR, beyond binding RNA-binding proteins and microRNAs, mediates heterotypic RNA-RNA interactions that confer cell type-specific gene expression, and demonstrate that this function is essential for GABAergic signaling in *Drosophila*. We show the vesicular GABA transporter *VGAT* is transcribed pan-neuronally but its translation is repressed in non-GABAergic neurons by miR-7 through three conserved 3′UTR sites. We identify a GABAergic neuron-specific lncRNA, GABA UnLocker (*Gul*), with sequences complementary to all three *VGAT* miR-7 sites that derepresses *VGAT* translation. *Gul* is co-expressed with the adjacent *Gad1* gene, which produces GABA, in all GABAergic neurons, such that a single GABAergic neuron-specific transcriptional input instructs both GABA synthesis and, via *Gul*, the transporter for GABA release. *Gul* also targets other mRNAs in GABA metabolism, suggesting that it coordinates GABAergic gene expression through 3′UTR RNA-RNA interactions.

## Introduction

Precise regulation of gene expression is fundamental to the development and function of the nervous system. Both transcriptional and post-transcriptional mechanisms can act in concert to determine where, when, and at what levels individual gene products ultimately accumulate ^1–4^. Yet the functional output of a neuron rarely depends on the precise expression of a single gene in isolation; rather, it emerges from the coordinated expression of suites of functionally related genes acting together within a shared molecular pathway. The execution of a specific neurotransmitter modality, for example, requires the molecular machinery for synthesis, packaging, and release of the neurotransmitter to be co-expressed in the same cell at appropriate relative levels, and disruptions in this balance contribute to a wide range of neurological and psychiatric disorders ^5–7^. While transcription factors orchestrate neuronal identity programs ^8,9^, the mechanisms by which individual components of a specific molecular pathway are coordinately regulated post-transcriptionally within a defined cell type remain poorly understood ^10–13^. Such regulation is particularly important when pathway components are transcribed broadly yet must be functionally restricted to, or required at different levels in, specific neuronal populations.

The 3′ untranslated region (3′UTR) of mRNAs is a key regulatory platform that integrates inputs from RNA-binding proteins (RBPs) and microRNAs (miRNAs) to control mRNA stability, translation efficiency, and subcellular localization ^2,14–19^. The combinatorial binding of these regulators to 3′UTR sequence elements allows a single transcript to be differentially regulated across cell types and developmental contexts, making the 3′UTR a central determinant of where and when a gene product is functionally expressed. Beyond these protein and miRNA inputs, 3′UTRs can also interact directly with one another: homotypic base-pairing between copies of the same 3′UTR drives dimerization and granule assembly that direct mRNA localization in the *Drosophila* female germline ^20–23^. A third class of post-transcriptional regulator are the long non-coding RNAs (lncRNAs). Although thousands are expressed in the mammalian brain, with many exhibiting striking neuronal subtype-specific expression patterns ^24–26^, relatively few have been demonstrated to function in the cytoplasm to regulate gene expression ^27–30^. Among the cytoplasmic functions described for lncRNAs, some act directly on target mRNAs through base-pairing; for example, half-STAU1-binding site RNAs (1/2-sbsRNAs) base-pair with complementary Alu elements in the 3′UTRs of specific target mRNAs to recruit the double-stranded RNA-binding protein STAU1 and trigger their decay ^31^. Others act as molecular sponges that sequester RBPs ^32,33^ or miRNAs away from their targets, most notably the circular RNA CDR1as (ciRS-7), which contains approximately 70 conserved binding sites for miR-7 and is highly expressed in the brain ^34^.

Here, we uncover a previously unrecognized mechanism of cell type-specific post-transcriptional gene regulation, in which a lncRNA whose expression is restricted to a specific cell population relieves miRNA-mediated silencing of a target mRNA at its 3′UTR. We show that this mechanism is essential for expression of the vesicular GABA transporter VGAT in GABAergic neurons in *Drosophila*, and thus for signaling of the inhibitory neurotransmitter GABA. *VGAT* mRNA is transcribed throughout the nervous system, yet VGAT protein is expressed exclusively in GABAergic neurons where it is required for packaging GABA into synaptic vesicles for release. We demonstrate that *VGAT* mRNA is repressed outside of GABAergic neurons by miR-7, which targets three conserved binding sites in the *VGAT* 3′UTR, and that escape from this repression within GABAergic neurons requires a GABAergic neuron-specific lncRNA, which we have named GABA UnLocker (*Gul*). *Gul* does not contain a miR-7 binding site and therefore cannot sequester the miRNA as CDR1as does; instead it mimics miR-7, carrying three copies of the miR-7 5′ end seed sequence, each complementary to one of the miR-7 target sites in the *VGAT* 3′UTR and, at one of them, extending over 22 uninterrupted bases. The *Gul* gene lies directly adjacent to, and is divergently transcribed from, *Gad1*, which encodes the essential enzyme for GABA synthesis, and both genes share a GABAergic neuron-specific expression pattern, likely due to shared transcriptional regulatory elements. This mechanism thereby links the transcriptional state of one locus (*Gad1*/*Gul*) to the translational output of a functionally related gene at a distinct locus (*VGAT*), through a lncRNA intermediary. Furthermore, we find that *Gul* contains complementary sequences to other mRNAs encoding proteins involved in nearly every aspect of GABA metabolism and signaling, with and without predicted miR-7 sites. The protein output of those that also carry a miR-7 site is preferentially regulated in GABAergic neurons. This suggests that a single transcriptional input at the *Gad1*/*Gul* locus is propagated post-transcriptionally in *trans* to functionally related genes at dispersed loci, achieving operon-like coordination without specific transcriptional regulation of the individual genes. Similar lncRNA-mediated coordination through 3′UTR complementarity might be more widely used in the nervous system and beyond.

## Results

### VGAT mRNA is transcribed in all neurons but translated only in GABAergic neurons

*VGAT* function is specifically required in GABAergic neurons, a population that accounts for only ∼10% of *Drosophila* neurons ^35^. Surprisingly, analysis of existing single cell RNA-seq datasets from the optic lobe ^36^, the ventral nerve cord ^37^, and central brain ^38^, revealed that all *Drosophila* neuronal clusters contain *VGAT* mRNA that is readily detectable even in non-GABAergic clusters (glutamatergic and cholinergic), in contrast to glial clusters, which do not transcribe it. However, in the three regions respectively, its levels are 5.3-, 4.0-, and 3.2-fold higher in GABAergic clusters compared to non-GABAergic clusters (**Fig. S1A-C**). Using RNA *in-situ* hybridization, we confirmed that *VGAT* mRNA is present in all the neurons of the optic lobe including lamina and T4/T5 motion sensing neurons that are exclusively non-GABAergic populations ^39–41^ **(Fig. 1A and Fig. S1D).**

**Figure 1.**
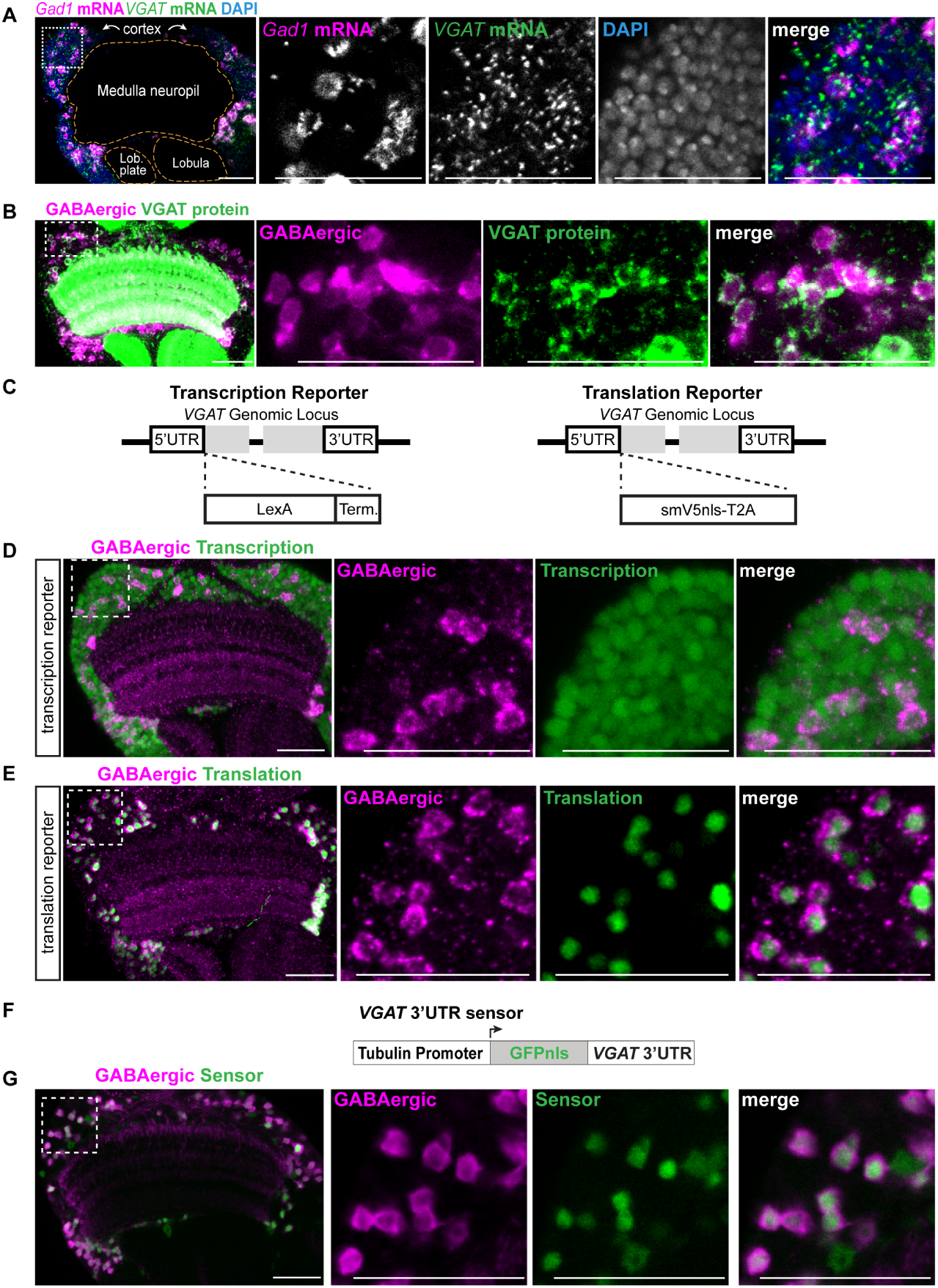
*VGAT* mRNA is transcribed in all neurons but translated only in GABAergic neurons. **(A)** HCR RNA-FISH of the adult *Drosophila* optic lobe showing the distribution of *VGAT* mRNA (green) relative to GABAergic neurons (magenta) marked by *Gad1* mRNA. **(B)** Immunofluorescence using VGAT antibody showing the distribution of VGAT protein (green) relative to GABAergic neurons (magenta) labeled using a *Gad1-LexA>myrCherry*. **(C)** Diagram of knock-in alleles separately characterize *VGAT* transcription and translation. The transcription reporter consists of a LexA coding sequence followed by Sv40 and Hsp70 terminators inserted into the N-terminus of the VGAT coding sequences. The translation reporter consists of a nuclear-localized spaghetti monster-V5 (smV5nls) and T2A sequence inserted into the N-terminus of the VGAT coding sequences. **(D)** LexA activity in the optic lobe of transcription reporter flies, read out with *LexAop-GFP* and GABAergic neurons (magenta) marked with a Gad1 antibody. **(E)** V5 immunofluorescence in translation reporter flies with GABAergic neurons (magenta) marked with a Gad1 antibody. **(F)** Diagram of *VGAT* 3′UTR sensor transgene consisting of a tubulin promoter driven GFPnls with the 3′UTR of *VGAT*. **(G)** Activity of *VGAT* 3′UTR sensor (green) in the optic lobe of wild type flies relative to GABAergic neurons (magenta) marked with *Gad1-LexA>myrCherry*. Scale bar: 20 μm. Inset: region of medulla cortex.

Given that *VGAT* functions only in GABAergic neurons, we examined its protein expression using a previously validated antibody ^42^. Staining in flies carrying the GABAergic marker *Gad1*-LexA>myrCherry showed that VGAT protein is present exclusively in GABAergic neurons **(Fig. 1B)**, indicating that *VGAT* mRNA is either repressed post-transcriptionally in other neuronal types or translated there and the protein rapidly degraded.

To determine whether the difference in *VGAT* mRNA levels between GABAergic and non-GABAergic neurons is mediated by differences in the level of transcription or post-transcriptional regulation, *e.g.* mRNA stability, we utilized an allele in which the coding sequence of the LexA transcriptional activator was inserted upstream of the endogenous *VGAT* coding sequence ^43^, hereafter referred to as the transcription reporter **(Fig. 1C)**. Importantly, the LexA coding sequence is immediately followed by an SV40 transcriptional terminator and a 3′UTR from the *Hsp70* locus and is therefore not subjected to post-transcriptional regulation through the *VGAT* 3′UTR, although any potential inputs of the 5′UTR are preserved. Imaging of LexA activity in the optic lobe of flies containing this allele using a LexAop-GFP reporter revealed a uniform expression pattern in all neurons **(Fig. 1D)**, providing strong evidence that *VGAT* is transcribed pan-neuronally and that post-transcriptional regulation accounts for the lack of protein expression and the increased mRNA abundance in GABAergic neurons. Having established that *VGAT* mRNA is present throughout the nervous system, we next asked whether this mRNA is translated into protein outside of GABAergic neurons.

To address this, we generated an allele containing a nuclear-localized ‘spaghetti monster’ V5 (smV5) tag followed by a self-cleaving T2A peptide, inserted in frame at the N-terminus of the endogenous *VGAT* coding sequence and leaving its 3′UTR intact, hereafter referred to as the translation reporter **(Fig. 1C)**. Immunofluorescence using V5 antibody in this line confirmed a GABAergic neuron-specific expression pattern **(Fig. 1E)**. As the smV5 protein is separated from VGAT via the 2A peptide cleavage signal, it is not subjected to any post-translational regulation that may affect VGAT. We can therefore conclude that *VGAT* mRNA is not translated outside of GABAergic neurons.

Finally, to confirm that post-transcriptional regulation through the 3′UTR is sufficient to restrict *VGAT* expression to GABAergic neurons, we engineered a *VGAT*-3′UTR sensor in which the tubulin promoter drives ubiquitous transcription of GFP with the 3′UTR of *VGAT* **(Fig. 1F)**. Expression of GFP protein from this reporter was restricted to the GABAergic lineage **(Fig. 1G)**, confirming that the *VGAT* 3′UTR alone is sufficient to spatially restrict expression to GABAergic neurons in the *Drosophila* nervous system. Together, these results demonstrate that while *VGAT* is transcribed pan-neuronally, its translation is restricted to GABAergic neurons through the action of its 3′UTR.

### miR-7 represses VGAT translation in non-GABAergic neurons

Having established that the *VGAT* 3′UTR is sufficient to restrict translation to GABAergic neurons (**Fig. 1G**), we next asked what mechanism confers this specificity. A 3′UTR can impose such regulation through binding RBPs or miRNAs. Either could repress *VGAT* outside the GABAergic population; an RNA-binding protein could also promote its expression within it. The sequence of the *VGAT* 3′UTR itself strongly implicated a specific miRNA: it contains three predicted binding sites (referred to as sites a, b, and c, 5′ to 3′; **Fig. 2A**) for the deeply conserved miR-7, a miRNA that regulates development and function of the nervous system from flies to mammals ^34,44–48^ and is highly expressed in the adult *Drosophila* nervous system based on existing small RNA sequencing ^49,50^ **(Fig. S2A)**. All three sites are completely conserved across the entire drosophilid group, spanning 50 million years **(Fig. 2A)**. The three predicted miR-7 target sites in the *VGAT* 3′UTR are the highest number in any gene in the *Drosophila* genome (**Fig. S2B; Supp. Table 3**). Moreover, each of these miR-7 sites is a high-affinity 8mer, i.e. a perfect complement to the miR-7 seed sequence followed by an adenosine that increases site efficacy **(Fig. 2A)** ^16,51^. Furthermore, two of the sites (b and c) are positioned only 15 nucleotides apart, which has been previously shown to be within the optimal spacing for synergistic regulation by miRNA targeting and strengthened mRNA repression ^52–54^. Predicted duplexes of each site to the full 23-base miR-7 sequence also revealed extensive base pairing in the 3′ region outside of the miR-7 seed **(Fig. S2C)**, which has been suggested to enhance repression by miRNAs by stabilizing the target-bound complex ^52,55^.

**Figure 2.**
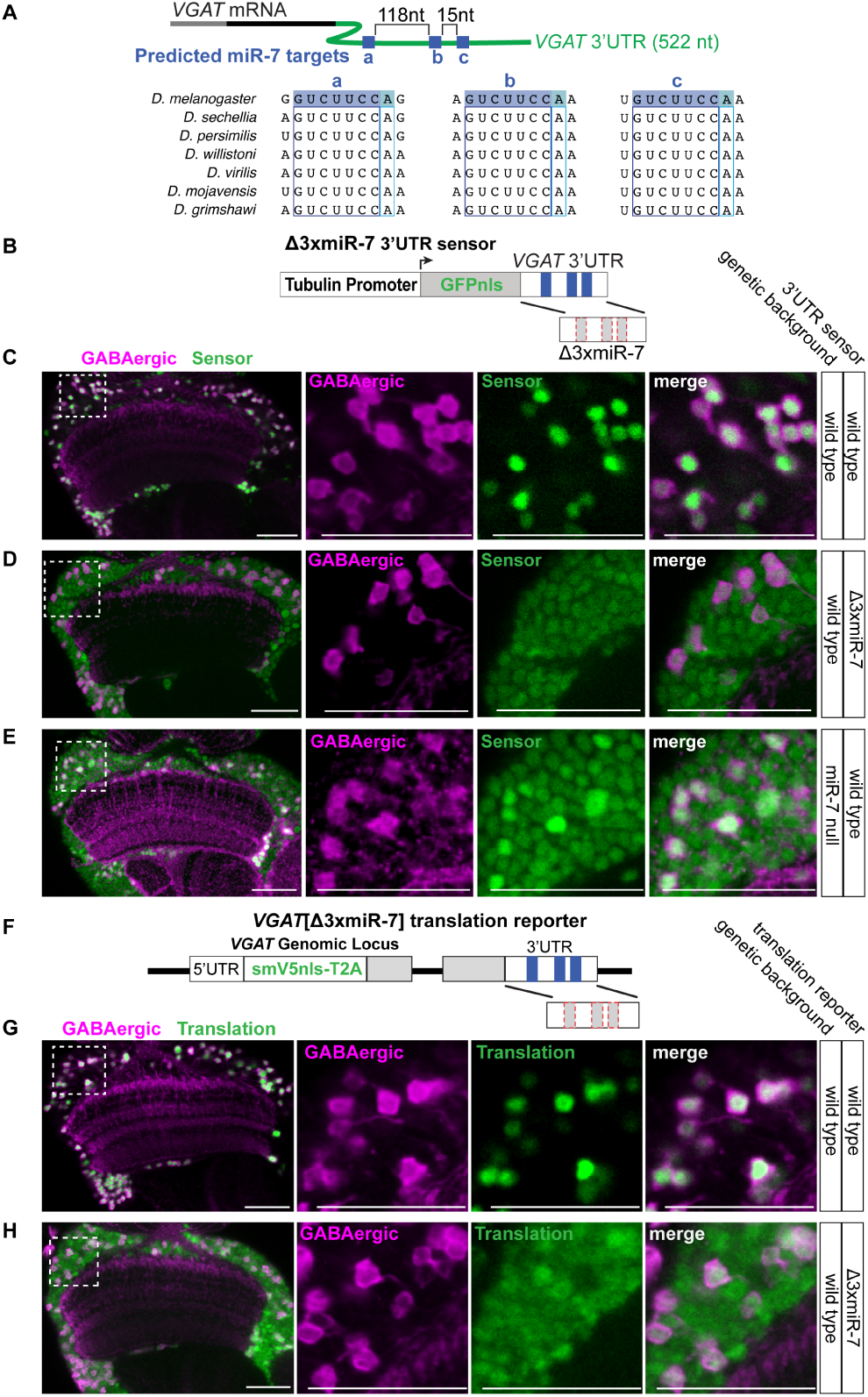
miR-7 represses *VGAT* translation in non-GABAergic neurons. **(A)** Diagram of the *VGAT* mRNA showing the three predicted miR-7 target sites (a, b and c, 5′ to 3′) in the 522 nt 3′UTR and the spacing between them, above an alignment of each site across seven Drosophilid species. The 8mer match to the miR-7 seed complement is highlighted in blue, comprising the 7mer recognition site plus the trailing adenosine colored in a lighter shade. **(B)** Diagram of the *VGAT* Δ3xmiR-7 3′UTR sensor transgene, the transgene in Fig. 1F with the three miR-7 target sites deleted. **(C)** Activity of the wild type *VGAT* 3′UTR sensor (green) in the optic lobe of wild type flies relative to GABAergic neurons (magenta) marked with *Gad1-LexA>myrCherry*. **(D)** Activity of the Δ3xmiR-7 sensor (green) in the optic lobe of wild type flies relative to GABAergic neurons (magenta) marked with *Gad1-LexA>myrCherry*. **(E)** Activity of the wild type *VGAT* 3′UTR sensor (green) in the optic lobe of miR-7 null flies relative to GABAergic neurons (magenta) marked with a GABA antibody. **(F)** Diagram of the endogenous *VGAT* Δ3xmiR-7 translation reporter allele, the translation reporter in Fig. 1C with the three miR-7 target sites deleted from the *VGAT* 3′UTR. **(G)** V5 immunofluorescence (green) in wild type translation reporter flies with GABAergic neurons (magenta) marked with *Gad1-LexA>myrCherry*. **(H)** V5 immunofluorescence (green) in Δ3xmiR-7 translation reporter flies with GABAergic neurons (magenta) marked with *Gad1-LexA>myrCherry*. Scale bar: 20 μm. Inset: region of medulla cortex.

To test whether miR-7 has a role in *VGAT* repression, we used both transgenic and genome engineering-based approaches. First, we modified the tubulin>GFP-*VGAT* 3′UTR transgenic sensor described above **(Fig. 1F-G)**, by deleting the three miR-7 8mer target sites in the *VGAT* 3′UTR (Δ3xmiR-7, **Fig. 2B)** and examining its expression in the optic lobe. In contrast to the wild type construct, which is repressed in non-GABAergic neurons **(Fig. 1F-G and Fig. 2C)**, GFP from the Δ3xmiR-7 construct was expressed in all neurons at a similar level **(Fig. 2D and Fig. S3; Supp. Table 4)**. We next examined the wild type sensor in flies homozygous for a miR-7 null allele ^56^. In this genetic background, the *VGAT* wild type 3′UTR sensor was also derepressed in all non-GABAergic neurons **(Fig. 2E)**, consistent with miR-7 being required for repression of the *VGAT* 3′UTR in non-GABAergic neurons. Notably, in this background GFP sensor levels retained a larger GABAergic neuron-bias compared to the Δ3xmiR-7 sensor level **(Fig. 2D-E and Fig. S3)**. Taken together, these results support a role for miR-7 in repressing *VGAT* in non-GABAergic neurons and suggest that the same sites are also acted on by an additional GABAergic neuron-specific factor that promotes higher expression in the GABAergic population (see below).

Next, we tested whether the loss of miR-7 input drives derepression of *VGAT* translation in the endogenous context. To do so, we edited the *VGAT* gene locus within the background of the smV5-T2A-*VGAT* translation reporter described above, generating a new allele with each of the three miR-7 target sites deleted (Δ3xmiR-7, **Fig. 2F)**. In contrast to the wild type 3′UTR allele **(Fig. 1E and Fig. 2G)**, loss of the miR-7 inputs caused pan-neuronal translation of smV5 **(Fig. 2H)**, consistent with our results using the transgenic sensor **(Fig. 2D)**. Furthermore, quantification of expression of this reporter between GABAergic and non-GABAergic neurons indicated similar levels, matching the result of the same Δ3xmiR-7 modification made in the context of the *VGAT* 3′UTR transgenic sensor **(Fig. S3)**, extending the finding that loss of those sites results in near-uniform *VGAT* expression to the endogenous locus.

Together, these findings show that the pan-neuronally transcribed *VGAT* mRNA is repressed in non-GABAergic neurons by miR-7, but they do not explain how *VGAT* escapes that repression in GABAergic neurons. The simplest explanation would be that miR-7 is less active in this population. Two observations argue against this possibility. First, a transgenic sensor that reports miR-7 activity at single-cell resolution (**Fig. S4A**) showed comparable activity in GABAergic and non-GABAergic neurons, with only a 1.35-fold bias in mean intensity toward the GABAergic population (**Fig. S4B-D**), whereas removing miR-7 input altogether, in both the sensor and the endogenous reporter, drove near-uniform translation of *VGAT* across neurons (**Fig. 2H** and **Fig. S3**). Therefore, a difference in miR-7 activity of the size we can measure cannot on its own account for that restriction. Second, a general reduction in miR-7 activity would relieve its targets as a class, yet conserved miR-7 targets co-varied with *Gad1* no more strongly than transcripts carrying no miR-7 site in any of the three regions of the adult CNS, regardless of site strength (**Fig. S4E**). Escape from miR-7 repression is therefore conferred locally, at the *VGAT* 3′UTR itself, rather than by a change in miR-7 availability.

### A GABAergic neuron-specific lncRNA contains complementary sequences to miR-7 target sites in the VGAT 3′UTR

Two additional classes of molecule could act at these miR-7 sites. RNA-binding proteins can modulate miRNA targeting at individual sites, either cooperatively ^57^ or by competing for access ^58^, and therefore an RBP depleted or enriched in GABAergic neurons could influence miR-7 activity locally at the *VGAT* 3′UTR. Screening 696 annotated RNA-binding proteins across the adult CNS ^36–38^, we found none with expression restricted to, or specifically absent from, GABAergic neurons: the most GABAergic-enriched RBP was 2.7-fold, while *Gad1* itself was 17-to 37-fold enriched (**Fig. S5; Supp. Table 5**). This does not exclude a broadly transcribed RBP whose abundance or activity is controlled post-transcriptionally, but it offered no candidate to test directly.

We therefore considered the second possibility, that the sites are occupied by another RNA, as antisense transcripts can stabilize their sense counterpart through direct base pairing ^59,60^, although only a small number of such cases have been described. The *VGAT* locus is not covered by any antisense transcript, but we hypothesized that an RNA transcribed elsewhere in the genome could act in *trans* to pair with the *VGAT* 3′UTR. We therefore performed a BLAST search using the full *VGAT* 3′UTR to identify transcripts capable of such an interaction. The top-scoring hit by both bit-score and E-value lay within the annotated 3′UTR of an uncharacterized gene, *CG14989*, which shares a 22-nucleotide stretch of uninterrupted complementarity centered on miR-7 target site-c (**Fig. 3A, Fig. S6A**). As a control, we repeated the search allowing sense-strand matches. This doubled the pool of hits, but *CG14989* remained the only hit in either orientation with a significant E-value, and was among the top three for both conservation and sequence complexity (**Fig. S6B; Supp. Table 6**). Further analysis of the surrounding *CG14989* sequence revealed two additional complementary sequences to the 8mer miR-7 seed target sequence, without the extended flanking complementary bases present around miR-7 site-c. All three sites were contained in a 66-base region of the 2,622-base *CG14989* transcript (referred to as sites c′, b′, and a′, 5′ to 3′; **Fig. 3B**). We emphasize that these are matches to the miR-7 guide sequence, not miR-7 target sites as found in the *VGAT* 3′UTR. *CG14989* therefore has the potential to base-pair with the *VGAT* 3′UTR and compete for miR-7 target sites if expressed in the same cell. Importantly, each of the three sites is deeply conserved across drosophilid species (**Fig. S6C**).

**Figure 3.**
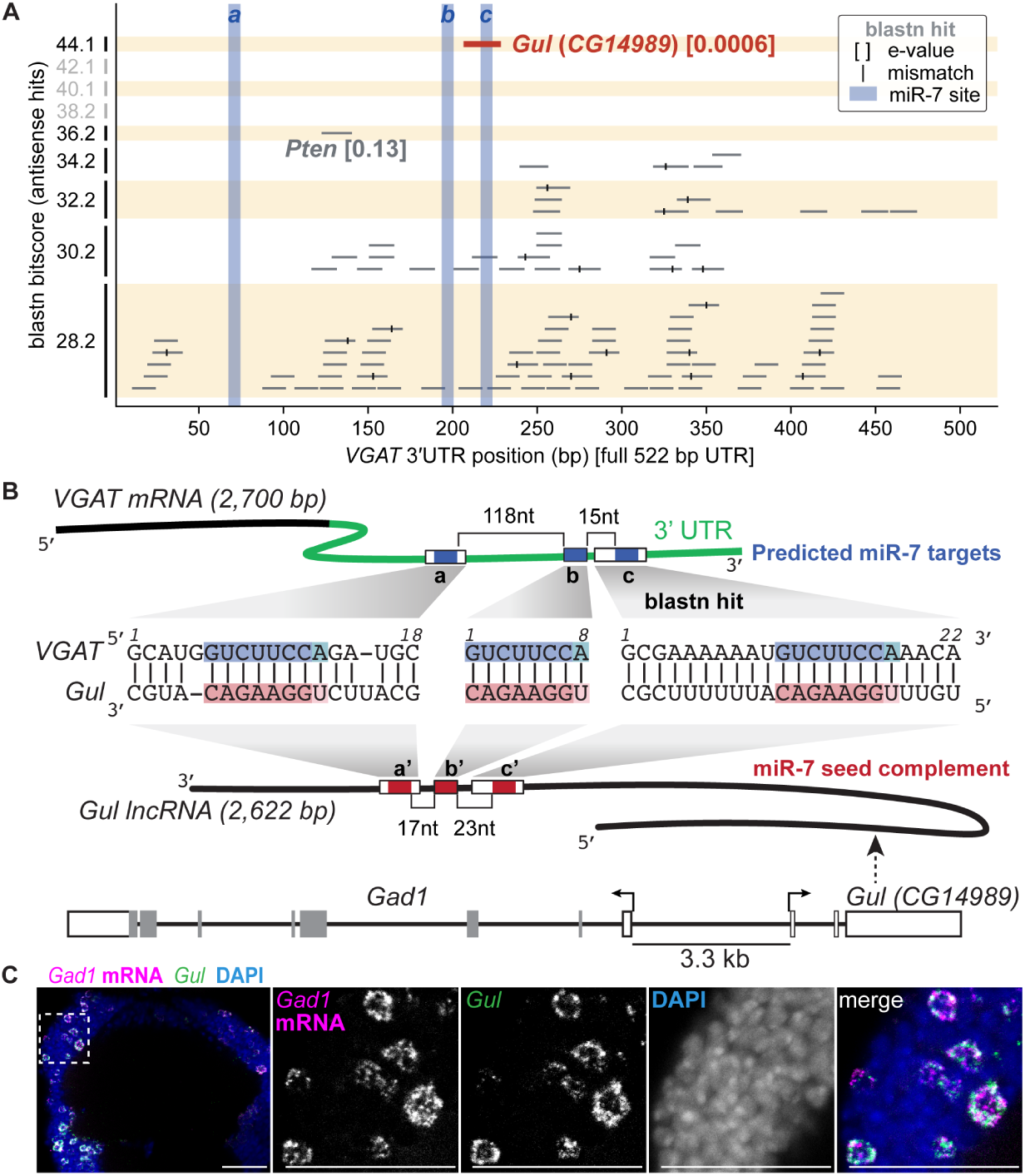
A GABAergic neuron-specific lncRNA carries sequences complementary to the *VGAT* miR-7 sites. **(A)** BLAST of the *VGAT* 3′UTR against annotated *Drosophila* transcripts, plotting each antisense hit at its position along the 522 bp 3′UTR against its blastn bit-score. Hits are grey and the miR-7 target sites are shaded blue; vertical ticks mark mismatches within a hit and bracketed values are E-values. The top hit, *Gul* (*CG14989*), is red. **(B)** Diagram of the *VGAT* mRNA and the *CG14989* transcript showing the predicted base-pairing at each of the three miR-7 guide matches (c′, b′ and a′, 5′ to 3′), the 66 nt window they occupy within the 2,622 nt transcript, and the divergent arrangement of *Gul* (*CG14989*) and *Gad1* at the locus. Predicted miR-7 target sites in *VGAT* are blue and the miR-7 seed complements in *CG14989* are red. **(C)** HCR RNA-FISH of the adult *Drosophila* optic lobe showing the distribution of *Gul* (green) relative to GABAergic neurons (magenta) marked by *Gad1* mRNA, with nuclei labeled by DAPI (blue). Scale bar: 20 μm. Inset: region of medulla cortex.

Strikingly, *CG14989* is positioned directly adjacent to, and transcribed divergently from, *Gad1*, which encodes the enzyme required for GABA synthesis and is transcribed only in GABAergic neurons. Analysis of scRNA-seq data from different brain regions ^36–38^ revealed that *CG14989* RNA shares the GABAergic neuron-restricted expression pattern of *Gad1* throughout the central nervous system (**Fig. S7A**), which we subsequently confirmed in the optic lobe using RNA in-situ hybridization (**Fig. 3C**), suggesting that the two genes share transcriptional regulatory elements. Indeed, of the 7,000-8,000 transcripts expressed in each brain region, *Gad1* and *CG14989* are the only two detected throughout the GABAergic population and almost nowhere else (present in 98-100% of GABAergic clusters and 0-6% of the remainder; **Fig. S7B**). The *CG14989* locus produces two isoforms: one expressed in the gut, which contains a long ORF with a conserved AUG codon, and one expressed specifically in neurons (**Fig. S8A; Supp. Table 7**). The neuronal isoform, which carries the *VGAT*-complementary region, is not present in the current genome annotation, but can be readily identified in bulk RNA-seq (**Fig. S8A**) and appears as a full-length transcript model in long-read sequencing of adult tissues ^61^. Unlike the gut isoform, this neuronal isoform does not contain a long ORF, with the longest theoretical one being 115 amino acids and beginning at a non-conserved AUG (**Fig. S8B**), and shows no evidence of translation by Ribo-seq or proteomics despite its high transcript abundance (**Fig. S8C**); this neuronal isoform is therefore likely to be a long non-coding RNA. Based on this expression pattern, and the functional analysis described below, we named *CG14989* as GABA UnLocker (*Gul*) and designate its gut and neuronal isoforms *Gul-G* and *Gul-N* respectively. The experiments that follow concern the neuronal isoform, and *Gul* refers to *Gul-N* from here on.

### Gul is necessary and sufficient for VGAT translation

Given the presence of three complementary regions to *VGAT* mRNA in *Gul* that overlap predicted miR-7 sites, and the expression of *Gul* exclusively in neurons where *VGAT* mRNA is translated into protein, we next tested whether *Gul* is necessary for *VGAT* translation in GABAergic neurons. We generated a deletion of the *Gul* gene body using CRISPR/Cas9, referred to as *ΔGul* **(Fig. S9A)**. This deletion is homozygous lethal, consistent with *VGAT* potentially being regulated by *Gul*, as *VGAT* expression is required for viability ^42^, but lethality could also be due to loss of the *Gul-G* isoform in the gut that is also deleted in this allele. To examine the expression of *VGAT* in neurons lacking *Gul*, we utilized MARCM clonal analysis ^62^ to generate homozygous *ΔGul* clones in an otherwise heterozygous animal. By co-staining *VGAT* and a GABAergic neuron subtype marker (Pm1/2/3 marked by the transcription factor Svp ^36,63^), we showed that loss of *Gul* leads to the absence of detectable VGAT protein in GABAergic neurons, confirming the necessity of *Gul* for *VGAT* expression **(Fig. 4A)**.

**Figure 4.**
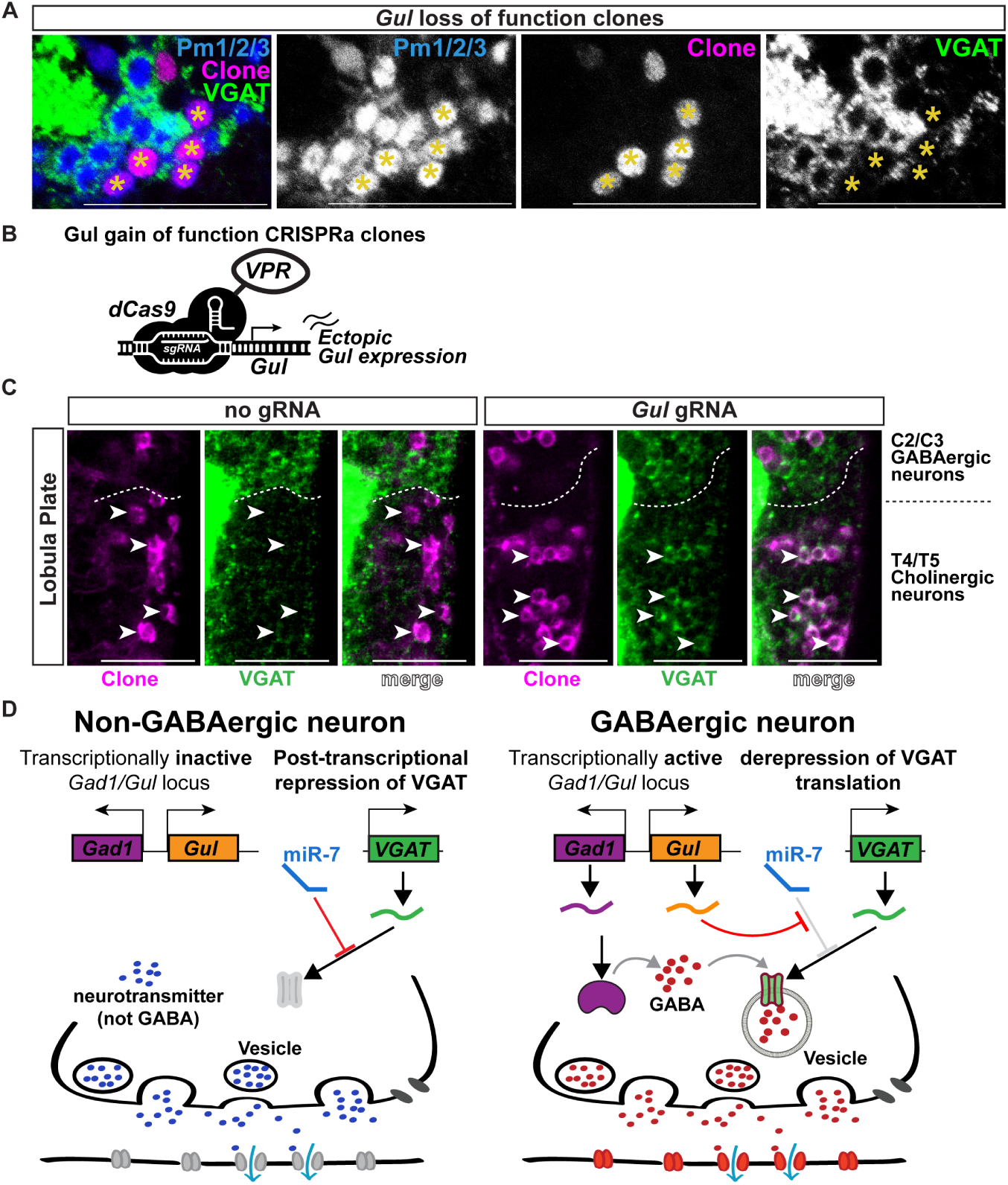
*Gul* is necessary and sufficient for *VGAT* translation. **(A)** Immunofluorescence of Δ*Gul* MARCM clones (magenta) in the optic lobe showing VGAT protein (green) relative to Pm1/2/3 neurons (blue) marked with a Svp antibody. Asterisks mark cells within a clone. **(B)** Diagram of the CRISPR activation system used to induce the endogenous *Gul* locus, in which a VPR activator fused to catalytically inactive Cas9 is targeted to the *Gul* promoter by ubiquitously expressed guide RNAs. **(C)** Immunofluorescence of *Gul* CRISPRa clones (magenta) in the lobula plate cortex showing VGAT protein (green), with the boundary between C2/C3 GABAergic neurons and T4/T5 cholinergic neurons marked. Arrowheads mark cells within a clone and the dashed line marks the boundary between the two populations. Clones generated without a guide RNA are shown as a control. **(D)** Model of *VGAT* regulation in non-GABAergic and GABAergic neurons. Scale bar: 20 μm.

We next asked whether non-GABAergic neurons, which transcribe *VGAT* mRNA but repress it through miR-7, are competent to express VGAT protein when *Gul* is supplied. We ectopically activated transcription of the endogenous *Gul* locus using CRISPR activation (CRISPRa), in which a transcriptional activator fused to catalytically inactive Cas9 is induced through the Gal4/UAS system and is targeted to the *Gul* promoter by sequence-specific guide RNAs (**Fig. 4B**) ^64,65^. Stochastic clones were generated throughout the visual system by combining UAS-CRISPRa with a heat-shock-inducible tubulin>FRT-STOP-FRT.Gal4 driver. We investigated *VGAT* protein levels in two of the non-GABAergic populations introduced above: the lamina cortex ^39^ and the T4/T5 region of the lobula plate cortex ^40,41^. We found widespread induction of *VGAT* protein within CRISPRa clones in both lobula plate neurons **(Fig. 4C)** and lamina neurons **(Fig. S9B)**, demonstrating that *Gul* is sufficient to derepress the translation of *VGAT* mRNA in non-GABAergic neurons. Notably, the levels of *VGAT* induction in the lobula plate T4/T5 neurons were comparable to an adjacent population of C2/C3 neurons, a naturally GABAergic population that expresses *Gul* and *VGAT*.

Together, these results establish that the GABAergic neuron-specific lncRNA *Gul* is both necessary and sufficient for *VGAT* translation in *Drosophila*, identifying it as the determinant of cell type specificity of *VGAT* expression. These results also resolve the alternative raised earlier, that GABAergic neurons might simply carry less miR-7 activity. Both manipulations are clonal and cell-autonomous, made in an otherwise wild-type background, so miR-7 is unaltered in either. Removing *Gul* abolishes *VGAT* protein in GABAergic neurons that retain whatever miR-7 activity those cells normally have, and supplying *Gul* induces *VGAT* protein in non-GABAergic neurons in which miR-7 remains present and active. A difference in miR-7 availability is therefore neither necessary nor sufficient to explain where *VGAT* is translated.

### Gul complementarity extends to multiple GABA-pathway mRNAs and biases their expression toward GABAergic neurons

If *Gul* relieves miR-7 repression by pairing with the *VGAT* 3′UTR, the same mechanism should apply to other transcripts it can pair with. We took two approaches to identify transcripts that *Gul* might act on along with *VGAT*. The first used sequence alone, in two BLAST searches against annotated 3′UTRs (**Fig. 5A** and **Fig. S10A**). Searching a 110 bp region carrying the three mimic sites returned four transcripts whose complementarity spans a complete miR-7 8mer: *VGAT*, the best hit in the transcriptome with 22 nt, and *sgll*, *ScsbetaG*, and *stol* with 15, 17, and 15 nt. Searching a larger ∼2 kb region surrounding the three mimic sites in the *Gul* transcript recovered three more transcripts that pair outside the mimic sites of *Gul* and carry no predicted miR-7 site: *P5CDh1*, *Gabat*, and *Glts*, over 28, 23, and 16 nt with no more than two mismatches. Across the seven transcripts, conservation of the matched site was well above background of all blast hits (median 0.89 against 0.26; Supp. Table 8), and all are compositionally complex, unlike the AT-rich repeats that account for much of the background (**Fig. S6B, S10B, S11)**. The four miR-7 targets are the only genes containing conserved 8mer targets in the genome with at least 7 nt of flanking complementarity to a *Gul* mimic site.

**Figure 5.**
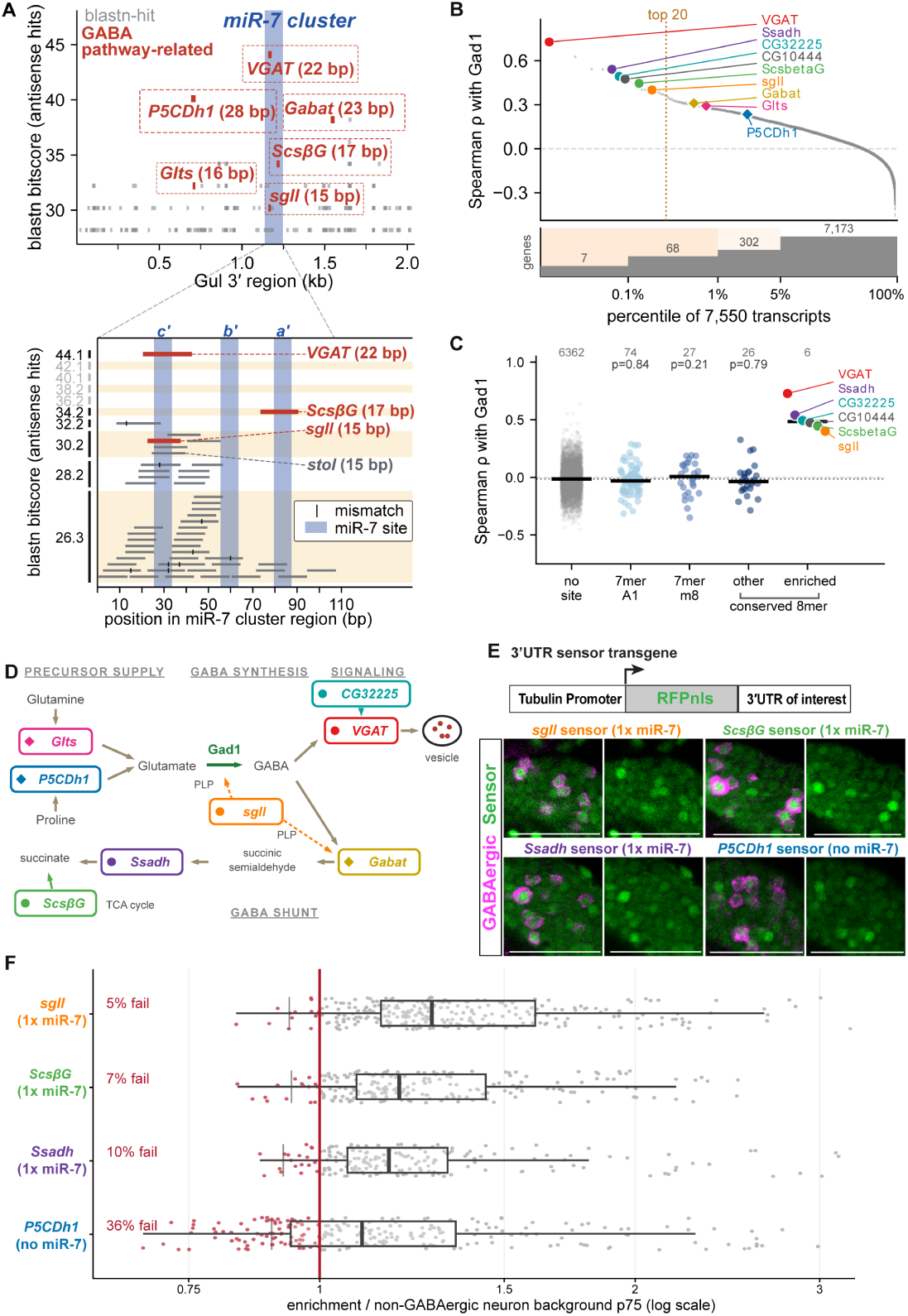
*Gul* complementarity extends across the GABA pathway. **(A)** BLAST of the *Gul* transcript against annotated 3′UTRs, plotting each hit at its position along the *Gul* 3′ region against its blastn bit-score, with hits in grey, the miR-7 sites shaded blue, and vertical ticks marking mismatches within a hit. GABA-pathway transcripts are red and labeled with the length of their match. The lower panel expands the miR-7 cluster region, showing the three mimic sites (c′, b′ and a′) and the hits that overlap them. **(B)** Every analyzed transcript ranked by its mean Spearman ρ with *Gad1* in the optic lobe, plotted against percentile. The dotted line marks the top 20, and the strip beneath gives the number of transcripts in each band. The nine candidate transcripts are labeled and colored. **(C)** Mean Spearman ρ with *Gad1* in the optic lobe for transcripts grouped by miR-7 site class, with the six enriched transcripts drawn as a separate column and the bracket marking that the last two columns are both conserved 8mers. Bars are class medians and n is given above each column, and the six are colored individually as in (B). **(D)** Diagram of the GABA pathway showing the transcripts recovered by the sequence and co-variation screens, colored as in (B). **(E)** Diagram of the 3′UTR sensor transgenes, consisting of a tubulin promoter driven RFPnls with the 3′UTR of interest, and their activity (green) in a subset of the optic lobe medulla cortex of wild type flies relative to GABAergic neurons (magenta) marked with a Gad1 antibody. **(F)** Per-GABAergic neuron sensor activity for each 3′UTR quantified in the entire optic lobe medulla cortex, expressed as enrichment over the local non-GABAergic background of the same image. The red line marks the 75^th^ percentile of the relative output of the non-GABAergic background; neurons at or below it are red and their percentage is given for each sensor. Scale bar: 20 μm.

The second approach ranked all transcripts by co-variation with the GABAergic marker *Gad1* across cluster-mean expression in the three regions of the adult CNS (optic lobe, ventral nerve cord, and central brain), restricted to the 7,550-10,728 transcripts detected in at least a quarter of clusters (**Fig. S12**). The two approaches share no input: one reads *Gul* sequence and takes no account of expression, the other reads GABAergic expression and takes no account of *Gul*. Because *Gul* and *Gad1* are themselves near-indistinguishable in expression (ρ = 0.88 to 0.93 across the three regions), a ranking anchored on *Gad1* cannot attribute co-variation to *Gul* specifically; it identifies transcripts that track GABAergic identity, of which *Gul* is one.

Strikingly, transcripts carrying a conserved 8mer miR-7 site are 0.4% of the expressed transcript group, but supply six of the twenty most strongly *Gad1*-correlated transcripts in every CNS region, a 65-to 81-fold enrichment (hypergeometric p < 10^-9^; **Fig. 5B; Supp. Table 9**), even though the conserved 8mer class as a whole shows no such enrichment (**Fig. 5C**), and it is the same six genes in optic lobe, ventral nerve cord, and central brain: *VGAT*, *Ssadh*, *CG32225*, *sgll*, *ScsbetaG*, and *CG10444*. Ranked among the 33 conserved 8mer targets detected in all three regions by co-variation with *Gad1*, these six take the first six places (mean ρ +0.72 to +0.48) ahead of a 1.8-fold gap to the seventh miR-7 target transcript (+0.26) (**Fig. 5C**). *VGAT* aside, two of the remaining five, *sgll* and *ScsbetaG*, had already been recovered by the sequence search (15/15 and 17/17 nt, centered on their miR-7 sites), which took no account of expression; the other three emerged only in the co-variation analysis, but their individual miR-7 8mer sites are complementary to the three consecutive mimic copies in the *Gul* sequence. The transcripts recovered by complementarity alone but carrying no miR-7 site, *P5CDh1*, *Glts*, and *Gabat*, were themselves strongly *Gad1*-correlated, ranking within the top 2.5% of transcripts in every CNS region (**Fig. 5B** and **Fig. S12A**).

Eight of the ten transcripts recovered by the two screens form a functional module related to GABA metabolism and signaling (**Fig. 5D**). VGAT loads GABA into synaptic vesicles, and *CG32225* is the *Drosophila* ortholog of LAMP5, which traffics VGAT. *sgll* encodes pyridox(am)ine 5′-phosphate oxidase (human PNPO), which produces pyridoxal 5′-phosphate, the cofactor required by both Gad1 for GABA synthesis and Gabat for its degradation. *Gabat*, also recovered here, encodes the transaminase (human ABAT) that initiates GABA degradation by converting GABA to succinic semialdehyde, and *Ssadh* encodes succinic semialdehyde dehydrogenase, which acts on that substrate as the terminal enzyme of the GABA shunt (the pathway that degrades GABA to succinate and returns it to the tricarboxylic acid cycle); both consecutive steps of the shunt are therefore represented in the set. *ScsbetaG* encodes the β subunit of succinyl-CoA synthetase (human SUCLG2), the canonical tricarboxylic acid cycle step that produces succinate, the metabolite on which the GABA shunt reconverges. *P5CDh1* and *Glts* both supply glutamate, the substrate for Gad1, as pyrroline-5-carboxylate dehydrogenase (human ALDH4A1) and glutamate synthetase respectively; notably, their *Gul*-complementary sites coincide, the *Glts* match lying entirely within the *P5CDh1* match, so the two transcripts are complementary to the same position on *Gul*. The remaining two are not part of this module: *CG10444* encodes an SLC5-family sodium:solute symporter, and *stol* a calcium-channel α2δ subunit without a known specific association to GABAergic neurons. The set therefore spans glutamate supply, cofactor production, vesicular loading and trafficking, and the two-step catabolic shunt, rather than representing an arbitrary collection of mRNAs (**Fig. 5D**).

A caveat to the analysis of gene expression from RNA-seq data is that transcriptional and post-transcriptional contributions cannot be disentangled when measuring steady state mRNA levels. To test whether these relationships reflect post-transcriptional regulation, we generated 3′UTR sensors (**Fig. 5E**) for *sgll*, *ScsbetaG*, *Ssadh*, and *P5CDh1*. Because the sensors are driven by the same tubulin promoter and differ only in their 3′UTR, any difference in protein output between neuronal populations is post-transcriptional in origin. The three sensors carrying a miR-7 site each drove higher protein output in GABAergic than in non-GABAergic neurons, with *sgll* the most enriched, whereas the *P5CDh1* sensor, which carries extensive *Gul* complementarity but no predicted miR-7 site, showed only a modest bias (**Fig. 5F**). Comparing each GABAergic neuron in the medulla region of the optic lobe with its non-GABAergic neighbors, the *P5CDh1* sensor failed to exceed them in 36% of GABAergic neurons, against 5-10% for the three sensors carrying a miR-7 site. It is therefore *Gul* complementarity overlapping a miR-7 site that confers a reliable GABAergic bias.

Across the 33 conserved 8mer targets, two features of the 3′UTR together accounted for roughly a third of the variance in whether a transcript registered a GABAergic difference: the number of other conserved miRNA families with sites in the same 3′UTR, which sets how large a share of that transcript’s miRNA input runs through the miR-7 site, and the length of uninterrupted complementarity to *Gul* across that site (**Fig. S13; Supp. Table 10**). *stol* and *sgll* illustrate the first: they carry the same single conserved 8mer and the same 15 bp of complementarity to *Gul*, but *stol* sits in a 3′UTR with thirty other conserved families where *sgll* has none, and only the latter registered a GABAergic difference. These features indicate which transcripts are susceptible to regulation through their miR-7 site, not what acts on that site; the evidence identifying *Gul* as the agent is the loss-and gain-of-function experiments above. Whether *Gul* can compete at these sites, particularly against miR-7/Argonaute complexes that bind their target mRNAs efficiently, may also depend on how much of it is present. Amongst the 12,000-16,000 transcripts detected in each CNS region, *Gul* ranks 2nd to 12th in abundance and carries three consecutive copies of the miR-7 5′ end, placing its mimic sites in approximately 70-to 100-fold excess over the sites they compete for (**Fig. S14; Supp. Table 11**). At this stoichiometry a minimal duplex may be sufficient to compete with miR-7 for binding.

Together, these results support a model in which the GABAergic neuron-specific lncRNA *Gul* post-transcriptionally regulates a functionally related set of transcripts, recovered both by complementarity to *Gul* and by co-variation of their expression with GABAergic identity, spanning glutamate supply, cofactor production, vesicular loading, and GABA breakdown.

## Discussion

Our findings reveal a novel post-transcriptional regulatory mechanism in which a lncRNA antagonizes miRNA-mediated repression to achieve cell type-specific expression of a target gene.

We demonstrate that the GABAergic neuron-specific lncRNA *Gul* is necessary and sufficient for the derepression of *VGAT*, which is transcribed pan-neuronally but is otherwise strongly silenced by miR-7 outside of the GABAergic population. This mechanism of lncRNA-mediated miRNA antagonism appears to extend beyond *VGAT*: *Gul* bears complementarity to mRNAs spanning nearly every aspect of GABA metabolism and signaling, and it is those also predicted to be miR-7 targets whose protein output is preferentially regulated, establishing *Gul* as an antagonist of miR-7-mediated repression across the GABA pathway. *Gul* is transcribed divergently from a shared regulatory region at the *Gad1* locus, so a single transcriptional input coordinates multiple essential components of GABA signaling in the correct neuronal context.

This mechanism also represents a distinct strategy for achieving cell type-specific regulation of miRNA activity. miR-7 is similarly active in both GABAergic and non-GABAergic neurons, and therefore cell type-specific derepression of *VGAT* is not achieved by regulating miR-7 expression or its general activity, as in the case of target-directed miRNA degradation ^66^ or miRNA sponges like ciRS-7 ^34^. Instead, a cell can deploy a competing transcript that acts locally, at the level of individual target sites, to achieve target-and cell-type-specific derepression, decoupling the regulatory output of a broadly active miRNA from its spatial (cell type) expression pattern. This strategy may depend on specific characteristics of *Gul*: it is a competitor of high abundance carrying three copies of the miR-7 5′ end, and a transcript of lower abundance, or one carrying a single copy, would not be expected to compete as effectively at the same degree of complementarity.

The organization of the *Gad1/Gul-VGAT* regulatory network, and potentially that of other GABA-related genes, is reminiscent of the logic of prokaryotic operons, in which a single regulatory input drives the coordinated expression of functionally related genes. In bacteria this is achieved through polycistronic transcription of pathway components from a shared promoter, and in eukaryotes, such coordination often comes from co-regulation of genes by the same transcription factor. Our findings suggest a third route: the GABAergic transcriptional program drives expression of both genes at the *Gad1/Gul* locus, and while *Gad1* mRNA is directly translated into the enzyme required for GABA synthesis, *Gul* acts *in trans* to relieve miR-7-mediated repression of multiple functionally related genes at distinct loci that are transcribed broadly in the nervous system. The *Gad1/Gul* locus is therefore a functional analog of an operon, co-regulating these dispersed pathway genes without the co-transcription that links the genes of a true operon. Our findings demonstrate that this strategy is an essential regulatory step for GABAergic neuronal signaling in *Drosophila* and suggest it may represent a general mechanism for coordinating pathway components across distinct loci in eukaryotes.

Coordination of this kind does not require every target to respond to the same degree. The GABAergic neuron-specific regulation we measure for the other targets of *Gul* besides *VGAT*, both in mRNA levels from scRNA-seq and using the transgenic 3′UTR sensors, varies quantitatively and is smaller than that of the *VGAT* sensor, which is close to fully repressed outside the GABAergic population (**Fig. 1G**). *VGAT* carries three conserved 8mer sites where each of the other transcripts carries only one; two of the three *VGAT* sites lie 15 nt apart, within the spacing over which neighboring miRNA sites act cooperatively, and the third lies 118 nt away and is expected to act independently. A near-binary response for *VGAT* alongside graded responses for single-site transcripts is therefore the expected consequence of site number and arrangement, rather than evidence of a different mechanism. *VGAT* is also likely the only one of these transcripts whose protein product is needed exclusively in GABAergic neurons; the others encode enzymes of cofactor supply, GABA catabolism, and the tricarboxylic acid cycle that are also required in other neurons and in glia, so a tuning of level rather than a switch may be all that is required at these transcripts. Future experiments will address whether *Gul* influences aspects of RNA regulation beyond the mRNA stability and translation measured here, such as subcellular localization.

Consistent with this regulatory strategy operating beyond GABA metabolism, we note that a similar arrangement has been described among the *Drosophila* neurogenic genes. The 3′UTRs of several of the Notch target genes in the *Enhancer of split* and *Bearded* complexes carry a miR-7 site while the 3′UTRs of the proneural genes *achaete*, *lethal of scute*, and *atonal* carry a conserved ‘proneural box’ that contains the 5′ end of miR-7; the two motifs are exactly complementary and are predicted to pair over extended regions, reaching 18 uninterrupted bases between *Bearded* and *atonal* ^67^. These Notch targets are directly activated by the proneural proteins^68^ as well as by Notch ^69,70^, and among them the Bearded family proteins compete with Delta for binding to Neuralized, reducing Delta signaling from the cells in which they are expressed ^71–73^. Because both sets of genes are expressed in the same proneural cluster cells, complementary base pairing between their transcripts could perform a coordinating role like the one we describe for *Gul* and *VGAT*; theirs between two mRNAs rather than between a lncRNA and an mRNA. Coupling the expression of functionally related genes through complementary sequences that overlap shared miRNA target sites may therefore be a broader feature of post-transcriptional gene regulation in *Drosophila*.

*CG14989/Gul* is present in all insects with conserved synteny and orientation relative to *Gad1* **(Fig. S7C)**. Furthermore, a lncRNA in the same relative genomic position (adjacent to and transcribed divergently from *Gad1*) is a common feature across animals including humans, even in the absence of detectable sequence homology with *Gul*. This lack of sequence conservation is consistent with the general observation that lncRNAs tend to evolve more rapidly than protein-coding genes at the sequence level while maintaining positional and functional conservation ^74,75^. The human *GAD1*-adjacent lncRNA, *GAD1-AS1*, is significantly correlated with *GAD1* expression, mirroring the relationship we describe between *Gul* and *Gad1* in *Drosophila* ^26^. We did not, however, detect significant complementary bases between *GAD1-AS1* and *VGAT* or other GABA-related mRNAs in human. Whether *GAD1-AS1* performs an analogous coordinating function in human GABAergic neurons, and whether it does so through a mechanism involving mRNA interaction, represents an important question for future investigation.

Beyond the mechanistic novelty of *Gul*-mediated miRNA antagonism, our findings suggest that *Gul* may represent a previously unrecognized type of lncRNA defined by its highly specialized role in establishing and maintaining the molecular identity of a specific class of cell (*e.g.* neurotransmitter identity). It is well established that many lncRNAs are transcribed in close proximity to protein-coding genes, and that their expression is frequently correlated with that of their neighboring gene ^26^. However, in most cases the functional significance of this relationship remains unclear, and the best characterized cases are lncRNAs that act within the nucleus to regulate chromatin and transcriptional state of neighboring genes *in cis* ^30^. Our findings raise the possibility that at least some of these lncRNAs may serve a dedicated role in enforcing the molecular program required by their neighboring coding gene by acting *in trans* in the cytoplasm to regulate the expression of specific functionally-related transcripts: in the case of *Gul*, it not only shares the exclusive GABAergic expression pattern of *Gad1*, but bears complementarity to the mRNAs of the metabolic pathway of GABA produced by Gad1. To our knowledge, a role for lncRNAs in reinforcing the molecular identity of a specific cell type by coordinating the expression of functionally related mRNAs has not been previously documented, and we suggest that systematically examining the sequences and targets of lncRNAs co-expressed with key cell type-defining genes may reveal additional examples of this regulatory logic.

In summary, our findings reveal a mechanism for cell type-specific post-transcriptional control of gene expression, in which a lncRNA pairs with the 3′UTRs of functionally related mRNAs and preferentially derepresses those that are predicted miR-7 targets. Because *Gul* is co-regulated with *Gad1*, a single transcriptional input is propagated through these 3′UTR interactions to genes at dispersed loci. *Gul* therefore acts as an operon-like coordinator of the GABA pathway in *Drosophila*, and we suggest that this represents a conserved strategy for achieving coordinated gene expression in a cell type-specific manner in eukaryotes.

## Limitations of the study

Our data establish that Gul is necessary and sufficient for VGAT translation and that it bears complementarity to the 3′UTRs of additional GABA-pathway transcripts, but several questions remain open. We have not visualized the Gul-target RNA duplex directly in vivo, and the precise step at which base-pairing relieves miR-7 repression, whether by competition for the shared site, a change in target-site accessibility, or recruitment of additional factors, remains to be defined. The extended target set is supported by complementarity and by co-variation with GABAergic identity, with functional validation for a subset; further targets, or interactions with transcripts we did not test, may exist. Finally, we measured mRNA abundance and protein output but did not assess whether Gul influences other aspects of RNA regulation, such as subcellular localization, and whether an analogous lncRNA-mediated mechanism operates in other cell types or species remains to be tested.

## Materials and Methods

### EXPERIMENTAL METHODS

#### Fly strains and fly rearing

Flies were reared on molasses-cornmeal-agar media at 25°C. The fly lines used are described in Supp. Table 1. *VGAT* 3′UTR sensor plasmids were injected by BestGene Inc. or WellGenetics in the 3rd chromosome (attP2 landing sites). For MARCM experiments, L1-L2 larvae were heat shocked for one hour at 38°C, and then reared at 25°C until flies were dissected in the adult. In all experiments at least 3 different brains were analyzed.

#### Immunohistochemistry

Flies were anesthetized on ice and brains dissected in ice-cold Schneider’s media and then transferred to Schneider’s media on ice. Brains were fixed using 4% paraformaldehyde diluted in PBS 1X for 20 min at room temperature. Then, they were rinsed 3 times in 0.5% Triton-X diluted in PBS 1X (PBTx), washed for 30 min in PBTx, and blocked for 30 min in PBTx with 5% goat serum (PBTx-block) at room temperature. They were then incubated at 4°C for 2 overnights in primary antibodies diluted in PBTx-block. They were then rinsed 3 times in PBTx, incubated two times for 30 min in PBTx, and incubated at 4°C overnight in secondary antibodies diluted in PBTx-block with 5 µg/mL DAPI. Finally, they were rinsed 3 times in PBTx, incubated two times for 30 min in PBTx, and mounted in Slowfade before imaging with either a Leica SP8 or Leica Stellaris confocal microscope using a 63x glycerol objective. Image stacks were acquired at 0.8-1 µm optical sections, and 0.5 µm sections for sensors that underwent segmentation using Cellpose (below) and quantification. Images were processed in Fiji and Adobe Illustrator. GABAergic neurons in miR-7 null brains (Fig. 2E) were identified by immunostaining with a rabbit anti-GABA antibody. The antibodies used are described in Supp. Table 1.

#### Hybridization Chain Reaction Fluorescence In Situ Hybridization (HCR-FISH) v3.0

Flies were anesthetized on ice and brains dissected in ice-cold PBS and then transferred to PBS on ice. Brains were fixed using 4% paraformaldehyde diluted in PBTx for 20 min at room temperature, and processed using the HCR-FISH v3.0 protocol as described previously ^76^. Samples were washed three times in PBTx for 15 min. They were then incubated at 37°C for 30 min in Hybridization solution (Molecular Instruments) with shaking. Following pre-hybridization, the brains were incubated in hybridization solution with RNA-specific probes (diluted 1:100) at 37°C overnight with shaking. The following day, brains were washed four times in Probe Wash Buffer (Molecular Instruments) at 37°C for 15 min on a rocker, followed by two 10 minute washes in 0.1% Tween-20 diluted in 5x sodium chloride sodium citrate (5XSSCt). They were then incubated for 30 min at room temperature in Amplification Buffer (Molecular Instruments) with rocking, followed by Amplification Buffer with fluorescently coupled amplifiers synthesized by Molecular Instruments (diluted 1:50) and incubated at 25°C for 4 hours in the dark. Next, they were washed 5 times for 15 min in 5XSSCt. Brains were then incubated in 5XSSCt containing 5 µg/mL DAPI before two final washes in PBTx for 30 min and mounting in SlowFade medium. The *VGAT* HCR-probe set was synthesized by Molecular Instruments. *Gad1* and *CG14989*/*Gul* probe sets were designed using HCR Probe Maker ^77^ with modifications (https://github.com/chenyenchung/insitu_probe_generator) and ordered as oligo pools from IDT. All buffers were made with DEPC-treated RNase-free water. Sequences of *Gad1* and *Gul* probes are listed in Supp. Table 2.

#### Antibody generation

A Guinea pig polyclonal antibody against Gad1 was generated by Genscript using the full Gad1 protein as antigen.

#### Molecular Biology

To generate the *VGAT* 3′UTR sensors, we first modified pJFRC177-10XUAS-FRT>-dSTOP-FRT>-myr::GFP (Addgene #32149) to make tublin>2xmsGFP2-PEST-NLS (Plasmid RLV72), which contains the *Drosophila* αTub84B promoter upstream of tandem copies of msGFP2 ^78^ with three copies of the SV40 NLS and a PEST domain, followed by an SV40 terminator flanked by SapI restriction sites. Wild-type and mutant versions of the *VGAT* 3′UTR were synthesized by Genewiz (Azenta) and inserted via Golden Gate SapI assembly. The plasmid sequences can be found in Supp. Table 2.

We also generated an alternative red sensor plasmid by replacing the 2xMsGFP2 with tandem copies of mScarlet-I3 ^79^ (RLV183). To generate the *sgll*, *ScsbetaG*, *Ssadh*, and *P5CDh1* 3′UTR sensors, the respective 3′UTR sequences were synthesized by Genewiz (Azenta) and inserted into RLV183 as described above via Golden Gate SapI assembly. The miR-7 sensor was also generated in RLV183, by inserting a synthesized fragment in which an SV40 terminator lies downstream of two complete 23-nucleotide complements to the mature miR-7-5p sequence, separated by 25 nucleotides.

#### CRISPR/Cas9-mediated deletion of *Gul*

To generate the Δ*Gul* deletion allele, guide RNAs targeting sequences flanking the *Gul* gene body were designed using the CRISPR optimal target finder and EvaluateCrispr tool from the *Drosophila* TRiP group and cloned into the pCFD5 gRNA expression plasmid ^80^. The donor plasmid was assembled using the pTV-Cherry plasmid (DGRC#1338) ^81^, containing an attP site and loxP-flanked 3XP3-mCherry selectable marker. Homology arms flanking the gRNA cut sites were synthesized as 200 bp fragments with a gRNA site (identical to one of the genomic gRNA sites) flanking to permit in-vivo linearization prior to homology-directed repair ^82^ and inserted using Infusion homology-based cloning. Guide RNA and homology arm sequences are listed in Supp. Table 2. Cas9 and guide RNA constructs were co-injected into a nos-Cas9 strain expressing a germline-specific Cas9 by WellGenetics. Surviving G0 flies were crossed to a balancer stock and individual G1 progeny were screened by mCherry fluorescence and subsequent PCR and Sanger sequencing to confirm the deletion boundaries by WellGenetics.

#### Generation of endogenous *VGAT* alleles

Endogenous *VGAT* alleles, including the smV5-T2A-*VGAT* wild type reporter and the smV5-T2A-*VGAT* 3X-miR-7 target site deletion, were generated by Recombinase-Mediated Cassette Exchange (RMCE) using a previously generated attP-flanked deletion of the *VGAT* locus ^83^. An initial RMCE Donor plasmid base (RLV154) was designed with the following: the desired *VGAT* genomic sequence covered in the initial deletion, flanked by attB sites, with a nuclear-localized spaghetti monster V5 (smV5) reporter fused in-frame to the N-terminus of *VGAT* coding sequence followed by T2A self-cleaving peptide was synthesized by Genewiz. RLV154 also contains a cloning cassette that permits insertion of modified 3′UTR sequence covering the miR-7-containing region. Wild type (RLV155) or miR-7 mutant (RLV157) donors were generated by inserting synthesized 3′UTR fragments into RLV154 using Infusion cloning. Note that RLV157 also contains a synonymous mutation to alter an additional (fourth) predicted miR-7 target site that resides in the coding region of *VGAT* adjacent to the endogenous STOP codon. RMCE plasmids were injected by BestGene Inc. into the *VGAT*-attP strain, screened by loss of red eye fluorescence, and Sanger sequenced to confirm successful integration and correct orientation. The plasmid sequences can be found in Supp. Table 2.

#### CRISPR activation (CRISPRa) of *Gul*

Ectopic activation of *Gul* in neurons was achieved using the CRISPRa system ^64,65^. A stable strain expressing Guide RNAs targeting the *Gul* promoter region were designed and generated by the TRiP group (BDSC#81666 ^65^). CRISPRa clones were induced by crossing flies containing a heat-shock-inducible tubulin>FRT-STOP-FRT.Gal4, UAS-dCas9-VPR, and a UAS-myr.tdTomato reporter to a line carrying the *Gul* promoter-targeting gRNAs. Pupae at ∼48 h APF were heat shocked at 38°C for 15 min to induce Flp-mediated removal of the STOP cassette, thereby activating Gal4 stochastically in individual cells. Animals were dissected one day after adult eclosion. VGAT protein levels were assessed by immunofluorescence as described above.

### COMPUTATIONAL ANALYSIS

#### Single-cell RNA-seq data analysis

Existing single-cell RNA-seq datasets from the *Drosophila* optic lobe ^36^ and ventral nerve cord and full brain regions ^37,38^ were used to assess cell type-specific expression of *VGAT*, *Gad1*, and *Gul*. Expression data was analyzed using Seurat ^84^ and visualized using a custom R script.

The RNA-binding protein screen was run across three regions of the adult CNS, on the same cluster-mean matrices described above, with glia excluded and GABAergic clusters defined by an Otsu split of the log *Gad1* distribution, as described below. The RNA-binding protein gene list was taken from the Gene Ontology term GO:0003723 (RNA binding) together with its molecular-function descendant terms, from the FlyBase GO annotation file and the GO Consortium ontology, and restricted to protein-coding genes using the FlyBase r6.68 annotation, giving 696 genes. Genes were retained if their mean expression across clusters exceeded 0.02 and their 90th-percentile expression exceeded 0.05. Two readouts were computed. For detection, a gene was called present in a cluster when its mean exceeded 10% of its own 90th-percentile value, GABAergic-restricted when present in at least 90% of GABAergic clusters and no more than 20% of the remainder, and GABAergic-excluded when the converse held. For level, genes were ranked by the log2 ratio of mean expression in GABAergic against other clusters.

#### *VGAT* expression by neuronal class

Fold differences in *VGAT* between GABAergic, non-GABAergic, and glial clusters were computed on the mean of log-normalized expression per cluster. Clusters were assigned to neuronal classes by neurotransmitter marker expression, and three sets were excluded before analysis: clusters meeting no confident class call, clusters over-called as glial, and clusters called GABAergic by the threshold but with *Gad1* detected in fewer than half their cells. That last exclusion matters because the threshold separates clusters containing GABAergic cells from those containing none, rather than identifying clusters that are GABAergic throughout: mixed clusters have a normal per-cell *Gad1* level with roughly a third of the detection rate, so their mean represents neither population. Group differences were tested by one-sided Mann-Whitney.

For this comparison GABAergic clusters were defined by an Otsu cut on the linear cluster mean of *Gad1*, which is not the definition used for the co-variation analyses described below, where the cut is taken on log *Gad1*. The two are separated deliberately: the linear cut is the more inclusive criterion and is appropriate for asking whether a class contains GABAergic cells at all, while the log cut is the stricter one used wherever GABAergic and non-GABAergic clusters are contrasted quantitatively.

#### 3′UTR sequence and miR-7 target annotation

3′UTR sequences were taken from the FlyBase r6.58 3′UTR set, using the longest annotated isoform per gene. Gene symbols were canonicalized against FlyBase r6.68 before analysis. Predicted miR-7 target genes and their site classes were taken from the TargetScanFly 7.2 per-gene conserved-target export for dme-miR-7-5p (seed GGAAGAC, species 7227), which lists 183 genes; each gene was assigned its best site class in the order 8mer > 7mer-m8 > 7mer-A1.

The number of conserved miR-7 sites per gene was taken from the TargetScanFly 7.2 per-gene predicted-target export for miR-7-5p (183 genes), using TargetScan’s own conserved and poorly-conserved calls, which are based on the probability of preferential conservation. Requiring preferential conservation rather than presence in the alignment alone is what makes the maximum site count unique to one gene, and site counts should be quoted against that criterion.

#### BLAST searches

A BLAST database was built from 30,283 FlyBase 3′UTR sequences (makeblastdb,-dbtype nucl). Three queries were searched against it with blastn-task blastn-short-evalue 1000-word_size 7-max_target_seqs 40000: the 522 bp *VGAT* 3′UTR; the full 2,045 bp *Gul* 3′ region; and a 110 bp slice of that region (positions 1,138 to 1,247) containing the three miR-7 mimic sites. Low-complexity sequence was removed with dustmasker at the default level of 20, applied as a hard per-hit interval filter rather than as query soft-masking; without it, AT-repeats and poly-A/T tracts within the *Gul* region seed large numbers of spurious short full-identity matches. Self-hits were excluded (the *VGAT* query against FBtr0087577, the *Gul* query against the four *Gul* isoforms), and where a gene contributed several transcripts only its highest-scoring hit was retained. Hits were ranked separately by strand; the antisense set, which is the base-pairing configuration the model predicts, is what is reported, with the sense set retained as a control. The three searches return 985, 995, and 500 genes respectively. Because blastn-short assigns bit-scores from a small set of discrete values, most hits fall into large ties within which rank is arbitrary, so tie-block membership rather than rank is quoted throughout. The A+T fraction of every matched segment was recorded alongside its bit-score, because the 3′UTR database is 62.9% A+T; no composition correction was applied and nothing was filtered or ranked on it.

Positional panels plot each gene’s best full-3′UTR hit against its position along the query, for the top 100 hits per strand, on a categorical bit-score axis with antisense and sense hits distinguished. The conservation-complexity panels plot two z-scored measures of each matched segment against each other: conservation as the drosophilid Branch Length Score, and complexity as the A+T fraction with its sign reversed. Both are standardized against a pooled reference combining the antisense background and the sense-strand control, so that the two strands sit on one common frame rather than each being centered on its own mean.

#### Conservation and sequence complexity

Two conservation measures were computed for each matched segment, both on the target gene’s own 3′UTR. Breadth across the genus was measured as a Branch Length Score, the fraction of total phylogenetic branch length spanned by species retaining the site, computed on the UCSC 27-way insect alignment restricted to its 23 *Drosophila* species, with a species counted as retaining the site at 80% or greater identity to dm6. Depth toward distant insects was measured as the mean phyloP score over the matched segment on the 124-way insect alignment. Sequence complexity was scored on the matched segment as the triplet-composition statistic underlying DUST, the sum over distinct 3-mers of c(c−1)/2 for a triplet occurring c times, divided by the segment length minus three. Conservation and complexity are reported as corroboration; no threshold on either was applied at any stage, and neither was used to select transcripts.

Conservation of each *Gul*-complementary site was scored with UCSC dm6 phyloP27way, retrieved per base from the UCSC REST track API (api.genome.ucsc.edu/getData/track). The 27-insect alignment was used here in preference to the 124-way because it is *Drosophila*-dense, comprising 23 *Drosophila* species and four outgroups, and the question is conservation across the genus; the 124-way alignment is used elsewhere in this study, where the question is instead depth of conservation toward distant insects, and the two are complementary rather than alternative.

For each gene the site was taken as the best antisense hit to *Gul* from the primary dustmasker-filtered BLAST table. BLAST subject coordinates are positions within each transcript’s stored 3′UTR, reverse-complemented for minus-strand genes, and were converted to dm6 genomic coordinates through a per-transcript coordinate table. Every mapping was verified against the reported BLAST mismatch count and against the genomic sequence at the mapped interval before any score was used. Each site was then compared against three references: a 50 bp genomic flank either side for the local profile, the full host 3′UTR as background, and an empirical null of 1,000 random windows of the same length drawn from fully covered positions within the same 3′UTR. Per-gene empirical p values were computed against that null.

#### Predicted miR-7 duplexes

Duplexes between mature dme-miR-7-5p (miRBase MIMAT0000102, 23 nt) and each of the three miR-7 target sites in the *VGAT* 3′UTR were predicted with ViennaRNA 2.7.2 through its Python bindings, using the Turner 2004 parameters at 37 °C with default dangling-end treatment. Two energies were computed for each site: the co-folded duplex minimum free energy, from which a binding energy was obtained by subtracting the two monomer folding energies, and the intermolecular-only duplex energy. Sites were taken as windows centered on each 8mer, and the co-folded and intermolecular energies agree at these sites because no intramolecular structure competes with the duplex over such a window.

#### Multi-species site alignments

Alignments span ten loci across seven species. For each site, the target-side and *Gul*-side arms are shown with their pairing register, positions identical to *D. melanogaster* in black and substitutions in red, and pairing scored as preserved or lost relative to the *D. melanogaster* duplex. Panel extent is set by where the alignment runs out, walking outward from the miR-7 8mer. Alignments were taken from a curated reference set where one was available, and otherwise by reference projection from a Cactus whole-genome alignment ^85^; *CG10444* required realignment with MUSCLE, because the Cactus projection drops *D. virilis*, *D. mojavensis*, and *D. grimshawi* at that locus. Sites recovered by co-variation rather than by sequence carry no *Gul*-side arm, since their 8mers are complementary to any of the three mimic copies and no specific *Gul* position is implied.

#### Co-variation with *Gad1* across cell types

Analyses used the cluster-mean expression matrices described above, covering three regions of the adult CNS: optic lobe, ventral nerve cord, and central brain. Glial clusters were excluded throughout, leaving 231, 115, and 346 neuronal clusters respectively, and GABAergic clusters were defined by an Otsu split of the log *Gad1* distribution; the cuts were 9.88, 8.05, and 3.65, giving 46, 50, and 68 GABAergic clusters. This one definition of “GABAergic” is used for every analysis in the study. Before correlating, each gene’s ranked cluster-mean curve was cut at the onset of its near-silent tail: a baseline was taken as the median of the lowest 20% of clusters, and clusters were retained above baseline plus 2% of the expression span. Genes retained in fewer than 25% of a region’s clusters were then dropped. Co-variation was computed as the Spearman correlation between each gene and *Gad1* over the clusters retained for that gene. Group comparisons were made on medians, by one-sided Mann-Whitney and by permutation with 20,000 iterations, with the transcripts under discussion held out of the background. The one-sided form reflects the directional prediction that these transcripts co-vary more positively than the background, not merely differently. Comparing medians rather than means is load-bearing rather than presentational: the conserved 8mer class is unshifted on medians while its extreme tail is strongly shifted, so a mean-based test on the same data supports a class-wide conclusion that these data do not license.

Within each region, transcripts passing the filters above were ranked by their Spearman correlation with *Gad1*, and enrichment of conserved 8mer miR-7 targets in the top 20 was tested by a one-sided hypergeometric test against all ranked transcripts. The top-20 cut was fixed in advance and class membership is defined by sequence, so the test is independent of which transcripts were chosen for follow-up. No test is reported for the six transcripts against their own site class, because they were selected for occupying the top of that same ranking.

#### The 33-target analysis set, the two-factor model, and *Gul* stoichiometry

The analysis set comprises conserved 8mer miR-7 targets that are detected in all three regions (39 genes) and carry a locatable GTCTTCCA 8mer in the FlyBase r6.58 longest-isoform 3′UTR (33 genes; six drop out because TargetScan’s 3′UTR model differs from FlyBase’s). Other miRNA input was quantified as the number of distinct miRNA families other than miR-7 with a conserved site in the same 3′UTR, from the TargetScanFly 7.2 conserved-family site file filtered to species 7227. Complementarity to *Gul* was quantified by scanning the *Gul* sequence against each 3′UTR for the longest contiguous run of Watson-Crick pairs covering the miR-7 8mer: pairing was extended in both directions from every possible register, runs shorter than the 8 nt site were discarded, and the longest surviving run was recorded. The same scan was also applied genome-wide. The response variable was the mean of the three per-region Spearman correlations with *Gad1*, glia excluded. The two predictors were fitted by ordinary least squares; model fit is reported as R², with a leave-one-out R² as an out-of-sample check and a permutation p obtained by refitting to 20,000 random permutations of the response. 3′UTR length and other-family count were separated by partial correlation and by comparison within bands of similar 3′UTR length.

Abundance was taken as mean expression across neuronal clusters within each region, on the same matrices and with the same glial exclusion as above. Mimic-site stoichiometry was computed in GABAergic clusters only, as the ratio of *Gul* sites to target sites, counting three mimic sites per *Gul* transcript and one miR-7 site per target transcript, summed over the six transcripts. Values are cluster-mean normalized expression and are therefore relative rather than molar.

#### Definition of the *Gul* isoforms

*Gul-G* is the annotated transcript FBtr0308363 (2,917 nt), whose 246-codon open reading frame occupies nucleotides 132 to 872. *Gul-N* is not present in the current FlyBase annotation and was inferred from RNA-seq: it takes the alternative transcription start site of FBtr0345137, which lies 164 nt downstream of the *Gul-G* start codon, extended to the longest annotated 3′ end. The resulting transcript is 2,622 nt, corresponds to nucleotides 296 to 2,917 of *Gul-G*, and is an exact substring of it; the three miR-7 mimic sites lie within the shared 3′ region and are therefore carried by both. This isoform is independently supported by long-read sequencing: transcript model G4015.3 of Alfonso-Gonzalez et al. (2023) uses the same transcription start site and the same two splice junctions, and extends 70 nt further 3′. Of the three long-read models at the locus, it is the only one initiating at the downstream start site, and it also carries the longest 3′ end. The 2,622 nt form defined above was used for the complementarity and mimic-site analyses; open-reading-frame enumeration (below) used the annotated neuronal transcript FBtr0345137.

#### Coding potential of the *Gul* isoforms

Transcript, CDS and protein sequences were retrieved from Ensembl REST for assembly dm6/BDGP6, and orthologues from Ensembl Compara. Open reading frames were enumerated in all three frames of each isoform; for the neuronal isoform this was the annotated transcript FBtr0345137 (1,218 nt). Start-codon context was scored by the *Drosophila* (Cavener) rule, as a purine at −3 and/or a G at +4. Conservation at each candidate start was taken from the UCSC 124-insect alignment as the mean per-base phyloP and phastCons over the three start-codon bases, and regional phyloP and phastCons were computed separately over the coding sequence, the 5′ leader, and the 3′UTR, with a reference distribution sampled from genes drawn from five 5 Mb genomic windows. Frame-aware coding-potential scoring was not applied, so the frame-sensitive evidence comes instead from orthologue residue identity: the residue corresponding to each candidate start was read across aligned orthologues, which distinguishes a conserved start codon from a conserved codon lying within a different reading frame.

#### Ribosome profiling analysis

To assess the translation efficiency of *Gul* in the nervous system, we analyzed ribosome profiling (Ribo-seq) data from adult *Drosophila* heads ^86^. Transcript abundance and translational efficiency were taken from its Supplementary File 1, which reports RNA-seq TPM, Ribo-seq CDS TPM, and translational efficiency (Ribo-seq CDS TPM divided by RNA-seq TPM) for 13,796 genes, one representative transcript per gene; all values reported here are from the whole-head dataset. Abundance rank was computed across transcripts with a non-zero RNA-seq value, and the translational-efficiency percentile across transcripts with non-zero, finite values for both measurements. *Gul* was additionally compared against an abundance-matched reference set, every transcript whose RNA-seq value falls within two-fold of *Gul*’s, and its translational efficiency is reported against the number of that set falling below it and against the set median. Because the table carries one transcript per gene, this analysis does not resolve *Gul-G* from *Gul-N*; the *Gul-N* transcription start site lies within the *Gul-G* coding sequence, so the annotated coding window over which ribosome occupancy is measured contains the open reading frames of both.

### QUANTIFICATION AND STATISTICAL ANALYSIS

#### Image analysis and quantification

Confocal image stacks were processed using Fiji (ImageJ) ^87^ and Adobe Illustrator. Quantification of reporter expression levels between GABAergic and non-GABAergic neurons was performed on confocal image stacks using a semi-automated pipeline consisting of three steps: nuclear segmentation, per-cell intensity quantification, and statistical comparison.

Nuclear segmentation was performed on 3D confocal stacks using Cellpose ^88^ under a custom nextflow workflow (https://github.com/chenyenchung/cellpose_nf) with the default pretrained model. Segmentation was run in 3D mode with anisotropy correction applied using the z-step size and xy pixel size extracted from image metadata. Segmentation parameters were as follows: cell probability threshold = 0, flow threshold = 0, stitch threshold = 0.25, minimum cell size = 100 voxels, and maximum size fraction = 0.02. Images were processed with channel axis set to axis 1 and z-axis set to axis 0.5.

Per-cell quantification was performed using a custom Python script. For each segmented nuclear ROI, the fraction of nuclear voxels overlapping the Gad1 cytoplasmic marker mask (to define GABAergic neurons) was computed, and cells were classified as GABAergic (cytoplasmic marker positive) or non-GABAergic (cytoplasmic marker negative) using an overlap threshold of 60%; glia were excluded on a looser 10% overlap with the glial nuclear-counterstain mask. Mean, maximum, sum, and standard deviation of reporter fluorescence intensity were measured within each nuclear ROI.

Statistical comparison of reporter intensity between GABAergic and non-GABAergic populations was performed in R using a two-sided Wilcoxon rank-sum test. Effect size was calculated as the rank-biserial correlation coefficient r. Fold change was computed as the ratio of median intensity in cytoplasmic marker-positive versus marker-negative cells, where each cell’s intensity is the mean reporter signal within its nuclear ROI. Results were visualized as violin plots with overlaid boxplots and jittered individual data points (subsampled to 400 cells per group for display). All analyses were performed across three biological replicates. Full analysis code is provided in Supplementary Scripts 3-5.

#### Local non-GABAergic baseline

This analysis used only the medulla region of the optic lobe for each sensor. For the per-neuron comparison each cell was scored against a local reference rather than the population as a whole. The neighborhood is defined by count, not radius: the K = 10 nearest cytoplasmic marker-negative cells, by Euclidean distance between segmentation-mask centroids in microns, using the stack’s xy pixel size and its z step from the acquisition metadata (0.50 or 1.00 µm). Distances are in microns rather than voxels because the z step is two to four times coarser than the xy pixel size. A cell entered the reference pool if it was marker-negative and its sensor intensity was at least 2.0 on the 8-bit scale; cells below that floor were excluded from the pool but still scored themselves. Queries were every segmented nucleus and, separately, every GABA ROI centroid, with the search leave-one-out for marker-negative queries so that a cell never anchors on itself. Neighbours were found with a k-d tree, k truncated to the pool size where fewer than ten eligible cells were present. The local reference is the median sensor intensity of those neighbors; each cell’s relative output is its own intensity divided by that reference, and a GABAergic neuron was counted as failing to exceed its neighbors when its relative output was at or below the 75th percentile of the relative outputs of the non-GABAergic reference cells in that region, a conservative threshold slightly above parity (approximately 1.05 to 1.12), set so that one-quarter of background cells fall below it by construction.

K was chosen on a leave-one-out null, marker-negative cells scored against their own neighbors, where a perfect reference returns one plus noise. Spread is minimized between K = 4 and K = 10 in every stack and is 10 to 15% worse at K = 50; K = 10 was preferred over K = 4 because a four-cell median is fragile. No distance cut was applied: a hard limit was tested and rejected because it removed real cells, and the distance to the tenth neighbor is retained per cell as a diagnostic, with a median of 6.4 to 8.9 µm and a 95th percentile of 11.5 to 36.9 µm across stacks.

## Lead contact

Further information and requests for resources and reagents should be directed to and will be fulfilled by the lead contact, Claude Desplan.

## Materials availability

Fly strains and plasmids generated in this study are available from the lead contact upon request.

## Data and code availability

All plasmid and oligonucleotide sequences are listed in Supp. Tables 1 and 2, and the per-cell, per-gene, and per-site data underlying the quantitative analyses are provided in Supp. Tables 3-11. All original code used for these analyses is deposited in a public repository ([repository DOI to be added]) and is publicly available as of the date of publication. Fly strains and the raw imaging data generated in this study are available from the corresponding authors upon request.

## Declaration of generative AI and AI-assisted technologies

Claude (Anthropic) was used to generate code in R and python for data processing and visualization. All output generated by AI was supervised, vetted, and edited by the authors before final use.

## Supporting information

supplemental tables and scripts

## Acknowledgements

We thank Ildar Gainetdinov, Richard Carthew, Eric Klann, and Isabel Holguera for thorough feedback on the manuscript and members of the Desplan lab for helpful discussions. We thank the Bloomington *Drosophila* Stock Center for stocks (NIH P40OD018537) and FlyBase ^89^ for genome annotation and sequence data (NHGRI U41 HG000739). We also thank Leslie Griffith for providing *VGAT*-AttP used for making the *VGAT* translation reporter, Julie Simpson for providing the *VGAT*-LexA transcription reporter, Andrea Brand for providing the miR-7 null allele, and Richard Carthew for providing miR-7 reporter stocks. We also thank David Krantz for providing VGAT antibody. The Desplan laboratory was supported by grants from the National Eye Institute R01EY017916 and R01EY13010. R.L. was supported by a Ruth L. Kirschstein National Research Service Award (NRSA) Individual Postdoctoral Fellowship (5F32EY035122).

## Author contributions

R.L. and C.D. conceived the project. R.L. and C.D. wrote the manuscript. R.L. designed and performed all the experiments. R.L. analyzed the data.

