## supplemental figures and legends for "Long non-coding RNA-3′UTR interactions coordinate GABA signaling in the brain"

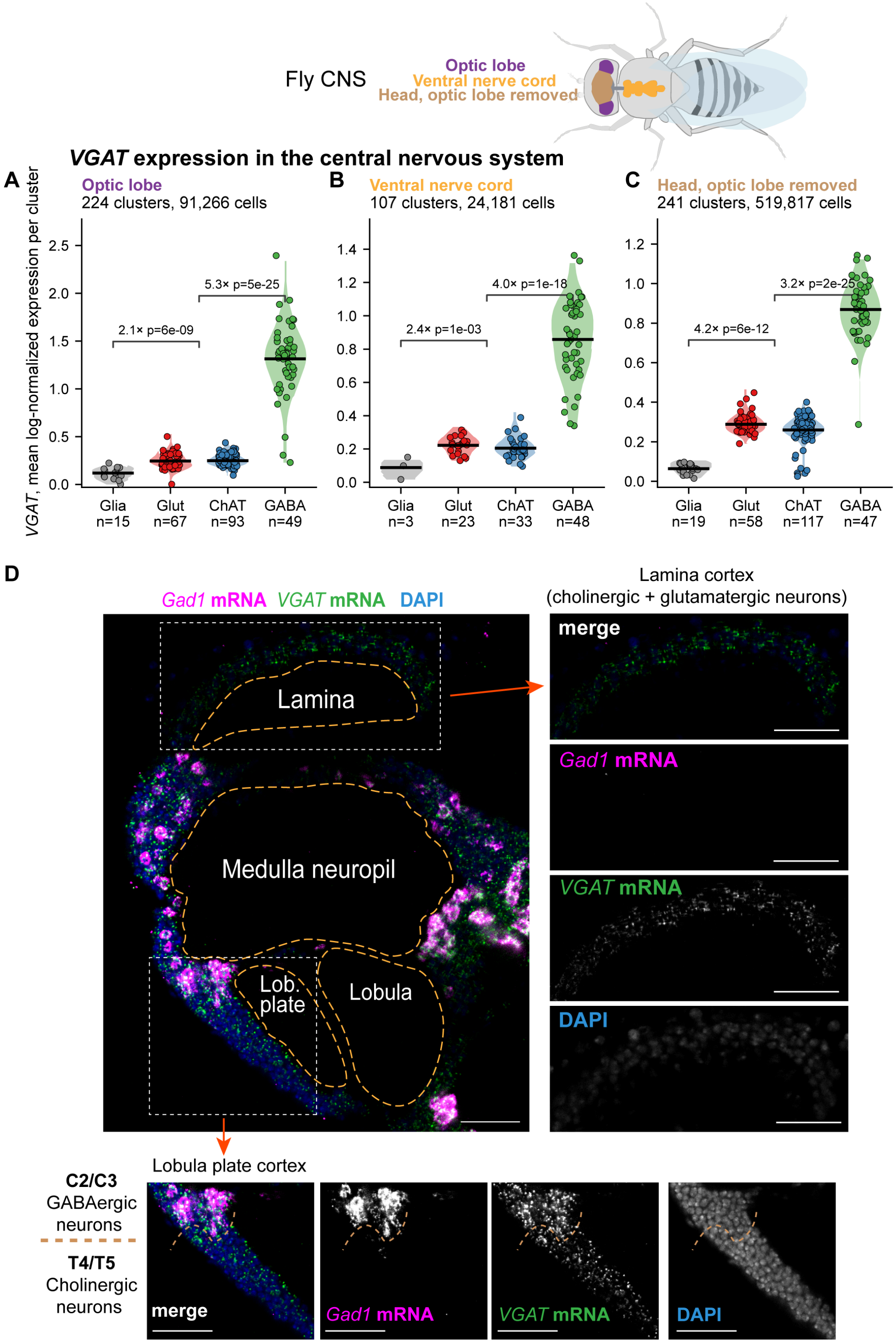


**Figure S1.** ***VGAT*** **mRNA is present in all neuronal classes throughout the central nervous system.**

**(A to C)** Mean *VGAT* expression per cluster in adult single-cell RNA-seq, grouped by neurotransmitter class, in the optic lobe (A), the ventral nerve cord (B) and the head with optic-lobe-labeled clusters removed (C). Each dot is one cluster, the bar is the class median, and the number of clusters per class is given below. Classes are colored glia grey, glutamatergic red, cholinergic blue and GABAergic green. Fold-differences and p values are shown for the GABAergic against cholinergic and the non-GABAergic against glial comparisons. Each panel carries its own y axis and the three cannot be compared with one another.

**(D)** HCR RNA-FISH of the adult *Drosophila* optic lobe showing the distribution of *VGAT* mRNA (green) relative to GABAergic neurons (magenta) marked by *Gad1* mRNA, with nuclei labeled by DAPI (blue). Insets show the lamina cortex, which contains only cholinergic and glutamatergic neurons and glia, and lobula plate cortex with the boundary between C2/C3 GABAergic and T4/T5 cholinergic neurons (dashed line). Scale bar: 20 µm.

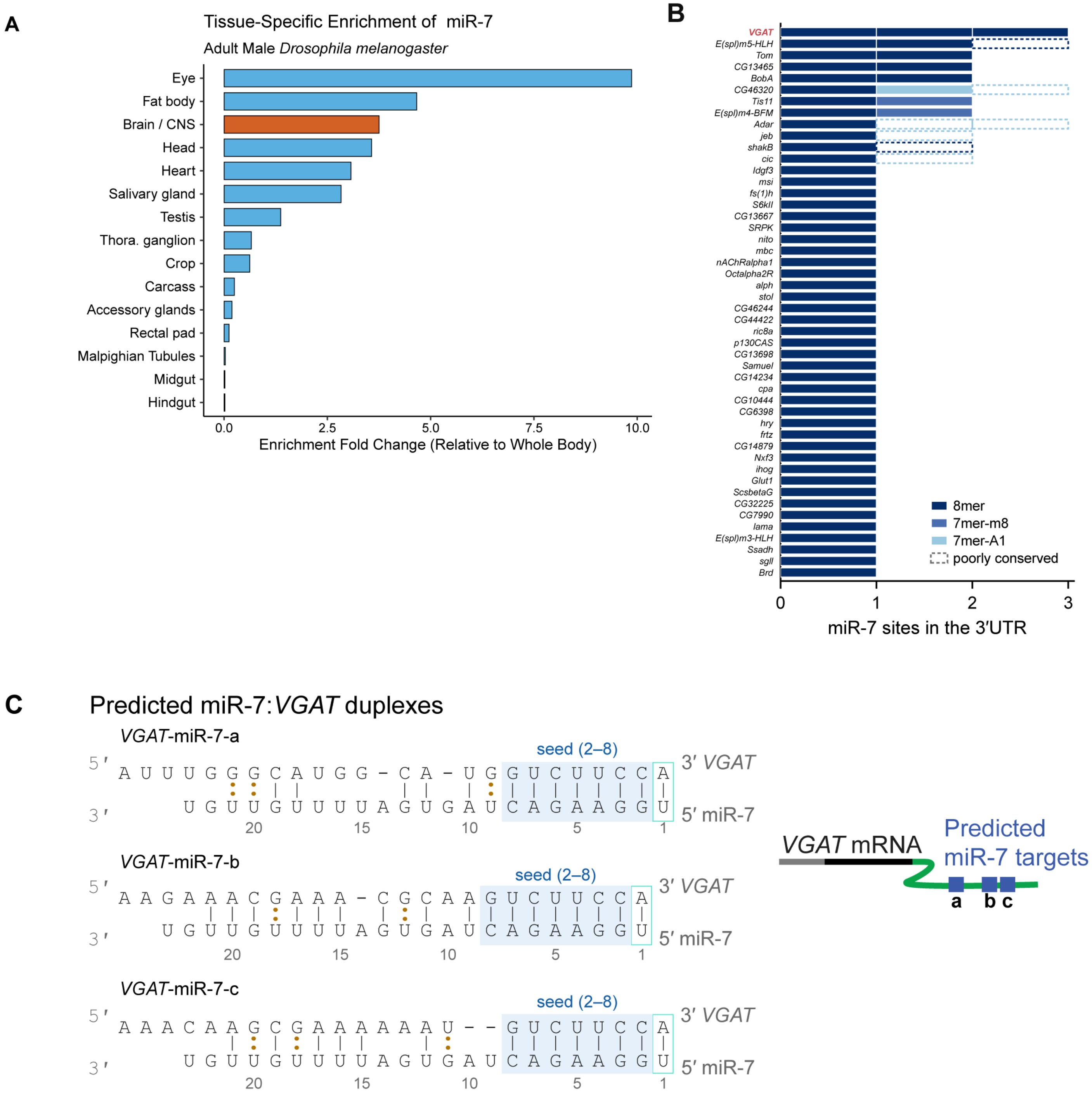


**Figure S2. miR-7 and its predicted target sites in the *VGAT* 3′UTR.**

**(A)** miR-7 abundance across adult male tissues from published small RNA sequencing, with brain and CNS in orange.

**(B)** The 48 genes in the *Drosophila* genome carrying a conserved miR-7 8mer, one row per gene, ordered by site number**.** Colors give the seed match type and a dashed open segment marks a site whose conservation is no better than chance for a site of its type. Conserved evaluation is based on TargetScan's PCT-based call.

**(C)** Predicted duplexes between each of the three *VGAT* miR-7 target sites and the full 23 nt miR-7 sequence, showing base-pairing outside the seed.

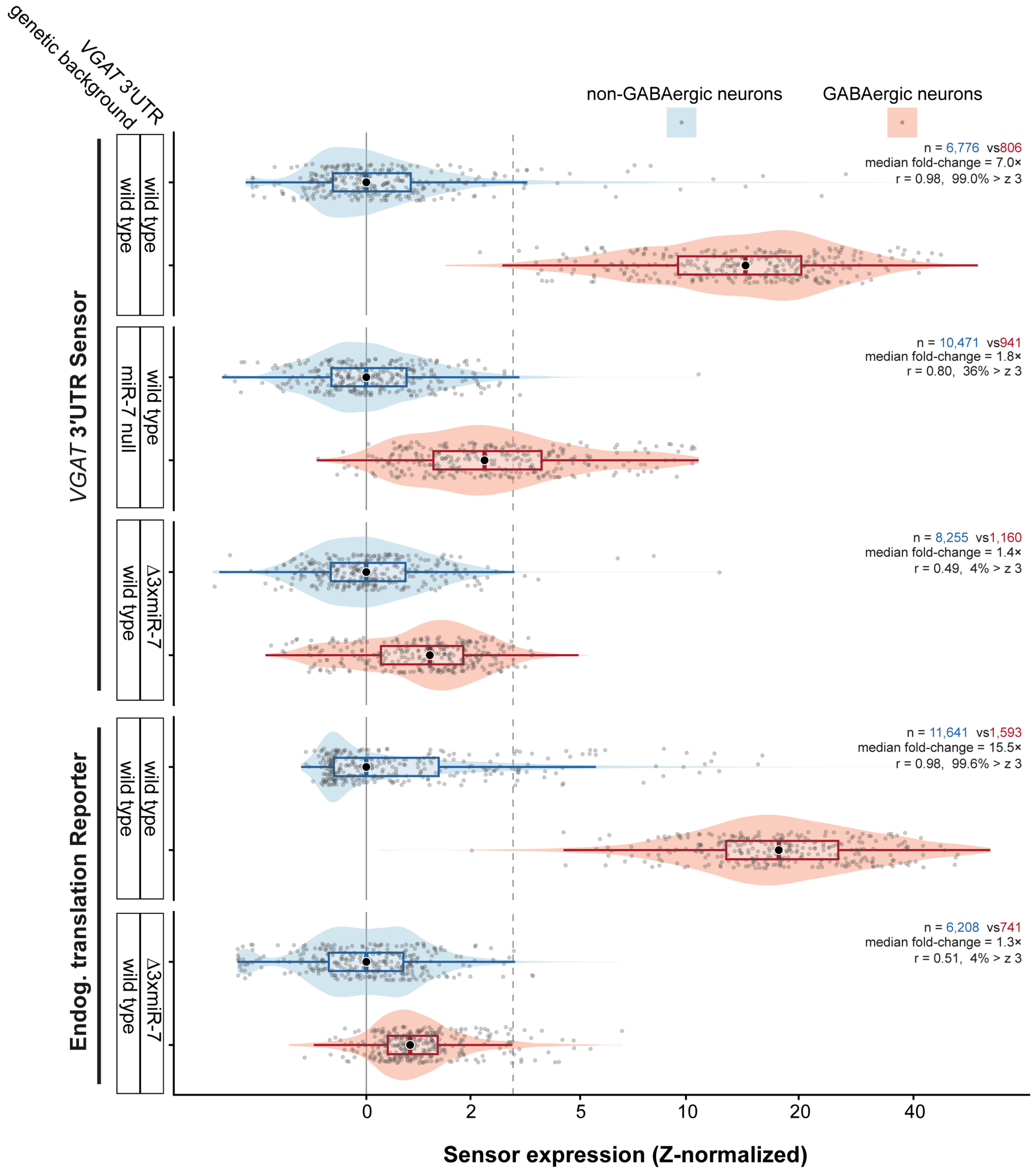


**Figure S3. Quantification of *VGAT* 3′UTR sensor and translation reporter activity.** Per-neuron intensity in GABAergic against non-GABAergic neurons for five genotypes, each Z-normalized to its own non-GABAergic population. For each genotype the panel gives the number of neurons of each class, the median fold-change between them, the correlation across paired measurements, and the percentage of GABAergic neurons exceeding z = 3.

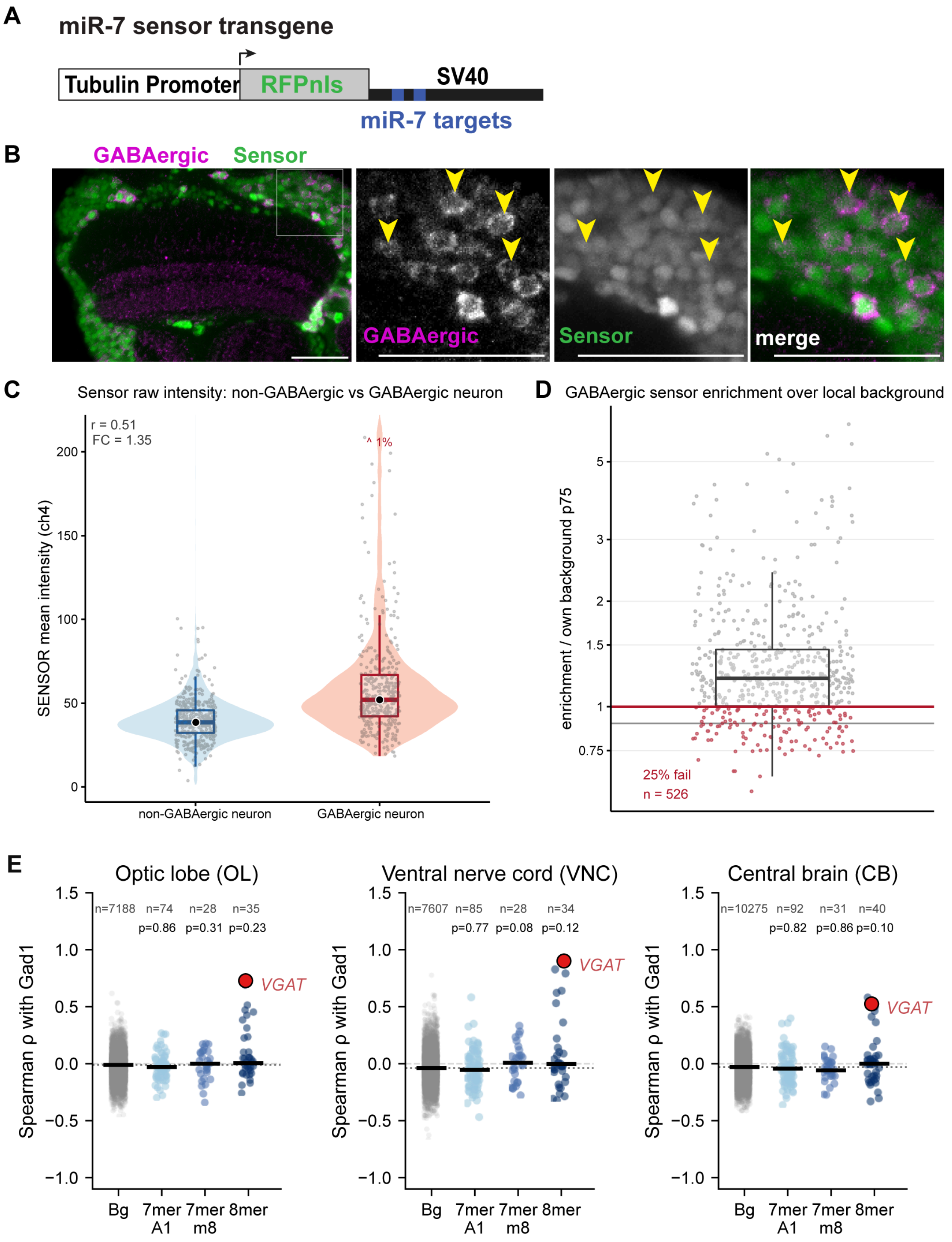


**Figure S4. miR-7 activity is comparable in GABAergic and non-GABAergic neurons.**

**(A)** Diagram of the miR-7 sensor transgene: a tubulin promoter driving nuclear-localized RFP, with two complete 23-nucleotide complements to mature miR-7-5p, separated by 25 nucleotides, upstream of an SV40 terminator. The sensor is red and is pseudocolored green in the panels below.

**(B)** Activity of the miR-7 sensor (green) in the optic lobe of wild type flies relative to GABAergic neurons (magenta) marked with a Gad1 antibody. Arrowheads mark GABAergic neurons marked with a Gad1 antibody. Scale bar: 20 µm. Inset: region of medulla cortex.

**(C)** Per-neuron sensor intensity in non-GABAergic against GABAergic neurons, with the correlation and the median fold-change between the two populations given.

**(D)** Per-neuron sensor enrichment over the local background of the same optic lobe as panel C, with the percentage of GABAergic neurons failing to exceed their background given.

**(E)** Mean Spearman ρ with *Gad1* for every analyzed transcript, grouped by miR-7 site class, in the optic lobe, ventral nerve cord and central brain. *VGAT* is marked, bars are class medians, and n and the p value against background are given above each column.

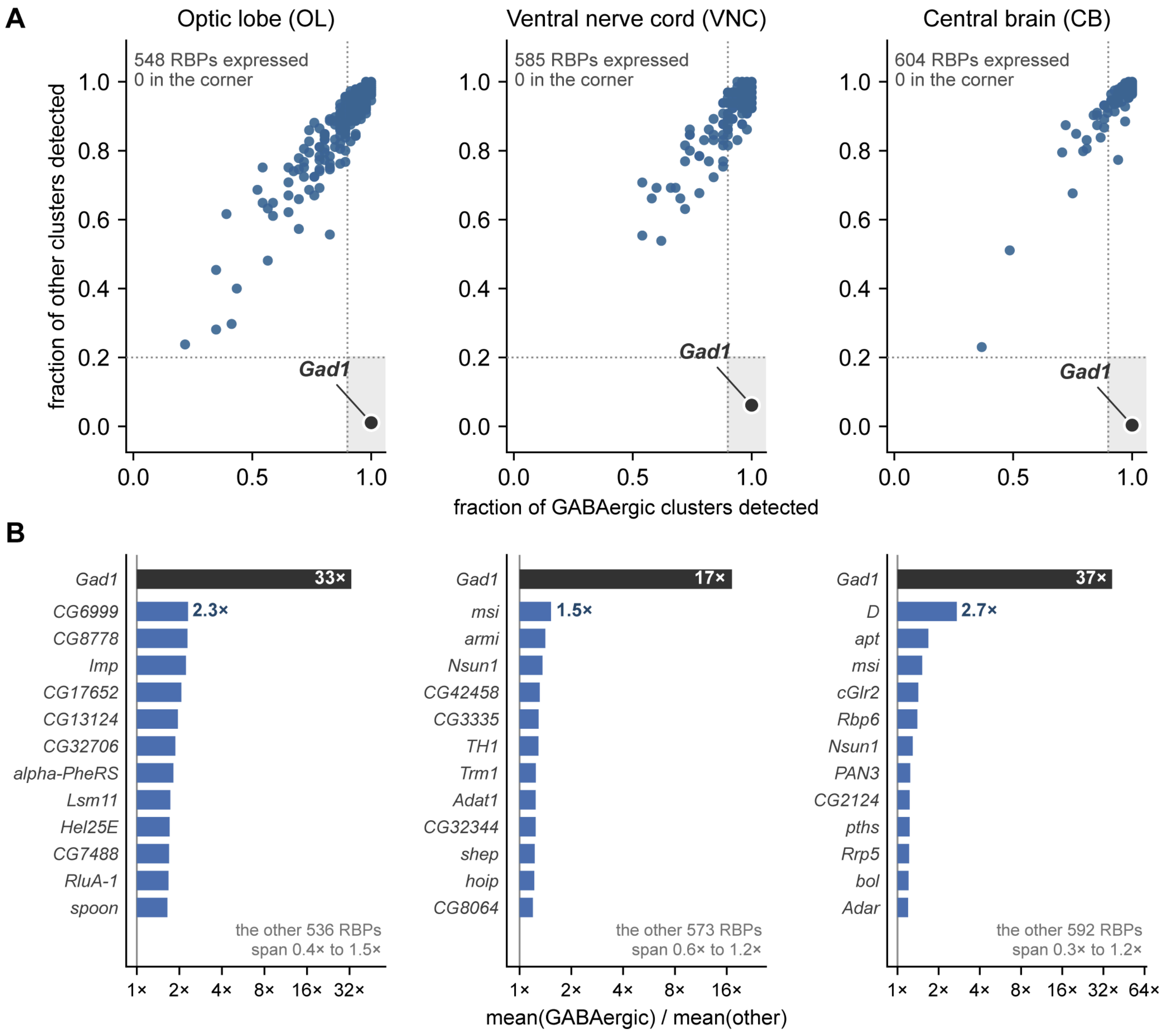


**Figure S5. No RNA-binding protein is restricted to, or excluded from, all GABAergic neurons.**

**(A)** Detection frequency of the 548, 585 and 604 annotated RNA-binding proteins expressed in the optic lobe, ventral nerve cord and central brain, plotted as the fraction of GABAergic clusters against the fraction of other clusters in which each is detected. The shaded corner is detection in ≥90% of GABAergic clusters and ≤20% of the others. *Gad1* is plotted as a positive control.

**(B)** The most GABAergic-enriched RNA-binding proteins in each region, one bar per gene, with *Gad1* shown for comparison.

GABAergic clusters are defined by an Otsu split of the log *Gad1* distribution and glia are excluded, the same cluster sets used in Fig. S7B.

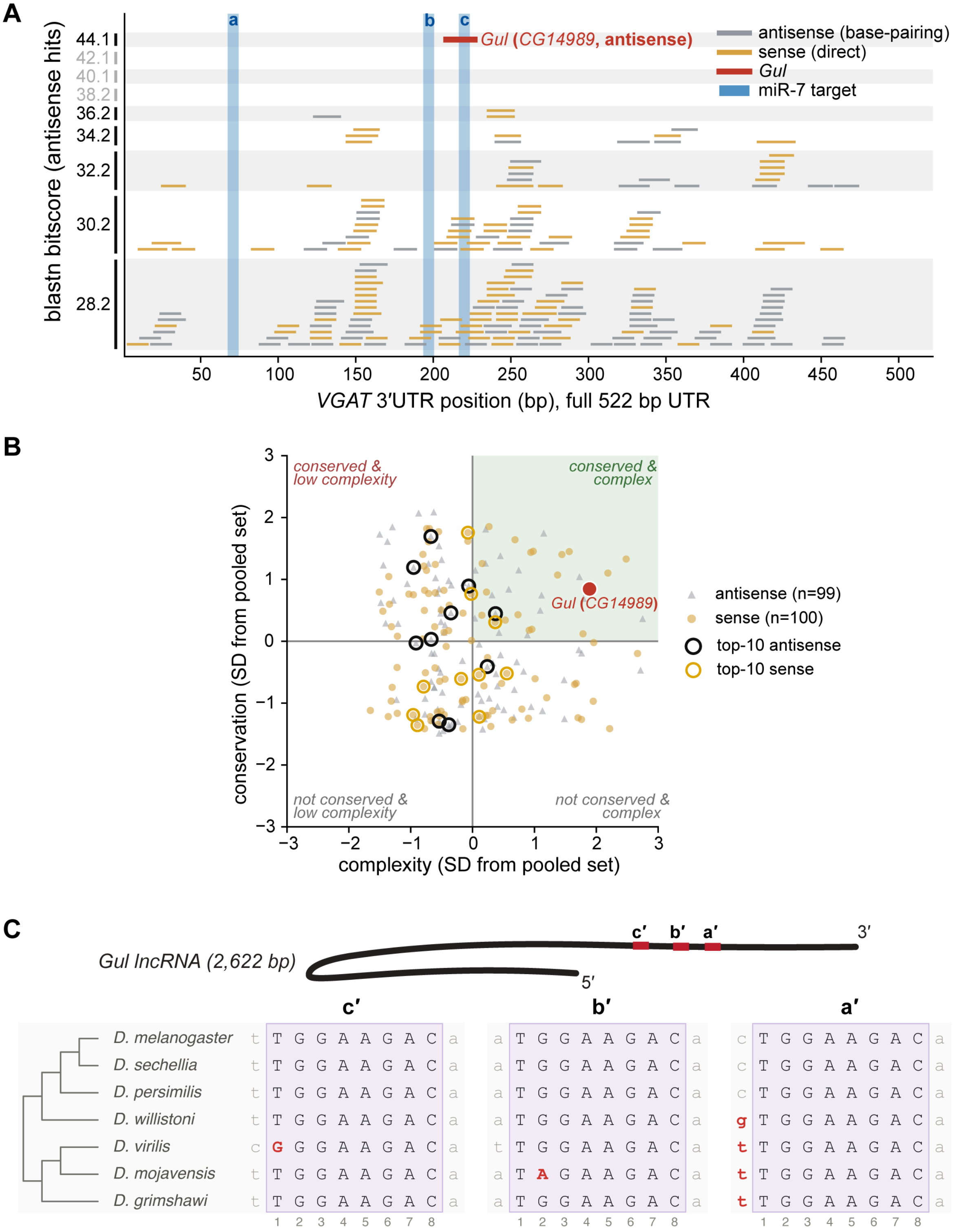


**Figure S6. BLAST of the *VGAT* 3′UTR against annotated transcripts.**

**(A)** Every hit plotted at its position along the 522 bp *VGAT* 3′UTR against its blastn bit-score. Antisense hits (grey) can base-pair with *VGAT*; sense hits (gold) are shown as a strand control, since a match in the same orientation cannot pair. The three miR-7 target sites are shaded. *Gul* (*CG14989*) is labeled with its bit-score and E-value.

**(B)** Drosophilid branch-length conservation against sequence complexity (the inverse of A+T fraction) for the top 100 hits of each orientation, both z-scored on one pooled reference: the antisense background with *Gul* excluded (n = 99) plus all sense hits (n = 100). Open circles mark genes with the top 10 blastn bitscore of each set from panel A. The quadrant is occupied equally by the two orientations, 20 of 99 antisense against 18 of 100 sense.

**(C)** The three *Gul* mimic sites c′, b′ and a′ aligned across seven Drosophilid species, with their positions on the 2,622 bp *Gul-N* transcript above, drawn 5′ to 3′. The 8mer is boxed and in upper case, flanking bases in lower case, and any base differing from *D. melanogaster* is red.

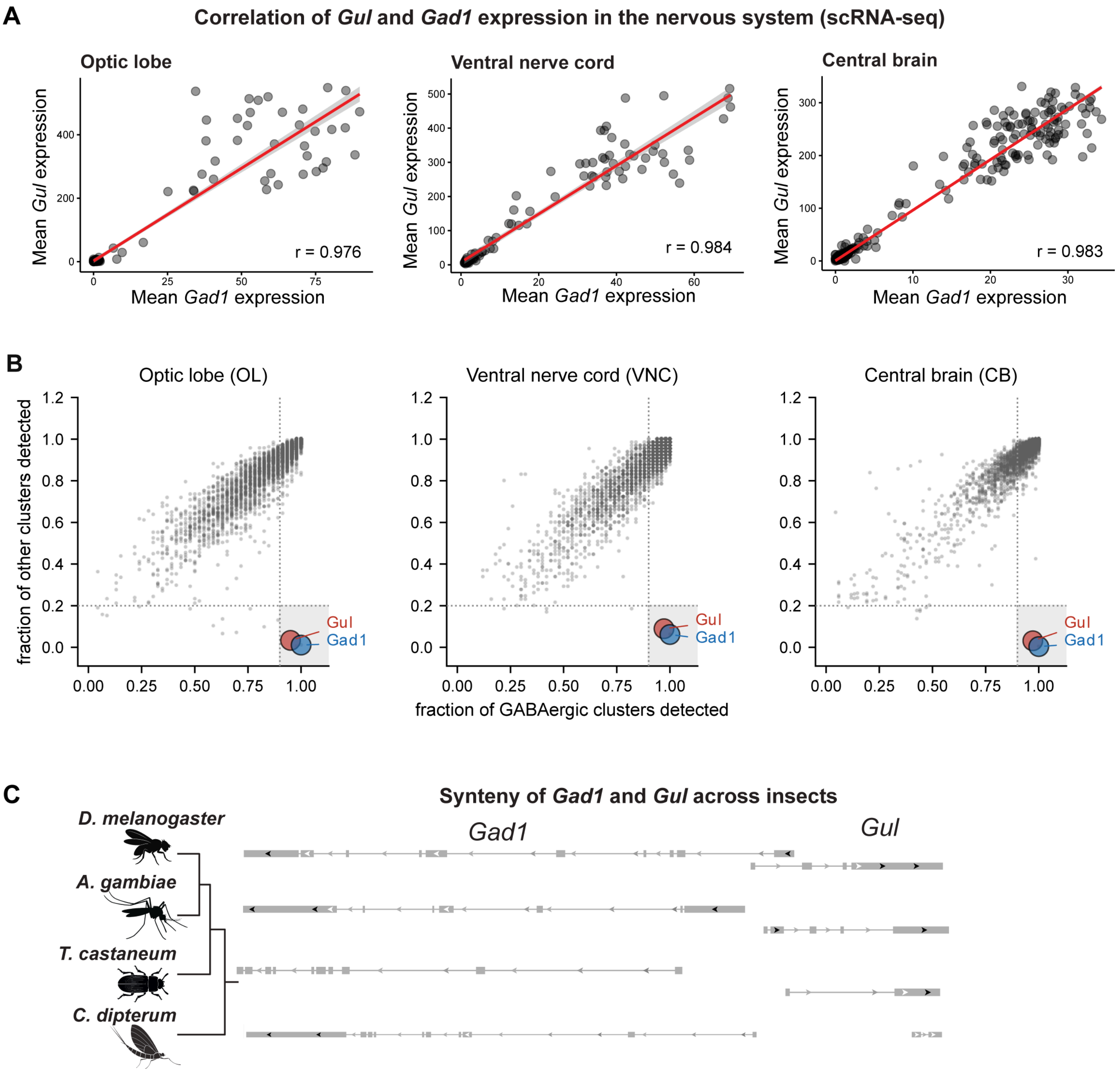


**Figure S7. *Gul* and *Gad1* are the only two transcripts restricted to the GABAergic population.**

**(A)** *Gul* expression per cluster across the optic lobe, ventral nerve cord and central brain, plotted against *Gad1*.

**(B)** Detection frequency of all expressed transcripts (7,073, 7,360 and 8,001 in the three regions), plotted as in Fig. S5A, with *Gul* (red) and *Gad1* (blue) marked. The shaded corner is detection in ≥90% of GABAergic clusters and ≤20% of the others, occupied by only *Gul* and *Gad1* in every region. *Gul* and *Gad1* are detected in identical fractions of clusters in the ventral nerve cord and central brain and differ by 0.02 in the optic lobe; their markers are semi-transparent and offset by 0.028 so that both are visible, and the separation drawn is not a measured difference.

**(C)** Synteny of the *Gul* and *Gad1* locus across insects.

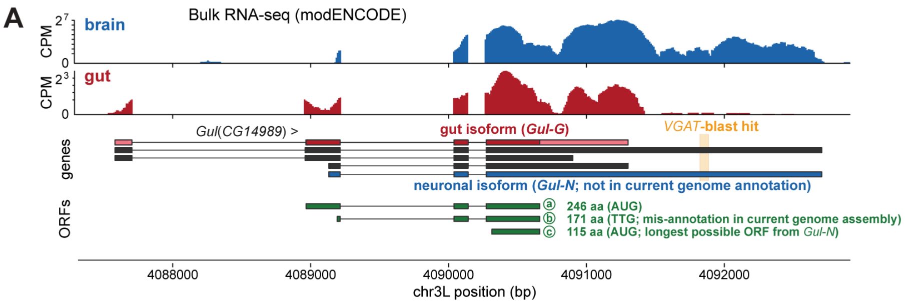


**Start-codon context and conservation**

| ORF | start codon | Kozak context | phyloP | phastCons |
|---|---|---|---|---|
| ⓐ 246 aa (annotated *Gul-G* start) | AUG | strong | +1.77 | 1.00 |
| ⓑ 171 aa (mis-annotated *Gul-N* start) | **TTG** | N.A (non-AUG) | +11.34 | 1.00 |
| ⓒ 115 aa (longest possible *Gul-N* ORF) | AUG | moderate | **−0.29** | **0.02** |

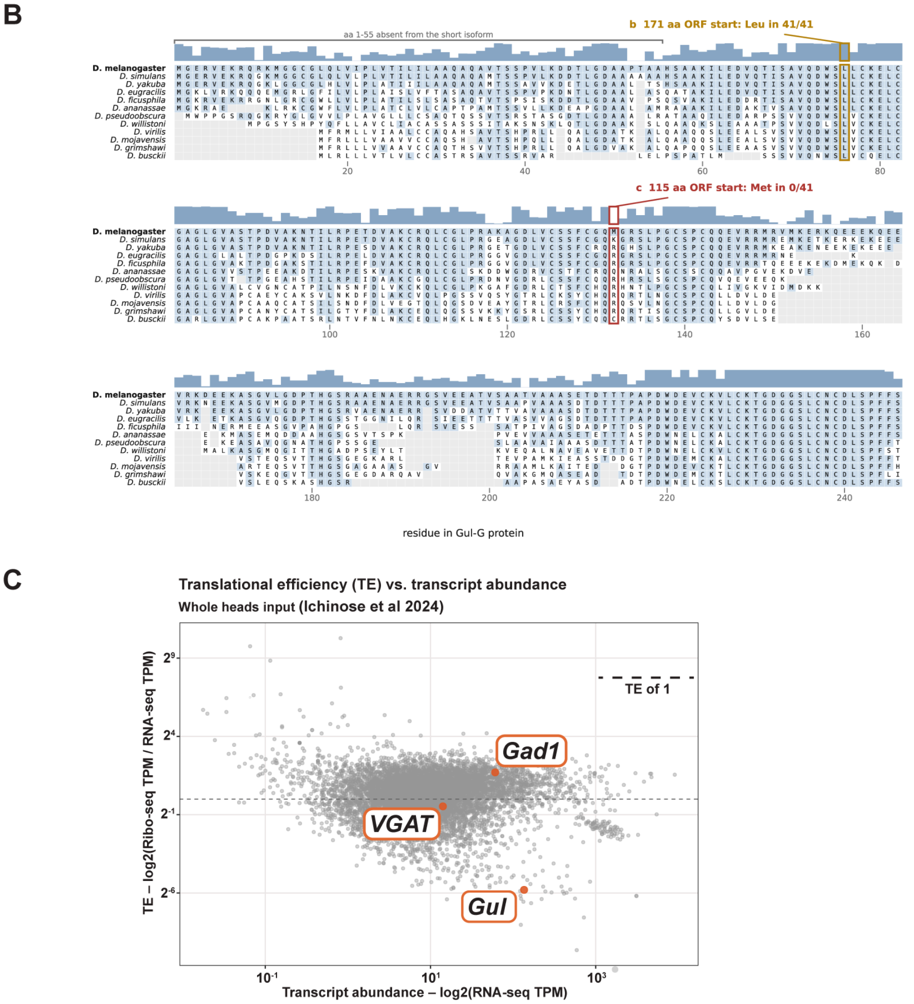


**Figure S8. The neuronal isoform *Gul-N* is non-coding.**

(A) Diagram of the two *Gul* isoforms showing their open reading frames, with a table of start-codon context and conservation for each. ORF a, in the gut isoform *Gul-G*, has an AUG in strong Kozak context that is conserved. ORF b is the start currently annotated for the neuronal isoform, but it is a TTG rather than an AUG and so has no Kozak context to score; it lies in frame within ORF a, so the conservation at that position reflects the *Gul-G* coding sequence rather than a genuine start site. ORF c, the longest ORF *Gul-N* could initiate from an AUG, has only moderate Kozak context and is not conserved.

**(B)** Alignment of the 246 aa *Gul-G* open reading frame across 41 drosophilid species (11 shown, spanning roughly 60 million years of divergence), with per-column conservation plotted above each block. The start of the annotated 171 aa neuronal ORF (b) aligns to a leucine that is conserved as Leu in all 41 species, a constrained residue within the *Gul-G* reading frame rather than a translation start; the first AUG available to the 115 aa ORF (c) aligns to a methionine found in 0 of 41. The first 55 residues of *Gul-G* are absent from the neuronal isoform.

**(C)** Ribo-seq data from the *Drosophila* head (Ichinose et al., 2024) plotting translational efficiency, *i.e.*, the ratio of ribosome protected fragments to transcripts, against transcript abundance, with *Gul*, *Gad1*, and *VGAT* marked. Of the 559 transcripts within two-fold of *Gul*'s abundance, only 11 have a lower translational efficiency; the median for that group is 1.66, against 0.018 for *Gul*.

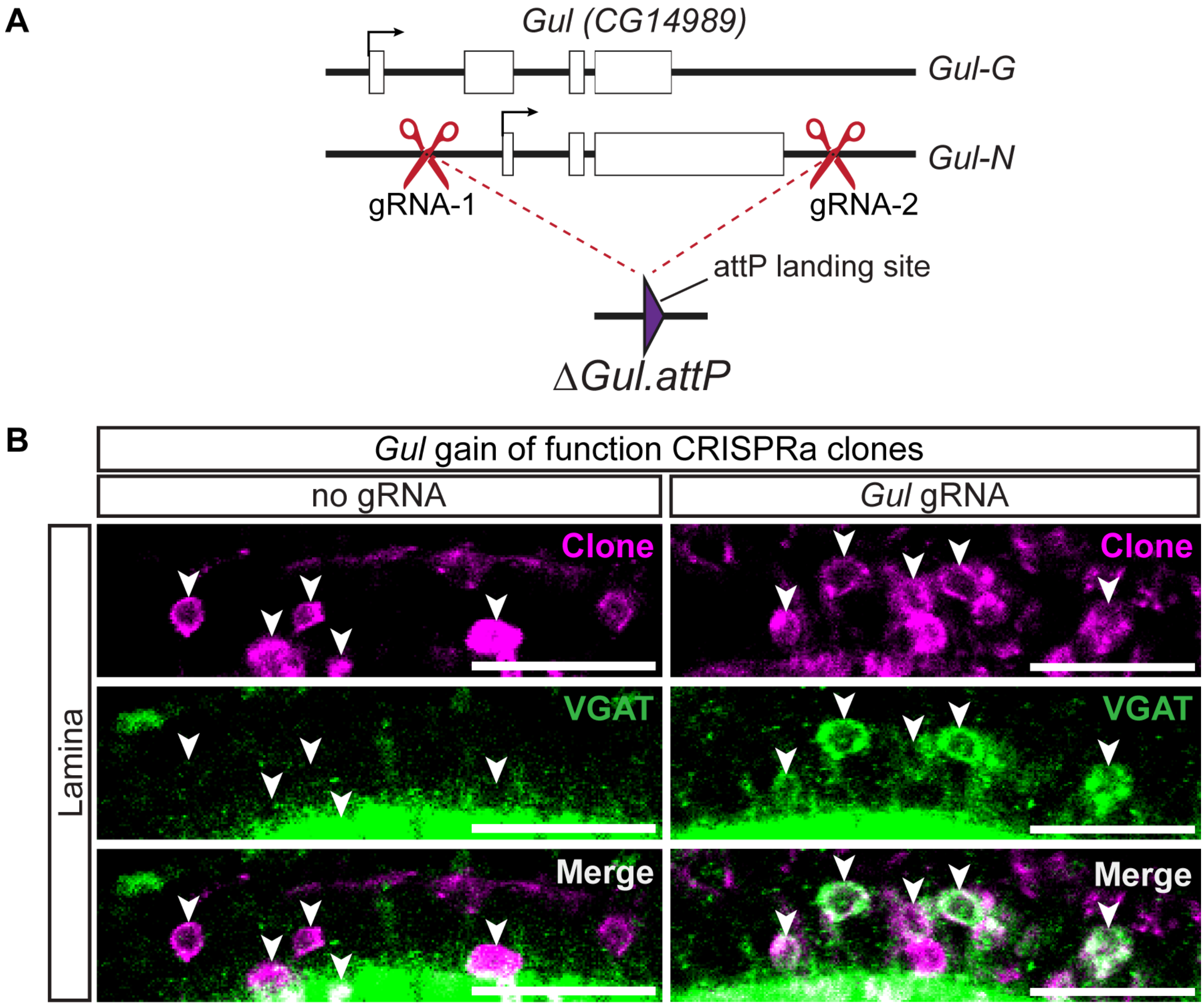


**Figure S9.** ***Gul*** **loss- and gain-of-function alleles.**

**(A)** Diagram of the Δ*Gul* deletion, in which two guide RNAs flanking the *Gul* gene body were used to replace it with an attP landing site. Both isoforms are removed.

**(B)** Immunofluorescence of *Gul* CRISPRa clones (magenta) in the lamina cortex showing VGAT protein (green). Arrowheads mark cells within a clone. Clones generated without a guide RNA are shown as a control. Scale bar: 20 µm.

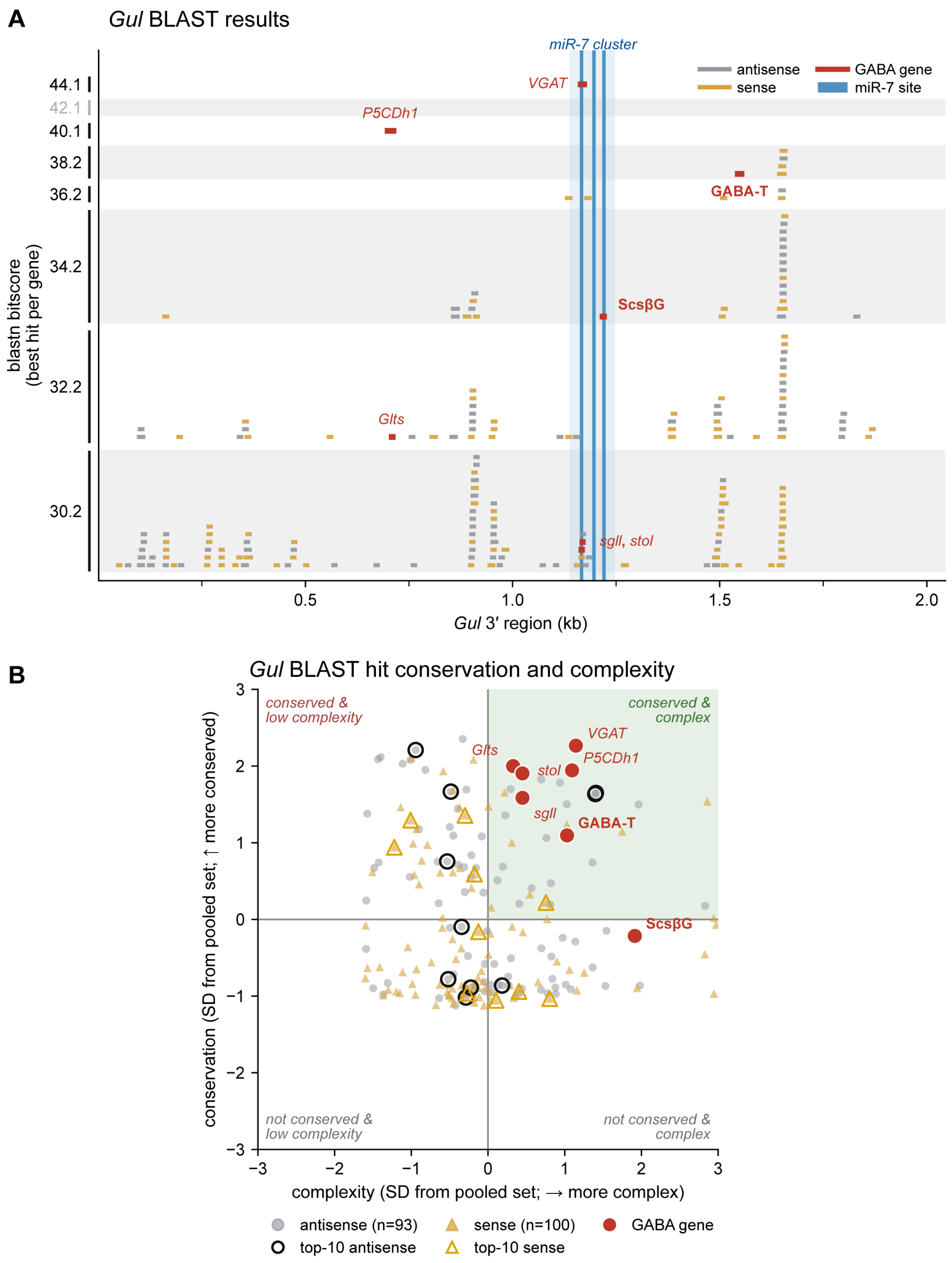


**Figure S10. BLAST of the *Gul* transcript against annotated 3′UTRs.**

**(A)** Every hit from the full-transcript search plotted at its position along the 2,045 bp *Gul* 3′ region against its blastn bit-score, one best hit per gene, top 100 per strand. **Antisense hits (grey) can base-pair with *Gul*; sense hits (gold) are shown as a strand control.** The miR-7 cluster and the three GGAAGACA mimic copies are shaded. The seven GABA-pathway transcripts are red and named; *sgll* and *stol* share one label, being the same 15 nt score at overlapping positions.

**(B)** Drosophilid branch-length conservation of the matched site against sequence complexity for the top 100 hits of each orientation, both z-scored on one pooled reference: the antisense set with the seven excluded (n = 93) plus all sense hits (n = 100). Open circles mark genes with the top 10 blastn bitscore of each set from panel A.

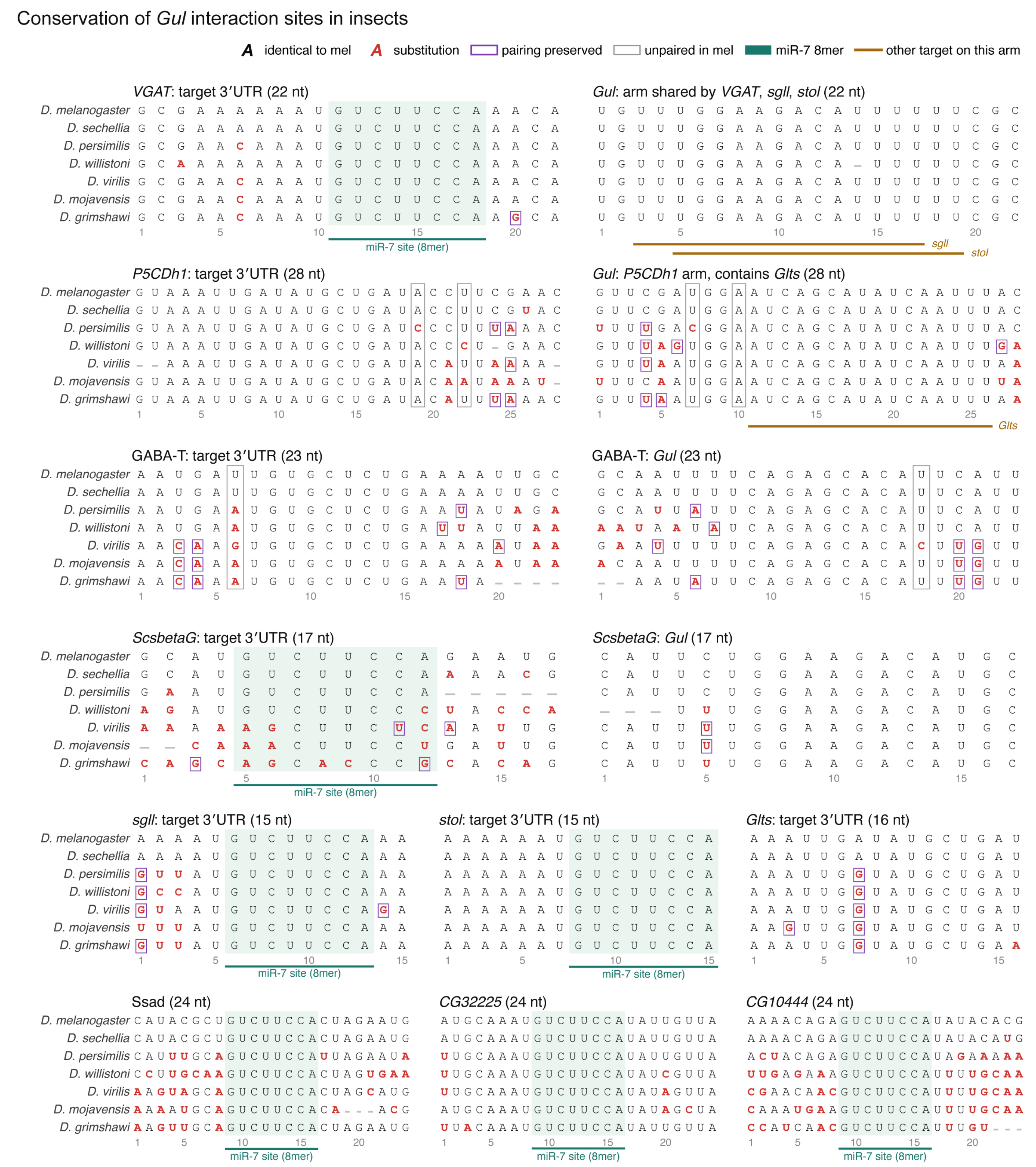


**Figure S11. Conservation of the *Gul*:target interaction sites.** Alignments across seven Drosophilid species for each transcript recovered by the two screens, showing the target-side site on the left and the paired *Gul*-side arm on the right. Each distinct *Gul* arm is drawn once, with the transcripts that share it marked as amber footprint bars. Bases are shown in register with *D. melanogaster*, substitutions are red, and purple marks positions where predicted pairing is preserved, which is not the same as covariation. Gaps are left unfilled and some are alignment artifacts. Teal shading marks a miR-7 8mer; *P5CDh1*, *Glts* and *GABA-T* carry no miR-7 site of any class and are therefore unshaded.

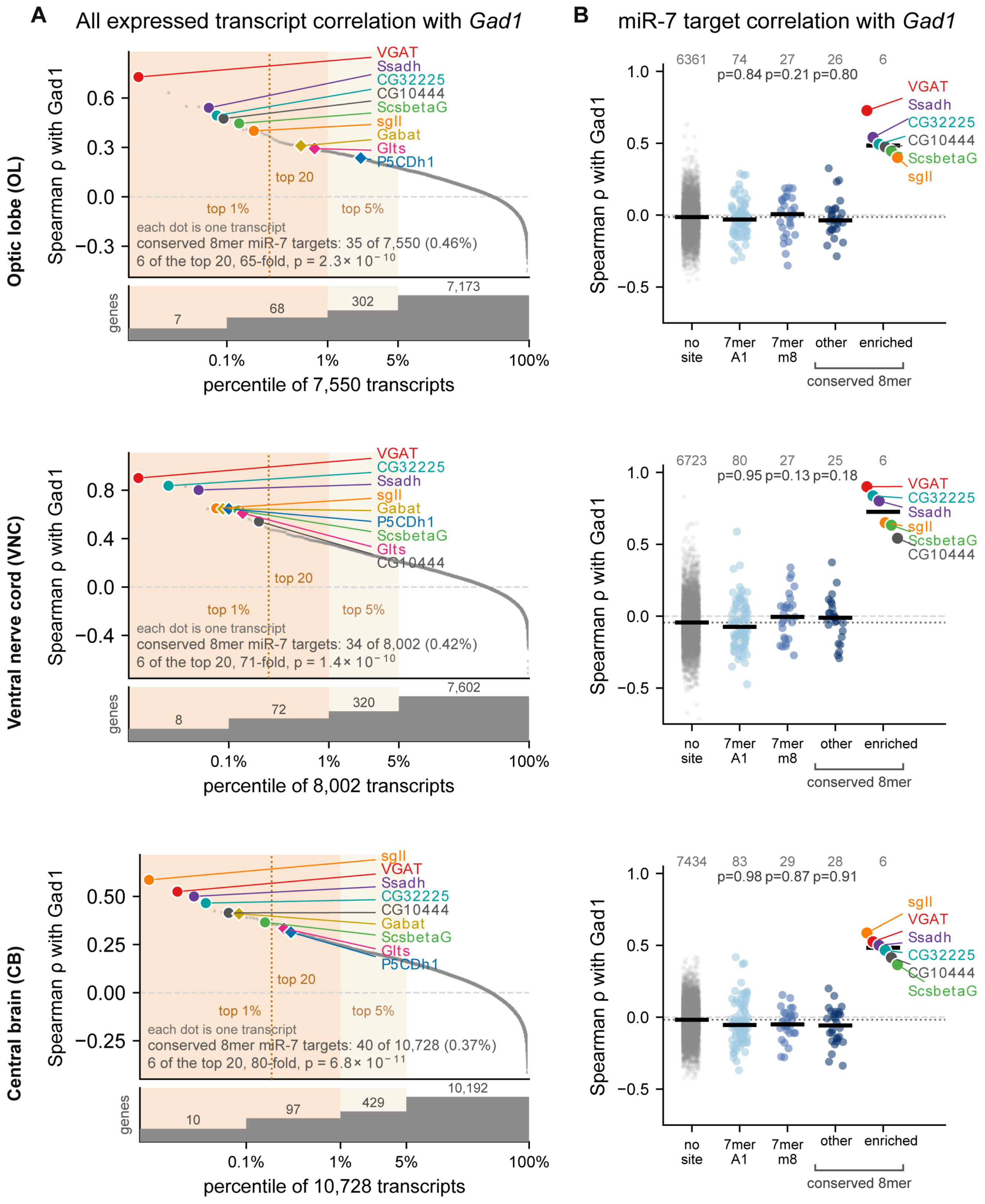


**Figure S12. Co-variation with *Gad1* across three CNS regions.** Rows are regions: optic lobe, ventral nerve cord and central brain. Glial clusters are excluded throughout and GABAergic clusters are defined by an Otsu split of the log *Gad1* distribution.

**(A)** Every analyzed transcript ranked by its mean Spearman ρ with *Gad1*, plotted against percentile. Shaded bands mark the top 1% and top 5%, and the strip beneath each panel gives the number of transcripts in each band. The nine candidate transcripts are labeled.

**(B)** The same regions grouped by miR-7 site class, with the six enriched transcripts drawn as a separate column. Bars are class medians and n is given above each column. No p value is drawn over the enriched column, because those transcripts were selected for sitting at the top of this same ranking. The enriched set is *VGAT*, *Ssadh*, *CG32225*, *sgll*, *ScsbetaG* and *CG10444*.

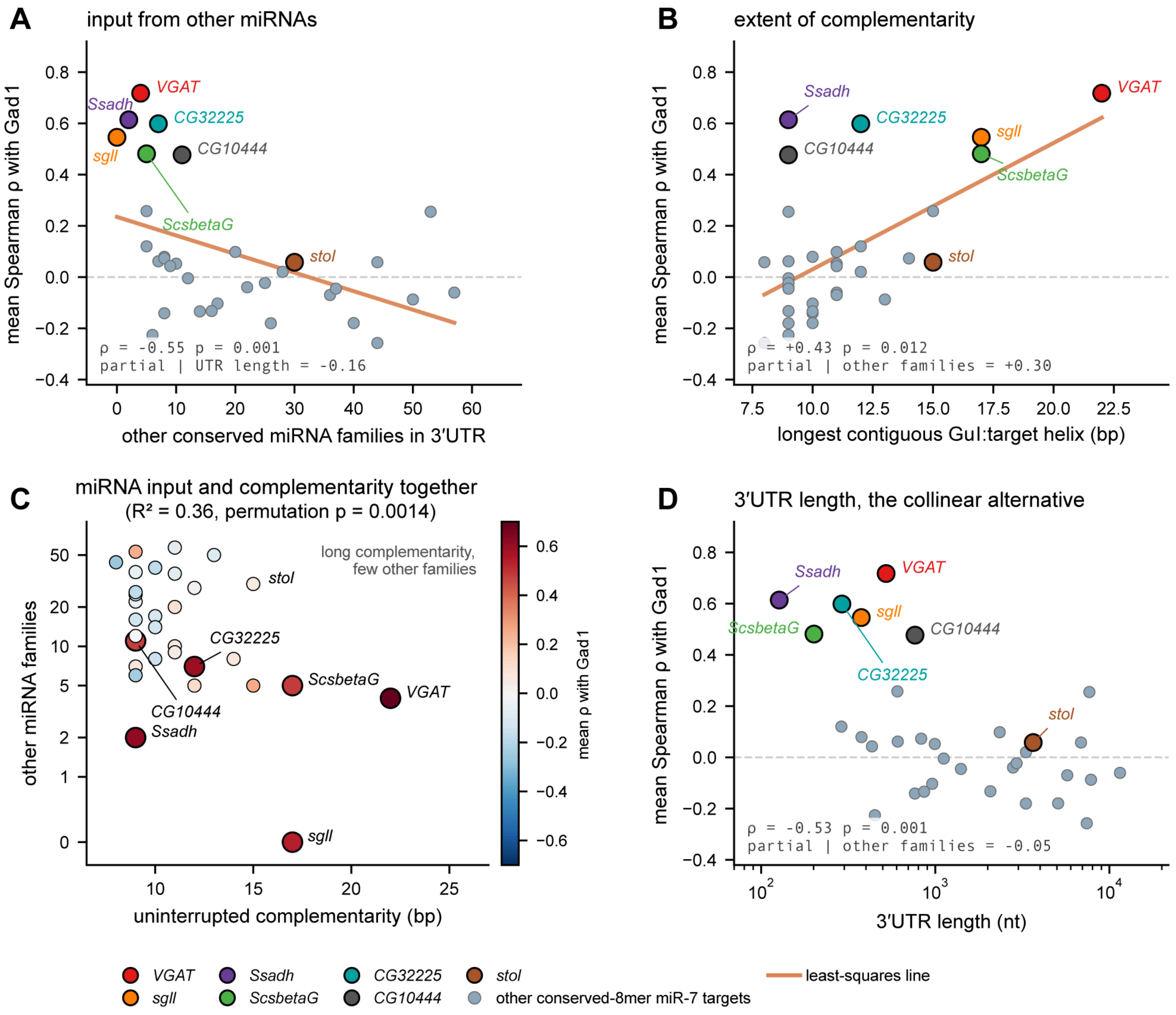


**Figure S13. Two 3′UTR features predict which conserved 8mer targets register a GABAergic difference.** All panels use the 33 conserved miR-7 8mer targets detected in all three CNS regions, plotted against mean Spearman ρ with *Gad1*.

**(A)** Number of other conserved miRNA families, that is, families other than miR-7 with a conserved site in the same 3′UTR (ρ = –0.55, p < 0.001).

**(B)** Length of the longest uninterrupted complementarity to *GuI* across the miR-7 site (ρ = +0.43, p = 0.012).

**(C)** The two features together ($R^2$ = 0.36, permutation p = 0.0014). **The x axis is symlog to include** *sgll*, which has zero other families and a log axis drops it.

**(D)** 3′UTR length, the collinear alternative to (A). *stol* is labeled as the counter-example, carrying the same single conserved 8mer and the same 15 bp of complementarity to *GuI* as *sgll* but a 3,652 nt 3′UTR with 30 other families.

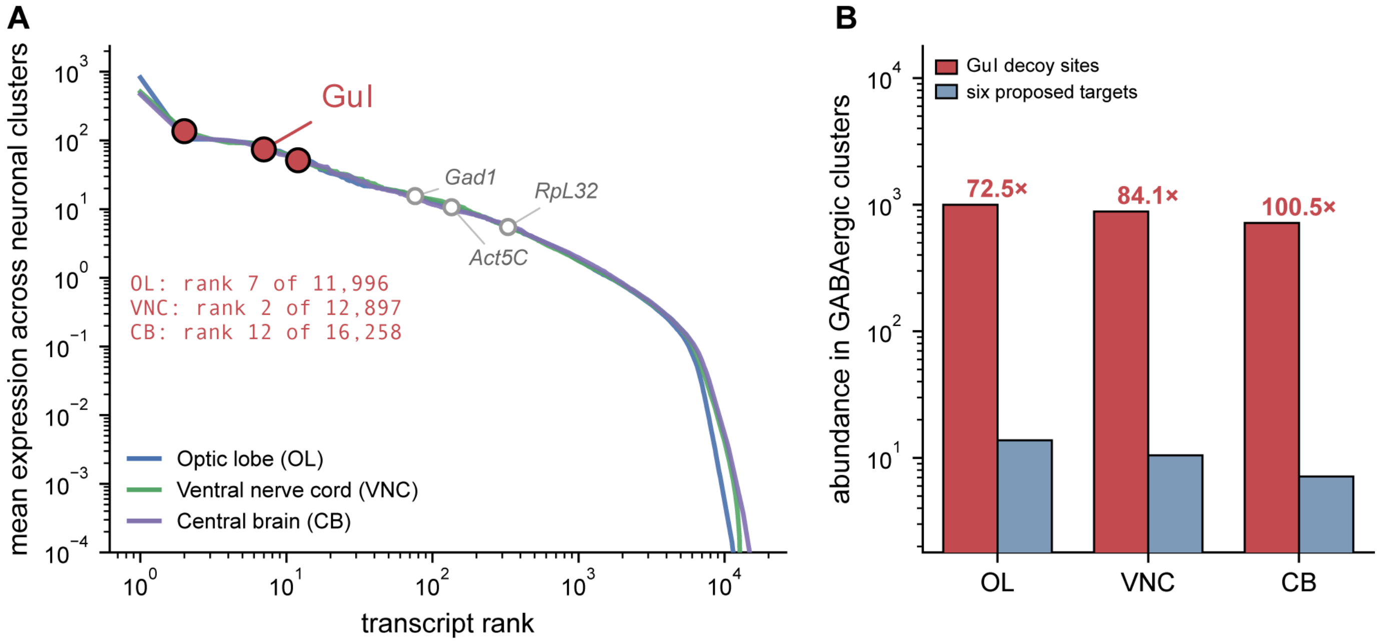


**Figure S14. *Gul* abundance and mimic-site stoichiometry.**

**(A)** Rank-abundance of all detected transcripts in each CNS region, with *Gul* (filled red) and three reference transcripts (open grey) marked. Ranks and totals are given for each region.

**(B)** *Gul* mimic-site abundance against the combined abundance of the six enriched transcripts (*VGAT*, *Ssadh*, *CG32225*, *sgll*, *ScsbetaG* and *CG10444*), in GABAergic clusters, with *Gul* in red and these six in blue. The two bars do not measure the same quantity: the *Gul* bar is its expression multiplied by the three tandem mimic copies, while the target bar is summed transcript abundance, one entry per gene.
