## supplemental tables and scripts for "Long non-coding RNA-3′UTR interactions coordinate GABA signaling in the brain": compute_bls.py

```
#!/usr/bin/env python3
"""
Branch Length Score (BLS) for the key sites, over the Drosophila subtree only.

BLS (Stark/Kheradpour) measures how much of a phylogeny a site is conserved
across, weighted by branch length -- so conservation among many *close* species
(tiny branches) barely counts, while reaching *distant* species counts a lot.
This is the standard metric for short regulatory-motif conservation in flies.

We compute it on the UCSC 27-way alignment but restricted to the 23 Drosophila
species (the 4 distant-insect outgroups -- musDom2/anoGam1/apiMel4/triCas2 -- are
dropped). Rationale: 3'UTR sequence rarely aligns to those outgroups anyway, and
dropping them gives ONE clean 0..1 value that distinguishes "conserved in a few
drosophilids" (low) from "conserved across all drosophilids" (=1) with full
dynamic range, regardless of whether distal insects have the site. Saturation at
1.0 for pan-drosophilid sites is intended.

Method:
  - tree: data/bls/dm6.27way.nh, pruned to the 23 Drosophila (branch lengths of
    collapsed nodes summed).
  - alignment: stream each chromosome's multiz27way MAF, extract the per-species
    columns over each site (cached to data/bls/alignments.json).
  - present: a drosophilid counts as conserved if it aligns over the site with
    >= BLS_MIN_ID identity to dm6 (gaps count as mismatch). dm6 is always present.
  - BLS_dro = (branch length of the minimal subtree spanning the present
    drosophilids) / (total Drosophila-subtree branch length).

Inputs:
  data/sites_coordinates.tsv    the curated sites (7 GABA complements + miR-7 motifs)
  data/bls/dm6.27way.nh         the tree with branch lengths
Output:
  data/bls_scores.tsv           per-site BLS_dro + per-clade presence
"""

import json
import re
import urllib.request
import zlib
from pathlib import Path

import pandas as pd

ROOT = Path(__file__).resolve().parent   # data/ sits beside these scripts
DATA = ROOT / "data"
BLS = DATA / "bls"
TREE = BLS / "dm6.27way.nh"
ALN_CACHE = BLS / "alignments.json"
OUT = DATA / "bls_scores.tsv"

MAF_URL = "https://hgdownload.soe.ucsc.edu/goldenPath/dm6/multiz27way/maf/{chrom}.maf.gz"
OUTGROUPS = {"musDom2", "anoGam1", "apiMel4", "triCas2"}
BLS_MIN_ID = 0.80          # a drosophilid is "conserved" at >= this identity to dm6

# drosophilid clades, roughly outward from D. mel, for the presence readout
CLADES = [
    ("mel_subgroup",  ["droSim1", "droSec1", "droYak3", "droEre2", "droBia2", "droSuz1"]),
    ("mel_group_rest", ["droAna3", "droBip2", "droEug2", "droEle2", "droKik2",
                        "droTak2", "droRho2", "droFic2"]),
    ("obscura",       ["droPse3", "droPer1", "droMir2"]),
    ("willistoni",    ["droWil2"]),
    ("Drosophila_subg", ["droVir3", "droMoj3", "droAlb1", "droGri2"]),
]


# ----------------------------------------------------------------- tree
def parse_newick(s):
    s = re.sub(r"\s+", "", s).rstrip(";")
    pos = 0

    def parse():
        nonlocal pos
        node = {"children": [], "name": None, "bl": 0.0}
        if s[pos] == "(":
            pos += 1
            while True:
                node["children"].append(parse())
                if s[pos] == ",":
                    pos += 1
                    continue
                if s[pos] == ")":
                    pos += 1
                    break
        m = re.match(r"[^,:()]+", s[pos:])
        if m:
            node["name"] = m.group(0)
            pos += len(m.group(0))
        if pos < len(s) and s[pos] == ":":
            pos += 1
            m = re.match(r"[0-9.eE+-]+", s[pos:])
            node["bl"] = float(m.group(0))
            pos += len(m.group(0))
        return node
    return parse()


def leaves(n):
    if not n["children"]:
        return [n["name"]]
    out = []
    for c in n["children"]:
        out += leaves(c)
    return out


def prune(n, drop):
    if not n["children"]:
        return None if n["name"] in drop else dict(n, children=[])
    kids = [prune(c, drop) for c in n["children"]]
    kids = [k for k in kids if k]
    if not kids:
        return None
    if len(kids) == 1:
        k = dict(kids[0])
        k["bl"] += n["bl"]
        return k
    return dict(n, children=kids)


def total_bl(n):
    return sum(c["bl"] + total_bl(c) for c in n["children"])


def spanning_bl(root, present):
    """Sum of edge lengths whose two sides each contain a present leaf."""
    total = 0.0

    def rec(node):
        nonlocal total
        for c in node["children"]:
            cnt = sum(1 for l in leaves(c) if l in present)
            if cnt >= 1 and (len(present) - cnt) >= 1:
                total += c["bl"]
            rec(c)
    rec(root)
    return total


# ----------------------------------------------------------------- MAF
def extract_sites(chrom, targets):
    """targets: list of (key, lo, hi) 1-based inclusive on this chrom.
    Returns {key: {species: aligned_seq_over_site}}.

    A site can straddle a MAF block boundary, so the per-species columns are
    ACCUMULATED per overlapping block and concatenated in genomic order at the
    end -- species missing from a block are padded with gaps so every species
    keeps the same column count. (Assigning per block instead, as this did
    originally, let a later partial block overwrite an earlier one: the site came
    back truncated, species that were absent from the later block silently kept a
    longer fragment, and a 1-base remnant then scored as trivially conserved.
    That inflated ~7-14% of records in every BLS table.)"""
    want = {k: (lo, hi) for k, lo, hi in targets}
    hi_max = max(hi for _, _, hi in targets)
    acc = {k: [] for k in want}          # key -> [(rstart, ncols, {sp: frag}), ...]
    d = zlib.decompressobj(zlib.MAX_WBITS | 16)
    buf, block = "", []

    def handle(block):
        ref = None
        rows = []
        for ln in block:
            if ln.startswith("s "):
                p = ln.split()
                sp = p[1].split(".")[0]
                start, size, seq = int(p[2]), int(p[3]), p[6]
                if sp == "dm6" and ref is None:
                    ref = (start, size, seq)
                rows.append((sp, seq))
        if not ref:
            return
        rstart, rsize, rseq = ref
        for key, (lo, hi) in want.items():
            if rstart > hi or rstart + rsize < lo:
                continue
            cols, g = [], rstart
            for i, ch in enumerate(rseq):
                if ch != "-":
                    if lo <= g + 1 <= hi:      # ref start is 0-based
                        cols.append(i)
                    g += 1
            if not cols:
                continue
            frag = {sp: "".join(seq[i] for i in cols).upper() for sp, seq in rows}
            acc[key].append((rstart, len(cols), frag))

    with urllib.request.urlopen(MAF_URL.format(chrom=chrom), timeout=90) as fh:
        done = False
        while not done:
            chunk = fh.read(1 << 16)
            if not chunk:
                break
            buf += d.decompress(chunk).decode("ascii", "replace")
            while "\n" in buf:
                line, buf = buf.split("\n", 1)
                if line.startswith("a "):
                    if block:
                        handle(block)
                        rs = next((int(l.split()[2]) for l in block
                                   if l.startswith("s dm6.")), 0)
                        if rs > hi_max:
                            done = True
                            break
                    block = [line]
                elif line[:2] in ("s ", "i ", "e ", "q "):
                    block.append(line)
            if done:
                break
        if block and not done:                 # the trailing block, else it is dropped
            handle(block)

    out = {}
    for key, parts in acc.items():
        parts.sort(key=lambda t: t[0])         # genomic order
        species = set()
        for _, _, f in parts:
            species |= set(f)
        out[key] = {sp: "".join(f.get(sp, "-" * n) for _, n, f in parts)
                    for sp in species}
    return out


def load_alignments(sites):
    if ALN_CACHE.exists():
        cached = json.loads(ALN_CACHE.read_text())
        if all(s["key"] in cached for s in sites):
            print(f"alignments: cached ({len(cached)} sites)")
            return cached
    by_chrom = {}
    for s in sites:
        by_chrom.setdefault(s["chrom"], []).append((s["key"], s["lo"], s["hi"]))
    aln = {}
    for chrom, tg in by_chrom.items():
        print(f"streaming {chrom} MAF for {len(tg)} sites (up to {max(h for _,_,h in tg):,}) ...",
              flush=True)
        aln.update(extract_sites(chrom, tg))
    ALN_CACHE.write_text(json.dumps(aln))
    return aln


# ----------------------------------------------------------------- main
def identity(a, b):
    if not a:
        return 0.0
    return sum(x == y for x, y in zip(a, b)) / len(b)


def main():
    tree = parse_newick(TREE.read_text())
    dtree = prune(tree, OUTGROUPS)
    dros = set(leaves(dtree))
    TOT = total_bl(dtree)

    sc = pd.read_csv(DATA / "sites_coordinates.tsv", sep="\t")
    keep = sc[sc.site_type.isin(["gaba_target_complement", "gul_mir7_sense"])].copy()
    sites = []
    for _, r in keep.iterrows():
        sites.append({"key": f"{r.site_type}:{r.gene}:{int(r.genome_start)}",
                      "name": r["name"], "gene": r.gene, "site_type": r.site_type,
                      "chrom": r.chrom, "lo": int(r.genome_start),
                      "hi": int(r.genome_end), "ref": r.sequence})

    aln = load_alignments(sites)

    rows = []
    for s in sites:
        a = aln[s["key"]]
        dm6 = a.get("dm6", s["ref"])
        present = {"dm6"}
        percl = {}
        for sp in dros:
            if sp == "dm6":
                continue
            seq = a.get(sp)
            if seq and identity(seq, dm6) >= BLS_MIN_ID:
                present.add(sp)
        score = spanning_bl(dtree, present) / TOT
        for cname, members in CLADES:
            n_pres = sum(1 for m in members if m in present)
            percl[cname] = f"{n_pres}/{len(members)}"
        rows.append({
            "site_type": s["site_type"], "name": s["name"], "gene": s["gene"],
            "chrom": s["chrom"], "genome_start": s["lo"], "genome_end": s["hi"],
            "ref_seq": dm6, "n_dro_aligned": sum(1 for sp in dros if sp in a and sp != "dm6"),
            "n_dro_conserved": len(present) - 1, "BLS_dro": round(score, 3),
            **percl})

    out = pd.DataFrame(rows)
    out.to_csv(OUT, sep="\t", index=False)
    print(f"\nwrote {OUT.relative_to(ROOT)}  (Drosophila-subtree BL = {TOT:.3f}, "
          f"identity threshold {BLS_MIN_ID:.0%})\n")
    show = out.sort_values("BLS_dro", ascending=False)
    print(show[["name", "n_dro_conserved", "BLS_dro"]
               + [c for c, _ in CLADES]].to_string(index=False))


if __name__ == "__main__":
    main()
```
