## supplemental tables and scripts for "Long non-coding RNA-3′UTR interactions coordinate GABA signaling in the brain": compute_bls_full.py

```
#!/usr/bin/env python3
"""
BLS_dro for ALL 995 panel-A hits (the full Gul full-UTR antisense set), so the
conservation x complexity quadrant can show the whole BLAST group.

Same method/tree as compute_bls.py (BLS over the 23-Drosophila subtree, a species
counts as conserved at >= BLS_MIN_ID identity to dm6 over the site). Target genome
coordinates come from all_sites_coordinates.tsv (already mapped). Streams each
chromosome's multiz27way MAF once, extracting every panel-A site on it; caches
the extracted alignments so re-runs are instant.

Output: data/bls_full.tsv  (rank, gene, bitscore, evalue, at_frac, is_gaba, n_dro_*, BLS_dro)
"""

import json
from pathlib import Path

import pandas as pd

from compute_bls import (parse_newick, prune, leaves, total_bl, spanning_bl,
                         extract_sites, identity, OUTGROUPS, BLS_MIN_ID, TREE)

ROOT = Path(__file__).resolve().parent   # data/ sits beside these scripts
DATA = ROOT / "data"
CACHE = DATA / "bls" / "alignments_full.json"
OUT = DATA / "bls_full.tsv"

GABA = {"VGAT", "P5CDh1", "Gabat", "ScsbetaG", "CG9674", "stol", "sgll"}


def main():
    tree = parse_newick(TREE.read_text())
    dtree = prune(tree, OUTGROUPS)
    dros = set(leaves(dtree))
    TOT = total_bl(dtree)

    a = pd.read_csv(DATA / "all_sites_coordinates.tsv", sep="\t")
    a = a[a.search == "Gul_fullUTR"].copy()
    sites = [{"key": str(int(r.rank)), "gene": r.gene, "rank": int(r.rank),
              "bitscore": r.bitscore, "evalue": r.evalue,
              "length": int(r.length), "at_frac": r.at_frac,
              "chrom": r.target_chrom, "lo": int(r.target_genome_start),
              "hi": int(r.target_genome_end), "ref": r.sequence}
             for r in a.itertuples(index=False)]

    aln = json.loads(CACHE.read_text()) if CACHE.exists() else {}
    todo = [s for s in sites if s["key"] not in aln]
    if todo:
        by_chrom = {}
        for s in todo:
            by_chrom.setdefault(s["chrom"], []).append((s["key"], s["lo"], s["hi"]))
        for chrom, tg in sorted(by_chrom.items()):
            print(f"streaming {chrom} MAF for {len(tg)} sites "
                  f"(up to {max(h for _, _, h in tg):,}) ...", flush=True)
            aln.update(extract_sites(chrom, tg))
            CACHE.write_text(json.dumps(aln))          # checkpoint per chromosome
    else:
        print(f"alignments cached ({len(aln)} sites)")

    rows = []
    for s in sites:
        al = aln.get(s["key"], {})
        dm6 = al.get("dm6", s["ref"])
        present = {"dm6"}
        for sp in dros:
            if sp == "dm6":
                continue
            seq = al.get(sp)
            if seq and identity(seq, dm6) >= BLS_MIN_ID:
                present.add(sp)
        rows.append({"rank": s["rank"], "gene": s["gene"], "bitscore": s["bitscore"],
                     "evalue": s["evalue"], "length": s["length"], "at_frac": s["at_frac"],
                     "is_gaba": s["gene"] in GABA,
                     "n_dro_aligned": sum(1 for sp in dros if sp in al and sp != "dm6"),
                     "n_dro_conserved": len(present) - 1,
                     "BLS_dro": round(spanning_bl(dtree, present) / TOT, 3)})
    out = pd.DataFrame(rows)
    out.to_csv(OUT, sep="\t", index=False)
    g, bg = out[out.is_gaba], out[~out.is_gaba]
    print(f"\nwrote {OUT.relative_to(ROOT)}: {len(out)} hits ({len(g)} GABA, {len(bg)} background)")
    print(f"BLS_dro: GABA median {g.BLS_dro.median():.3f}, background median {bg.BLS_dro.median():.3f}")


if __name__ == "__main__":
    main()
```
