## supplemental tables and scripts for "Long non-coding RNA-3′UTR interactions coordinate GABA signaling in the brain": compute_bls_sense.py

```
#!/usr/bin/env python3
"""
BLS_dro for the top 100 SENSE hits (plus-strand / direct-match) of BOTH queries --
VGAT and Gul -- the companions to compute_bls_vgat.py / compute_bls_full.py, so the
sense matches can be placed in the same conservation x complexity space as the
antisense set. One CONFIG entry per query.

Same method/tree as compute_bls.py (BLS over the 23-Drosophila subtree; a species
counts as conserved at >= BLS_MIN_ID identity to dm6 over the site). Unlike the
antisense BLS scripts, the sense hits are NOT in all_sites_coordinates.tsv (that
table covers the four antisense searches only), so this script maps each hit's
target segment to dm6 coordinates inline from utr_coords.tsv -- exactly the mapping
make_all_sites_table.py uses -- and streams each chromosome's multiz27way MAF once,
caching per query so re-runs are instant.

There is no reciprocal partner on the sense strand (the VGAT<->Gul pair is
antisense-only), so no is_partner column: every hit is background.

Outputs: data/bls_sense_top100.tsv (VGAT), data/bls_gul_sense_top100.tsv (Gul)
         (rank, gene, bitscore, evalue, length, at_frac, n_dro_*, BLS_dro)
"""

import json
from pathlib import Path

import pandas as pd

from compute_bls import (parse_newick, prune, leaves, total_bl, spanning_bl,
                         extract_sites, identity, OUTGROUPS, BLS_MIN_ID, TREE)

ROOT = Path(__file__).resolve().parent   # data/ sits beside these scripts
DATA = ROOT / "data"
COORDS = DATA / "utr_coords.tsv"
TOP_N = 100

# one config per query side; VGAT keeps its original output name (bls_sense_top100.tsv)
CONFIGS = [
    dict(ranked="ranked_VGAT_sense_DUSTnative.csv", out="bls_sense_top100.tsv",
         cache="alignments_sense.json"),
    dict(ranked="ranked_Gul_sense_DUSTnative.csv", out="bls_gul_sense_top100.tsv",
         cache="alignments_gul_sense.json"),
]


def span_to_genome(rec, u_lo, u_hi):
    """UTR positions [u_lo, u_hi] (1-based) -> dm6 [start, end] (1-based incl)."""
    if rec["strand"] == "+":
        g1, g2 = rec["start"] + (u_lo - 1), rec["start"] + (u_hi - 1)
    else:
        g1, g2 = rec["end"] - (u_lo - 1), rec["end"] - (u_hi - 1)
    return min(g1, g2), max(g1, g2)


def compute(cfg, coords, dtree, dros, TOT):
    CACHE = DATA / "bls" / cfg["cache"]
    OUT = DATA / cfg["out"]
    s = pd.read_csv(DATA / cfg["ranked"]).sort_values("rank").head(TOP_N)
    sites, missing = [], 0
    for h in s.itertuples(index=False):
        rec = coords.get(h.sseqid)
        if rec is None:
            missing += 1
            continue
        t_lo, t_hi = min(h.sstart, h.send), max(h.sstart, h.send)
        gs, ge = span_to_genome(rec, t_lo, t_hi)
        sites.append({"key": str(int(h.rank)), "gene": h.symbol, "rank": int(h.rank),
                      "bitscore": h.bitscore, "evalue": h.evalue, "length": int(h.length),
                      "at_frac": h.at_frac, "chrom": rec["chrom"], "lo": gs, "hi": ge,
                      "ref": h.seg})
    if missing:
        print(f"  {missing} hits skipped (no transcript coords)")

    aln = json.loads(CACHE.read_text()) if CACHE.exists() else {}
    todo = [x for x in sites if x["key"] not in aln]
    if todo:
        by_chrom = {}
        for x in todo:
            by_chrom.setdefault(x["chrom"], []).append((x["key"], x["lo"], x["hi"]))
        for chrom, tg in sorted(by_chrom.items()):
            print(f"streaming {chrom} MAF for {len(tg)} sites "
                  f"(up to {max(h for _, _, h in tg):,}) ...", flush=True)
            aln.update(extract_sites(chrom, tg))
            CACHE.write_text(json.dumps(aln))          # checkpoint per chromosome
    else:
        print(f"alignments cached ({len(aln)} sites)")

    rows = []
    for x in sites:
        al = aln.get(x["key"], {})
        dm6 = al.get("dm6", x["ref"])
        present = {"dm6"}
        for sp in dros:
            if sp != "dm6" and al.get(sp) and identity(al[sp], dm6) >= BLS_MIN_ID:
                present.add(sp)
        rows.append({"rank": x["rank"], "gene": x["gene"], "bitscore": x["bitscore"],
                     "evalue": x["evalue"], "length": x["length"], "at_frac": x["at_frac"],
                     "n_dro_aligned": sum(1 for sp in dros if sp in al and sp != "dm6"),
                     "n_dro_conserved": len(present) - 1,
                     "BLS_dro": round(spanning_bl(dtree, present) / TOT, 3)})
    out = pd.DataFrame(rows)
    out.to_csv(OUT, sep="\t", index=False)
    print(f"wrote {OUT.relative_to(ROOT)}: {len(out)} sense hits | "
          f"BLS_dro median {out.BLS_dro.median():.3f}, at_frac median {out.at_frac.median():.3f}\n")


def main():
    tree = parse_newick(TREE.read_text())
    dtree = prune(tree, OUTGROUPS)
    dros = set(leaves(dtree))
    TOT = total_bl(dtree)
    coords = pd.read_csv(COORDS, sep="\t").set_index("transcript_id")
    coords = coords[["chrom", "strand", "start", "end"]].to_dict("index")
    for cfg in CONFIGS:
        print(f"=== {cfg['ranked']} -> {cfg['out']} ===")
        compute(cfg, coords, dtree, dros, TOT)


if __name__ == "__main__":
    main()
```
