## supplemental tables and scripts for "Long non-coding RNA-3′UTR interactions coordinate GABA signaling in the brain": fetch_conservation.py

```
#!/usr/bin/env python3
"""
Fetch conservation for every plotted BLAST hit (Gul query).

An additive layer on top of the existing analysis -- it reads the ranked tables
and writes one new file; nothing else is touched. For each hit it maps the
matched segment in the *target* gene's 3'UTR to dm6 genome coordinates and pulls
per-base scores there from the UCSC REST API, for two tracks:

  phyloP27way    per-base evolutionary rate (signed; + = slower than neutral =
                 conserved) over the 27-way tree -- mostly Drosophila, so it
                 weights recent conservation among close relatives.
  phyloP124way   the same, over the 124-way insect tree (mosquitoes, bees,
                 beetles; ~350 My). A base conserved here is conserved DEEP, in
                 old species, not just among flies. Two summaries are derived:
                   phylop124_mean   mean over the whole matched segment
                   phylop124_core6  best CORE_K-bp window mean -- rewards a
                                    short preserved core even if the flanks are
                                    not conserved. The figure colours by this.
  phastCons27way probability a base is inside a conserved element (0..1). Kept
                 for reference only -- do NOT rank sites by it. It saturates at
                 ~1 for any base in an element, so two sites that both sit in
                 elements read the same and the mean is set by their unconserved
                 flank, which is length-sensitive and mis-orders real sites.

Why phyloP124way core, not phyloP27way mean: a 27-way window mean can call a
site highly conserved when it merely has many bases shared among recent species.
The deep tree + a core statistic emphasise conservation in old species even when
only a short core is preserved -- e.g. it demotes Glts and P5CDh1 (shallow,
Drosophila-only cores) relative to VGAT (cores preserved across insects).

Coordinate mapping (verified: all 30,283 UTRs are single contiguous blocks,
1-based inclusive, so subject-relative positions map to the genome directly):
  + strand:  genome = utr.start + (pos - 1)
  - strand:  genome = utr.end   - (pos - 1)
The UCSC API is 0-based half-open, so a genomic span [gmin, gmax] (1-based
inclusive) is requested as start=gmin-1, end=gmax.

Inputs:
  data/ranked_Gul_antisense_DUSTnative.csv   panel A hits (all 995 plotted)
  data/ranked_Gul_mir7_antisense.csv         panel B hits (bitscore >= 26.3)
  data/utr_coords.tsv   FBtr -> dm6 chrom, strand, start, end

Output:
  data/conservation_gul.tsv   one row per (source, sseqid, sstart, send) with
                              chrom, genomic span, n scored, phylop_mean/max
                              (27-way), phylop124_mean/core6 (deep), and
                              phastcons_mean/min (reference).

Cached: rows already present (keyed by source+sseqid+sstart+send) are not
re-fetched, so re-runs are cheap and interrupted runs resume. A cache written by
an older version is ignored if it lacks the phylop124_core6 column.
"""

import json
import sys
import time
import urllib.error
import urllib.request
from pathlib import Path

import pandas as pd

ROOT = Path(__file__).resolve().parent   # data/ sits beside these scripts
DATA = ROOT / "data"
COORDS = DATA / "utr_coords.tsv"
OUT = DATA / "conservation_gul.tsv"

TRACKS = ["phyloP27way", "phyloP124way", "phastCons27way"]
CORE_K = 6                         # deep-core window: best CORE_K-bp mean
GENOME = "dm6"
PANEL_B_MIN_BITSCORE = 26.3        # bitscore floor for the panel-B hit set
API = "https://api.genome.ucsc.edu/getData/track"
PAUSE = 0.15                       # be polite to the UCSC API
RETRIES = 3

OUT_COLS = ["source", "symbol", "sseqid", "sstart", "send", "chrom",
            "gmin", "gmax", "n_scored",
            "phylop_mean", "phylop_max",           # 27-way (recent-weighted)
            "phylop124_mean", "phylop124_core6",   # 124-way (deep); core = best window
            "phastcons_mean", "phastcons_min"]     # reference only


def load_coords():
    co = pd.read_csv(COORDS, sep="\t").set_index("transcript_id")
    return co[["chrom", "strand", "start", "end"]].to_dict("index")


def to_genome(rec, pos):
    """Subject-UTR position (1-based) -> dm6 coordinate (1-based)."""
    return rec["start"] + (pos - 1) if rec["strand"] == "+" \
        else rec["end"] - (pos - 1)


def fetch_track(chrom, gmin, gmax, track):
    """Per-base track values over [gmin, gmax] (1-based inclusive)."""
    url = (f"{API}?genome={GENOME};track={track};chrom={chrom}"
           f";start={gmin - 1};end={gmax}")
    for attempt in range(RETRIES):
        try:
            with urllib.request.urlopen(url, timeout=45) as fh:
                d = json.load(fh)
            return [x["value"] for x in d.get(track, []) if isinstance(x, dict)]
        except (urllib.error.URLError, TimeoutError, json.JSONDecodeError):
            if attempt == RETRIES - 1:
                raise
            time.sleep(1.5 * (attempt + 1))


def best_core(vals, k=CORE_K):
    """Best k-bp window mean -- the most conserved short core in the segment.
    Direction-independent, so genome vs gene order does not matter."""
    if not vals:
        return float("nan")
    if len(vals) < k:
        return sum(vals) / len(vals)
    return max(sum(vals[i:i + k]) / k for i in range(len(vals) - k + 1))


def hits_to_fetch():
    """The exact set of hits the figure plots, tagged by panel."""
    a = pd.read_csv(DATA / "ranked_Gul_antisense_DUSTnative.csv")
    a = a.assign(source="panelA")
    b = pd.read_csv(DATA / "ranked_Gul_mir7_antisense.csv")
    b = b[b.bitscore >= PANEL_B_MIN_BITSCORE].assign(source="panelB")
    cols = ["source", "symbol", "sseqid", "sstart", "send"]
    return pd.concat([a[cols], b[cols]], ignore_index=True)


def main():
    if not COORDS.exists():
        sys.exit(f"missing coordinate file: {COORDS}")
    coords = load_coords()
    want = hits_to_fetch()

    done = {}
    if OUT.exists():
        prev = pd.read_csv(OUT, sep="\t")
        if "phylop124_core6" not in prev.columns:
            print(f"cache: {OUT.name} predates the deep (124-way) columns -- ignoring, refetching")
        else:
            for _, r in prev.iterrows():
                done[(r.source, r.sseqid, int(r.sstart), int(r.send))] = r
            print(f"cache: {len(done)} hits already fetched in {OUT.name}")

    # genomic-interval memo, so identical intervals are fetched once (per track)
    interval_cache = {}
    rows, n_new, n_err = [], 0, 0
    for i, h in enumerate(want.itertuples(index=False)):
        key = (h.source, h.sseqid, int(h.sstart), int(h.send))
        if key in done:
            rows.append(done[key])
            continue
        rec = coords.get(h.sseqid)
        if rec is None:
            n_err += 1
            continue
        slo, shi = min(h.sstart, h.send), max(h.sstart, h.send)
        g1, g2 = to_genome(rec, slo), to_genome(rec, shi)
        gmin, gmax = min(g1, g2), max(g1, g2)
        ivkey = (rec["chrom"], gmin, gmax)
        try:
            vals = interval_cache.get(ivkey)
            if vals is None:
                vals = {t: fetch_track(rec["chrom"], gmin, gmax, t) for t in TRACKS}
                interval_cache[ivkey] = vals
                time.sleep(PAUSE)
            pp, pp124, pc = (vals["phyloP27way"], vals["phyloP124way"],
                             vals["phastCons27way"])
            f = lambda xs, g: round(g(xs), 4) if xs else float("nan")
            rows.append(pd.Series({
                "source": h.source, "symbol": h.symbol, "sseqid": h.sseqid,
                "sstart": int(h.sstart), "send": int(h.send),
                "chrom": rec["chrom"], "gmin": gmin, "gmax": gmax,
                "n_scored": len(pp),
                "phylop_mean": f(pp, lambda x: sum(x) / len(x)),
                "phylop_max": f(pp, max),
                "phylop124_mean": f(pp124, lambda x: sum(x) / len(x)),
                "phylop124_core6": round(best_core(pp124), 4) if pp124 else float("nan"),
                "phastcons_mean": f(pc, lambda x: sum(x) / len(x)),
                "phastcons_min": f(pc, min)}))
            n_new += 1
        except Exception as e:                       # network hiccup -> skip, resume next run
            print(f"  ! {h.sseqid} {slo}-{shi}: {type(e).__name__} {e}")
            n_err += 1
        if n_new and n_new % 100 == 0:
            print(f"  fetched {n_new} new ({i + 1}/{len(want)}) ...", flush=True)
            pd.DataFrame(rows)[OUT_COLS].to_csv(OUT, sep="\t", index=False)  # checkpoint

    df = pd.DataFrame(rows)[OUT_COLS]
    df.to_csv(OUT, sep="\t", index=False)
    print(f"\nwrote {OUT.name}: {len(df)} hits "
          f"({n_new} newly fetched, {n_err} skipped)")
    print(f"phyloP124 core6: {df.phylop124_core6.min():.2f}-{df.phylop124_core6.max():.2f}, "
          f"median {df.phylop124_core6.median():.2f}  (deep-core; the figure colours by this)")
    print(f"phyloP124 mean:  {df.phylop124_mean.min():.2f}-{df.phylop124_mean.max():.2f}, "
          f"median {df.phylop124_mean.median():.2f}  (deep window)")
    print(f"phyloP27 mean:   {df.phylop_mean.min():.2f}-{df.phylop_mean.max():.2f}, "
          f"median {df.phylop_mean.median():.2f}  (recent-weighted)")


if __name__ == "__main__":
    main()
```
