## supplemental tables and scripts for "Long non-coding RNA-3′UTR interactions coordinate GABA signaling in the brain": make_all_sites_table.py

```
#!/usr/bin/env python3
"""
Genome coordinates for every hit (= site) in the analysis.

Each ranked table is one row per gene's best antisense complement. This adds dm6
genome coordinates to all of them, for BOTH ends of the duplex:

  target_*  the matched segment in the target gene's 3'UTR (the subject)
  query_*   the paired segment on the query UTR (Gul or VGAT)

covering all four searches:
  Gul_fullUTR       ranked_Gul_antisense_DUSTnative.csv    (995)
  VGAT_fullUTR      ranked_VGAT_antisense_DUSTnative.csv   (985)
  Gul_mir7region    ranked_Gul_mir7_antisense.csv          (500)
  VGAT_mir7region   ranked_VGAT_mir7region_antisense.csv  (1007)

Coordinates are 1-based inclusive (matching 00_inputs/utr_coords.tsv). The
mapping is the one used for the conservation layer and site table, verified there
by round-tripping the genomic sequence. Pure offline computation -- no network.
For the curated key sites (miR-7 motifs + the 8 manuscript complements) see
make_sites_table.py / sites_coordinates.tsv; this is the exhaustive per-hit view.

Output:
  data/all_sites_coordinates.tsv
"""

from pathlib import Path

import pandas as pd

ROOT = Path(__file__).resolve().parent   # data/ sits beside these scripts
DATA, INP = ROOT / "data", ROOT / "data" / "inputs"
COORDS = DATA / "utr_coords.tsv"
OUT = DATA / "all_sites_coordinates.tsv"

GUL_FBTR, VGAT_FBTR = "FBtr0308363", "FBtr0087577"

# per search: the ranked table, the query transcript, and the offset that turns
# a query-local position into a position in that transcript's full UTR
SEARCHES = {
    "Gul_fullUTR":     ("ranked_Gul_antisense_DUSTnative.csv",  GUL_FBTR,  0),
    "VGAT_fullUTR":    ("ranked_VGAT_antisense_DUSTnative.csv", VGAT_FBTR, 0),
    "Gul_mir7region":  ("ranked_Gul_mir7_antisense.csv",        GUL_FBTR,  1137),
    "VGAT_mir7region": ("ranked_VGAT_mir7region_antisense.csv", VGAT_FBTR, 47),
}

OUT_COLS = ["search", "rank", "gene", "target_fbtr", "sequence", "length",
            "pident", "bitscore", "evalue", "at_frac",
            "target_utr_start", "target_utr_end", "target_strand",
            "target_chrom", "target_genome_start", "target_genome_end",
            "query_utr_start", "query_utr_end", "query_strand",
            "query_chrom", "query_genome_start", "query_genome_end",
            "phylop124_core6", "phylop124_mean", "phylop_mean", "phastcons_mean"]


def span_to_genome(rec, u_lo, u_hi):
    """UTR positions [u_lo, u_hi] (1-based) -> dm6 [start, end] (1-based incl)."""
    if rec["strand"] == "+":
        g1, g2 = rec["start"] + (u_lo - 1), rec["start"] + (u_hi - 1)
    else:
        g1, g2 = rec["end"] - (u_lo - 1), rec["end"] - (u_hi - 1)
    return min(g1, g2), max(g1, g2)


def main():
    coords = pd.read_csv(COORDS, sep="\t").set_index("transcript_id")
    coords = coords[["chrom", "strand", "start", "end"]].to_dict("index")

    # conservation was only fetched for the plotted Gul-query hits; join where present
    cons = pd.read_csv(DATA / "conservation_gul.tsv", sep="\t")
    cons_by = {
        "Gul_fullUTR": cons[cons.source == "panelA"],
        "Gul_mir7region": cons[cons.source == "panelB"],
    }

    rows, missing = [], 0
    for search, (fname, q_fbtr, q_off) in SEARCHES.items():
        df = pd.read_csv(DATA / fname)
        qrec = coords[q_fbtr]
        cons_tbl = cons_by.get(search)
        for h in df.itertuples(index=False):
            trec = coords.get(h.sseqid)
            if trec is None:
                missing += 1
                continue
            # target (subject) side
            t_lo, t_hi = min(h.sstart, h.send), max(h.sstart, h.send)
            tgs, tge = span_to_genome(trec, t_lo, t_hi)
            # query side: query-local -> query full-UTR position -> genome
            q_lo, q_hi = h.qstart + q_off, h.qend + q_off
            q_lo, q_hi = min(q_lo, q_hi), max(q_lo, q_hi)
            qgs, qge = span_to_genome(qrec, q_lo, q_hi)
            # conservation, if this hit was fetched
            dc = dm = pp = pc = None
            if cons_tbl is not None:
                c = cons_tbl[(cons_tbl.sseqid == h.sseqid)
                             & (cons_tbl.sstart == h.sstart)
                             & (cons_tbl.send == h.send)]
                if len(c):
                    row = c.iloc[0]
                    dc, dm = row.phylop124_core6, row.phylop124_mean
                    pp, pc = row.phylop_mean, row.phastcons_mean
            rows.append({
                "search": search, "rank": int(h.rank), "gene": h.symbol,
                "target_fbtr": h.sseqid, "sequence": h.seg, "length": int(h.length),
                "pident": h.pident, "bitscore": h.bitscore, "evalue": h.evalue,
                "at_frac": getattr(h, "at_frac", None),
                "target_utr_start": t_lo, "target_utr_end": t_hi,
                "target_strand": trec["strand"], "target_chrom": trec["chrom"],
                "target_genome_start": tgs, "target_genome_end": tge,
                "query_utr_start": q_lo, "query_utr_end": q_hi,
                "query_strand": qrec["strand"], "query_chrom": qrec["chrom"],
                "query_genome_start": qgs, "query_genome_end": qge,
                "phylop124_core6": dc, "phylop124_mean": dm,
                "phylop_mean": pp, "phastcons_mean": pc})

    out = pd.DataFrame(rows)[OUT_COLS]
    out.to_csv(OUT, sep="\t", index=False)
    print(f"wrote {OUT.relative_to(ROOT)}: {len(out)} sites"
          + (f" ({missing} skipped, no coords)" if missing else ""))
    for s in SEARCHES:
        n = (out.search == s).sum()
        nc = out[(out.search == s)].phylop_mean.notna().sum()
        print(f"  {s:16s} {n:4d} sites"
              + (f", {nc} with conservation" if nc else ""))


if __name__ == "__main__":
    main()
```
