## supplemental tables and scripts for "Long non-coding RNA-3′UTR interactions coordinate GABA signaling in the brain": make_sites_table.py

```
#!/usr/bin/env python3
"""
Assemble one coordinate table for every "site" in the analysis.

Three kinds of site, tagged by `site_type`:

  gaba_target_complement  the best antisense complement to the full Gul 3'UTR in
                          each of the seven manuscript GABA-pathway genes (the
                          BLAST hits highlighted in the figures). These live in
                          the *target* gene's UTR.
  gul_mir7_sense          the three GGAAGACA miR-7 sense sites in Gul's own UTR.
  vgat_mir7_complement    the three GTCTTCC complement cores in VGAT's UTR; the
                          one that is the full 8-mer TGTCTTCC is flagged.

Every row carries UTR-relative coordinates AND dm6 genome coordinates (1-based
inclusive, matching 00_inputs/utr_coords.tsv), the sequence, and — where it was
already fetched — mean phyloP/phastCons over the site. The genome mapping is the
same one used for the conservation layer and was checked by pulling the genomic
sequence back and matching it to the site base-for-base (re-verified here).

Inputs:
  data/ranked_Gul_antisense_DUSTnative.csv   the 7 GABA target hits
  data/conservation_gul.tsv                  their genome coords + conservation
  data/inputs/3utr_goi.fa                     Gul + VGAT query UTRs
  data/utr_coords.tsv   FBtr -> dm6 chrom, strand, start, end

Output:
  data/sites_coordinates.tsv
"""

from pathlib import Path

import pandas as pd

ROOT = Path(__file__).resolve().parent   # data/ sits beside these scripts
DATA, INP = ROOT / "data", ROOT / "data" / "inputs"
COORDS = DATA / "utr_coords.tsv"
OUT = DATA / "sites_coordinates.tsv"

# the two query UTRs resolve to exactly one transcriptome isoform each
GUL_FBTR, VGAT_FBTR = "FBtr0308363", "FBtr0087577"
SENSE, CORE, CORE_FULL = "GGAAGACA", "GTCTTCC", "TGTCTTCC"

GABA_DISPLAY = {
    "VGAT": "VGAT", "P5CDh1": "P5CDh1", "Gabat": "GABA-T", "ScsbetaG": "ScsβG",
    "CG9674": "Glts", "stol": "stol", "sgll": "sgll",
}

COLUMNS = ["site_type", "name", "gene", "fbtr", "sequence", "length",
           "utr_start", "utr_end", "strand", "chrom", "genome_start",
           "genome_end", "bitscore", "evalue", "phylop124_core6", "phylop124_mean",
           "phylop_mean", "phastcons_mean", "note"]


def read_fasta(path):
    seqs, k = {}, None
    for line in open(path):
        line = line.strip()
        if line.startswith(">"):
            k = line[1:].split()[0]
            seqs[k] = ""
        elif line:
            seqs[k] += line
    return seqs


def find_all(seq, motif):
    out, i = [], seq.find(motif)
    while i != -1:
        out.append(i + 1)                       # 1-based
        i = seq.find(motif, i + 1)
    return out


def utr_span_to_genome(rec, u_lo, u_hi):
    """UTR positions [u_lo, u_hi] (1-based) -> dm6 [gstart, gend] (1-based incl)."""
    if rec["strand"] == "+":
        g1, g2 = rec["start"] + (u_lo - 1), rec["start"] + (u_hi - 1)
    else:
        g1, g2 = rec["end"] - (u_lo - 1), rec["end"] - (u_hi - 1)
    return min(g1, g2), max(g1, g2)


def main():
    coords = pd.read_csv(COORDS, sep="\t").set_index("transcript_id")
    coords = coords[["chrom", "strand", "start", "end"]].to_dict("index")
    goi = read_fasta(INP / "3utr_goi.fa")
    rows = []

    # ---- 1. GABA-pathway target complement sites (the 7 BLAST hits) ---------
    hits = pd.read_csv(DATA / "ranked_Gul_antisense_DUSTnative.csv")
    cons = pd.read_csv(DATA / "conservation_gul.tsv", sep="\t")
    cons = cons[cons.source == "panelA"]
    for sym, disp in GABA_DISPLAY.items():
        h = hits[hits.symbol == sym].iloc[0]
        rec = coords[h.sseqid]
        u_lo, u_hi = min(h.sstart, h.send), max(h.sstart, h.send)
        gs, ge = utr_span_to_genome(rec, u_lo, u_hi)
        c = cons[(cons.sseqid == h.sseqid) & (cons.sstart == h.sstart)
                 & (cons.send == h.send)]
        cr = c.iloc[0] if len(c) else None
        rows.append({
            "site_type": "gaba_target_complement",
            "name": f"{disp} complement to Gul", "gene": disp, "fbtr": h.sseqid,
            "sequence": h.seg, "length": int(h.length),
            "utr_start": u_lo, "utr_end": u_hi, "strand": rec["strand"],
            "chrom": rec["chrom"], "genome_start": gs, "genome_end": ge,
            "bitscore": h.bitscore, "evalue": h.evalue,
            "phylop124_core6": None if cr is None else cr.phylop124_core6,
            "phylop124_mean": None if cr is None else cr.phylop124_mean,
            "phylop_mean": None if cr is None else cr.phylop_mean,
            "phastcons_mean": None if cr is None else cr.phastcons_mean,
            "note": "query seg is the Gul-side sequence; antisense (minus strand)"})

    # ---- 2. Gul miR-7 sense sites (in Gul's own UTR) ------------------------
    gul, rec = goi["Gul"], coords[GUL_FBTR]
    for i, s in enumerate(find_all(gul, SENSE), 1):
        e = s + len(SENSE) - 1
        gs, ge = utr_span_to_genome(rec, s, e)
        rows.append({
            "site_type": "gul_mir7_sense", "name": f"Gul miR-7 sense site {i}",
            "gene": "Gul", "fbtr": GUL_FBTR, "sequence": SENSE, "length": len(SENSE),
            "utr_start": s, "utr_end": e, "strand": rec["strand"],
            "chrom": rec["chrom"], "genome_start": gs, "genome_end": ge,
            "bitscore": None, "evalue": None,
            "phylop124_core6": None, "phylop124_mean": None,
            "phylop_mean": None, "phastcons_mean": None,
            "note": f"{SENSE} motif in Gul UTR"})

    # ---- 3. VGAT miR-7 complement cores (in VGAT's UTR) ---------------------
    vgat, rec = goi["VGAT"], coords[VGAT_FBTR]
    full_starts = set(find_all(vgat, CORE_FULL))
    for i, s in enumerate(find_all(vgat, CORE), 1):
        e = s + len(CORE) - 1
        gs, ge = utr_span_to_genome(rec, s, e)
        is_full = (s - 1) in full_starts          # TGTCTTCC starts one base 5'
        rows.append({
            "site_type": "vgat_mir7_complement",
            "name": f"VGAT complement core {i}" + (" (8-mer)" if is_full else ""),
            "gene": "VGAT", "fbtr": VGAT_FBTR, "sequence": CORE, "length": len(CORE),
            "utr_start": s, "utr_end": e, "strand": rec["strand"],
            "chrom": rec["chrom"], "genome_start": gs, "genome_end": ge,
            "bitscore": None, "evalue": None,
            "phylop124_core6": None, "phylop124_mean": None,
            "phylop_mean": None, "phastcons_mean": None,
            "note": f"{CORE} core; full {CORE_FULL} 8-mer" if is_full
                    else f"{CORE} core (7-mer; non-matching flank)"})

    df = pd.DataFrame(rows)[COLUMNS]
    df.to_csv(OUT, sep="\t", index=False)
    print(f"wrote {OUT.relative_to(ROOT)}: {len(df)} sites")
    for t, n in df.site_type.value_counts().items():
        print(f"  {t}: {n}")
    with pd.option_context("display.width", 200, "display.max_columns", None):
        print()
        print(df[["site_type", "name", "gene", "utr_start", "utr_end",
                  "chrom", "genome_start", "genome_end", "bitscore",
                  "phylop124_core6"]].to_string(index=False))


if __name__ == "__main__":
    main()
```
