## supplemental tables and scripts for "Long non-coding RNA-3′UTR interactions coordinate GABA signaling in the brain": run_analysis.py

```
#!/usr/bin/env python3
"""Regenerate the BLAST specificity analysis from primary inputs.

    blastn + dustmasker  ->  ranked gene tables

Four searches, all blastn-short -evalue 1000 -word_size 7 -dust no:
  [1] both queries (Gul, VGAT) vs the transcriptome, then the dustmasker level-20
      hard filter (drop any hit whose matched segment touches a masked interval)
  [2] the Gul miR-7 sense-site cluster vs the transcriptome
  [3] the VGAT miR-7 complement region vs the transcriptome
  [4] the VGAT region vs the Gul UTR alone -- pairwise, at a permissive evalue,
      so short site-to-site pairings are reported rather than cut off

Inputs (data/inputs/):
  3utr_goi.fa             query 3'UTRs: VGAT (522 bp), Gul (2045 bp)
  gul_mir7.fa             110 bp miR-7 sense-site cluster, Gul UTR 1138..1247
  VGAT_mir7_region.fa     197 bp miR-7 complement region, VGAT UTR 48..244
  transcriptome_3utr.fa   30,283 FlyBase FBtr 3'UTRs (BLAST db; built on demand)
  fbtr_symbol_map.tsv     FBtr -> FBgn, gene symbol, description

Outputs (data/), ranked tables one row per gene (best antisense hit):
  dust_intervals_L20.txt                dustmasker intervals on the queries
  blast_raw_strand.tsv                  [1] raw hits: sstrand, coords, qseq, BTOP
  blast_mir7_btop.tsv                   [2] raw hits with BTOP
  blast_vgat_region_btop.tsv            [3] raw hits with BTOP
  blast_vgat_region_vs_gul.tsv          [4] every pairwise HSP
  gul_utr.fa                            Gul UTR alone, the subject for [4]
  ranked_Gul_antisense_DUSTnative.csv   [1] Gul query, antisense
  ranked_VGAT_antisense_DUSTnative.csv  [1] VGAT query, antisense
  ranked_Gul_sense_DUSTnative.csv       [1] Gul query, plus strand (sense control)
  ranked_VGAT_sense_DUSTnative.csv      [1] VGAT query, plus strand (sense control)
  ranked_Gul_mir7_antisense.csv         [2] antisense
  ranked_Gul_mir7_sense.csv             [2] plus strand
  ranked_VGAT_mir7region_antisense.csv  [3] antisense

at_frac column (diagnostic only; nothing filters or ranks on it). blastn-short
scores as if all four bases were equally likely, but the 3'UTR db is 62.9% A+T, so
AT-rich matches are cheaper by chance than their bitscore implies, and DUST cannot
see this (it scores triplet repetition, not composition). Each ranked table carries
at_frac, the A+T fraction of the matched query segment; compare it against the 62.9%
background printed at run time. The seven manuscript genes sit at 53-75% A+T (at
background) while the 995-gene background runs to a median of 82%.

Tie handling: hits are sorted (bitscore desc, qstart asc, sseqid asc) so the output
is deterministic. Within a block of hits sharing a bitscore the rank order is
arbitrary in the data, so ranks inside a tie block need not match any earlier CSV;
everything that is not a tie is reproduced exactly, including all seven GABA-pathway
genes.
"""

import re
import subprocess
import sys
from collections import Counter
from pathlib import Path

import pandas as pd

ROOT = Path(__file__).resolve().parent   # data/ sits beside these scripts
INP, DATA = ROOT / "data" / "inputs", ROOT / "data"

DB = INP / "transcriptome_3utr.fa"
QUERIES, MIR7_Q = INP / "3utr_goi.fa", INP / "gul_mir7.fa"
VGAT_REGION = INP / "VGAT_mir7_region.fa"
SYMBOL_MAP = INP / "fbtr_symbol_map.tsv"

REGION_START, REGION_END = 1138, 1247      # miR-7 cluster within the Gul UTR
DUST_LEVEL = 20

# for the VGAT-region-vs-Gul pairwise: report every HSP, however weak, so the
# short site-to-site pairings are visible rather than silently cut off
PAIRWISE_EVALUE = "10000"

# self-hits to exclude: the query's own transcript(s)
SELF_HITS = {
    "VGAT": {"FBtr0087577"},
    "Gul": {"FBtr0308363", "FBtr0308361", "FBtr0308362", "FBtr0345137"},
}
SELF_HITS["Gul_mir7"] = SELF_HITS["Gul"]

BLAST_FMT = ("qseqid sseqid pident length mismatch gapopen qstart qend sstart "
             "send evalue bitscore sstrand")
BLAST_OPTS = ["-task", "blastn-short", "-evalue", "1000", "-word_size", "7",
              "-dust", "no", "-soft_masking", "false",
              "-max_target_seqs", "40000"]

# ordering used for every tie: makes the output deterministic
SORT_BY = ["bitscore", "qstart", "sseqid"]
SORT_ASC = [False, True, True]


def run(cmd, **kw):
    print("  $", " ".join(str(c) for c in cmd[:6]), "...", flush=True)
    return subprocess.run(cmd, check=True, **kw)


def ensure_db():
    if DB.with_suffix(".fa.nsq").exists():
        return
    print("building BLAST db (indexes absent)")
    run(["makeblastdb", "-in", str(DB), "-dbtype", "nucl"],
        stdout=subprocess.DEVNULL)


def blastn(query, out, extra_fields="", subject=None, evalue=None):
    """Run blastn against the db, or against `subject` for a pairwise search."""
    fmt = f"6 {BLAST_FMT}{(' ' + extra_fields) if extra_fields else ''}"
    opts = [o for o in BLAST_OPTS]
    if evalue:
        opts[opts.index("-evalue") + 1] = evalue
    if subject:                       # -max_target_seqs is db-only
        i = opts.index("-max_target_seqs")
        del opts[i:i + 2]
        target = ["-subject", str(subject)]
    else:
        target = ["-db", str(DB)]
    with open(out, "w") as fh:
        run(["blastn", "-query", str(query), *target, *opts, "-outfmt", fmt],
            stdout=fh)
    cols = fmt.split()[1:]
    return pd.read_csv(out, sep="\t", names=cols)


def dust_intervals(fasta, out):
    """dustmasker -outfmt interval -> {query: [(start, end), ...]} 1-based."""
    with open(out, "w") as fh:
        run(["dustmasker", "-in", str(fasta), "-level", str(DUST_LEVEL),
             "-outfmt", "interval"], stdout=fh)
    iv, cur = {}, None
    for line in open(out):
        line = line.strip()
        if line.startswith(">"):
            cur = line[1:].strip()
            iv[cur] = []
        elif line:
            a, b = re.split(r"\s*-\s*", line)
            iv[cur].append((int(a) + 1, int(b) + 1))   # dustmasker is 0-based
    return iv


def lcfrac(iv, q, start, end):
    """Fraction of a matched query segment inside a low-complexity interval."""
    masked = sum(max(0, min(end, b) - max(start, a) + 1) for a, b in iv.get(q, []))
    return masked / (end - start + 1)


def base_freqs(fasta):
    """Base composition of the whole 3'UTR db -- the background blastn-short
    does NOT use (it scores as if all four bases were equally likely)."""
    n = Counter()
    with open(fasta) as fh:
        for line in fh:
            if not line.startswith(">"):
                n.update(line.strip().upper())
    tot = sum(n[c] for c in "ACGT")
    return {c: n[c] / tot for c in "ACGT"}


def rank_genes(hits, query, symbols, strand="minus"):
    """Best hit per gene on one strand, ranked. Self-hits removed.
    strand='minus' = antisense complement (the base-pairing hits, default);
    strand='plus'  = sense / same-orientation direct matches (supplement control)."""
    a = hits[(hits.qseqid == query) & (hits.sstrand == strand)
             & (~hits.sseqid.isin(SELF_HITS[query]))].copy()
    a = a.merge(symbols, left_on="sseqid", right_on="FBtr", how="left")
    unmapped = a.gene_symbol.isna().sum()
    assert unmapped == 0, f"{unmapped} hits did not map to a gene symbol"

    a = a.sort_values(SORT_BY, ascending=SORT_ASC, kind="stable")
    best = a.drop_duplicates("gene_symbol").reset_index(drop=True)
    best["rank"] = range(1, len(best) + 1)
    best["n_tied_at_bitscore"] = best.bitscore.map(best.bitscore.value_counts())
    best = best.rename(columns={"gene_symbol": "symbol", "qseq": "seg"})

    # A+T fraction of the matched segment -- reported beside the bitscore,
    # never used to rank or filter. Compare against the db background (62.9%).
    best["at_frac"] = best.seg.map(
        lambda s: round(sum(c in "AT" for c in s) / len(s), 3))
    return best


OUT_COLS = ["rank", "symbol", "FBgn", "sseqid", "sstrand", "pident", "length",
            "mismatch", "qstart", "qend", "sstart", "send", "seg", "evalue",
            "bitscore", "at_frac", "n_tied_at_bitscore", "gene_description"]


def main():
    for f in (DB, QUERIES, MIR7_Q, SYMBOL_MAP):
        if not f.exists():
            sys.exit(f"missing input: {f}")
    ensure_db()
    symbols = pd.read_csv(SYMBOL_MAP, sep="\t")[
        ["FBtr", "FBgn", "gene_symbol", "gene_description"]]
    freqs = base_freqs(DB)
    print(f"db base composition: A+T = {freqs['A'] + freqs['T']:.1%} "
          f"(blastn-short scores as if it were 50%) -- compare at_frac to this")

    # the miR-7 query must be exactly the stated slice of the Gul UTR
    fa = {}
    for line in open(QUERIES):
        if line.startswith(">"):
            k = line[1:].strip(); fa[k] = ""
        else:
            fa[k] += line.strip()
    mir7 = "".join(l.strip() for l in open(MIR7_Q) if not l.startswith(">"))
    assert fa["Gul"][REGION_START - 1:REGION_END] == mir7, \
        f"gul_mir7.fa is not Gul[{REGION_START}:{REGION_END}]"
    print(f"queries: VGAT {len(fa['VGAT'])} bp, Gul {len(fa['Gul'])} bp, "
          f"miR-7 region {len(mir7)} bp (= Gul {REGION_START}..{REGION_END})")

    # ---- panel A: full UTR, dustmasker-native hard filter -----------------
    print("\n[1/4] full-UTR BLAST + dustmasker filter")
    # BTOP is carried alongside qseq so the same file supplies both the query
    # segments (ranking/at_frac) and the mismatch positions (fig_vgat_panelA ticks)
    full = blastn(QUERIES, DATA / "blast_raw_strand.tsv", extra_fields="qseq btop")
    iv = dust_intervals(QUERIES, DATA / "dust_intervals_L20.txt")
    full["lcfrac_L20"] = [lcfrac(iv, r.qseqid, r.qstart, r.qend)
                          for r in full.itertuples()]
    kept = full[full.lcfrac_L20 == 0]        # drop any hit touching a masked interval
    print(f"  {len(full)} hits -> {len(kept)} after hard dust filter (level {DUST_LEVEL})")

    dust_cols = OUT_COLS[:12] + ["lcfrac_L20"] + OUT_COLS[12:]
    for query, out in [("Gul", "ranked_Gul_antisense_DUSTnative.csv"),
                       ("VGAT", "ranked_VGAT_antisense_DUSTnative.csv")]:
        t = rank_genes(kept, query, symbols)
        t[dust_cols].to_csv(DATA / out, index=False)
        best = t.iloc[0]
        print(f"  {out}: {len(t)} genes | rank 1 = {best.symbol} "
              f"(bitscore {best.bitscore}, {int(best.n_tied_at_bitscore)} tied)")

    # sense-strand companions (supplement control): each gene's best PLUS-strand
    # (direct, same-orientation) hit, same DUST filter. Nothing tops bitscore ~36 --
    # no reciprocal miR-7 partner -- so the VGAT<->Gul pairing is shown to be
    # antisense-specific, not generic sequence similarity. One per query.
    for query, out in [("VGAT", "ranked_VGAT_sense_DUSTnative.csv"),
                       ("Gul", "ranked_Gul_sense_DUSTnative.csv")]:
        ts = rank_genes(kept, query, symbols, strand="plus")
        ts[dust_cols].to_csv(DATA / out, index=False)
        print(f"  {out}: {len(ts)} genes | rank 1 = {ts.iloc[0].symbol} "
              f"(bitscore {ts.iloc[0].bitscore}; top antisense was 44.1)")

    # ---- panel B: miR-7 region --------------------------------------------
    print("\n[2/4] Gul miR-7 region BLAST")
    mir = blastn(MIR7_Q, DATA / "blast_mir7_btop.tsv", extra_fields="btop qseq sseq")
    reg = rank_genes(mir, "Gul_mir7", symbols)
    reg[OUT_COLS].to_csv(DATA / "ranked_Gul_mir7_antisense.csv", index=False)
    print(f"  ranked_Gul_mir7_antisense.csv: {len(reg)} genes")
    print(f"  genes at bitscore >= 26.3 (panel B): {(reg.bitscore >= 26.3).sum()}")
    # sense-strand companion for the miR-7 region (supplement zoom): direct matches
    # to the 110 bp cluster, both strands shown together in the combined figure.
    reg_s = rank_genes(mir, "Gul_mir7", symbols, strand="plus")
    reg_s[OUT_COLS].to_csv(DATA / "ranked_Gul_mir7_sense.csv", index=False)
    print(f"  ranked_Gul_mir7_sense.csv: {len(reg_s)} genes "
          f"(rank 1 = {reg_s.iloc[0].symbol}, bitscore {reg_s.iloc[0].bitscore})")

    # ---- the VGAT-side mirror: its miR-7 complement region ----------------
    # dustmasker calls no intervals here (the only VGAT interval is 478-484,
    # outside the region), so no complexity filter is applied -- same as Gul's.
    print("\n[3/4] VGAT miR-7 complement region vs transcriptome")
    vreg = blastn(VGAT_REGION, DATA / "blast_vgat_region_btop.tsv",
                  extra_fields="btop qseq sseq")
    vr = rank_genes(vreg, "VGAT", symbols)
    vr[OUT_COLS].to_csv(DATA / "ranked_VGAT_mir7region_antisense.csv", index=False)
    print(f"  ranked_VGAT_mir7region_antisense.csv: {len(vr)} genes | "
          f"rank 1 = {vr.iloc[0].symbol} (bitscore {vr.iloc[0].bitscore})")
    print(f"  background floor: {(vr.bitscore <= 22.3).sum()} of {len(vr)} genes "
          f"sit at bitscore <= 22.3")

    # ---- pairwise: VGAT region vs the Gul UTR alone, every HSP ------------
    # This is what shows whether Gul's *other* sense sites pair with VGAT at
    # all. They are far too short to survive a transcriptome-wide search, so
    # they only become visible in a two-sequence comparison.
    print("\n[4/4] VGAT miR-7 region vs Gul UTR (pairwise, every HSP)")
    gul_fa = DATA / "gul_utr.fa"
    gul_fa.write_text(f">Gul\n{fa['Gul']}\n")
    pw = blastn(VGAT_REGION, DATA / "blast_vgat_region_vs_gul.tsv",
                extra_fields="btop qseq sseq", subject=gul_fa,
                evalue=PAIRWISE_EVALUE)
    anti = pw[pw.sstrand == "minus"]
    print(f"  {len(pw)} HSPs ({len(anti)} antisense), "
          f"bitscore {anti.bitscore.min()}-{anti.bitscore.max()}")

    print("\ndone -- ranked tables written to data/")


if __name__ == "__main__":
    main()
```
