## supplemental tables and scripts for "Long non-coding RNA-3′UTR interactions coordinate GABA signaling in the brain": build_rbp_list.py

```
#!/usr/bin/env python3
"""Build the RNA-binding protein gene list: GO:0003723 plus its molecular_function
descendants, from the FlyBase GO annotation file.

GO:0003723 is propagated to its molecular_function descendants (equivalent to GO.db's
GOMFOFFSPRING over the FlyBase GAF). Propagation is not optional: GO:0003723 has 163
molecular_function descendants, and direct annotations alone give 507 genes against
1,160 with the subtree -- most RBPs are annotated to a child term such as mRNA binding
(GO:0003729), not to the parent.

Both source files are downloaded from the GO Consortium and cached next to the output.
They are dated releases, so the list is reproducible only against the fetch date recorded
in the output header; re-running later may legitimately give a different count.

Usage:
    python3 build_rbp_list.py                      # writes rbp_go0003723.{tsv,csv}
    python3 build_rbp_list.py --keep-cache         # keep fb.gaf.gz / go-basic.obo
"""
import argparse, collections, gzip, os, urllib.request

import pandas as pd

import paths as P

GAF_URL = "https://current.geneontology.org/annotations/fb.gaf.gz"
OBO_URL = "https://current.geneontology.org/ontology/go-basic.obo"
ROOT = "GO:0003723"
OUT = "rbp_go0003723.tsv"


def fetch(url, dest):
    if os.path.exists(dest):
        print(f"  cached {os.path.basename(dest)}")
        return dest
    print(f"  downloading {url}")
    # the GO Consortium CDN 403s the default python-urllib User-Agent
    req = urllib.request.Request(url, headers={"User-Agent": "curl/8.0"})
    with urllib.request.urlopen(req, timeout=300) as r, open(dest, "wb") as fh:
        while chunk := r.read(1 << 20):
            fh.write(chunk)
    return dest


def parse_obo(path):
    """{GO id: term} from go-basic.obo, plus {alt_id: primary id}."""
    terms, cur = {}, None

    def flush(c):
        if c and c.get("id"):
            terms[c["id"]] = c

    for line in open(path):
        line = line.rstrip("\n")
        if line.startswith("["):
            flush(cur)
            cur = ({"id": None, "ns": None, "parents": [], "obsolete": False, "alt": []}
                   if line == "[Term]" else None)
        elif cur is not None:
            if line.startswith("id: GO:"):
                cur["id"] = line[4:]
            elif line.startswith("alt_id: GO:"):
                cur["alt"].append(line[8:])
            elif line.startswith("namespace: "):
                cur["ns"] = line[11:]
            elif line.startswith("is_a: GO:"):
                cur["parents"].append(line[6:16])
            elif line.startswith("relationship: part_of GO:"):
                cur["parents"].append(line[22:32])
            elif line.startswith("is_obsolete: true"):
                cur["obsolete"] = True
    flush(cur)
    alt = {a: t["id"] for t in terms.values() for a in t["alt"]}
    return terms, alt


def descendants(terms, root):
    """`root` and everything reachable from it by is_a / part_of, child-ward."""
    kids = collections.defaultdict(list)
    for t in terms.values():
        for p in t["parents"]:
            kids[p].append(t["id"])
    seen, stack = {root}, [root]
    while stack:
        for k in kids[stack.pop()]:
            if k not in seen:
                seen.add(k)
                stack.append(k)
    return seen


def coding_genes(datadir=None):
    """FBgns with an mRNA or CDS feature in the FlyBase r6.68 GTF, and all gene FBgns.

    GO:0003723 is "RNA binding", and plenty of *RNAs* bind RNA: the subtree pulls in 290
    tRNA genes, 93 miRNA genes, and assorted snRNA / snoRNA / scaRNA loci -- 37% of the
    raw list. Those are not RNA-binding proteins, they are not captured by polyA 10x
    anyway, and leaving them in inflates the denominator the paper quotes. Filtering on
    coding status rather than on symbol prefixes catches classes nobody thought to name.
    """
    coding, allg = set(), set()
    with gzip.open(P.find("dmel-all-r6.68.gtf.gz", datadir), "rt", errors="replace") as fh:
        for line in fh:
            if line.startswith("#"):
                continue
            f = line.split("\t")
            if len(f) < 9:
                continue
            i = f[8].find('gene_id "')
            if i < 0:
                continue
            fbgn = f[8][i + 9:f[8].find('"', i + 9)]
            allg.add(fbgn)
            if f[2] in ("mRNA", "CDS"):
                coding.add(fbgn)
    return coding, allg


def run(cache_dir=None, keep_cache=False):
    cache = cache_dir or os.path.join(P.ROOT, "data", "genome")
    os.makedirs(cache, exist_ok=True)
    gaf = fetch(GAF_URL, os.path.join(cache, "fb.gaf.gz"))
    obo = fetch(OBO_URL, os.path.join(cache, "go-basic.obo"))

    terms, alt = parse_obo(obo)
    seen = descendants(terms, ROOT)
    mf = {t for t in seen
          if terms.get(t, {}).get("ns") == "molecular_function" and not terms[t]["obsolete"]}
    print(f"  ontology: {len(terms):,} terms; {ROOT} + descendants = {len(seen)} "
          f"({len(mf)} non-obsolete molecular_function)")

    # Keyed on FBgn, not symbol. Symbols drift between FlyBase releases and the three
    # expression matrices were built against different ones; FBgn is stable, and the CG
    # annotation ID carried in the GAF synonym field is nearly as stable and is what the
    # matrices use for uncharacterised genes. Emitting all three lets the screen match on
    # identity rather than on string similarity -- which is what keeps DCP1 (decapping
    # protein 1, FBgn0034921, a real RBP) apart from Dcp-1 (Death caspase-1, FBgn0010501,
    # not one). They differ only in punctuation as symbols.
    genes, direct, skipped = {}, set(), set()
    n = 0
    with gzip.open(gaf, "rt", errors="replace") as fh:
        for line in fh:
            if line.startswith("!"):
                continue
            f = line.rstrip("\n").split("\t")
            n += 1
            # aspect F = molecular function; a NOT qualifier asserts the gene does *not*
            # have the activity, so those annotations must not add the gene to the list
            if len(f) < 11 or f[8] != "F" or "NOT" in f[3]:
                continue
            go = alt.get(f[4], f[4])
            # the FlyBase GAF also carries Complex Portal objects (CPX-*: RISC, RNA Pol II,
            # the exosome, NELF...). They are complexes, not genes, can never match a row in
            # a gene-by-cluster matrix, and would silently inflate the list size the paper
            # quotes -- 21 of them under this GO subtree.
            if not f[1].startswith("FBgn"):
                skipped.add(f[1])
                continue
            if go in seen:
                fbgn, sym = f[1], f[2]
                cg = next((x for x in f[10].split("|") if x.startswith("CG")), "")
                rec = genes.setdefault(fbgn, [sym, cg, set()])
                rec[2].add(go)          # keep the specific terms, not just membership
                if go == ROOT:
                    direct.add(fbgn)
    print(f"  GAF: {n:,} annotation lines")
    print(f"  direct {ROOT} only: {len(direct)} genes")
    print(f"  with descendants  : {len(genes)} genes")
    print(f"  with a CG annotation ID: {sum(1 for v in genes.values() if v[1])}")
    print(f"  non-gene objects skipped (Complex Portal etc.): {len(skipped)}")

    coding, allg = coding_genes()
    absent = {g for g in genes if g not in allg}
    noncoding = {g for g in genes if g in allg and g not in coding}
    for g in noncoding | absent:
        del genes[g]
    print(f"  dropped {len(noncoding)} non-coding RNA genes (tRNA / miRNA / snRNA / "
          f"snoRNA / scaRNA loci -- RNAs that bind RNA, not RNA-binding proteins)")
    print(f"  dropped {len(absent)} genes absent from the r6.68 GTF")
    print(f"  protein-coding RBPs: {len(genes)}")

    out = os.path.join(cache, OUT)
    with open(out, "w") as fh:
        fh.write(f"# RNA-binding proteins: {ROOT} 'RNA binding' plus all "
                 f"molecular_function descendants\n"
                 f"# source: {GAF_URL}\n#         {OBO_URL}\n"
                 f"# aspect F; NOT-qualified annotations excluded\n"
                 f"# {len(seen)} GO terms -> {len(genes)} protein-coding genes\n"
                 f"# non-coding RNA genes annotated with these terms (tRNA, miRNA, snRNA,\n"
                 f"# snoRNA, scaRNA loci) are excluded: they bind RNA but are not RBPs\n"
                 f"# regenerate with scripts/build_rbp_list.py\n"
                 f"# match on FBgn or CG; `symbol` is for display only\n")
        fh.write("FBgn\tsymbol\tCG\tGO_terms\n")
        for fbgn in sorted(genes, key=lambda k: genes[k][0]):
            sym, cg, gos = genes[fbgn]
            fh.write(f"{fbgn}\t{sym}\t{cg}\t{';'.join(sorted(gos))}\n")
    print(f"  wrote {out}")

    # Same content as a plain CSV for sharing and for opening in a spreadsheet: header
    # row only, no comment lines, since Excel renders those as data rows. The .tsv above
    # stays the canonical input because it carries the provenance header.
    out_csv = os.path.splitext(out)[0] + ".csv"
    pd.DataFrame([(fb, genes[fb][0], genes[fb][1], ";".join(sorted(genes[fb][2])))
                  for fb in sorted(genes, key=lambda k: genes[k][0])],
                 columns=["FBgn", "symbol", "CG", "GO_terms"]).to_csv(out_csv, index=False)
    print(f"  wrote {out_csv}")

    if not keep_cache:
        for p in (gaf, obo):
            os.remove(p)
        print("  removed the cached GAF and OBO (--keep-cache to retain)")
    return genes


if __name__ == "__main__":
    ap = argparse.ArgumentParser(description=__doc__,
                                 formatter_class=argparse.RawDescriptionHelpFormatter)
    ap.add_argument("--cache-dir", default=None)
    ap.add_argument("--keep-cache", action="store_true")
    a = ap.parse_args()
    run(a.cache_dir, a.keep_cache)
```
