## supplemental tables and scripts for "Long non-coding RNA-3′UTR interactions coordinate GABA signaling in the brain": extract_matrix.R

```
#!/usr/bin/env Rscript
# ---------------------------------------------------------------------------
# extract_matrix.R
# Run ONCE on the HPC (Torch) inside an interactive, Duo-authenticated session.
#
#   module load r        # or activate the conda env that has Seurat
#   Rscript extract_matrix.R <rds> [cluster_column] [out_csv] [subset_expr]
#
#   Rscript extract_matrix.R Adult.rds                       # MultiomeAnnotated
#   Rscript extract_matrix.R meta_head.rds RNA_snn_res.20 meanexpr_head.csv
#   Rscript extract_matrix.R meta_head.rds RNA_snn_res.20 meanexpr_cb.csv \
#           'experiment == "Allen2025" & region != "optic lobe"'
#
# subset_expr is evaluated against the object metadata, so it can filter on experiment,
# tissue, region, cell class or anything else in o[[]]. It is optional but it is the
# step that was invisible before: meta_head holds the whole head, so the central brain
# matrix required removing the optic lobe, and nothing on disk recorded that. The number
# of cells before and after is printed and belongs in the job log.
#
# The cluster column is an argument because the objects do not agree on one: the optic
# lobe matrix was built on MultiomeAnnotated (named cell types), while the VNC and CB
# matrices carry bare numeric cluster IDs, i.e. a resolution column. Hardcoding it meant
# editing the script per object, which is how the three matrices ended up on different
# conventions without anything recording it.
#
# Produces four files, stemmed on the output name (third argument):
#   <name>.csv            AverageExpression: mean(expm1(x)), linear scale
#   <name>_logmean.csv    mean of the log-normalised values
#   <name>_pctexpr.csv    fraction of cells with non-zero counts
#   <name>_clusters.csv   per-cluster cell counts and metadata composition
# The three matrices are different statistics and none is derivable from the others.
# It will not overwrite an existing file -- pass a distinct name per object.
# Bring that CSV back to the analysis directory; the Python scripts in this
# whole miR-7 correlation test locally.
#
# What it does: collapses the per-cell log-normalized RNA expression to a
# mean-normalized-expression value per (gene, cluster) --
# the same per-cluster summary the manual neset_plotting script used, applied
# to ALL genes at once. AverageExpression on the 'data' layer un-logs, averages
# in linear space, and returns that per cluster.
# ---------------------------------------------------------------------------

suppressMessages({
  library(Seurat)
})

args <- commandArgs(trailingOnly = TRUE)
if (length(args) < 1) stop("Usage: Rscript extract_matrix.R <rds> [cluster_column] [out_csv]")
rds_path  <- args[[1]]
group_col <- if (length(args) >= 2 && nzchar(args[[2]])) args[[2]] else "MultiomeAnnotated"
out_csv   <- if (length(args) >= 3 && nzchar(args[[3]])) args[[3]] else
             "gene_by_cluster_meanexpr.csv"
subset_expr <- if (length(args) >= 4) args[[4]] else ""
if (file.exists(out_csv))
  stop("Refusing to overwrite ", out_csv, " -- give a different name as the third argument.")
cat("Cluster column:", group_col, "\nOutput:", out_csv, "\n")
if (nzchar(subset_expr)) cat("Subset:", subset_expr, "\n")

cat("Reading:", rds_path, "\n"); flush.console()
if (!file.exists(rds_path)) stop("File not found: ", rds_path)

o <- readRDS(rds_path)
cat("Loaded object of class:", class(o)[1], "\n")
DefaultAssay(o) <- "RNA"

# Optional metadata filter. Evaluated with `with()` against o[[]] rather than through
# Seurat's non-standard evaluation, so any expression over the metadata columns works and
# a typo fails loudly instead of silently keeping everything.
if (nzchar(subset_expr)) {
  md <- o[[]]
  keep <- tryCatch(with(md, eval(parse(text = subset_expr))),
                   error = function(e) stop("subset_expr failed: ", conditionMessage(e),
                                            "\nAvailable columns: ",
                                            paste(colnames(md), collapse = ", ")))
  if (!is.logical(keep)) stop("subset_expr must evaluate to TRUE/FALSE per cell.")
  keep[is.na(keep)] <- FALSE
  cat("Subsetting:", sum(keep), "of", length(keep), "cells kept\n")
  if (!sum(keep)) stop("subset_expr kept no cells.")
  o <- subset(o, cells = rownames(md)[keep])
}

# confirm the cluster annotation column exists
md_cols <- colnames(o[[]])
if (!(group_col %in% md_cols)) {
  cat("\nAvailable metadata columns:\n"); print(md_cols)
  stop("Column '", group_col, "' not found in object metadata (see list above).")
}
n_clusters <- length(unique(o[[]][[group_col]]))
cat("Cluster column '", group_col, "' has ", n_clusters, " levels.\n", sep = "")

# ---------------------------------------------------------------------------
# All three statistics in one pass, by sparse matrix multiply against a cell x cluster
# indicator. Computed here rather than via AverageExpression so the Seurat object can be
# dropped first: holding the object, the expression matrix and per-cluster column subsets
# at once was killed by the OOM reaper at 32G on the full head object.
#
#   linear   mean(expm1(x))   what AverageExpression(layer="data") returns
#   logmean  mean(x)          on the log-normalised values
#   pctexpr  mean(x > 0)      detection rate
#
# expm1() keeps sparsity (expm1(0) == 0), so nothing is densified until the result.
# ---------------------------------------------------------------------------
cat("Extracting the data layer...\n"); flush.console()
dat <- GetAssayData(o, layer = "data")
md  <- o[[]]
rm(o); gc(verbose = FALSE)
cat("Matrix:", nrow(dat), "genes x", ncol(dat), "cells\n")

gf <- factor(as.character(md[[group_col]]))
# Seurat prefixes "g" when cluster labels start with a digit; match that, so these outputs
# stay comparable with the matrices already in data/expression/ (which carry g0, g1, ...).
lev <- levels(gf)
# Seurat mangles cluster labels two ways, and both must be reproduced exactly or the
# matrix cannot be joined to the ones already in data/expression/. It replaces "_" with
# "-", and it prefixes "g" to labels that START WITH A DIGIT -- only those. An earlier
# version prefixed every label whenever any one of them started with a digit, which turned
# the whole Desplan optic lobe into gDm4, gG_Astro, ... and made a matrix of the same 254
# clusters share zero column names with the one it was meant to match.
lev <- gsub("_", "-", lev, fixed = TRUE)
n_pre <- sum(grepl("^[0-9]", lev))
if (n_pre) cat("Prefixing 'g' to", n_pre, "labels that start with a digit, as Seurat does.\n")
lev <- ifelse(grepl("^[0-9]", lev), paste0("g", lev), lev)
ind <- Matrix::sparse.model.matrix(~ 0 + gf)
colnames(ind) <- lev
n_cells <- Matrix::colSums(ind)

per_cluster <- function(x) {
  out <- as.matrix(x %*% ind)
  out <- sweep(out, 2, n_cells, "/")
  rownames(out) <- rownames(dat); colnames(out) <- lev
  out
}
cat("Computing mean(expm1) ...\n"); flush.console()
m <- per_cluster(expm1(dat))
cat("Computing mean(log) ...\n"); flush.console()
lm_ <- per_cluster(dat)
cat("Computing detection rate ...\n"); flush.console()
pct <- per_cluster(as(dat > 0, "dMatrix"))
gc(verbose = FALSE)

cat("Matrix:", nrow(m), "genes x", ncol(m), "clusters\n")
cat("CG14989 (Gul) present:", "CG14989" %in% rownames(m), "\n")
for (g in c("VGAT","Gad1","repo","CG32032","AnxB9","GstE12","elav","nSyb")) {
  cat("  ", g, ":", g %in% rownames(m), "\n")
}

log_csv <- sub("\\.csv$", "_logmean.csv", out_csv)
write.csv(lm_, log_csv); cat("WROTE:", normalizePath(log_csv), "\n")
pct_csv <- sub("\\.csv$", "_pctexpr.csv", out_csv)
write.csv(round(pct, 5), pct_csv); cat("WROTE:", normalizePath(pct_csv), "\n")
rm(lm_, pct); gc(verbose = FALSE)

# ---------------------------------------------------------------------------
# Companion table: what each cluster is made of. Without it a cluster-mean matrix cannot
# be filtered after the fact -- there is no way to tell whether a cluster is one region or
# a mixture of two, and a mixed cluster's mean cannot be unmixed. Export everything once,
# subset locally against this.
# ---------------------------------------------------------------------------
raw  <- levels(gf)
info <- data.frame(cluster = lev, n_cells = as.integer(n_cells[lev]),
                   stringsAsFactors = FALSE)
for (col in colnames(md)) {
  v <- md[[col]]
  if (!(is.character(v) || is.factor(v))) next
  v <- as.character(v)
  if (length(unique(v)) > 50 || col == group_col) next
  tab  <- table(factor(as.character(md[[group_col]]), levels = raw), v)
  info[[paste0(col, "__top")]]    <- colnames(tab)[max.col(tab, ties.method = "first")]
  info[[paste0(col, "__purity")]] <- round(as.numeric(
    apply(tab, 1, function(r) if (sum(r)) max(r) / sum(r) else NA_real_)), 3)
}

info_csv <- sub("\\.csv$", "_clusters.csv", out_csv)
write.csv(info, info_csv, row.names = FALSE)
cat("WROTE:", normalizePath(info_csv), " (", nrow(info), "clusters x", ncol(info), "columns )\n")
cat("Cluster purity by column (fraction of clusters that are >=95% one value):\n")
for (col in grep("__purity$", colnames(info), value = TRUE))
  cat("   ", sub("__purity$", "", col), ":", round(mean(info[[col]] >= 0.95), 2), "\n")

write.csv(m, out_csv)
cat("\nWROTE:", normalizePath(out_csv), "\n")
cat("Copy this file to the analysis directory alongside the Python scripts.\n")
```
