## supplemental tables and scripts for "Long non-coding RNA-3′UTR interactions coordinate GABA signaling in the brain": fig_class_vs_six.py

```
#!/usr/bin/env python3
"""The conserved miR-7 target classes are flat; the six proposed targets are not.

One argument on two independent readouts of the same gene classes:

  top row     Spearman rho with Gad1          (co-variation)
  bottom row  log2 fold change, GABAergic vs other neurons   (expression level)

The six are pulled out of the 8mer column into their own; left inside it they
carry the stratum's mean, so the 8mer column would read as elevated only because
of them. Split out, the 8mer column means the class excluding the proposed
targets, and the six are visibly the exception rather than hidden in it.

CIRCULARITY: no p is drawn over the "six" column, and none should be. They were
selected for sitting at the top of the Gad1 ranking, so testing them against the
rest of their own class is circular, and fold change is close enough to rho that
the same objection carries. The tested claims are the four class columns against
background; the tail claim belongs to fig_gad1_enrichment, which needs neither a
held-out set nor a choice of summary statistic.

Genes are the intersection of both analyses, so a gene contributes to both rows
or to neither. Glia excluded throughout.

Usage:
    python3 fig_class_vs_six.py            # 3 regions, 2 rows
    python3 fig_class_vs_six.py --tissues OL
"""
import argparse
import numpy as np, pandas as pd
import matplotlib
matplotlib.use("Agg")
import matplotlib.pyplot as plt
from matplotlib.lines import Line2D
from matplotlib.transforms import blended_transform_factory

import paths as P
import genes as GN
import gul_mir7_correlation as G

SIX = list(GN.TARGET_POOL)
COLS = ["background", "7mer-A1", "7mer-m8", "8mer", "six"]
LABEL = {"background": "no\nsite", "7mer-A1": "7mer\nA1", "7mer-m8": "7mer\nm8",
         "8mer": "other", "six": "enriched"}
COLOR = {"background": "#8A8A8A", "7mer-A1": "#9ECAE1", "7mer-m8": "#4C72B0",
         "8mer": "#08306B", "six": "#C44E52"}
# "the six" is project shorthand and means nothing to a reader. The axis label names what
# the group IS -- the transcripts this paper proposes Gul acts on -- rather than why they sit
# high, which would restate the ranking the panel is plotting and be circular. Kept to two
# The alternatives, so they are not retried blind. "Gul targets" asserts in the figure the
# very thing the figure is evidence for. A GABA-worded label read as "GABA pathway" is wrong
# for CG10444, which para 39 places outside the module. "proposed" was the earlier choice
# here. The live objection to "enriched" is that it restates the y axis, rho with Gad1, so
# the column is named by its own outcome -- which is why the LEGEND entry below, and the
# caption, must define the set rather than leaving the axis label to do it.
#
# Width is the binding constraint: the five columns get only ~28 pt each, so both lines must
# stay short. "proposed" fits only because the neighbour was cut from "8mer (others)" to
# "other 8mer"; at 8 pt with the old neighbour the two labels touched.
LEGEND = {"background": "no annotated miR-7 site", "7mer-A1": "conserved 7mer-A1",
          "7mer-m8": "conserved 7mer-m8",
          "8mer": "conserved 8mer, excluding the six proposed targets",
          "six": "the six proposed $\\it{Gul}$ targets"}
ROWS = [("rho", "Spearman ρ with $\\it{Gad1}$", (-0.75, 1.05), 0.0),
        ("lfc", "log$_2$ fold change\nGABAergic / other", (-1.7, 3.15), 0.0)]


def build(datadir=None):
    R = pd.read_csv(P.find("gad1_correlation_by_sitetype_pergene.csv", datadir))
    L = pd.read_csv(P.find("gaba_lfc_per_gene.csv", datadir))
    D = R.merge(L[["tissue", "gene", "lfc"]], on=["tissue", "gene"], how="inner")
    D["col"] = np.where(D.gene.isin(SIX), "six", D.stratum)
    return D[D.col.isin(COLS)]


def run(tissues=None, outdir=None, datadir=None, rows=None, panel=False,
        panel_h=2.33, panel_w=3.61):
    def O(n): return P.out(n, outdir)
    cols = COLS
    plt.rcParams.update({
        "figure.dpi": 110, "savefig.dpi": 200,
        "axes.spines.top": False, "axes.spines.right": False,
        "svg.fonttype": "none", "font.family": "sans-serif",
        "font.sans-serif": ["Arial"]})

    D = build(datadir)
    order = tissues or ["OL", "VNC", "CB"]
    n = len(order)
    use = [r for r in ROWS if r[0] in (rows or [r0[0] for r0 in ROWS])]
    fig, axes = plt.subplots(len(use), n,
                             figsize=((panel_w * n, panel_h * len(use)) if panel
                                      else (3.5 * n + 0.8, 3.5 * len(use) + 0.4)),
                             squeeze=False)
    rows_out = []

    for r, (val, ylab, ylim, ref) in enumerate(use):
        for c, tag in enumerate(order):
            ax = axes[r][c]
            d = D[D.tissue == tag]
            bg = d[d.col == "background"][val].values
            ax.axhline(np.median(bg), color="0.45", lw=0.9, ls=":", zorder=2)
            ax.axhline(ref, color="0.85", lw=0.8, ls="--", zorder=1)
            for i, col in enumerate(cols):
                sub = d[d.col == col]
                v = sub[val].values
                if not len(v): continue
                big = col == "background"
                if col == "six":
                    # per-gene colours from genes.COLORS, so a reader can carry the
                    # same colour between this panel and fig_gad1_enrichment.
                    g_ = sub.sort_values(val, ascending=False)
                    # spread wide enough that markers clear each other: five of
                    # the six sit within 0.15 of each other on rho, so vertical
                    # separation cannot do the work.
                    xs = np.linspace(i - 0.34, i + 0.30, len(g_))
                    lo, hi = float(np.min(v)), float(np.max(v))
                    span = max(hi - lo, 0.24 * (ylim[1] - ylim[0]))
                    mid = 0.5 * (hi + lo)
                    top = min(mid + 0.72 * span, ylim[1] * 0.86)
                    bot = max(mid - 0.72 * span, ylim[0] + 0.06 * (ylim[1] - ylim[0]))
                    yls = np.linspace(top, bot, len(g_))
                    for (gene, yv), x, yl in zip(zip(g_.gene, g_[val]), xs, yls):
                        cc = GN.COLORS.get(gene, COLOR["six"])
                        ax.plot([x], [yv], "o", ms=6.2, mfc=cc, mec="white",
                                mew=0.7, zorder=7)
                        ax.annotate(f"$\\it{{{gene}}}$", (x, yv), xycoords="data",
                                    xytext=(len(cols) - 0.48, yl), textcoords="data",
                                    fontsize=7.2, color=cc, va="center", ha="left",
                                    zorder=8,
                                    arrowprops=dict(arrowstyle="-", color=cc,
                                                    lw=0.6, shrinkA=1, shrinkB=3))
                else:
                    jit = (np.random.default_rng(i).random(len(v)) - 0.5) * 0.34
                    ax.scatter(np.full(len(v), i) + jit, v,
                               s=(5 if big else 15),
                               alpha=(0.10 if big else 0.55),
                               color=COLOR[col], edgecolor="none",
                               zorder=(3 if big else 5), rasterized=big)
                ax.plot([i - 0.31, i + 0.31], [np.median(v)] * 2,
                        color="black", lw=2.0, zorder=6)
                if col in ("background", "six"):
                    p = np.nan            # see the circularity note in the docstring
                else:
                    _, p = G.perm_test(v, bg)
                if np.isfinite(p):
                    ax.text(i, ylim[1] * 0.90, f"p={p:.2f}", ha="center",
                            fontsize=7.2, color="0.25")
                ax.text(i, ylim[1] * 1.00, f"{len(v)}", ha="center", fontsize=6.8,
                        color="0.45")
                rows_out.append(dict(tissue=tag, measure=val, column=col, n=len(v),
                                     median=np.median(v), perm_p=p))
            ax.set_xticks(range(len(cols)))
            # 6.9, measured not guessed: the column pitch is 26.0 pt, and at 7.2 pt
            # "other" and "proposed" leave a 0.9 pt gap, which renders as touching.
            # 6.9 pt with "other" rather than "others" opens it to ~3.5 pt.
            ax.set_xticklabels([LABEL[c_] for c_ in cols], fontsize=6.9)
            # A bracket under the last two columns naming both as conserved 8mer. Without
            # it the panel reads as though the proposed targets were a site class of their
            # own, when they are a subset pulled out of the 8mer column -- which is the
            # whole point of the split. It also frees the column labels to be one short
            # word each, since "8mer" no longer has to appear twice.
            if "8mer" in cols and "six" in cols:
                i0, i1 = cols.index("8mer"), cols.index("six")
                tr = blended_transform_factory(ax.transData, ax.transAxes)
                yb, tick = -0.175, 0.020
                ax.plot([i0 - 0.42, i1 + 0.42], [yb, yb], transform=tr, color="0.35",
                        lw=0.9, clip_on=False, zorder=7)
                for xx in (i0 - 0.42, i1 + 0.42):
                    ax.plot([xx, xx], [yb, yb + tick], transform=tr, color="0.35",
                            lw=0.9, clip_on=False, zorder=7)
                ax.text((i0 + i1) / 2.0, yb - 0.028, "conserved 8mer", transform=tr,
                        ha="center", va="top", fontsize=7.2, color="0.25")
            ax.set_xlim(-0.6, len(cols) + 0.30); ax.set_ylim(*ylim)
            ax.tick_params(axis="y", labelsize=8)
            if c == 0: ax.set_ylabel(ylab, fontsize=9.5)
            if r == 0 and not panel:
                ax.set_title(f"{G.TISSUE_NAME.get(tag, tag)} ({tag})", fontsize=10.5)

    nrow = len(use)
    if not panel:
        h = [Line2D([], [], marker="o", ls="none",
                    mfc=("white" if c_ == "six" else COLOR[c_]),
                    mec=("0.35" if c_ == "six" else "none"),
                    mew=(0.9 if c_ == "six" else 0), ms=7,
                    label=(LEGEND[c_] + " (coloured per gene)" if c_ == "six"
                           else LEGEND[c_])) for c_ in cols]
        h.append(Line2D([], [], color="0.45", lw=0.9, ls=":", label="background median"))
        fig.legend(handles=h, frameon=False, ncol=(3 if n > 1 else 2),
                   loc="lower center",
                   bbox_to_anchor=(0.5, 0.055 if nrow == 1 else -0.05), fontsize=8.5)
        fig.suptitle("Neither miR-7 site class is elevated in GABAergic neurons; the six are",
                     fontsize=11.5)
        foot = ("p, one-sided permutation vs the no-site background "
                "(20,000 iterations, difference of medians). No p is shown "
                "for the six:\nthey were selected for sitting at the top of "
                "this ranking, so testing them against their own class would "
                "be circular.")
        fig.text(0.5, 0.010 if nrow == 1 else 0.015, foot, ha="center",
                 fontsize=7.4, color="0.35")
    if panel:
        fig.tight_layout(pad=0.4)
    else:
        fig.tight_layout(rect=[0, 0.20 if nrow == 1 else 0.08, 1,
                               0.90 if nrow == 1 else 0.94])
    suffix = ("" if not tissues else "_" + "".join(order)) + \
             ("" if not rows else "_" + "".join(rows)) + ("_panel" if panel else "")
    path = O(f"fig_class_vs_six{suffix}.png")
    fig.savefig(path, bbox_inches="tight")
    fig.savefig(path.replace(".png", ".svg"), bbox_inches="tight")
    plt.close(fig)

    T = pd.DataFrame(rows_out)
    T.to_csv(O(f"class_vs_six{suffix}.csv"), index=False)
    print(T.pivot_table(index=["measure", "column"], columns="tissue",
                        values=["median", "perm_p"])
           .to_string(float_format=lambda v: f"{v:.3f}"))
    return T


if __name__ == "__main__":
    ap = argparse.ArgumentParser(description=__doc__,
                                 formatter_class=argparse.RawDescriptionHelpFormatter)
    ap.add_argument("--tissues", nargs="+", default=None)
    ap.add_argument("--rows", nargs="+", default=None, choices=["rho", "lfc"],
                   help="which readouts to draw; default both")
    ap.add_argument("--panel", action="store_true",
                   help="strip title/legend/footer for use as a figure panel")
    ap.add_argument("--panel-height", dest="panel_h", type=float, default=2.33,
                   help="panel canvas height per row, inches. The main figure "
                        "uses 2.16 so the right column matches panel A once "
                        "panel C takes 2.50 for its count strip.")
    ap.add_argument("--outdir", default=None)
    ap.add_argument("--datadir", default=None)
    a = ap.parse_args()
    run(a.tissues, a.outdir, a.datadir, a.rows, a.panel, a.panel_h)
```
