## supplemental tables and scripts for "Long non-coding RNA-3′UTR interactions coordinate GABA signaling in the brain": fig_gad1_enrichment.py

```
#!/usr/bin/env python3
"""Where the candidate transcripts sit in the genome-wide Gad1 ranking.

The section's least circular statistic: it uses expression and TargetScan
annotation only - no Gul, no BLAST, no choice of which genes to call targets.
Conserved-8mer transcripts are 0.37-0.46% of those tested but six of the twenty
most Gad1-correlated in every region, and it is the same six genes in optic
lobe, ventral nerve cord and central brain.

One panel per region. x is the correlation itself, increasing to the right; y is
the fraction of transcripts whose correlation is at least x, on a log scale.

DEFAULT: one curve, over every transcript analysed, with all nine candidates
marked on it. The miR-7 site classes are not split out here (that is fig_class_vs_six). The
class statistic is the corner annotation plus the top-20 rule, which together are
the claim: 0.4% of transcripts supply 6 of the 20 most Gad1-correlated.

`--split-classes` draws the two-curve form (conserved 8mer targets against
everything else), where the vertical gap at any rho is the fold enrichment at
that threshold.

Marker shapes match the key used throughout the figure: circle = one of the six,
diamond = Gul-complementary with no miR-7 site (P5CDh1, Glts, Gabat).

Glia excluded and genes retained in < 25% of clusters dropped, matching the
ranking quoted in the text.

Usage:
    python3 fig_gad1_enrichment.py
    python3 fig_gad1_enrichment.py --tissues OL --panel
    python3 fig_gad1_enrichment.py --split-classes        # supplementary form
"""
import argparse
import numpy as np, pandas as pd
from scipy.stats import hypergeom
import matplotlib
matplotlib.use("Agg")
import matplotlib.pyplot as plt
from matplotlib.lines import Line2D
from mpl_toolkits.axes_grid1 import make_axes_locatable

import paths as P
import genes as GN
import gul_mir7_correlation as G
from sitetype_stratification import load_site_types, MIN_RETAIN

TISSUES = [("OL", "gene_by_cluster_meanexpr.csv"),
           ("VNC", "gene_by_cluster_meanexpr_VNC.csv"),
           ("CB", "gene_by_cluster_meanexpr_CB.csv")]
TOPN = 20
COL_8MER = "#08306B"
COL_BAND = "#FDD9B5"
SIX = set(GN.TARGET_POOL)
NO_SITE = ["Gabat", "P5CDh1", "Glts"]
#: rank-style: one set of percentile boundaries, used as the shaded band edges,
#: the count-strip bin edges and the x ticks, so nothing in the panel is drawn
#: against a boundary the rest of the panel does not share
EDGES = [0.1, 1.0, 5.0, 100.0]


def collect(datadir=None):
    site, _ = load_site_types(datadir)
    out = {}
    for tag, csv in TISSUES:
        M = GN.canon_index(pd.read_csv(P.find(csv, datadir), index_col=0))
        M = G.drop_glia(M, tag)
        res = G.analyze(tag, M, GN.GAD1, [], [], set())["res"]
        f = res[res.n_clusters >= MIN_RETAIN * M.shape[1]].sort_values(
            "rho", ascending=False)
        f = f.assign(rank=np.arange(1, len(f) + 1),
                     is8=[site.get(g) == "8mer" for g in f.index])
        out[tag] = f
    return out


def run(outdir=None, datadir=None, tissues=None, panel=False, split=False,
        style="rank", figsize=None):
    def O(n): return P.out(n, outdir)
    plt.rcParams.update({
        "figure.dpi": 110, "savefig.dpi": 200,
        "axes.spines.top": False, "axes.spines.right": False,
        "svg.fonttype": "none", "font.family": "sans-serif",
        "font.sans-serif": ["Arial"]})

    D = collect(datadir)
    order = tissues or ["OL", "VNC", "CB"]
    n = len(order)
    #: top20 needs the rows; rank needs room for the count strip under the plot
    ph = {"top20": 2.66, "rank": 2.50}.get(style, 2.33)
    fig, axes = plt.subplots(1, n, figsize=figsize or ((4.0 * n, 4.3) if n > 1
                             else ((3.61, ph) if panel else (4.6, 3.4))))
    axes = np.atleast_1d(axes)
    rows = []

    def surv(v, grid):
        """fraction of v at or above each point of grid"""
        v = np.sort(np.asarray(v))
        return 1.0 - np.searchsorted(v, grid, side="left") / len(v)

    def sci(v):
        """2.34e-10 -> mathtext 2.3x10^-10. Matches how p is written in the text."""
        e = int(np.floor(np.log10(v)))
        return f"${v / 10 ** e:.1f}{{\\times}}10^{{{e}}}$"

    for i, tag in enumerate(order):
        ax = axes[i]
        f = D[tag]
        N, K = len(f), int(f.is8.sum())
        hits = f[f.is8]
        k = int((hits["rank"] <= TOPN).sum())
        p = hypergeom.sf(k - 1, N, K, TOPN)
        fold = (k / TOPN) / (K / N)
        rows.append(dict(tissue=tag, n_genes=N, n_8mer=K, in_top20=k,
                         fold=fold, hypergeom_p=p))

        bg = f[~f.is8]
        if style == "top20":
            # The head of the ranking itself. Position is not an axis here: a
            # transcript is either in the picture or it is not, so nothing can
            # land in the middle and read as mid-pack.
            head_ = f.head(TOPN)
            extra = sorted((g for g in NO_SITE
                            if g in f.index and int(f.loc[g, "rank"]) > TOPN),
                           key=lambda g: -f.loc[g, "rho"])
            names = list(head_.index) + extra
            ys = np.concatenate([np.arange(TOPN, dtype=float),
                                 TOPN + 0.9 + np.arange(len(extra), dtype=float)])
            for g, yy in zip(names, ys):
                if g in SIX or g in NO_SITE:
                    c = GN.COLORS.get(g, COL_8MER)
                elif f.loc[g, "is8"]:
                    c = COL_8MER
                else:
                    c = "0.80"
                ax.barh(yy, f.loc[g, "rho"], height=0.70, color=c, zorder=3)
            for g, yy in zip(extra, ys[TOPN:]):
                ax.text(f.loc[g, "rho"] + 0.010, yy, f"#{int(f.loc[g, 'rank'])}",
                        va="center", ha="left", fontsize=6.4, color="0.45")
            if len(extra):
                ax.axhline(TOPN - 0.15, color="0.75", lw=0.8, ls=(0, (3, 2)),
                           zorder=5)
            ax.set_yticks(ys)
            ax.set_yticklabels(names, fontsize=6.2)
            for t, g in zip(ax.get_yticklabels(), names):
                t.set_fontstyle("italic")
                t.set_color(GN.COLORS[g] if g in GN.COLORS else "0.35")
            ax.set_ylim(ys[-1] + 0.85, -0.85)
            ax.set_xlim(0, f.rho.max() * 1.20)
            ax.tick_params(axis="x", labelsize=8)
            ax.tick_params(axis="y", length=0)
            ax.set_xlabel("Spearman ρ with $\\it{Gad1}$", fontsize=9.5)
            if not panel:
                ax.set_title(f"{G.TISSUE_NAME.get(tag, tag)} ({tag})", fontsize=10.5)
            # no stats block: the bars are already the statistic, and anything
            # placed here lands on the appended rows. The caption carries
            # "K of N (x%) carry a conserved 8mer site; k of these 20 do".
            continue

        if style == "scatter":
            # Two independent readouts of the same transcripts, so neither axis is
            # a position: the bulk falls in the middle and the candidates fall in
            # a corner. Nothing here can make a top-2% transcript look mid-pack.
            L = pd.read_csv(P.find("gaba_lfc_per_gene.csv", datadir))
            L = L[L.tissue == tag].set_index("gene")
            d = f.join(L["lfc"], how="inner")
            ax.axhline(0, color="0.88", lw=0.8, ls="--", zorder=1)
            ax.axvline(0, color="0.88", lw=0.8, ls="--", zorder=1)
            rest = d[~d.is8]
            ax.scatter(rest.rho, rest.lfc, s=3.0, alpha=0.16, color="0.55",
                       edgecolor="none", zorder=2, rasterized=True)
            hit = d[d.is8]
            ax.scatter(hit.rho, hit.lfc, s=13, alpha=0.85, color=COL_8MER,
                       edgecolor="none", zorder=4)
            xlo_, xhi_ = d.rho.min(), d.rho.max()
            ylo_, yhi_ = d.lfc.min(), d.lfc.max()
            ax.set_xlim(xlo_ - 0.03 * (xhi_ - xlo_), xhi_ + 0.42 * (xhi_ - xlo_))
            ax.set_ylim(ylo_ - 0.04 * (yhi_ - ylo_), yhi_ + 0.06 * (yhi_ - ylo_))
            ax.tick_params(labelsize=8)
            ax.set_xlabel("Spearman ρ with $\\it{Gad1}$", fontsize=9.5)
            if i == 0:
                ax.set_ylabel("log$_2$ fold change\nGABAergic / other neurons",
                              fontsize=9.5)
            if not panel:
                ax.set_title(f"{G.TISSUE_NAME.get(tag, tag)} ({tag})", fontsize=10.5)
            marks = [(g, d.loc[g, "rho"], d.loc[g, "lfc"], "o" if g in SIX else "D")
                     for g in list(SIX) + NO_SITE if g in d.index]
            marks.sort(key=lambda t: -t[2])
            ys = np.linspace(0.97, 0.30, len(marks))
            for (g, x, y, mk), yl in zip(marks, ys):
                col = GN.COLORS.get(g, COL_8MER)
                ax.plot([x], [y], mk, ms=(6.0 if mk == "o" else 5.2), mfc=col,
                        mec="white", mew=0.9, zorder=6)
                ax.annotate(f"$\\it{{{g}}}$", (x, y), xycoords="data",
                            xytext=(0.80, yl), textcoords="axes fraction",
                            fontsize=7.4, color=col, va="center", ha="left",
                            zorder=7,
                            arrowprops=dict(arrowstyle="-", color=col, lw=0.6,
                                            shrinkA=1, shrinkB=3))
            ax.text(0.02, 0.03,
                    f"{len(d):,} transcripts; {int(d.is8.sum())} carry a conserved\n"
                    f"8mer miR-7 site ({100*d.is8.mean():.2f}%)",
                    transform=ax.transAxes, ha="left", va="bottom", fontsize=7.4,
                    color="0.35")
            continue

        if style == "rank":
            # rho against its position in the ranking, as a percentile rather than
            # a raw rank. On raw log rank, rank 161 of 7,550 lands mid-panel and
            # reads as middling when it is the top 2%; the percentile axis and the
            # top-1% / top-5% bands give the eye a ruler instead.
            pct = 100.0 * f["rank"].values / N
            # A strip under the plot counting transcripts per bin. The dots alone
            # do not carry it: past ~1% they merge into a line that reads as one
            # object rather than as thousands of genes. The bars put the number
            # on the page.
            cax = make_axes_locatable(ax).append_axes("bottom", size="24%",
                                                      pad=0.06, sharex=ax)
            # one dot per transcript, not a line, so the sparse end is visibly
            # a few dozen genes rather than a continuous curve
            ax.scatter(pct, f.rho.values, s=2.2, alpha=0.45, color="0.55",
                       edgecolor="none", rasterized=True, zorder=2)
            # the bands lead: their edges become the strip's bin edges and the x
            # ticks below, so every boundary drawn in this panel is the same set
            ax.axvspan(1e-3, 1.0, color=COL_BAND, alpha=0.55, lw=0, zorder=1)
            ax.axvspan(1.0, 5.0, color=COL_BAND, alpha=0.26, lw=0, zorder=1)
            ax.axhline(0.0, color="0.85", lw=0.8, ls="--", zorder=1)
            ax.set_xscale("log")
            xlo = 100.0 / N / 1.25          # just clear of the rank-1 dot
            ax.set_xlim(xlo, 100.0)
            lo, hi = f.rho.min(), f.rho.max()
            ax.set_ylim(lo - 0.03 * (hi - lo), hi + 0.13 * (hi - lo))
            # band labels go low, in the empty wedge under the curve: the top of
            # the panel belongs to the leader lines
            xt = ax.get_xaxis_transform()      # x in data, y in axes fraction
            ax.text(np.sqrt(xlo * 1.0), 0.30, "top 1%", color="#B87020",
                    fontsize=6.8, ha="center", va="center", transform=xt)
            ax.text(np.sqrt(1.0 * 5.0), 0.30, "top 5%", color="#C08048",
                    fontsize=6.8, ha="center", va="center", transform=xt)
            x20 = 100.0 * TOPN / N
            ax.axvline(x20, color="#B87020", lw=0.9, ls=":", zorder=4)
            ax.text(x20 * 1.15, 0.44, f"top {TOPN}", color="#B87020", fontsize=6.8,
                    ha="left", va="center", transform=xt)
            # each tick carries how many transcripts are at or above it, so the
            # axis states its own selectivity instead of implying it
            # Bins are the band edges plus 0.1%, so each shaded band in the plot
            # above spans a whole number of bars. On a log count axis with the
            # count printed: a linear axis rendered the 7-gene bin as zero.
            dec = [xlo] + [t for t in EDGES if t > xlo]
            cnt, _ = np.histogram(pct, bins=dec)
            cax.axvspan(1e-3, 1.0, color=COL_BAND, alpha=0.55, lw=0, zorder=1)
            cax.axvspan(1.0, 5.0, color=COL_BAND, alpha=0.26, lw=0, zorder=1)
            # no edgecolor: a stroke is centred on the bin edge, so it puts half
            # its width either side of 1% and 5% and the bars stop lining up with
            # the bands above. The height steps separate the bars on their own.
            cax.bar(dec[:-1], cnt, width=np.diff(dec), align="edge",
                    color="0.55", lw=0, zorder=3)
            cax.set_yscale("log")
            cax.set_ylim(0.6, cnt.max() * 7)
            cax.set_yticks([])
            cax.set_ylabel("genes", fontsize=7.2, color="0.35")
            for x0, x1, c in zip(dec[:-1], dec[1:], cnt):
                cax.text(np.sqrt(x0 * x1), c * 1.45, f"{c:,}", ha="center",
                         va="bottom", fontsize=6.6, color="0.30", zorder=4)
            ax.text(0.02, 0.19, "each dot is one transcript", transform=ax.transAxes,
                    fontsize=6.6, ha="left", va="bottom", color="0.45")
            # counts live on the strip's bars, so the ticks carry the percentile
            # alone; they used to repeat the running total under each label
            ticks = [t for t in EDGES if t * N / 100 >= 1]
            cax.set_xticks(ticks)
            cax.set_xticklabels([f"{t:g}%" for t in ticks])
            cax.minorticks_off()
            cax.tick_params(labelsize=8)
            ax.tick_params(axis="x", labelbottom=False, length=0)
            ax.yaxis.set_major_locator(plt.MaxNLocator(5))
            cax.set_xlabel(f"percentile of {N:,} transcripts", fontsize=9.5)
            if i == 0:
                ax.set_ylabel("Spearman ρ with $\\it{Gad1}$", fontsize=9.5)
            marks = [(g, 100.0 * f.loc[g, "rank"] / N, f.loc[g, "rho"],
                      "o" if g in SIX else "D")
                     for g in list(SIX) + NO_SITE if g in f.index]
            # Five of the nine sit within 0.14 of each other on rho, so the labels
            # have to leave the curve. The column sits just right of the last
            # marker, in the empty wedge above the tail, to keep leaders short.
            marks.sort(key=lambda t: -t[2])
            ys = np.linspace(0.985, 0.50, len(marks))
            for (g, x, y, mk), yl in zip(marks, ys):
                col = GN.COLORS.get(g, COL_8MER)
                ax.plot([x], [y], mk, ms=(5.8 if mk == "o" else 5.0), mfc=col,
                        mec="white", mew=0.8, zorder=6)
                ax.annotate(f"$\\it{{{g}}}$", (x, y), xycoords="data",
                            xytext=(0.62, yl), textcoords="axes fraction",
                            fontsize=7.4, color=col, va="center", ha="left",
                            zorder=7,
                            arrowprops=dict(arrowstyle="-", color=col, lw=0.6,
                                            shrinkA=1, shrinkB=3))
            if not panel:
                ax.set_title(f"{G.TISSUE_NAME.get(tag, tag)} ({tag})", fontsize=10.5)
            # single spaces only: SVG collapses runs of whitespace, so anything
            # aligned with padding here falls apart in Illustrator
            ax.text(0.02, 0.03,
                    f"conserved 8mer miR-7 targets: {K} of {N:,} ({100*K/N:.2f}%)\n"
                    f"{k} of the top {TOPN}, {fold:.0f}-fold, p = {sci(p)}",
                    transform=ax.transAxes, ha="left", va="bottom", fontsize=7.4,
                    color="0.35")
            continue

        grid = np.linspace(f.rho.min(), f.rho.max(), 600)
        if split:
            ax.plot(grid, surv(bg.rho.values, grid), color="0.6", lw=1.5, zorder=2,
                    label="no conserved 8mer site")
            ax.plot(grid, surv(hits.rho.values, grid), color=COL_8MER, lw=2.0,
                    zorder=3, label="conserved 8mer miR-7 target")
        else:
            ax.plot(grid, surv(f.rho.values, grid), color="0.35", lw=1.8, zorder=2,
                    label="all transcripts analysed")
        rho20 = f.rho.iloc[TOPN - 1]
        ax.axvline(rho20, color="#E0A050", lw=1.0, ls="--", zorder=1)
        ax.text(rho20, 1.35, f"top {TOPN}", color="#B87020", fontsize=7,
                ha="center", va="bottom")

        # Every candidate sits on the curve it belongs to: one curve by default,
        # the 8mer / no-8mer pair under --split-classes. Label ys are assigned in
        # curve-height order, not rho order, or the leader lines cross.
        marks = []
        for g in list(SIX) + NO_SITE:
            if g not in f.index:
                continue
            x = f.loc[g, "rho"]
            src = ((hits if g in SIX else bg) if split else f).rho.values
            marks.append((g, x, surv(src, np.array([x]))[0], "o" if g in SIX else "D"))
        marks.sort(key=lambda t: -t[2])
        ys = np.linspace(0.96, 0.28, len(marks))
        for (g, x, y, mk), yl in zip(marks, ys):
            col = GN.COLORS.get(g, COL_8MER)
            ax.plot([x], [y], mk, ms=(5.5 if mk == "o" else 4.8), mfc=col,
                    mec="white", mew=0.8, zorder=6)
            ax.annotate(f"$\\it{{{g}}}$", (x, y), xycoords="data",
                        xytext=(0.80, yl), textcoords="axes fraction",
                        fontsize=7.4, color=col, va="center", ha="left", zorder=7,
                        arrowprops=dict(arrowstyle="-", color=col, lw=0.6,
                                        shrinkA=1, shrinkB=3))

        ax.set_yscale("log"); ax.set_ylim(0.5 / N, 1.9)
        ax.set_xlim(f.rho.min(), f.rho.min() + (f.rho.max() - f.rho.min()) * 1.62)
        ax.tick_params(labelsize=8)
        if not panel:
            ax.set_title(f"{G.TISSUE_NAME.get(tag, tag)} ({tag})", fontsize=10.5)
        ax.set_xlabel("Spearman ρ with $\\it{Gad1}$", fontsize=9.5)
        if i == 0:
            ax.set_ylabel("fraction of group above this ρ" if split else
                          "fraction of transcripts above this ρ", fontsize=9.5)
        note = (f"{K} of {N:,} ({100*K/N:.2f}%)\n"
                f"{k} in the top {TOPN}, {fold:.0f}-fold, p = {sci(p)}" if split else
                f"conserved 8mer miR-7 targets: {K} of {N:,} ({100*K/N:.2f}%)\n"
                f"{k} of the top {TOPN}, {fold:.0f}-fold, p = {sci(p)}")
        ax.text(0.02, 0.03, note, transform=ax.transAxes, ha="left", va="bottom",
                fontsize=7.4, color="0.35")

    if not panel:
        curves = ([Line2D([], [], color=COL_8MER, lw=2.0,
                          label="conserved 8mer miR-7 targets"),
                   Line2D([], [], color="0.6", lw=1.5, label="all other transcripts")]
                  if split else
                  [Line2D([], [], color=("0.55" if style == "rank" else "0.35"),
                          lw=1.8, label="all transcripts analysed")])
        h = curves + [
            Line2D([], [], marker="o", ls="none", mfc="0.35", mec="white", ms=6,
                   label="the six"),
            Line2D([], [], marker="D", ls="none", mfc="0.35", mec="white", ms=5.5,
                   label="$\\it{Gul}$-complementary, no miR-7 site")]
        fig.legend(handles=h, frameon=False, ncol=(4 if n > 1 else 2),
                   loc="lower center",
                   bbox_to_anchor=(0.5, -0.07 if n > 1 else -0.13), fontsize=8.4)
        fig.suptitle("Conserved miR-7 8mer targets are concentrated among the most "
                     "$\\it{Gad1}$-correlated transcripts", fontsize=11.5)
        fig.text(0.5, -0.005 if n > 1 else -0.26,
                 ("The vertical gap between the two curves at any \u03c1 is the "
                  "fold enrichment of 8mer targets at that threshold." if split else
                  "Conserved 8mer miR-7 targets are 0.4% of the transcripts tested "
                  "and supply 6 of the 20 most $\\it{Gad1}$-correlated."),
                 ha="center", fontsize=7.6, color="0.35")
    fig.tight_layout(pad=0.4) if panel else fig.tight_layout(rect=[0, 0.04, 1, 0.92])
    # the style is always in the filename: four of them coexist while the panel
    # is being chosen, and a bare name would silently mean whichever ran last
    suffix = ("" if not tissues else "_" + "".join(order)) + \
             ("_classes" if split else "") + "_" + style + \
             ("_panel" if panel else "")
    path = O(f"fig_gad1_enrichment{suffix}.png")
    fig.savefig(path, bbox_inches="tight")
    fig.savefig(path.replace(".png", ".svg"), bbox_inches="tight")
    plt.close(fig)

    T = pd.DataFrame(rows)
    T.to_csv(O("gad1_enrichment_top20.csv"), index=False)
    print(T.to_string(index=False, float_format=lambda v: f"{v:.3g}"))
    for tag in order:
        f = D[tag]
        print(f"\n{tag} top {TOPN} 8mer targets: "
              + ", ".join(f"{g} (#{int(r)})" for g, r in
                          zip(f[f.is8].index[:8], f[f.is8]['rank'][:8])
                          if r <= TOPN))
    return T


if __name__ == "__main__":
    ap = argparse.ArgumentParser(description=__doc__,
                                 formatter_class=argparse.RawDescriptionHelpFormatter)
    ap.add_argument("--tissues", nargs="+", default=None)
    ap.add_argument("--panel", action="store_true",
                   help="strip title/legend/footer for use as a figure panel")
    ap.add_argument("--style", default="scatter",
                   choices=["scatter", "top20", "rank", "survival"],
                   help="scatter: rho against GABAergic fold change, no position "
                        "axis. top20: the head of the ranking as bars. rank: rho "
                        "against percentile. survival: the old decay curve.")
    ap.add_argument("--split-classes", dest="split", action="store_true",
                   help="survival style only: two curves, 8mer targets against "
                        "everything else (the earlier form; for a supplement)")
    ap.add_argument("--outdir", default=None)
    ap.add_argument("--datadir", default=None)
    a = ap.parse_args()
    run(a.outdir, a.datadir, a.tissues, a.panel, a.split,
        "survival" if a.split else a.style)
```
