## supplemental tables and scripts for "Long non-coding RNA-3′UTR interactions coordinate GABA signaling in the brain": fig_gul_abundance.py

```
#!/usr/bin/env python3
"""
Manuscript panel: Gul is among the most abundant transcripts in the CNS, and is
present in large excess over the miR-7 8mer target pool it would compete with.

A  rank-abundance curve per tissue (glia excluded), with Gul and reference
   transcripts marked.
B  stoichiometry -- Gul site abundance (expression x 3 tandem TGGAAGACA copies)
   against the summed site abundance of the six proposed 8mer targets
   (genes.TARGET_POOL), in GABAergic clusters where the mechanism is proposed to
   act. GABAergic = Otsu cut on log Gad1, the paper-wide definition.

Values are AverageExpression on the RNA 'data' layer: comparable across genes
within a tissue, but not absolute molar concentrations.

Usage:
    python3 fig_gul_abundance.py          # writes to results/{tables,figures}
"""
import os, argparse
import numpy as np, pandas as pd
import paths as P
import genes as GN
import matplotlib
matplotlib.use("Agg")
import matplotlib.pyplot as plt
from matplotlib.lines import Line2D
from fig_switchlike_gaba import gad1_threshold

GUL = "CG14989"
GUL_COPIES = 3                      # tandem TGGAAGACA repeats in the Gul 3'UTR
# The proposed target pool, defined once in genes.py -- the six conserved-8mer miR-7
# targets at the top of the Gad1 ranking:
#   VGAT      vesicular GABA transporter
#   sgll      pyridox(am)ine 5'-phosphate oxidase -> PLP, cofactor for Gad1 and Gabat
#   Ssadh     succinic semialdehyde dehydrogenase (GABA shunt)
#   ScsbetaG  succinyl-CoA synthetase beta (converges on succinate)
#   CG32225   uncharacterised; reported LAMP5 orthologue, traffics VGAT
#   CG10444   SLC5-family symporter -- in the ranking, NOT in the GABA module
TARGET_POOL = list(GN.TARGET_POOL)
TISSUES = [("gene_by_cluster_meanexpr.csv", "OL", "Optic lobe"),
           ("gene_by_cluster_meanexpr_VNC.csv", "VNC", "Ventral nerve cord"),
           ("gene_by_cluster_meanexpr_CB.csv", "CB", "Central brain")]
TCOL = {"OL": "#4C72B0", "VNC": "#55A868", "CB": "#8172B2"}
GUL_C, REF_C = "#C44E52", "#9A9A9A"

#: Set by --panel / --figsize. The full form is 9.10 x 4.02 in, wider than A4, so it could
#: only ever have been placed shrunk. PANEL sends the two explanatory legends to the caption
#: and keeps only the tissue key, which is what lets the panels narrow without collisions.
PANEL = False
FIGSIZE = None
REFS = [("RpL32", "RpL32"), ("Act5C", "Act5C"), ("Gad1", "Gad1")]


def neuronal(csv):
    M = GN.canon_index(pd.read_csv(csv, index_col=0))
    repo = M.loc["repo"]
    glial = set(repo[repo >= 0.17].index) | {c for c in M.columns if c.startswith("G-")}
    return M[[c for c in M.columns if c not in glial]]


def run(outdir=None, datadir=None):
    def O(name): return P.out(name, outdir)
    plt.rcParams.update({
        "figure.dpi": 110, "savefig.dpi": 300,
        "axes.spines.top": False, "axes.spines.right": False,
        "svg.fonttype": "none", "font.family": "sans-serif",
        "font.sans-serif": ["Arial"], "axes.linewidth": 0.8,
    })
    pool = set(TARGET_POOL)

    fig, axes = plt.subplots(1, 2, figsize=FIGSIZE or (9.4, 4.2),
                             gridspec_kw=dict(width_ratios=[1.5, 1]))
    ax = axes[0]
    summary, stoich = [], []
    for csv, tag, full in TISSUES:
        N = neuronal(P.find(csv, datadir))
        mean = N.mean(axis=1).sort_values(ascending=False)
        mean = mean[mean > 0]
        rank = np.arange(1, len(mean) + 1)
        ax.plot(rank, mean.values, color=TCOL[tag], lw=1.6, alpha=0.9,
                zorder=3, label=f"{full} ({tag})")
        r_gul = int(np.where(mean.index == GUL)[0][0]) + 1
        ax.scatter([r_gul], [mean[GUL]], s=64, facecolor=GUL_C, edgecolor="black",
                   linewidth=1.0, zorder=7)
        summary.append(dict(tissue=tag, gul=mean[GUL], rank=r_gul, n=len(mean),
                            pct=100 * (1 - r_gul / len(mean))))
        # stoichiometry in GABAergic clusters. The split is the Otsu cut on log Gad1,
        # the same rule fig_switchlike_gaba and the sensitivity analysis use, so one
        # definition of "GABAergic" serves the whole paper.
        gad = N.loc[GN.GAD1]; gaba = gad[gad >= gad1_threshold(gad.values)].index
        gul_ab = N.loc[GUL, gaba].mean() * GUL_COPIES
        present = [g for g in pool if g in N.index and g != GUL]
        per = N.loc[present, gaba].mean(axis=1)
        tgt_ab = per.sum()
        stoich.append(dict(tissue=tag, gul_sites=gul_ab, target_sites=tgt_ab,
                           ratio=gul_ab / tgt_ab, n_targets=len(present),
                           genes=",".join(sorted(present)),
                           **{f"expr_{g}": per.get(g, np.nan) for g in TARGET_POOL}))
    S = pd.DataFrame(summary); ST = pd.DataFrame(stoich)

    # reference genes: mark on the OL curve only, to keep the panel clean
    N0 = neuronal(P.find(TISSUES[0][0], datadir))
    m0 = N0.mean(axis=1).sort_values(ascending=False); m0 = m0[m0 > 0]
    # de-collide the reference labels. Act5C needs the largest drop: it sits in the
    # steepest stretch of the curve, so a shallow offset puts the line through the text.
    offs = [(9, 10), (6, -20), (10, 4)]
    for (sym, lab), off in zip(REFS, offs):
        if sym not in m0.index: continue
        r = int(np.where(m0.index == sym)[0][0]) + 1
        ax.scatter([r], [m0[sym]], s=26, facecolor="white", edgecolor=REF_C,
                   linewidth=1.2, zorder=6)
        # RpL32, Act5C and Gad1 are gene symbols, so they take italic. style="italic"
        # rather than mathtext: mathtext leaves the digits upright and splits the label
        # into one tspan per glyph, which Illustrator cannot edit.
        ax.annotate(lab, (r, m0[sym]), textcoords="offset points", xytext=off,
                    fontsize=7.5, color="0.4", style="italic",
                    arrowprops=dict(arrowstyle="-", color="0.75", lw=0.6,
                                    shrinkA=0, shrinkB=2))
    g0 = S[S.tissue == "OL"].iloc[0]
    ax.annotate("$\\it{Gul}$", (g0["rank"], g0["gul"]), textcoords="offset points",
                xytext=(20, 14), fontsize=11, color=GUL_C, fontweight="bold",
                arrowprops=dict(arrowstyle="-", color=GUL_C, lw=1.0))
    ranks = "\n".join(f"  {r.tissue}:  rank {int(r['rank'])} of {int(r['n']):,}"
                      for _, r in S.iterrows())
    # This block used to sit at (0.985, 0.62), directly on the rank-abundance curve.
    # The empty region is the lower left: over the first two decades of rank the curve
    # occupies only the top ~30% of the axes, so the text clears it and the tissue legend.
    ax.text(0.03, 0.55, ranks, transform=ax.transAxes, ha="left", va="top",
            fontsize=7.5, color=GUL_C, family="monospace")
    ax.set_xscale("log"); ax.set_yscale("log")
    ax.set_ylim(1e-4, None)                       # drop the uninformative tail
    ax.set_xlabel("transcript rank", fontsize=9.5)
    ax.set_ylabel("mean expression across neuronal clusters", fontsize=9.5)
    ax.tick_params(labelsize=8)
    ax.set_title("A", fontsize=11, loc="left", fontweight="bold")
    leg = ax.legend(frameon=False, fontsize=7.5, loc="lower left", handlelength=1.4)
    ax.add_artist(leg)
    if not PANEL:                     # Gul and the references are labelled on the plot
            ax.legend(handles=[Line2D([], [], marker="o", ls="none", mfc=GUL_C, mec="black",
                                  mew=1.0, ms=7, label="$\\it{Gul}$"),
                           Line2D([], [], marker="o", ls="none", mfc="white", mec=REF_C,
                                  mew=1.2, ms=6, label="reference transcripts")],
                  frameon=False, fontsize=7.5, loc="upper right")

    # ---- B: stoichiometry ----
    ax = axes[1]
    x = np.arange(len(ST)); w = 0.36
    ax.bar(x - w/2, ST.gul_sites, w, color=GUL_C, edgecolor="black", linewidth=0.7,
           label=("$\\it{Gul}$ mimic sites" if PANEL else
                  "$\\it{Gul}$ mimic sites\n(expression × 3 copies)"), zorder=3)
    ax.bar(x + w/2, ST.target_sites, w, color="#7F9DB9", edgecolor="black",
           linewidth=0.7, zorder=3,
           label="six proposed targets" if PANEL else "the six proposed 8mer targets")
    for i, r in ST.iterrows():
        top = max(r.gul_sites, r.target_sites)
        ax.text(i, top * 1.15, f"{r.ratio:.1f}×", ha="center", fontsize=8.5,
                fontweight="bold", color=GUL_C)
    ax.set_yscale("log")
    ax.set_xticks(x); ax.set_xticklabels(ST.tissue, fontsize=9)
    # NOT "site abundance": only the Gul bar counts sites (expression x 3 tandem copies).
    # The target bar is summed TRANSCRIPT abundance, one entry per gene, so VGAT contributes
    # once despite carrying three conserved 8mers.
    ax.set_ylabel("abundance in GABAergic clusters", fontsize=9.5)
    ax.tick_params(axis="y", labelsize=8)
    ax.set_title("B", fontsize=11, loc="left", fontweight="bold")
    if PANEL:
        # Inside the axes, in the empty band above the bars. Dropping this legend when the
        # figure was page-sized left the two colours undecodable -- RL asked what the blue
        # bar was. The labels are shortened to fit there; "expression x 3 tandem copies"
        # and "combined transcript abundance" move to the caption.
        ax.legend(frameon=False, fontsize=6.6, loc="upper left", handlelength=1.1,
                  handletextpad=0.5, labelspacing=0.35, borderaxespad=0.3)
    else:
        ax.legend(frameon=False, fontsize=7.5, loc="upper center",
                  bbox_to_anchor=(0.5, -0.14), ncol=2)
    ax.margins(y=0.28)
    if PANEL:
        # headroom for the legend, added at the TOP only. margins() pads both ends, which
        # on a log axis buys the legend one decade of space and costs a dead decade under
        # the blue bars, making them look smaller than they are.
        lo, hi = ax.get_ylim()
        ax.set_ylim(lo, hi * 4.5)

    fig.tight_layout(w_pad=1.4 if PANEL else 2.0)
    p = O("fig_gul_abundance_panel.png" if PANEL else "fig_gul_abundance.png")
    fig.savefig(p, bbox_inches="tight"); fig.savefig(p.replace(".png", ".svg"), bbox_inches="tight")
    plt.close(fig)
    S.to_csv(O("gul_abundance_summary.csv"), index=False)
    ST.to_csv(O("gul_stoichiometry.csv"), index=False)
    print(S.to_string(index=False, float_format=lambda v: f"{v:,.2f}"))
    print()
    print(ST.to_string(index=False, float_format=lambda v: f"{v:,.2f}"))


if __name__ == "__main__":
    ap = argparse.ArgumentParser(description=__doc__,
                                 formatter_class=argparse.RawDescriptionHelpFormatter)
    ap.add_argument("--outdir", default=None,
                    help="write everything here; default routes to results/{tables,figures}")
    ap.add_argument("--datadir", default=None, help="extra directory to search for inputs")
    ap.add_argument("--panel", action="store_true",
                    help="send the two explanatory legends to the caption; writes "
                         "*_panel.svg")
    ap.add_argument("--figsize", nargs=2, type=float, default=None, metavar=("W", "H"),
                    help="figure size in inches; generate at the placed size")
    a = ap.parse_args()
    FIGSIZE, PANEL = (tuple(a.figsize) if a.figsize else None), a.panel
    run(a.outdir, a.datadir)
```
