## supplemental tables and scripts for "Long non-coding RNA-3′UTR interactions coordinate GABA signaling in the brain": fig_switchlike_gaba.py

```
#!/usr/bin/env python3
"""Which genes are in every GABAergic cluster and essentially no other?

Over ~7,000-8,000 expressed genes per region, the genes detected in >= 90% of
GABAergic clusters and <= 20% of the rest are Gul and Gad1 in every region (both at
~100% of GABAergic clusters against 1-9% elsewhere). Lim3 enters the OL corner only
under a looser cutoff rule and is anti-correlated with Gad1 in VNC and CB.

Cutoff sensitivity (written to gaba_cutoff_sensitivity.csv): sweeping the Gad1 cut
from the 60th to the 90th percentile, the corner is sometimes empty but never
contains any gene other than Gul, Gad1 and -- at one OL cutoff -- Lim3.

Definitions, over non-glial clusters (repo < 0.17, plus OL's named G- clusters):
  GABAergic  Gad1 above a per-region Otsu threshold on the bimodal log Gad1
             distribution (the cut lands at 9.9 / 8.1 / 3.7)
  detected   cluster mean above 10% of the gene's own 90th percentile

Usage:
    python3 fig_switchlike_gaba.py
"""
import argparse
import numpy as np, pandas as pd
import matplotlib
matplotlib.use("Agg")
import matplotlib.pyplot as plt
from matplotlib.lines import Line2D
import paths as P
import genes as GN
import gul_mir7_correlation as G

TISSUES = [("gene_by_cluster_meanexpr.csv", "OL"),
           ("gene_by_cluster_meanexpr_VNC.csv", "VNC"),
           ("gene_by_cluster_meanexpr_CB.csv", "CB")]

GABA_CUT, OTHER_CUT = 0.90, 0.20     # the "pan-GABAergic and specific" corner
#: Axis padding. Marker size is fixed in POINTS, so the same marker covers more of the
#: data range as the axes shrink: at 3.9 in per sub-panel the 0.05 margin was wider than
#: the Gul/Gad1 marker's radius, at 2.5 in it is not, and markers sitting at exactly
#: x = 1.0 get clipped by the spine. Panel mode pads further. The corner rectangle is
#: drawn to the same limit so it still reaches the edge.
def _pad():
    return (1.13, -0.09) if PANEL else (1.05, -0.05)
MIN_MEAN, MIN_P90 = 0.02, 0.05

#: Gul and Gad1 are EXACTLY coincident in VNC and CB (det_gaba 1.000, det_other 0.0036
#: and 0.0615 for both) and 0.022 apart in OL, well inside one marker. Drawn as equal
#: filled discs they read as a single point, which hides the very thing the panel claims:
#: that TWO transcripts occupy the corner. So Gad1 is a large open ring and Gul a smaller
#: filled disc inside it -- both at their true positions, no offset, and unambiguously two
#: marks. The third field is the marker size.
ANCHORS = {"CG14989": ("Gul", "#C0392B", 96), "Gad1": ("Gad1", "#1F6FB4", 96)}
TARGETS = ["VGAT", "CG32225", "Ssadh", "sgll", "ScsbetaG", "Gabat", "P5CDh1", "Glts", "stol"]


def gad1_threshold(g, nbins=256):
    """Otsu split of the bimodal log Gad1 distribution: maximise between-class variance.

    Replaced an earlier largest-upper-half-gap rule, which in OL put the cut at 28.9,
    inside the high plateau, rather than at the distribution's actual step near 10 (see
    fig_gaba_cutoff.py, which draws both). Otsu is outcome-blind and gives closely
    matched cluster counts across regions: 46 / 50 / 68 against the gap rule's 42 / 48 / 65.
    """
    x = np.log10(np.asarray(g, float) + 0.01)
    hist, edges = np.histogram(x, bins=nbins)
    p = hist / hist.sum(); c = np.cumsum(p)
    mids = (edges[:-1] + edges[1:]) / 2
    mu = np.cumsum(p * mids); mut = mu[-1]
    with np.errstate(invalid="ignore", divide="ignore"):
        sb = (mut * c - mu) ** 2 / (c * (1 - c))
    return 10 ** mids[np.nanargmax(sb)] - 0.01


def largest_gap_threshold(g):
    """The earlier rule, kept for the comparison drawn in fig_gaba_cutoff.py."""
    v = np.sort(np.log10(np.asarray(g, float) + 0.01))
    k = len(v) // 2
    i = int(np.argmax(np.diff(v[k:])))
    return 10 ** ((v[k:][i] + v[k:][i + 1]) / 2) - 0.01


def table(csv):
    M = GN.canon_index(pd.read_csv(P.find(csv), index_col=0))
    repo = M.loc["repo"]
    glial = set(repo[repo >= 0.17].index) | {c for c in M.columns if c.startswith("G-")}
    N = M[[c for c in M.columns if c not in glial]]
    g = N.loc[GN.GAD1]
    cut = gad1_threshold(g.values)
    gaba, other = g[g >= cut].index, g[g < cut].index
    p90 = N.quantile(.90, axis=1)
    thr = 0.1 * p90
    d = pd.DataFrame(dict(det_gaba=N[gaba].gt(thr, axis=0).mean(axis=1),
                          det_other=N[other].gt(thr, axis=0).mean(axis=1)))
    keep = (N.mean(axis=1) > MIN_MEAN) & (p90 > MIN_P90)
    return d[keep], cut, len(gaba), len(other)


def declutter(ys, gap, lo, hi):
    idx = np.argsort(ys); y0 = np.sort(np.asarray(ys, float)); y = y0.copy()
    for i in range(1, len(y)):
        y[i] = max(y[i], y[i-1] + gap)
    y -= (y.mean() - y0.mean())
    if y[-1] > hi: y -= y[-1] - hi
    if y[0] < lo:  y += lo - y[0]
    out = np.empty_like(y); out[idx] = y
    return out


def make_fig(D, meta, path):
    plt.rcParams.update({
        "figure.dpi": 110, "savefig.dpi": 200,
        "axes.spines.top": False, "axes.spines.right": False,
        "svg.fonttype": "none", "font.family": "sans-serif", "font.sans-serif": ["Arial"],
    })
    fig, axes = plt.subplots(1, 3, figsize=(FIGSIZE or (11.8, 4.4)))
    for i, (ax, (tag, d)) in enumerate(zip(np.atleast_1d(axes), D.items())):
        cut, n_gaba, n_other, corner = meta[tag]
        XHI, YLO = _pad()
        ax.add_patch(plt.Rectangle((GABA_CUT, YLO), XHI - GABA_CUT, OTHER_CUT - YLO,
                                   color="0.92", zorder=0))
        # alpha 0.09 was tuned for the dense band and made isolated genes almost
        # invisible, which is where the informative outliers are. Darker and less
        # transparent, with a smaller marker so the dense band reads by coverage
        # rather than by opacity and still does not saturate into a blob.
        ax.scatter(d.det_gaba, d.det_other, s=3.2, alpha=0.34, color="#5F5F5F",
                   edgecolor="none", rasterized=True, zorder=2)
        ax.axvline(GABA_CUT, color="0.55", lw=0.8, ls=":", zorder=3)
        ax.axhline(OTHER_CUT, color="0.55", lw=0.8, ls=":", zorder=3)

        marks = []
        # Gul and Gad1 are EXACTLY coincident in VNC and CB and 0.022 apart in OL, so one
        # marker hides the other and the panel's whole claim -- that TWO transcripts occupy
        # the corner -- becomes invisible. RL's solution: contrasting hues plus alpha, so
        # the overlap blends to a third colour and the two read as two even at heavy
        # overlap. That means the displacement can stay small. It is one-sided and inward
        # because both axes are fractions bounded at 0 and 1 and both anchors sit on those
        # bounds: a symmetric jitter would put one at x > 1 or y < 0, i.e. more than 100%
        # of clusters or a negative fraction. Caption must declare the offset.
        JITTER = 0.028
        for g, (name, col, ms) in sorted(ANCHORS.items(), key=lambda kv: -kv[1][2]):
            if g not in d.index: continue
            x, y = d.det_gaba[g], d.det_other[g]
            px, py = (x - JITTER, y + JITTER) if g == "CG14989" else (x, y)
            ax.scatter([px], [py], s=ms, facecolor=col, edgecolor="black",
                       linewidth=0.7, alpha=0.68, zorder=7)
            marks.append((px, py, name, col))
        # NARRATIVE CONSTRAINT. As Fig. S8 this panel is cited at paragraph 30, in the
        # section that introduces Gul -- every gene in TARGETS except VGAT is a section 5
        # result, so marking them here shows the reader the answer several pages early.
        # Paragraph 30's claim is only that Gul and Gad1 occupy the corner, which the
        # ANCHORS above already carry. Same reasoning as rbp_gaba_screen.py's rule that
        # Gul goes unlabelled in Fig. S2, running the other way.
        # most targets sit on top of each other at (1, ~1); label those as one group
        ubiq = []
        for g in ([] if PANEL else TARGETS):
            if g not in d.index: continue
            ax.scatter([d.det_gaba[g]], [d.det_other[g]], s=36,
                       color=GN.COLORS.get(g, "#4C72B0"), edgecolor="black",
                       linewidth=0.5, zorder=5)
            if d.det_other[g] < 0.95:
                marks.append((d.det_gaba[g], d.det_other[g], g, GN.COLORS.get(g, "#4C72B0")))
            else:
                ubiq.append(g)

        ax.set_xlim(-0.05, XHI); ax.set_ylim(YLO, 1.20)
        marks.sort(key=lambda m: -m[1])
        ys = declutter([m[1] for m in marks], 0.085, 0.02, 1.00)
        for (x, y, name, col), yl in zip(marks, ys):
            ax.annotate(f"$\\it{{{name}}}$", (x, y), xycoords="data",
                        xytext=(1.12, yl), textcoords="data", fontsize=8, color=col,
                        ha="left", va="center", zorder=7, annotation_clip=False,
                        arrowprops=dict(arrowstyle="-", color=col, lw=0.6,
                                        shrinkA=2, shrinkB=3))
        if ubiq:
            ax.annotate(f"{len(ubiq)} putative targets\n(detected in ~all clusters)",
                        (1.0, 1.0), xycoords="data", xytext=(1.12, 1.13),
                        textcoords="data", fontsize=7.5, color="0.35", ha="left",
                        va="center", zorder=7, annotation_clip=False,
                        arrowprops=dict(arrowstyle="-", color="0.55", lw=0.6,
                                        shrinkA=2, shrinkB=3))
        if not PANEL:
            names = ", ".join(f"$\\it{{{n}}}$" for n in corner)
            ax.text(0.03, 0.30,
                    f"{len(corner)} of {len(d):,} expressed\ngenes in this corner:\n{names}",
                    transform=ax.transAxes, ha="left", va="top", fontsize=7.5, color="0.3")
        # the three axes share their labels; in a composite show each once, which also
        # stops the leftmost x label overhanging the figure and being clipped
        if not PANEL or i == 1:
            ax.set_xlabel("fraction of GABAergic clusters detected", fontsize=9)
        if not PANEL or i == 0:
            ax.set_ylabel("fraction of other clusters detected", fontsize=9)
        ax.tick_params(labelsize=8)
        ax.set_title(f"{G.TISSUE_NAME.get(tag, tag)} ({tag})" if PANEL else
                     (f"{G.TISSUE_NAME.get(tag, tag)} ({tag})\n"
                      f"{n_gaba} GABAergic / {n_other} other clusters "
                      f"(Gad1 $\\geq$ {cut:.1f})"), fontsize=9.5, pad=8)
    handles = [Line2D([], [], marker="o", ls="none", mfc="#8A8A8A", mec="none", ms=6,
                      label="all expressed genes"),
               Line2D([], [], marker="o", ls="none", mfc="#C44E52", mec="black", mew=0.8,
                      ms=8, label="Gul / Gad1"),
               Line2D([], [], marker="o", ls="none", mfc="#4DAF4A", mec="black", mew=0.5,
                      ms=7, label="putative targets")]
    if PANEL:
        # suptitle and legend go to the caption, the same convention the other panel
        # scripts follow; it also recovers the height a composite needs
        fig.tight_layout(w_pad=2.0)
        path = path.replace(".png", "_panel.png")
    else:
        fig.legend(handles=handles, frameon=False, ncol=3, loc="lower center",
                   bbox_to_anchor=(0.5, 0.0), fontsize=8.5)
        fig.suptitle("Gul and Gad1 are the only transcripts present across the whole "
                     "GABAergic population and almost nowhere else", fontsize=11)
        fig.tight_layout(rect=[0, 0.08, 1, 0.90], w_pad=3.4)
    fig.savefig(path, bbox_inches="tight")
    fig.savefig(path.replace(".png", ".svg"), bbox_inches="tight")
    plt.close(fig)


def sweep(outdir=None):
    """Corner membership across Gad1 cutoffs; the identity of the answer is what matters."""
    rows = []
    for csv, tag in TISSUES:
        M = GN.canon_index(pd.read_csv(P.find(csv), index_col=0))
        repo = M.loc["repo"]
        glial = set(repo[repo >= 0.17].index) | {c for c in M.columns if c.startswith("G-")}
        N = M[[c for c in M.columns if c not in glial]]
        g = N.loc[GN.GAD1].astype(float)
        p90 = N.quantile(.90, axis=1); thr = 0.1 * p90
        exp = (N.mean(axis=1) > MIN_MEAN) & (p90 > MIN_P90)
        for q in [0.60, 0.65, 0.70, 0.75, 0.80, 0.85, 0.90]:
            cut = float(np.quantile(g, q)); gaba = g[g >= cut].index; other = g[g < cut].index
            if len(gaba) < 8: continue
            dg = N[gaba].gt(thr, axis=0).mean(axis=1); do = N[other].gt(thr, axis=0).mean(axis=1)
            c = exp & (dg >= GABA_CUT) & (do <= OTHER_CUT)
            names = [ANCHORS.get(x, (x,))[0] for x in do[c].sort_values().index]
            rows.append(dict(tissue=tag, quantile=q, gad1_cut=cut, n_gaba=len(gaba),
                             n_in_corner=int(c.sum()), members="; ".join(names)))
    S = pd.DataFrame(rows)
    S.to_csv(P.out("gaba_cutoff_sensitivity.csv", outdir), index=False)
    return S


FIGSIZE = None
PANEL = False


def run(outdir=None):
    def O(n): return P.out(n, outdir)
    D, meta, rows = {}, {}, []
    for csv, tag in TISSUES:
        d, cut, n_gaba, n_other = table(csv)
        c = d[(d.det_gaba >= GABA_CUT) & (d.det_other <= OTHER_CUT)].sort_values("det_other")
        corner = [ANCHORS.get(g, (g,))[0] for g in c.index]
        D[tag] = d; meta[tag] = (cut, n_gaba, n_other, corner)
        for g in list(ANCHORS) + TARGETS:
            if g in d.index:
                rows.append(dict(tissue=tag, gene=ANCHORS.get(g, (g,))[0],
                                 det_gaba=d.det_gaba[g], det_other=d.det_other[g],
                                 in_corner=bool(d.det_gaba[g] >= GABA_CUT
                                                and d.det_other[g] <= OTHER_CUT)))
        rows.append(dict(tissue=tag, gene="[corner members]", det_gaba=np.nan,
                         det_other=np.nan, in_corner="; ".join(corner)))
        print(f"{tag}: Gad1 cut {cut:.2f} -> {n_gaba} GABAergic / {n_other} other clusters; "
              f"{len(c)} of {len(d)} expressed genes in the corner: "
              + ", ".join(f"{n} ({r.det_gaba:.0%}/{r.det_other:.0%})"
                          for n, (_, r) in zip(corner, c.iterrows())))
    T = pd.DataFrame(rows)
    T.to_csv(O("switchlike_gaba_genes.csv"), index=False)
    make_fig(D, meta, O("fig_switchlike_gaba.png"))
    S = sweep(outdir)
    print("\ncutoff sensitivity — the corner is sometimes empty, but never holds another gene:")
    print(S.to_string(index=False, float_format=lambda x: f"{x:.2f}"))
    return T


if __name__ == "__main__":
    ap = argparse.ArgumentParser(description=__doc__,
                                 formatter_class=argparse.RawDescriptionHelpFormatter)
    ap.add_argument("--outdir", default=None)
    ap.add_argument("--panel", action="store_true",
                    help="strip the suptitle and legend for a composite; they belong in "
                         "the caption")
    ap.add_argument("--figsize", nargs=2, type=float, default=None, metavar=("W", "H"))
    a = ap.parse_args()
    FIGSIZE, PANEL = (tuple(a.figsize) if a.figsize else None), a.panel
    run(a.outdir)
```
