## supplemental tables and scripts for "Long non-coding RNA-3′UTR interactions coordinate GABA signaling in the brain": fig_vgat_site_count.py

```
#!/usr/bin/env python3
"""
Manuscript panel: no gene in the Drosophila genome carries more conserved miR-7
sites than VGAT, and VGAT's three are all 8mers.

A  How many conserved miR-7 sites each predicted target carries, across all 183
   TargetScanFly 7.2 miR-7-5p predicted targets. 170 genes carry one, 12 carry
   two, and VGAT alone carries three.
B  The top of that ranking, with each gene's sites broken out by seed match.
   Filled segments are conserved sites; open segments are sites TargetScan calls
   poorly conserved, shown so the near-misses are visible rather than cropped.

Both panels use TargetScan's own per-gene conserved / poorly-conserved calls
(`TargetScan7.2__miR-7-5p.predicted_targets.txt`), which are PCT-based, with
genes.TRANSCRIPT_OVERRIDE applied: where TargetScan's representative transcript
is not the expressed one, the gene is scored on the designated transcript
instead. That drops Glts, whose sites are carried only by the long isoform
FBtr0345074 and not by the expressed FBtr0075344.

Caveat: the raw alignment table
(`dmel_conserved_sites.tsv`, the filtered Conserved_Family_Info) calls a site
conserved on alignment alone, ignoring PCT. By that looser definition 13 genes
carry three miR-7 sites and *two* carry three 8mers -- VGAT and E(spl)m5-HLH.
E(spl)m5-HLH's third 8mer has PCT = 0.00, i.e. conserved no better than chance
for a site of its type, which is why TargetScan's own export scores that gene
2 conserved + 1 poorly conserved and leaves VGAT the unique maximum. Panel B
shows that gene with its open segment for exactly this reason.

Usage:
    python3 fig_vgat_site_count.py        # writes to results/{tables,figures}
"""
import argparse
import re
import numpy as np, pandas as pd
import paths as P
import genes as GN
import matplotlib
matplotlib.use("Agg")
import matplotlib.pyplot as plt
from matplotlib.patches import Patch
from matplotlib.lines import Line2D

TARGETS = "TargetScan7.2__miR-7-5p.predicted_targets.txt"
SITES   = "dmel_conserved_sites.tsv"
MIR7_SEED = "GGAAGAC"                  # miR-7-5p family, per mir7_family_reference.txt
FOCUS   = "VGAT"
TYPES   = ["8mer", "7mer-m8", "7mer-A1"]
COLOR   = {"8mer": "#08306B", "7mer-m8": "#4C72B0", "7mer-A1": "#9ECAE1"}
FOCUS_C = "#C44E52"
N_TOP   = 12
NEAR_MISS = "E(spl)m5-HLH"   # the only other gene reaching three 8mers, on a PCT = 0 site


def load(datadir=None):
    """Per-gene site counts by type and conservation call, TargetScan's own."""
    T = pd.read_csv(P.find(TARGETS, datadir), sep="\t")
    keep = {"Target gene": "gene"}
    for t in TYPES:
        keep[f"Conserved {t.replace('A1', 'A1')} sites"] = f"cons_{t}"
        keep[f"Poorly conserved {t} sites"] = f"poor_{t}"
    T = T.rename(columns=keep)[list(keep.values())]
    T["gene"] = [GN.canon(g) for g in T.gene]
    T["conserved"] = T[[f"cons_{t}" for t in TYPES]].sum(axis=1)
    T["poorly"] = T[[f"poor_{t}" for t in TYPES]].sum(axis=1)
    return apply_overrides(T, datadir)


def apply_overrides(T, datadir=None):
    """Re-score genes whose expressed transcript is not TargetScan's representative.

    Counted off the designated transcript in the per-transcript conserved-site
    table; a gene left with no conserved site is not a predicted target and is
    dropped from the set rather than plotted as a zero.
    """
    if not GN.TRANSCRIPT_OVERRIDE:
        return T
    S = pd.read_csv(P.find(SITES, datadir), sep="\t")
    S.columns = [c.strip() for c in S.columns]
    S = S[S["miR Family"] == MIR7_SEED]
    drop = []
    for gene, tx in GN.TRANSCRIPT_OVERRIDE.items():
        i = T.index[T.gene == GN.canon(gene)]
        if not len(i):
            continue
        d = S[S["Transcript ID"].astype(str) == str(tx)]
        n = d["Seed match"].str.lower().replace({"7mer-a1": "7mer-A1"}).value_counts()
        for t in TYPES:
            T.loc[i, f"cons_{t}"] = int(n.get(t, 0))
            T.loc[i, f"poor_{t}"] = 0        # the site table holds conserved sites only
        T.loc[i, "conserved"] = int(n.sum())
        T.loc[i, "poorly"] = 0
        if not n.sum():
            drop.extend(i)
            print(f"note: {gene} scored on {tx} carries no conserved miR-7 site "
                  f"-- dropped from the predicted-target set")
    return T.drop(index=drop).reset_index(drop=True)


def alignment_counts(datadir=None):
    """The looser definition, for the caveat: conserved-by-alignment, PCT ignored."""
    S = pd.read_csv(P.find(SITES, datadir), sep="\t")
    S = S[S["miR Family"] == MIR7_SEED].copy()
    S["Gene Symbol"] = [GN.canon(g) for g in S["Gene Symbol"]]
    S["type"] = S["Seed match"].str.lower().replace({"7mer-a1": "7mer-A1"})
    tot = S.groupby("Gene Symbol").size()
    e8 = S[S.type == "8mer"].groupby("Gene Symbol").size()
    return S, tot, e8


ARMS = ["X", "2L", "2R", "3L", "3R", "4"]
GTF = "dmel-all-r6.68.gtf.gz"


def gene_coords(datadir=None):
    """{canonical symbol: (arm, start)} for every gene on a major arm, from the GTF."""
    import gzip
    out = {}
    with gzip.open(P.find(GTF, datadir), "rt") as fh:
        for ln in fh:
            if ln.startswith("#"):
                continue
            f = ln.split("\t")
            if len(f) < 9 or f[2] != "gene" or f[0] not in ARMS:
                continue
            m = re.search(r'gene_symbol "([^"]+)"', f[8]) or \
                re.search(r'gene_id "([^"]+)"', f[8])
            if m:
                out[GN.canon(m.group(1))] = (f[0], int(f[3]))
    return out


def lineup(outdir=None, datadir=None, width=3.45, height=None, select="8mer",
           ncol=1, tag=None, panel=True):
    """Fig. S3C: every gene in the genome with more than one conserved miR-7 site.

    One horizontal row per gene, segmented by seed match, VGAT at the top with
    three sites and the only gene whose three are all 8mers.

    HORIZONTAL, and that is the whole point. Three vertical designs were built
    first and every one failed the same way: a column is one gene wide, so the
    170 single-site genes crowd into solid blocks and out-weigh the tallest bar.
    RL, on the Manhattan version: "visually the opposite of what we want because
    VGAT has the smallest column". In rows, VGAT's bar is simply the longest mark
    on the page and its name sits beside it. Rejected, so they are not revisited:

      * one column per gene ranked by height -- a staircase, VGAT the first of a
        descending run rather than a lone spike
      * the same by genomic position, Manhattan-style -- position is irrelevant
        to the claim, so the axis spends its width on nothing, and that width is
        exactly what shrinks VGAT
      * the same as dots -- fixes the area problem, but y takes four values and a
        Manhattan needs a continuous y over orders of magnitude to read at all

    The set is COMPLETE, not a top-N: every gene with min_sites or more conserved
    sites. A cut at "more than one" is a criterion a reader can check; "the top
    12" is a number chosen to fit the page. The 170 genes with a single site are
    given as a count, since 182 rows would run to 16 inches.

    ONE THING THIS PANEL MUST NOT HIDE. "Conserved" is TargetScan's own PCT-based
    call. On the looser alignment-only definition (dmel_conserved_sites.tsv, PCT
    ignored) E(spl)m5-HLH also reaches three 8mers and VGAT ties rather than
    leads; the difference is a site with PCT = 0.00, conserved no better than
    chance for its type. Every poorly conserved site is drawn as an open segment,
    so the near-miss is visible rather than cropped.
    """
    def O(name): return P.out(name, outdir)
    plt.rcParams.update({
        "figure.dpi": 110, "savefig.dpi": 300,
        "axes.spines.top": False, "axes.spines.right": False,
        "svg.fonttype": "none", "font.family": "sans-serif",
        "font.sans-serif": ["Arial"], "axes.linewidth": 0.8})
    T = load(datadir)
    n_one = int((T.conserved == 1).sum())
    #: which genes get a row. "multi" is the complete set with more than one
    #: conserved site (12 rows); "8mer" every gene carrying a conserved 8mer of
    #: any number (48); "all" every predicted target (182, needs ncol > 1).
    SELECT = {
        "multi": (lambda d: d.conserved >= 2,
                  "every gene in the genome with more than one conserved miR-7 site",
                  lambda: f"({n_one} of the {len(T)} predicted targets carry a single site)"),
        "8mer":  (lambda d: d.cons_8mer >= 1,
                  "every gene in the genome carrying a conserved miR-7 8mer",
                  lambda: f"(of {len(T)} predicted targets; the rest carry 7mer sites only)"),
        "all":   (lambda d: d.conserved >= 1,
                  "all predicted miR-7 target genes in the genome",
                  lambda: f"({len(T)} genes, ranked by site count then seed match)"),
    }
    pick, title, sub = SELECT[select]
    D = T[pick(T)].sort_values(["conserved", "cons_8mer", "poorly"]).reset_index(drop=True)
    per = int(np.ceil(len(D) / ncol))
    # rows are ~0.145 in at 12 and tighten as the list grows; below ~0.075 the gene
    # names stop being legible, which is the real ceiling on how many fit
    row_h = 0.145 if per <= 14 else max(0.075, 2.05 / per)
    if height is None:
        # --panel drops the title and legend to the caption, as fig_gad1_enrichment
        # and fig_class_vs_six do, which is what lets this sit in a composite: the
        # two-line title also overruns the panel once the width drops below ~4.5 in
        height = per * row_h + (0.62 if panel else 1.30)
    fig, axs = plt.subplots(1, ncol, figsize=(width * ncol, height), squeeze=False)
    axs = axs[0]
    ax = axs[0]

    # itertuples renames "cons_7mer-m8" (the hyphen is not an identifier), so the
    # columns are read positionally off the row instead
    y = np.arange(len(D), dtype=float)
    lab_fs = 7.6 if per <= 14 else max(4.2, 90.0 / per)
    bar_h = 0.68 if per <= 14 else 0.80
    for i, r in D.iterrows():
        ax = axs[min(i // per, ncol - 1)]
        i = i % per
        left = 0.0
        for t in TYPES:                       # conserved sites, filled
            for _ in range(int(r[f"cons_{t}"])):
                ax.barh(i, 1, left=left, height=bar_h, color=COLOR[t],
                        edgecolor="white", linewidth=(0.9 if per <= 14 else 0.5), zorder=3)
                left += 1
        for t in TYPES:                       # poorly conserved, open
            for _ in range(int(r[f"poor_{t}"])):
                ax.barh(i, 1, left=left, height=bar_h, facecolor="none",
                        edgecolor=COLOR[t], linewidth=0.8,
                        linestyle=(0, (1.8, 1.4)), zorder=3)
                left += 1

    xmax = int((D.conserved + D.poorly).max())
    for c, a in enumerate(axs):
        names = list(D.gene[c * per:(c + 1) * per])
        a.set_yticks(np.arange(len(names)))
        a.set_yticklabels(names, fontsize=lab_fs)
        for t, g in zip(a.get_yticklabels(), names):
            t.set_fontstyle("italic")
            if g == FOCUS:
                t.set_color(FOCUS_C); t.set_fontweight("bold")
        a.tick_params(axis="y", length=0)
        a.set_ylim(-0.75, per - 0.25)
        a.set_xlim(0, xmax + 0.06)
        a.set_xticks(range(xmax + 1))
        a.tick_params(axis="x", labelsize=8)
        a.set_xlabel("miR-7 sites in the 3′UTR", fontsize=8.5)
    if not panel:
        axs[0].set_title(f"{title}\n{sub()}", fontsize=8, loc="left", color="0.3")

    j = int(D.index[D.gene == FOCUS][0])
    if per <= 14:
        axs[min(j // per, ncol - 1)].annotate(
            "three sites, all 8mers", (3.0, j % per), textcoords="offset points",
            xytext=(-7, 0), ha="right", va="center", fontsize=7.2,
            color="white", fontweight="bold", zorder=6)
    ax = axs[0]

    h = [Patch(facecolor=COLOR[t], edgecolor="white", linewidth=0.9, label=t)
         for t in TYPES]
    h += [Patch(facecolor="none", edgecolor="0.45", linewidth=0.8,
                linestyle=(0, (1.8, 1.4)), label="poorly conserved")]
    if not panel:
        axs[0].legend(handles=h, frameon=False, fontsize=6.9, ncol=4, loc="lower left",
                      bbox_to_anchor=(-0.11, -0.335 * (3.35 / height)),
                      handlelength=1.3, handletextpad=0.4, columnspacing=1.1)
    else:
        # In panel mode the legend has to live INSIDE the axes: the full-figure form
        # anchors it below the plot, which needs bottom margin the composite does not
        # have, and that is why panel mode used to drop it entirely and leave the four
        # encodings undecodable. The lower right is empty by construction -- the list is
        # sorted by site count, so the long tail of single-site genes leaves everything
        # past x = 1 blank for roughly three quarters of the rows.
        axs[-1].legend(handles=h, frameon=False, fontsize=6.4, loc="lower right",
                       handlelength=1.2, handletextpad=0.4, labelspacing=0.34,
                       borderaxespad=0.7)

    path = O(f"figS3C_mir7_site_ranking{tag or ''}.png")
    fig.tight_layout(rect=[0, (0.0 if panel else 0.085 * (3.35 / height)), 1, 1])
    fig.savefig(path); fig.savefig(path.replace(".png", ".svg"))
    plt.close(fig)
    T.to_csv(O("mir7_sites_per_target.csv"), index=False)
    print(f"wrote {path.replace('.png', '.svg')}  ({width:.2f} x {height:.2f} in)")
    print(f"  select={select}: {len(D)} rows ({per} per column x {ncol}), "
          f"{n_one} genes carry a single site, {len(T)} predicted targets")
    return path


def run(outdir=None, datadir=None):
    def O(name): return P.out(name, outdir)
    plt.rcParams.update({
        "figure.dpi": 110, "savefig.dpi": 300,
        "axes.spines.top": False, "axes.spines.right": False,
        "svg.fonttype": "none", "font.family": "sans-serif",
        "font.sans-serif": ["Arial"], "axes.linewidth": 0.8,
    })
    T = load(datadir)
    _, aln_tot, aln_8 = alignment_counts(datadir)

    fig, axes = plt.subplots(1, 2, figsize=(9.6, 4.3),
                             gridspec_kw=dict(width_ratios=[1, 1.45]))

    # ---- A: how many conserved sites per predicted target ----
    ax = axes[0]
    # carry the axis two counts past the maximum, so where the distribution stops
    # is something the reader sees rather than infers from the last bar drawn
    top_n = int(T.conserved.max())
    x = np.arange(1, top_n + 3)
    dist = T.conserved.value_counts().reindex(x, fill_value=0).sort_index()
    # split each bar into genes whose conserved sites are all 8mers, and the rest
    all8 = T[T.conserved > 0].assign(pure=lambda d: d["cons_8mer"] == d.conserved)
    pure = all8.groupby(["conserved", "pure"]).size().unstack(fill_value=0)
    b1 = pure.get(True, pd.Series(0, index=x)).reindex(x, fill_value=0).values
    b0 = pure.get(False, pd.Series(0, index=x)).reindex(x, fill_value=0).values
    ax.bar(x, b1, color=COLOR["8mer"], edgecolor="black", linewidth=0.6,
           label="all sites 8mer", zorder=3)
    ax.bar(x, b0, bottom=b1, color="#C8D3DE", edgecolor="black", linewidth=0.6,
           label="7mer site(s) included", zorder=3)
    for xi, n in zip(x, dist.values):
        ax.text(xi, max(n, 0.62) * 1.14, f"{n}", ha="center", fontsize=9,
                color="0.25" if n else "0.6")
    ax.annotate(f"three is the most any\n3'UTR carries ($\\it{{{FOCUS}}}$)",
                (top_n, dist[top_n]), textcoords="offset points", xytext=(-6, 46),
                ha="center", fontsize=9, color=FOCUS_C, linespacing=1.5,
                arrowprops=dict(arrowstyle="-", color=FOCUS_C, lw=1.0,
                                shrinkA=3, shrinkB=13))
    ax.set_yscale("log")
    ax.set_ylim(0.6, dist.max() * 4)
    ax.set_xticks(x)
    ax.set_xlabel("conserved miR-7 sites per gene", fontsize=9.5)
    ax.set_ylabel(f"genes (of {len(T)} predicted miR-7 targets)", fontsize=9.5)
    ax.tick_params(labelsize=8)
    ax.set_title("A", fontsize=11, loc="left", fontweight="bold")
    ax.legend(frameon=False, fontsize=7.5, loc="upper right")

    # ---- B: the top of the ranking, sites broken out by seed match ----
    ax = axes[1]
    top = T.sort_values(["conserved", "cons_8mer", "poorly"],
                        ascending=[False, False, False]).head(N_TOP)
    top = top.iloc[::-1]                                   # best at the top of the axis
    y = np.arange(len(top))
    left = np.zeros(len(top))
    for t in TYPES:
        w = top[f"cons_{t}"].values
        ax.barh(y, w, left=left, height=0.72, color=COLOR[t], edgecolor="black",
                linewidth=0.6, zorder=3)
        left += w
    for t in TYPES:
        w = top[f"poor_{t}"].values
        ax.barh(y, w, left=left, height=0.72, facecolor="white", edgecolor=COLOR[t],
                linewidth=0.9, linestyle=(0, (2.2, 1.4)), zorder=3)
        left += w
    ax.axvline(top_n, color=FOCUS_C, lw=0.9, ls=(0, (3.5, 2)), alpha=0.8, zorder=2)
    ax.text(top_n, len(top) - 0.25, f"{top_n} sites", ha="center", va="bottom",
            fontsize=8, color=FOCUS_C)
    ax.set_yticks(y)
    ax.set_yticklabels(top.gene, fontsize=8.5, style="italic")
    for tick, g in zip(ax.get_yticklabels(), top.gene):
        if g == FOCUS:
            tick.set_color(FOCUS_C); tick.set_fontweight("bold"); tick.set_fontsize(10)
    ax.set_xlabel("miR-7 sites in the 3'UTR", fontsize=9.5)
    ax.set_xticks(np.arange(0, int(left.max()) + 1))
    ax.tick_params(axis="x", labelsize=8)
    ax.margins(x=0.02)
    ax.set_title("B", fontsize=11, loc="left", fontweight="bold")
    handles = [Patch(facecolor=COLOR[t], edgecolor="black", linewidth=0.6, label=t)
               for t in TYPES]
    handles.append(Patch(facecolor="white", edgecolor="0.35", linewidth=0.9,
                         linestyle=(0, (2.2, 1.4)), label="poorly conserved"))
    ax.legend(handles=handles, frameon=False, fontsize=7.5, ncol=4,
              loc="upper center", bbox_to_anchor=(0.5, -0.14))

    fig.tight_layout(w_pad=2.2)
    p = O("fig_vgat_site_count.png")
    fig.savefig(p, bbox_inches="tight")
    fig.savefig(p.replace(".png", ".svg"), bbox_inches="tight")
    plt.close(fig)

    out = T.sort_values(["conserved", "cons_8mer"], ascending=False).copy()
    out["aln_sites_total"] = out.gene.map(aln_tot).fillna(0).astype(int)
    out["aln_sites_8mer"] = out.gene.map(aln_8).fillna(0).astype(int)
    out.to_csv(O("vgat_site_count.csv"), index=False)

    print(f"predicted miR-7 targets: {len(T)}")
    print(dist.rename("genes").to_frame().T.to_string())
    print()
    print(out.head(8).to_string(index=False))
    print()
    print("conserved-by-alignment (PCT ignored), for the caveat:")
    print(f"  genes with >=3 sites: {sorted(aln_tot[aln_tot >= 3].index.tolist())}")
    print(f"  genes with >=3 8mers: {sorted(aln_8[aln_8 >= 3].index.tolist())}")


if __name__ == "__main__":
    ap = argparse.ArgumentParser(description=__doc__,
                                 formatter_class=argparse.RawDescriptionHelpFormatter)
    ap.add_argument("--outdir", default=None,
                    help="write everything here; default routes to results/{tables,figures}")
    ap.add_argument("--datadir", default=None, help="extra directory to search for inputs")
    ap.add_argument("--select", choices=["multi", "8mer", "all"], default="8mer",
                    help="which genes get a row: >1 conserved site (default, 12), "
                         "any conserved 8mer (48), or every predicted target (182)")
    ap.add_argument("--ncol", type=int, default=1,
                    help="split the rows over this many columns")
    ap.add_argument("--width", type=float, default=3.45,
                    help="panel width in inches; the x range is only 0-3 sites, so "
                         "most of the width is gene names and this narrows well")
    ap.add_argument("--standalone", dest="panel", action="store_false",
                    help="add the title and legend back; they are off by default "
                         "because they belong in the caption, and the two-line title "
                         "overruns the panel below about 4.5 in wide")
    ap.add_argument("--lineup", action="store_true",
                    help="draw Fig. S3C instead: one bar per predicted target, "
                         "ranked by conserved 8mer count")
    a = ap.parse_args()
    if a.lineup:
        lineup(a.outdir, a.datadir, width=a.width, select=a.select, ncol=a.ncol,
               panel=a.panel,
               tag=(("" if a.select == "multi" else f"_{a.select}")
                    + ("_panel" if a.panel else "")) or None)
        # defaults reproduce the figure in use; other variants are reachable via flags
        raise SystemExit
    run(a.outdir, a.datadir)
```
