## supplemental tables and scripts for "Long non-coding RNA-3′UTR interactions coordinate GABA signaling in the brain": figS13_predictors.py

```
#!/usr/bin/env python3
"""Fig. S13: what predicts GABAergic co-variation among the 33 conserved 8mer targets.

Four panels on one A4 page (width 6.85 in, matching Fig. S12):
    A  other conserved miRNA families vs co-variation        rho = -0.55
    B  longest uninterrupted Gul complementarity vs co-variation   rho = +0.43
    C  the two factors together, coloured by co-variation    R2 = 0.36
    D  3'UTR length vs co-variation (the collinear alternative)

All panels are anchored on Gad1 with glia excluded, via
gul_duplex_model.build(anchor="Gad1", exclude_glia=True). One table supplies all
panels, so none can drift from another.

Two limits the panels are annotated with:
  - 3'UTR length and competing-family count are ~0.9 collinear and both predict the
    response (D is nearly as strong as A). At n = 33 the partials do not cleanly
    separate them (families | length = -0.16, length | families = -0.05), so the
    figure claims only that families is the interpretable variable and length adds
    nothing on top of it, not "families, not length".
  - Ssadh and CG32225 were found by ranking co-variation, so their position here is
    not independent evidence; the exclusion test (drop the four motivating genes) is
    printed by gul_duplex_model.py and belongs in the caption.

Usage:
    python3 figS13_predictors.py
"""
import argparse

import numpy as np
import matplotlib
matplotlib.use("Agg")
import matplotlib.pyplot as plt
from matplotlib.lines import Line2D
from scipy.stats import spearmanr

import paths as P
import genes as GN
import gul_duplex_model as DM

PAGE_W, PAGE_H = 6.85, 5.90
#: one type scale for the page; panels are ~3.2 in wide, so these are the sizes
#: that survive at print scale without the page having to be enlarged
SZ = dict(title=8.5, label=8.0, tick=7.0, gene=6.6, note=6.4)
FOCUS = list(GN.TARGET_POOL)
COL_OTH = "#8FA8BD"
COL_TREND = "#DD8452"
COUNTER = "stol"          # paragraph 40's counter-example


def gcolor(g):
    return GN.COLORS.get(g, COL_OTH)


#: candidate label offsets in points, tried in order. The six cluster tightly at
#: the top of every panel, so a fixed (+4.5, +3) offset stacks four names on top
#: of one another; this walks outward until the box is clear.
OFFSETS = [(6, 2), (6, -9), (6, 11), (6, -19), (6, 20), (6, -29), (6, 29),
           (-6, 2), (-6, -9), (-6, 11), (-6, -19), (-6, 20), (6, 38), (-6, 29)]
#: past this vertical displacement the label no longer reads as belonging to its
#: dot, so it gets a leader line -- the same fix the rank panel uses
LEADER_AT = 8


def place_labels(ax, items, fontsize, points=()):
    """Annotate (x, y, text, colour) avoiding overlap, in display coordinates.

    Box sizes are estimated from the string rather than measured: mathtext
    italics make get_window_extent expensive and the estimate only has to be
    good enough to pick between candidate offsets.

    `points` seeds the occupied set with the markers themselves, so a label
    cannot be parked on top of another gene's dot -- checking label against
    label alone left sgll sitting on CG10444.
    """
    ax.figure.canvas.draw()
    # transData returns PIXELS; offsets and font sizes are in POINTS. Mixing the
    # two made every estimated box ~35% too small, so labels were cleared against
    # boxes narrower than the text actually is and still collided.
    u = ax.figure.dpi / 72.0
    placed = []
    for x, y, r in points:
        px, py = ax.transData.transform((x, y))
        placed.append((px - r * u, py - r * u, px + r * u, py + r * u))
    for x, y, txt, col in items:
        px, py = ax.transData.transform((x, y))
        w, h = 0.58 * fontsize * len(txt) * u, 1.25 * fontsize * u
        for dx, dy in OFFSETS:
            x0 = px + dx * u if dx > 0 else px + dx * u - w
            y0 = py + dy * u - h / 2
            box = (x0, y0, x0 + w, y0 + h)
            if all(box[2] < b[0] or box[0] > b[2] or
                   box[3] < b[1] or box[1] > b[3] for b in placed):
                break
        placed.append(box)
        kw = {}
        if abs(dy) > LEADER_AT:
            kw["arrowprops"] = dict(arrowstyle="-", color=col, lw=0.5,
                                    shrinkA=0.5, shrinkB=2.5)
        ax.annotate(txt, (x, y), textcoords="offset points",
                    xytext=(dx, dy), va="center", style="italic",
                    ha=("left" if dx > 0 else "right"),
                    fontsize=fontsize, color=col, zorder=9, **kw)


def scatter(ax, R, xcol, xlabel, logx=False, label_extra=()):
    """One dot per transcript; the six and the counter-example carry their own colour."""
    named = set(FOCUS) | set(label_extra)
    lab, pts = [], []
    for g, r in R.iterrows():
        f = g in named
        ax.scatter([r[xcol]], [r.rho], s=(46 if f else 20),
                   facecolor=(gcolor(g) if f else COL_OTH),
                   edgecolor=("black" if f else "0.45"),
                   linewidth=(0.8 if f else 0.4), zorder=(6 if f else 4))
        if f:
            lab.append((r[xcol], r.rho, g, gcolor(g)))
        pts.append((r[xcol], r.rho, 5.2 if f else 3.8))
    ax.axhline(0, color="0.82", lw=0.8, ls="--", zorder=1)
    if logx:
        ax.set_xscale("log")
    ax.set_xlabel(xlabel, fontsize=SZ["label"])
    ax.set_ylabel("mean Spearman ρ with $\\it{Gad1}$", fontsize=SZ["label"])
    ax.tick_params(labelsize=SZ["tick"])
    ax.set_ylim(-0.42, 0.92)
    if logx:
        ax.set_xlim(R[xcol].min() * 0.55, R[xcol].max() * 1.9)
    else:
        pad = 0.09 * (R[xcol].max() - R[xcol].min())
        ax.set_xlim(R[xcol].min() - pad, R[xcol].max() + 2.2 * pad)
    place_labels(ax, lab, SZ["gene"], pts)


def trend(ax, x, y, logx=False):
    """Least-squares line for orientation only; every quoted statistic is rank-based."""
    xs = np.log10(np.asarray(x, float)) if logx else np.asarray(x, float)
    b = np.polyfit(xs, np.asarray(y, float), 1)
    g = np.linspace(xs.min(), xs.max(), 50)
    ax.plot(10 ** g if logx else g, np.polyval(b, g),
            color=COL_TREND, lw=1.6, zorder=2, alpha=0.9)


def stat_note(ax, text):
    ax.text(0.03, 0.03, text, transform=ax.transAxes, fontsize=SZ["note"],
            va="bottom", ha="left", family="monospace", color="0.25", zorder=10,
            bbox=dict(facecolor="white", alpha=0.82, edgecolor="none", pad=1.4))


def run(outdir=None, datadir=None):
    plt.rcParams.update({
        "figure.dpi": 110, "savefig.dpi": 200,
        "axes.spines.top": False, "axes.spines.right": False,
        "svg.fonttype": "none", "font.family": "sans-serif",
        "font.sans-serif": ["Arial"]})

    R, _, _ = DM.build(datadir, anchor="Gad1", exclude_glia=True)
    D = DM.diagnostics(R)
    r_fam, p_fam = spearmanr(R.n_families, R.rho)
    r_hel, p_hel = spearmanr(R.helix_gu, R.rho)
    r_len, p_len = spearmanr(R.UTR3_len, R.rho)

    fig, axes = plt.subplots(2, 2, figsize=(PAGE_W, PAGE_H))

    # ---- A  competing families -------------------------------------------------
    # Linear x, not log: miR-7 is excluded from the count, so sgll sits at zero.
    ax = axes[0][0]
    scatter(ax, R, "n_families", "other conserved miRNA families in 3′UTR",
            label_extra=[COUNTER])
    trend(ax, R.n_families, R.rho)
    ax.set_title("A   input from other miRNAs", fontsize=SZ["title"], loc="left")
    stat_note(ax, f"ρ = {r_fam:+.2f}  p = {p_fam:.3f}\n"
                  f"partial | UTR length = {DM.partial(R.rho, R.n_families, R.UTR3_len):+.2f}")

    # ---- B  duplex length ------------------------------------------------------
    ax = axes[0][1]
    scatter(ax, R, "helix_gu", "uninterrupted $\\it{Gul}$ complementarity (bp)",
            label_extra=[COUNTER])
    trend(ax, R.helix_gu, R.rho)
    ax.set_title("B   extent of $\\it{Gul}$ complementarity", fontsize=SZ["title"], loc="left")
    stat_note(ax, f"ρ = {r_hel:+.2f}  p = {p_hel:.3f}\n"
                  f"partial | other families = {DM.partial(R.rho, R.helix_gu, R.n_families):+.2f}")

    # ---- C  the two factors together ------------------------------------------
    ax = axes[1][0]
    sc = ax.scatter(R.helix_gu, R.n_families, c=R.rho, cmap="RdBu_r",
                    vmin=-0.7, vmax=0.7,
                    s=[62 if g in FOCUS else 28 for g in R.index],
                    edgecolor="black",
                    linewidth=[0.8 if g in FOCUS else 0.4 for g in R.index], zorder=4)
    place_labels(ax,
                 [(r.helix_gu, r.n_families, g, "black") for g, r in R.iterrows()
                  if g in FOCUS or g == COUNTER],
                 SZ["gene"],
                 [(r.helix_gu, r.n_families, 5.6 if g in FOCUS else 4.0)
                  for g, r in R.iterrows()])
    # symlog, NOT log: miR-7 is excluded from the competing-family count, so sgll
    # sits at exactly 0 and a log axis silently DROPS IT -- the one transcript with
    # no competitors at all, missing from the panel that argues competitors matter.
    # linthresh=1 keeps 0 on the axis while the 1-57 range still reads logarithmically.
    ax.set_yscale("symlog", linthresh=1, linscale=0.45)
    ax.set_yticks([0, 1, 2, 5, 10, 20, 50])
    ax.yaxis.set_major_formatter(matplotlib.ticker.FuncFormatter(
        lambda v, _: f"{v:.0f}"))
    ax.yaxis.set_minor_locator(matplotlib.ticker.NullLocator())
    ax.set_ylim(-0.35, 90)
    ax.set_xlim(R.helix_gu.min() - 1.5, R.helix_gu.max() + 5.0)
    ax.set_xlabel("uninterrupted complementarity (bp)", fontsize=SZ["label"])
    ax.set_ylabel("other miRNA families", fontsize=SZ["label"])
    ax.tick_params(labelsize=SZ["tick"])
    ax.set_title(f"C   miRNA input and complementarity together    R² = {D['r2']:.2f}, "
                 f"permutation p = {D['perm_p']:.4f}",
                 fontsize=SZ["title"], loc="left")
    ax.annotate("long complementarity,\nfew other families", xy=(0.97, 0.93), xycoords="axes fraction",
                ha="right", va="top", fontsize=SZ["note"], color="0.35")
    cb = fig.colorbar(sc, ax=ax, fraction=0.045, pad=0.03)
    cb.set_label("mean ρ with $\\it{Gad1}$", fontsize=SZ["note"])
    cb.ax.tick_params(labelsize=SZ["note"])
    # LOO R2 and the bootstrap CI go in the caption; the panel carries the two
    # numbers the text quotes

    # ---- D  the collinear alternative -----------------------------------------
    ax = axes[1][1]
    scatter(ax, R, "UTR3_len", "3′UTR length (nt)", logx=True, label_extra=[COUNTER])
    ax.set_title("D   3′UTR length, the collinear alternative",
                 fontsize=SZ["title"], loc="left")
    stat_note(ax, f"ρ = {r_len:+.2f}  p = {p_len:.3f}\n"
                  f"partial | other families = {DM.partial(R.rho, R.UTR3_len, R.n_families):+.2f}")

    h = [Line2D([], [], marker="o", ls="none", mfc=gcolor(g), mec="black", mew=0.8,
                ms=6, label=g) for g in FOCUS + [COUNTER]]
    h += [Line2D([], [], marker="o", ls="none", mfc=COL_OTH, mec="0.45", mew=0.4,
                 ms=5, label="other conserved-8mer miR-7 targets"),
          Line2D([], [], color=COL_TREND, lw=1.6,
                 label="least-squares line (orientation only)")]
    fig.legend(handles=h, frameon=False, ncol=5, loc="lower center",
               bbox_to_anchor=(0.5, -0.006), fontsize=SZ["note"],
               handletextpad=0.35, columnspacing=1.1)
    fig.tight_layout(rect=[0, 0.072, 1, 1.0], h_pad=1.5, w_pad=1.8)

    path = P.out("figS13_covariation_predictors.png", outdir)
    fig.savefig(path)
    fig.savefig(path.replace(".png", ".svg"))
    plt.close(fig)
    print(f"wrote {path.replace('.png', '.svg')}  ({PAGE_W:.2f} x {PAGE_H:.2f} in)")
    print(f"  A4 is 8.27 x 11.69 in -> margins {(8.27-PAGE_W)/2:.2f} in each side, "
          f"{(11.69-PAGE_H)/2:.2f} in top and bottom")
    print(f"  n = {len(R)} conserved-8mer targets, anchor Gad1, glia excluded")
    print(f"  families {r_fam:+.3f}   helix {r_hel:+.3f}   UTR length {r_len:+.3f}")
    print(f"  R2 {D['r2']:.3f}  LOO {D['loo']:.3f}  perm p {D['perm_p']:.4f}  "
          f"boot 95% CI [{D['boot_lo']:.3f}, {D['boot_hi']:.3f}]")
    return path


if __name__ == "__main__":
    ap = argparse.ArgumentParser(description=__doc__,
                                 formatter_class=argparse.RawDescriptionHelpFormatter)
    ap.add_argument("--outdir", default=None)
    ap.add_argument("--datadir", default=None)
    a = ap.parse_args()
    run(a.outdir, a.datadir)
```
