## supplemental tables and scripts for "Long non-coding RNA-3′UTR interactions coordinate GABA signaling in the brain": genes.py

```
#!/usr/bin/env python3
"""One canonical symbol per gene, and the gene sets the analyses share.

The three cluster-mean matrices were built against different FlyBase releases,
so the *same gene* carries different symbols in different tissues. CB is the
newest: it says `ScsbetaG`, `Gabat`, `Regnase-1` where OL and VNC still say
`Sucb`, `CG7433`, `CG10889`. Of the 549 conserved miR-7 targets, **28 are split
this way**, and a further 30 were being missed in CB entirely because the
TargetScan site-type lookup is keyed on the old symbols — those genes were
silently counted as background.

`canon()` maps every known alias to the FlyBase r6.68 symbol, so all three
tissues, the TargetScan tables and the figure labels use one name per gene.
Apply it to a matrix index with `canon_index()` before anything else touches it.

The alias table is derived, not hand-written: FBgn -> r6.68 symbol comes from
`data/genome/fbgn_symbol_r6.68.tsv` (extracted from the r6.68 GTF), and
FBgn -> old symbol from the TargetScanFly 7.2 conserved-target list. MANUAL
covers the handful of genes that are not miR-7 targets and so appear in neither.

No matrix contains two aliases of one gene, and no canonical name collides with
a different gene's old symbol, so canon() is a pure relabel with nothing to merge.
"""
import pandas as pd
import paths as P

SYMBOL_TSV = "fbgn_symbol_r6.68.tsv"
TARGET_TSV = "mir7_targetscanfly_conserved_targets.tsv"

# genes that are not miR-7 targets, so the TargetScan list cannot supply them
MANUAL = {
    "CG7433": "Gabat",        # FBgn0036927 - OL uses the old symbol
    "Cha": "ChAT",            # FBgn0000303 - OL uses the old symbol; VNC and CB say ChAT
}

# ---------------------------------------------------------------------------
# the shared gene sets: define them ONCE, in canonical symbols
# ---------------------------------------------------------------------------
ANCHOR = "CG14989"            # Gul (FBgn0035495); still the current symbol
GAD1   = "Gad1"

#: the highlighted genes, in figure order. Gfrl is excluded: not a conserved 8mer
#: target with a locatable site, its apparent Gul match tracing to a low-complexity
#: AT-repeat.
SHOWN = ["VGAT", "sgll", "ScsbetaG", "Glts", "stol", "P5CDh1", "Gabat"]
#: which of SHOWN are genuine conserved miR-7 targets. Glts is excluded: TargetScan
#: calls it a 7mer-m8 target on a long representative transcript, but the expressed
#: 837 nt 3'UTR carries no miR-7 seed match of any class. It remains a shown
#: gene and a putative Gul target; only its miR-7 status changed.
SHOWN_TARGETS = ["VGAT", "sgll", "ScsbetaG", "stol"]
#: shown genes that are NOT miR-7 targets (drawn with a diamond marker)
NONTARGET = ["Glts", "P5CDh1", "Gabat"]

#: TargetScan annotates one representative transcript per gene, and for some genes
#: that transcript is not the expressed one. Sites for these are re-derived from
#: the per-transcript conserved-site table using the transcript named here rather
#: than taken from the per-gene summary.
#:
#: Glts: TargetScan uses FBtr0345074, a long isoform carrying three miR-7 7mer-m8
#: matches upstream. The expressed transcript is FBtr0075344, whose 837 nt 3'UTR
#: carries no miR-7 seed match of any class - and which TargetScan annotates with
#: no conserved site for any miRNA family, so it belongs in the no-site background.
TRANSCRIPT_OVERRIDE = {"Glts": "FBtr0075344"}
MIR7_FAMILY = "GGAAGAC"


def override_site_types(datadir=None):
    """{gene: best conserved miR-7 site type on its designated transcript, or None}."""
    if not TRANSCRIPT_OVERRIDE:
        return {}
    S = pd.read_csv(P.find("dmel_conserved_sites.tsv", datadir), sep="\t")
    S.columns = [c.strip() for c in S.columns]
    S = S[S["miR Family"] == MIR7_FAMILY]
    order = {"8mer": 0, "7mer-m8": 1, "7mer-a1": 2, "7mer-A1": 2}
    out = {}
    for g, tx in TRANSCRIPT_OVERRIDE.items():
        d = S[S["Transcript ID"].astype(str) == str(tx)]
        if not len(d):
            out[g] = None
            continue
        best = sorted(d["Seed match"].astype(str), key=lambda x: order.get(x, 9))[0]
        out[g] = "7mer-A1" if best.lower() == "7mer-a1" else best
    return out


#: genes labelled in the site-type panels
PANEL = ["VGAT", "sgll", "ScsbetaG", "stol", "Ssadh", "CG32225", "CG10444"]

#: the six conserved-8mer targets at the top of the Gad1 ranking, glia excluded.
#: CG10444 belongs here on every expression criterion but is NOT part of the GABA
#: functional module.
TARGET_POOL = ["VGAT", "sgll", "Ssadh", "ScsbetaG", "CG32225", "CG10444"]

#: the five of TARGET_POOL that are in the GABA pathway; used where the claim is
#: about the module rather than about the ranking
MODULE = ["VGAT", "sgll", "Ssadh", "ScsbetaG", "CG32225"]

#: fixed colour per gene, shared by every figure that labels genes. All twelve must
#: stay mutually distinguishable: fig_switchlike_gaba shows them in one panel.
COLORS = {"VGAT": "#E41A1C", "sgll": "#FF7F00", "ScsbetaG": "#4DAF4A",
          "Glts": "#E7298A", "stol": "#A65628",
          "P5CDh1": "#0072B2", "Gabat": "#C9A400", "Ssadh": "#6A3D9A",
          "CG32225": "#00A0A0", "CG10444": "#595959"}
# CG10444 is deliberately grey, not a hue: it belongs to the ranking but not to the
# GABA module, and an olive/gold would have collided with Gabat's #C9A400.


def _build():
    cur = pd.read_csv(P.find(SYMBOL_TSV), sep="\t")
    fb2sym = dict(zip(cur.FBgn.astype(str), cur.symbol.astype(str)))
    t = pd.read_csv(P.find(TARGET_TSV), sep="\t")
    old = {str(a).split(".")[0]: str(b) for a, b in zip(t.iloc[:, 0], t.iloc[:, -1])}
    m = {o: fb2sym[f] for f, o in old.items() if f in fb2sym and fb2sym[f] != o}
    m.update(MANUAL)
    return m


#: {alias: canonical symbol}; canonical symbols are absent (they map to themselves)
ALIASES = _build()


def canon(sym):
    """Canonical FlyBase r6.68 symbol for one gene symbol."""
    return ALIASES.get(str(sym), str(sym))


def canon_all(syms):
    """Canonicalise any iterable of symbols, preserving type where it matters."""
    out = [canon(s) for s in syms]
    return set(out) if isinstance(syms, (set, frozenset)) else out


def canon_index(df):
    """Relabel a genes-x-clusters matrix index in place-ish (returns a new frame).

    Raises if a rename would collide, which would mean two rows for one gene and
    a silent choice about which to keep.
    """
    idx = [canon(g) for g in df.index]
    dup = pd.Index(idx)[pd.Index(idx).duplicated()]
    if len(dup):
        raise ValueError(f"canonicalising created duplicate rows: {sorted(set(dup))}")
    out = df.copy()
    out.index = idx
    return out


def canon_keys(d):
    """Canonicalise the keys of a {symbol: value} dict."""
    return {canon(k): v for k, v in d.items()}


if __name__ == "__main__":
    print(f"{len(ALIASES)} aliases known; e.g.")
    for k in sorted(ALIASES)[:10]:
        print(f"  {k:16s} -> {ALIASES[k]}")
    print(f"\nSHOWN          {SHOWN}")
    print(f"SHOWN_TARGETS  {SHOWN_TARGETS}")
    print(f"PANEL          {PANEL}")
    print(f"TARGET_POOL    {TARGET_POOL}")
    print(f"MODULE         {MODULE}")
```
