## supplemental tables and scripts for "Long non-coding RNA-3′UTR interactions coordinate GABA signaling in the brain": gul_duplex_model.py

```
#!/usr/bin/env python3
"""
Two-factor model for Gul sensitivity of conserved-8mer miR-7 targets.

Factor 1  DUPLEX  -- length of the longest contiguous Gul:target helix anchored
                     on the miR-7 8mer site (Watson-Crick, optionally G:U).
                     Gul's 3'UTR carries three tandem copies of TGGAAGACA, a
                     miR-7 5'-end mimic, so it competes for the target site
                     rather than sequestering the miRNA (Gul contains no miR-7
                     binding site at all).
Factor 2  COMPETITION -- number of distinct conserved miRNA families with sites
                     in the same 3'UTR; more competing input dilutes any single
                     miRNA's contribution.

The model: a target is Gul-sensitive when Gul can form a decent helix AND the
UTR is not dominated by other miRNA inputs. Neither factor alone suffices.

The response variable is the mean of three per-region Spearman correlations. Two
choices about it are exposed as flags, because both move the result:

  --anchor {Gad1,Gul}   which gene the targets are correlated against. Gad1 is
                        the default and what the manuscript uses (the two are
                        0.88-0.93 collinear, and Gad1 leaves the sequence screen
                        and the co-variation screen with no variable in common).
  --keep-glia           retain glial clusters. The default drops them (repo >=
                        0.17, plus OL's named G- clusters), which is the
                        manuscript specification; the claim is about GABAergic
                        vs non-GABAergic *neurons*.

Usage:
    python3 gul_duplex_model.py                          # default (Gad1, glia excluded)
    python3 gul_duplex_model.py --anchor Gul --keep-glia # alternative setting
"""
import os, re, argparse
from collections import defaultdict
import numpy as np, pandas as pd
import paths as P
import genes as GN
import gul_mir7_correlation as G
import matplotlib
matplotlib.use("Agg")
import matplotlib.pyplot as plt
from matplotlib.lines import Line2D
from scipy.stats import spearmanr, rankdata

UTR_FASTA = P.find("dmel-all-three_prime_UTR-r6.58.fasta")
GUL_FB = "FBgn0035495"
SITE = "GTCTTCCA"
FOCUS = list(GN.TARGET_POOL)
WC = {("A","T"),("T","A"),("G","C"),("C","G")}
WOB = {("G","T"),("T","G")}


def utr_seqs():
    seqs = defaultdict(str); name=None; buf=[]
    for line in open(UTR_FASTA):
        if line.startswith(">"):
            if name:
                q="".join(buf).upper()
                if len(q)>len(seqs[name]): seqs[name]=q
            m=re.search(r"parent=(FBgn\d+)",line); name=m.group(1) if m else None; buf=[]
        elif name: buf.append(line.strip())
    if name:
        q="".join(buf).upper()
        if len(q)>len(seqs[name]): seqs[name]=q
    return seqs


def helix_len(t, site, gr, wobble):
    """longest contiguous run of valid pairs covering the miR-7 site"""
    def ok(a,b): return (a,b) in WC or (wobble and (a,b) in WOB)
    best = 0
    for j in range(len(gr)):
        i2,j2,L = site,j,0
        while i2 < len(t) and j2 < len(gr) and ok(t[i2],gr[j2]): i2+=1; j2+=1; L+=1
        if L < 8: continue          # must at least cover the 8mer
        i3,j3 = site-1, j-1
        while i3>=0 and j3>=0 and ok(t[i3],gr[j3]): i3-=1; j3-=1; L+=1
        best = max(best, L)
    return best


ANCHORS = {"Gul": "CG14989", "Gad1": "Gad1"}
TISSUES = [("OL", "gene_by_cluster_meanexpr.csv"),
           ("VNC", "gene_by_cluster_meanexpr_VNC.csv"),
           ("CB", "gene_by_cluster_meanexpr_CB.csv")]


def response(anchor, exclude_glia, datadir=None):
    """Per-region Spearman rho against the anchor, plus expression-pattern covariates.

    Returns (rho: genes x {OL, VNC, CB}, cov: genes x four measures averaged over the
    three regions).

    The covariates exist to answer the obvious objection to the helix predictor -- that a
    longer Gul:target helix might simply mark genes with a particular expression pattern,
    so the correlation would be a confound rather than a mechanism. All four are computed
    here rather than ad hoc, because that is how they went stale: they were last run on the
    Gul-anchored, glia-included response and were the only analysis in the paper still on
    that specification.

      breadth   fraction of clusters in which the gene is detected (mean above 10% of its
                own 90th-percentile value) -- the same rule broad_class_test.py uses
      dyn       p90 / p10 dynamic range, the switch-like-ness measure from section 1
      retained  clusters kept by the dead-tail filter, as a fraction of the region's total
      expr      mean expression across clusters
    """
    cols, covs = {}, {}
    for tag, csv in TISSUES:
        M = GN.canon_index(pd.read_csv(P.find(csv, datadir), index_col=0))
        if exclude_glia:
            M = G.drop_glia(M, tag)
        A = G.analyze(tag, M, ANCHORS[anchor], [], [], set())["res"]
        cols[tag] = A.rho
        p90 = M.quantile(.90, axis=1).reindex(A.index)
        p10 = M.quantile(.10, axis=1).reindex(A.index)
        det = M.gt(0.1 * M.quantile(.90, axis=1), axis=0).mean(axis=1).reindex(A.index)
        covs[tag] = pd.DataFrame(dict(breadth=det,
                                      dyn=(p90 + 1e-9) / (p10 + 1e-9),
                                      retained=A.n_clusters / M.shape[1],
                                      expr=A.mean_expr))
    RR = pd.DataFrame(cols).dropna()
    CV = sum(covs[t].reindex(RR.index) for t, _ in TISSUES) / len(TISSUES)
    return RR, CV


COVARIATES = [("breadth", "detection breadth"), ("dyn", "dynamic range (p90/p10)"),
              ("retained", "clusters retained"), ("expr", "mean expression")]


def controls(R, CV):
    """Is either predictor a proxy for an expression pattern? Rows: one covariate each."""
    C = CV.reindex(R.index)
    rows = []
    for key, label in COVARIATES:
        z = C[key].values
        r_h, p_h = spearmanr(R.helix_gu, z)
        r_f, p_f = spearmanr(R.n_families, z)
        rows.append(dict(covariate=label,
                         rho_helix_cov=r_h, p_helix_cov=p_h,
                         rho_families_cov=r_f, p_families_cov=p_f,
                         helix_partial=partial(R.helix_gu, R.rho, z),
                         families_partial=partial(R.n_families, R.rho, z)))
    return pd.DataFrame(rows)


def build(datadir, anchor="Gul", exclude_glia=False):
    seqs = utr_seqs(); gr = seqs[GUL_FB][::-1]
    tgt = pd.read_csv(P.find("mir7_targetscanfly_conserved_targets.tsv", datadir), sep="\t")
    S2F = {GN.canon(s): fb for s, fb in zip(tgt.GeneSymbol.astype(str),
                                            tgt.GeneID.astype(str))}
    F = pd.read_csv(P.find("gul_8mer_mirna_competition.csv", datadir), index_col=0)
    # the competition table supplies n_families / UTR length; the response is
    # recomputed here so --anchor and --exclude-glia actually take effect
    RR, CV = response(anchor, exclude_glia, datadir)
    F = F[F.index.isin(RR.index)].copy()
    F["rho_mean"] = RR.mean(axis=1).reindex(F.index)
    rows=[]
    for g in F.index:
        s = seqs.get(S2F.get(g,""), "")
        pos = [m.start() for m in re.finditer(SITE, s)] if s else []
        if not pos: continue
        rows.append(dict(gene=g, rho=F.loc[g,"rho_mean"],
                         helix_wc=max(helix_len(s,p,gr,False) for p in pos),
                         helix_gu=max(helix_len(s,p,gr,True) for p in pos),
                         n_families=F.loc[g,"n_families"],
                         UTR3_len=F.loc[g,"UTR3_len"],
                         n_sites_8mer=len(pos)))
    return pd.DataFrame(rows).set_index("gene"), RR, CV


def sp(a,b): return spearmanr(a,b)[0]
def partial(x,y,z):
    rxy,rxz,ryz = sp(x,y),sp(x,z),sp(y,z)
    return (rxy-rxz*ryz)/np.sqrt(max((1-rxz**2)*(1-ryz**2),1e-12))


def model(R):
    """rank-based two-factor regression + interaction"""
    y = rankdata(R.rho)
    h = rankdata(R.helix_gu); c = rankdata(R.n_families)
    h = (h-h.mean())/h.std(); c = (c-c.mean())/c.std(); y=(y-y.mean())/y.std()
    X1 = np.column_stack([np.ones(len(y)), h])
    X2 = np.column_stack([np.ones(len(y)), c])
    X3 = np.column_stack([np.ones(len(y)), h, c])
    X4 = np.column_stack([np.ones(len(y)), h, c, h*c])
    out={}
    for nm,X in [("helix only",X1),("competition only",X2),
                 ("helix + competition",X3),("+ interaction",X4)]:
        b,*_ = np.linalg.lstsq(X,y,rcond=None)
        resid = y - X@b
        r2 = 1 - resid.var()/y.var()
        n,k = len(y), X.shape[1]-1
        adj = 1-(1-r2)*(n-1)/(n-k-1)
        out[nm]=(r2,adj,b)
    return out


def _design(R):
    y = rankdata(R.rho); h = rankdata(R.helix_gu); c = rankdata(R.n_families)
    z = lambda v: (v - v.mean()) / v.std()
    return z(y), np.column_stack([np.ones(len(y)), z(h), z(c)])


def _r2(y, X):
    b, *_ = np.linalg.lstsq(X, y, rcond=None)
    return 1 - (y - X @ b).var() / y.var()


def diagnostics(R, nboot=2000, nperm=20000, seed=0):
    """Out-of-sample and null checks on the two-factor model."""
    y, X = _design(R); n = len(y)
    pred = np.empty(n)
    for i in range(n):
        m = np.ones(n, bool); m[i] = False
        b, *_ = np.linalg.lstsq(X[m], y[m], rcond=None)
        pred[i] = X[i] @ b
    loo = 1 - ((y - pred) ** 2).sum() / ((y - y.mean()) ** 2).sum()

    rng = np.random.default_rng(seed)
    obs = _r2(y, X)
    null = np.array([_r2(rng.permutation(y), X) for _ in range(nperm)])
    boot = []
    for _ in range(nboot):
        i = rng.integers(0, n, n)
        if len(np.unique(i)) < 5: continue
        try: boot.append(_r2(y[i], X[i]))
        except np.linalg.LinAlgError: pass
    boot = np.array(boot)
    return dict(r2=obs, loo=loo, perm_p=(null >= obs).mean(),
                null_med=np.median(null), null_p95=np.quantile(null, .95),
                boot_lo=np.quantile(boot, .025), boot_hi=np.quantile(boot, .975))


def noise_ceiling(RR, genes):
    """Spearman-Brown reliability of the 3-region mean response."""
    S = RR.reindex(genes).dropna()
    pairs = [("OL", "VNC"), ("OL", "CB"), ("VNC", "CB")]
    rs = {f"{a}-{b}": spearmanr(S[a], S[b])[0] for a, b in pairs}
    rbar = np.mean(list(rs.values()))
    return rs, rbar, 3 * rbar / (1 + 2 * rbar)


def make_fig(R, path, anchor='Gul'):
    plt.rcParams.update({
        "figure.dpi":110,"savefig.dpi":200,"axes.spines.top":False,
        "axes.spines.right":False,"svg.fonttype":"none",
        "font.family":"sans-serif","font.sans-serif":["Arial"]})
    CT,CO,CTR = "#C44E52","#5B7C99","#DD8452"
    fig,axes = plt.subplots(1,3,figsize=(12.2,4.4))

    def trend(ax, x, y, logx=False):
        """Least-squares line, drawn for orientation only.

        The model and every quoted statistic are rank-based (Spearman, and a rank
        regression), so this line is not the fit being tested - it just shows the
        direction and rough steepness. On the competition panel it is fitted in
        log10(x), matching that axis.
        """
        xs = np.log10(np.asarray(x, float)) if logx else np.asarray(x, float)
        b = np.polyfit(xs, np.asarray(y, float), 1)
        g = np.linspace(xs.min(), xs.max(), 50)
        ax.plot(10**g if logx else g, np.polyval(b, g),
                color=CTR, lw=2, zorder=2, alpha=0.9)

    def base(ax,x,xlabel,logx=False):
        for g,r in R.iterrows():
            f = g in FOCUS
            ax.scatter([r[x]],[r.rho],s=(78 if f else 34),
                       facecolor=(CT if f else CO),edgecolor="black",
                       linewidth=(1.1 if f else 0.5),zorder=(6 if f else 4))
            if f or abs(r.rho)>0.15:
                ax.annotate(f"$\\it{{{g}}}$",(r[x],r.rho),textcoords="offset points",
                            xytext=(6,4),fontsize=7,zorder=8,
                            color=(CT if f else "0.35"))
        ax.axhline(0,color="0.8",lw=0.9,ls="--",zorder=1)
        if logx: ax.set_xscale("log")
        ax.set_xlabel(xlabel,fontsize=9.5); ax.set_ylabel(f"mean Spearman ρ with {anchor}",fontsize=9.5)
        ax.tick_params(labelsize=8); ax.set_ylim(-0.42,0.82)

    ax=axes[0]; base(ax,"helix_gu","longest contiguous Gul:target helix (bp)")
    trend(ax, R.helix_gu, R.rho)
    r,p = spearmanr(R.helix_gu,R.rho)
    ax.set_title(f"A  duplex length\nρ = {r:+.3f}, p = {p:.3f};  partial | competition = "
                 f"{partial(R.rho,R.helix_gu,R.n_families):+.3f}",
                 fontsize=10,loc="left")

    # linear, not log: miR-7 is excluded from the competing-family count, so sgll
    # sits at zero and log10(0) is undefined. Counts span 0-31.
    ax=axes[1]; base(ax,"n_families","competing conserved miRNA families")
    trend(ax, R.n_families, R.rho)
    r,p = spearmanr(R.n_families,R.rho)
    ax.set_title(f"B  competing miRNA input\nρ = {r:+.3f}, p = {p:.3f};  partial | duplex = "
                 f"{partial(R.rho,R.n_families,R.helix_gu):+.3f}",
                 fontsize=10,loc="left")

    # C: two-factor plane
    ax=axes[2]
    sc=ax.scatter(R.helix_gu,R.n_families,c=R.rho,cmap="RdBu_r",vmin=-0.7,vmax=0.7,
                  s=[110 if g in FOCUS else 55 for g in R.index],
                  edgecolor="black",linewidth=[1.1 if g in FOCUS else 0.5 for g in R.index],
                  zorder=4)
    for g,r in R.iterrows():
        if g in FOCUS or abs(r.rho)>0.15:
            ax.annotate(f"$\\it{{{g}}}$",(r.helix_gu,r.n_families),
                        textcoords="offset points",xytext=(7,3),fontsize=7,
                        color=("black" if g in FOCUS else "0.4"),zorder=8)
    # symlog, not log: sgll carries ZERO competing families (miR-7 excluded from the
    # count), and a log axis would drop it silently -- the one gene with no competitors
    # at all, absent from the panel that argues competitors matter.
    ax.set_yscale("symlog", linthresh=1, linscale=0.45)
    ax.set_yticks([0, 1, 2, 5, 10, 20, 50])
    ax.yaxis.set_major_formatter(matplotlib.ticker.FuncFormatter(lambda v, _: f"{v:.0f}"))
    ax.yaxis.set_minor_locator(matplotlib.ticker.NullLocator())
    ax.set_ylim(-0.35, 90)
    ax.set_xlim(R.helix_gu.min()-1.5, R.helix_gu.max()+4.5)   # room for the VGAT label
    ax.set_xlabel("longest contiguous helix (bp)",fontsize=9.5)
    ax.set_ylabel("competing miRNA families",fontsize=9.5)
    ax.set_title("C  the two factors together\n ",fontsize=10,loc="left")
    ax.tick_params(labelsize=8)
    ax.annotate("Gul-sensitive corner →\n(long helix, few competitors)",
                xy=(0.97,0.90),xycoords="axes fraction",ha="right",fontsize=7,color="0.35")
    cb=fig.colorbar(sc,ax=ax,fraction=0.046,pad=0.03); cb.set_label(f"mean ρ with {anchor}",fontsize=8)
    cb.ax.tick_params(labelsize=7)

    h=[Line2D([],[],marker="o",ls="none",mfc=CT,mec="black",mew=1.1,ms=8,
              label=", ".join(FOCUS)),
       Line2D([],[],marker="o",ls="none",mfc=CO,mec="black",mew=0.5,ms=7,
              label="other conserved-8mer miR-7 targets"),
       Line2D([],[],color=CTR,lw=2,label="least-squares line (orientation only; "
              "quoted statistics are rank-based)")]
    fig.legend(handles=h,frameon=False,ncol=3,loc="lower center",
               bbox_to_anchor=(0.5,-0.03),fontsize=8.5)
    fig.suptitle("Gul sensitivity requires both a viable duplex and low competing miRNA input",
                 fontsize=11)
    fig.tight_layout(rect=[0,0.07,1,0.93],w_pad=1.3)
    fig.savefig(path,bbox_inches="tight"); fig.savefig(path.replace(".png",".svg"),bbox_inches="tight")
    plt.close(fig)


if __name__ == "__main__":
    ap=argparse.ArgumentParser(description=__doc__,
        formatter_class=argparse.RawDescriptionHelpFormatter)
    ap.add_argument("--outdir", default=None,
                    help="write everything here; default routes to results/{tables,figures}")
    ap.add_argument("--datadir", default=None, help="extra directory to search for inputs")
    ap.add_argument("--anchor", choices=sorted(ANCHORS), default="Gad1",
                    help="correlate targets against this gene (default Gad1)")
    ap.add_argument("--keep-glia", action="store_true",
                    help="retain glial clusters; the default drops repo-positive and named G- clusters, which is the manuscript specification")
    a=ap.parse_args()
    a.exclude_glia = not a.keep_glia
    O = lambda n: P.out(n, a.outdir)
    tag = "" if (a.anchor == "Gad1" and a.exclude_glia) else \
          f"_{a.anchor.lower()}" + ("" if a.exclude_glia else "_glia")
    R, RR, CV = build(a.datadir, a.anchor, a.exclude_glia)
    R.sort_values("rho",ascending=False).to_csv(O(f"gul_duplex_scan{tag}.csv"))
    make_fig(R,O(f"fig_gul_two_factor{tag}.png"), a.anchor)
    print(f"n={len(R)} conserved-8mer targets   anchor={a.anchor}  "
          f"glia={'excluded' if a.exclude_glia else 'included'}\n")
    print("SINGLE FACTORS")
    for c in ["helix_wc","helix_gu","n_families"]:
        r,p=spearmanr(R[c],R.rho); print(f"  {c:<12} rho={r:+.3f} p={p:.4f}")
    print("\nPARTIALS")
    print(f"  helix | competition    {partial(R.rho,R.helix_gu,R.n_families):+.3f}")
    print(f"  competition | helix    {partial(R.rho,R.n_families,R.helix_gu):+.3f}")
    print(f"  (helix vs competition collinearity: {sp(R.helix_gu,R.n_families):+.3f})")
    print("\nRANK REGRESSION (standardised)")
    for nm,(r2,adj,b) in model(R).items():
        print(f"  {nm:<22} R2={r2:.3f}  adjR2={adj:.3f}  coef={np.round(b[1:],3)}")
    print("\nEXCLUSION (drop the 4 motivating genes)")
    d=R.drop(["Ssadh","CG32225","sgll","ScsbetaG"],errors="ignore")
    for c in ["helix_gu","n_families"]:
        r,p=spearmanr(d[c],d.rho); print(f"  {c:<12} rho={r:+.3f} p={p:.3f} n={len(d)}")

    CT = controls(R, CV)
    CT.to_csv(O(f"gul_expression_controls{tag}.csv"), index=False)
    print("\nEXPRESSION-PATTERN CONTROLS (is either predictor a proxy for a pattern?)")
    print(f"  raw:  helix {sp(R.helix_gu, R.rho):+.3f}   families {sp(R.n_families, R.rho):+.3f}")
    print(f"  {'covariate':<24}{'rho w/ helix':>14}{'rho w/ fam':>12}"
          f"{'helix|cov':>12}{'families|cov':>14}")
    for _, r in CT.iterrows():
        print(f"  {r.covariate:<24}{r.rho_helix_cov:>+9.3f} (p{r.p_helix_cov:>4.2f})"
              f"{r.rho_families_cov:>+7.3f}{r.helix_partial:>+12.3f}{r.families_partial:>+14.3f}")

    D = diagnostics(R)
    print("\nDIAGNOSTICS (helix + competition)")
    print(f"  in-sample R2           {D['r2']:.3f}")
    print(f"  leave-one-out R2       {D['loo']:.3f}")
    print(f"  permutation p          {D['perm_p']:.4f}  "
          f"(null median {D['null_med']:.3f}, 95th pct {D['null_p95']:.3f})")
    print(f"  bootstrap 95% CI       [{D['boot_lo']:.3f}, {D['boot_hi']:.3f}]")
    rs, rbar, rel = noise_ceiling(RR, R.index)
    print("\nNOISE CEILING")
    print("  between-region  " + "  ".join(f"{k} {v:+.3f}" for k, v in rs.items())
          + f"   mean {rbar:+.3f}")
    print(f"  Spearman-Brown reliability of the 3-region mean  {rel:.3f}"
          f"   -> R2 is {100*D['r2']/rel:.0f}% of the ceiling")

    print("\nTOP OF THE RANKING")
    print(R.sort_values("rho", ascending=False)
           .head(8)[["rho", "helix_gu", "n_families", "UTR3_len"]]
           .to_string(float_format=lambda v: f"{v:.3f}"))
```
