## supplemental tables and scripts for "Long non-coding RNA-3′UTR interactions coordinate GABA signaling in the brain": gul_mir7_correlation.py

```
#!/usr/bin/env python3
"""
Gul (CG14989) vs miR-7 target correlation across FCA cluster-mean tables.

Tests whether a hand-picked set of "shown" genes co-varies with the anchor
gene Gul more strongly than (a) non-target background genes and (b) the miR-7
target class as a whole, at the level of per-cluster mean expression.

Pipeline (identical for every gene, foreground and background):
  1. Rank a gene's cluster-mean expression descending.
  2. "Dead-tail onset" inclusive filter: keep clusters whose mean is above
     baseline + 2%*span, where baseline = median of the bottom 20% of ranked
     clusters. This drops only the flat, near-zero tail and keeps the whole
     informative range (deliberately inclusive, not a strict knee).
  3. Spearman rho between Gul and the gene over the retained clusters.
  4. Group comparisons via one-sided Mann-Whitney + a 20,000-iteration
     permutation test on the difference of medians, plus a cluster-count-matched
     null control (shown genes retain more clusters on average, so matched
     background genes are drawn at the same breadth).

Inputs are cluster-mean expression matrices (genes x clusters, gene symbols in
the first column) as written by AverageExpression() on the RNA "data" layer.

Usage:
    python3 gul_mir7_correlation.py                 # writes to results/{tables,figures}
    python3 gul_mir7_correlation.py --outdir ./out  # or all into one directory
    python3 gul_mir7_correlation.py --config my.json

Input files are located by name (see paths.py); see DEFAULT_CONFIG at the bottom
for the expected config shape.
"""
import os, sys, json, argparse
import numpy as np
import pandas as pd
from scipy.stats import spearmanr, mannwhitneyu
import paths as P
import genes as GN
import matplotlib
matplotlib.use("Agg")
import matplotlib.pyplot as plt
from matplotlib.lines import Line2D

# ----------------------------------------------------------------------------
# Constants
# ----------------------------------------------------------------------------
MIN_CL   = 5           # never keep fewer than this many clusters
DEAD_FRAC = 0.02       # tail threshold = baseline + DEAD_FRAC * span
N_PERM   = 20000       # permutation iterations
PERM_SEED = 0
MATCH_SEED = 7
MATCH_TOL  = 25        # +/- n_clusters window for the matched-null control
MATCH_DRAWS = 20000
META_GREY = "#8A8A8A"
COL_MIR7  = "#4C72B0"  # blue  - miR-7 target class
COL_SHOWN = "#C44E52"  # red   - shown genes

def _style():
    plt.rcParams.update({
        "figure.dpi": 110, "savefig.dpi": 200,
        "axes.spines.top": False, "axes.spines.right": False,
        "axes.titlesize": 9, "axes.labelsize": 8,
        "xtick.labelsize": 7, "ytick.labelsize": 7,
        "font.family": "sans-serif", "svg.fonttype": "none",
    })

# ----------------------------------------------------------------------------
# Core filter + correlation
# ----------------------------------------------------------------------------
def chord_d(s):
    """Chord distance of a descending curve (for the strict-knee reference line)."""
    y = np.asarray(s, float); n = len(y); span = y[0] - y[-1]
    if span <= 0: return np.zeros(n)
    xr = np.arange(n) / (n - 1); yn = (y - y[-1]) / span
    return (1 - xr) - yn

def knee_argmax(s):
    return int(np.argmax(chord_d(s)))

def dead_tail_k(s):
    """Number of leading clusters to keep under the dead-tail onset rule."""
    base = np.median(s[int(0.8 * len(s)):]); span = s[0] - base
    if span <= 0: return len(s)
    return int((s >= base + DEAD_FRAC * span).sum())

def perm_test(fg, bg, n=N_PERM, seed=PERM_SEED, stat="median"):
    """One-sided permutation p-value: P(shuffled difference >= observed).

    ``stat`` selects the summary being compared.

    **median** (default) matches what the figures draw as their group summary
    bar. Use it for any claim about whether the typical gene in a class differs
    from background.

    **mean** answers a different question, being moved by a handful of extreme
    genes: the conserved-8mer stratum gives p = 0.0000/0.0000/0.0015 on means and
    0.23/0.12/0.10 on medians, the difference being entirely the six proposed
    targets at rho +0.4 to +0.9. Both are real; keep them apart.

    The pool is **sorted** before the first shuffle, so the p-value depends only
    on the two multisets and the seed, never on the order the rows arrived in.
    Without that, ``gaba_derepression_class.py`` and ``fig_class_vs_six.py`` --
    which run the identical test on the identical genes, but reach them through
    a merge that reorders rows -- would otherwise disagree in the third decimal
    (e.g. VNC 7mer-m8 0.0594 vs 0.0582).
    """
    f = np.mean if stat == "mean" else np.median
    rng = np.random.default_rng(seed); obs = f(fg) - f(bg)
    pool = np.sort(np.concatenate([fg, bg])); k = len(fg); c = 0
    for _ in range(n):
        rng.shuffle(pool)
        if f(pool[:k]) - f(pool[k:]) >= obs: c += 1
    return obs, (c + 1) / (n + 1)

# ----------------------------------------------------------------------------
# Analysis for one matrix
# ----------------------------------------------------------------------------
def analyze(tag, M, anchor, shown_list, target_ids, mir7_symbols):
    """
    tag           : label for outputs ("OL"/"VNC"/"CB")
    M             : DataFrame genes x clusters
    anchor        : anchor gene symbol (Gul = CG14989)
    shown_list    : the 8 highlighted genes for this table (order preserved)
    target_ids    : which of shown_list are genuine miR-7 targets (by identity)
    mir7_symbols  : full miR-7 target symbol set (TargetScanFly conserved)
    """
    gul = M.loc[anchor].values.astype(float)
    mir7_in = set(mir7_symbols) & set(M.index)

    def corr_inc(gene, detail=False):
        v = M.loc[gene].values.astype(float); order = np.argsort(-v); s = v[order]
        k = int(min(max(dead_tail_k(s), MIN_CL), len(s))); keep = order[:k]
        if np.std(v[keep]) == 0 or np.std(gul[keep]) == 0: return None
        rho, p = spearmanr(gul[keep], v[keep])
        if np.isnan(rho): return None
        o = dict(gene=gene, rho=rho, p=p, n_clusters=k, cutoff=float(s[k-1]))
        if detail: o.update(s=s, k=k)
        return o

    res = pd.DataFrame([r for r in (corr_inc(g) for g in M.index if g != anchor) if r]).set_index("gene")
    res["is_mir7"]  = res.index.isin(mir7_in)
    res["is_shown"] = res.index.isin(shown_list)
    res["mean_expr"] = M.loc[res.index].mean(axis=1).values
    # shown genes: set target status by explicit identity, not raw symbol lookup
    for g in shown_list:
        if g in res.index: res.loc[g, "is_mir7"] = g in target_ids

    bg  = res[~res.is_mir7 & ~res.is_shown]
    mir = res[res.is_mir7]
    sh  = res.loc[[g for g in shown_list if g in res.index]]
    sh_t = sh.loc[[g for g in target_ids if g in sh.index]]

    tests = []
    for name, fg, bs in [("miR-7 targets vs background", mir, bg),
                         ("shown vs background", sh, bg),
                         ("shown vs other miR-7", sh, mir[~mir.index.isin(sh.index)]),
                         ("shown miR-7 targets vs background", sh_t, bg)]:
        pm = mannwhitneyu(fg.rho, bs.rho, alternative="greater").pvalue
        _, pp = perm_test(fg.rho.values, bs.rho.values)                  # medians
        _, pmean = perm_test(fg.rho.values, bs.rho.values, stat="mean")  # tail-sensitive
        tests.append(dict(comparison=name, n_fg=len(fg), med_fg=fg.rho.median(),
                          med_bg=bs.rho.median(), mannwhitney_p=pm, perm_p=pp,
                          perm_p_mean=pmean))

    # cluster-count-matched null: draw background genes at matched breadth
    rng = np.random.default_rng(MATCH_SEED)
    def matched(fg, tol=MATCH_TOL, nd=MATCH_DRAWS):
        pools = {}
        for g in fg.index:
            k = fg.loc[g, "n_clusters"]
            cand = bg[(bg.n_clusters >= k - tol) & (bg.n_clusters <= k + tol)]
            pools[g] = cand.rho.values if len(cand) >= 20 else bg.rho.values
        null = np.array([np.median([rng.choice(pools[g]) for g in fg.index]) for _ in range(nd)])
        obs = fg.rho.median()
        return obs, np.median(null), np.quantile(null, .95), max((null >= obs).mean(), 1/(nd+1))
    o8, m8, q8, p8 = matched(sh); o6, m6, q6, p6 = matched(sh_t)
    tests += [dict(comparison="shown vs cluster-matched null", n_fg=len(sh),
                   med_fg=o8, med_bg=m8, mannwhitney_p=np.nan, perm_p=p8),
              dict(comparison="shown miR-7 targets vs cluster-matched null", n_fg=len(sh_t),
                   med_fg=o6, med_bg=m6, mannwhitney_p=np.nan, perm_p=p6)]

    return dict(tag=tag, res=res, sh=sh, sh_t=sh_t, bg=bg, mir=mir,
                mir7_in=mir7_in, gul=gul, ncl=M.shape[1],
                T=pd.DataFrame(tests), corr_inc=corr_inc)

# ----------------------------------------------------------------------------
# Figures
# ----------------------------------------------------------------------------
def ecdf(v):
    v = np.sort(v); return v, np.arange(1, len(v)+1) / len(v)

def _spread(ys, gap):
    """Push labels apart to a minimum vertical gap, then recenter the block on
    the original values so labels stay next to their dots (no upward drift)."""
    order = np.argsort(ys); s = ys[order].astype(float)
    for i in range(1, len(s)):
        if s[i] - s[i-1] < gap: s[i] = s[i-1] + gap
    s += ys.mean() - s.mean()
    out = np.empty_like(s); out[order] = s
    return out

TISSUE_NAME = {"OL": "Optic lobe", "VNC": "Ventral nerve cord", "CB": "Central brain"}

# symbols are canonicalised at load (genes.py), so there is one name per gene
# fixed colour per shown gene (blue/grey reserved for the group clouds)
GENE_ORDER  = list(GN.SHOWN)
GENE_COLORS = dict(GN.COLORS)
NONTARGET   = set(GN.NONTARGET)            # shown but not miR-7 targets -> diamond marker

def make_combined_fig(results, order, path):
    """Compact horizontal 3-tissue group-strip (fits A4 width). Shown genes are
    colour-coded via one shared legend instead of per-panel text labels."""
    _style()
    plt.rcParams.update({"font.family": "sans-serif", "font.sans-serif": ["Arial"]})
    XP = [0.0, 0.9, 1.8]                       # tightened column spacing
    fig, axes = plt.subplots(1, 3, figsize=(8.0, 5.0))
    for ax, tag in zip(np.atleast_1d(axes), order):
        A = results[tag]; T = A["T"]; bg = A["bg"]; mir = A["mir"]; sh = A["sh"]
        for i, (vals, c) in enumerate([(bg.rho.values, META_GREY), (mir.rho.values, COL_MIR7)]):
            jit = (np.random.default_rng(i).random(len(vals)) - 0.5) * 0.30
            ax.scatter(np.full(len(vals), XP[i]) + jit, vals, s=8, alpha=0.15,
                       color=c, edgecolor="none", zorder=3, rasterized=True)
            ax.plot([XP[i]-0.22, XP[i]+0.22], [np.median(vals)]*2, color="black", lw=1.8, zorder=4)
        ax.plot([XP[2]-0.22, XP[2]+0.22], [np.median(sh.rho.values)]*2, color="black", lw=1.8, zorder=4)
        genes = list(sh.index)
        rng = np.random.default_rng(2)
        xj = XP[2] + (rng.random(len(genes)) - 0.5) * 0.26
        for gx, g in zip(xj, genes):
            t7 = bool(sh.loc[g, "is_mir7"])
            ax.scatter([gx], [sh.loc[g, "rho"]], s=46, zorder=6, linewidth=0.6,
                       marker=("o" if t7 else "D"),
                       facecolor=GENE_COLORS[g], edgecolor="black")
        ax.axhline(0, color="0.8", lw=0.9, ls="--")
        ax.set_xticks(XP); ax.set_xticklabels(["Bg", "miR-7", "shown"], fontsize=9)
        ax.set_xlim(-0.4, 2.2); ax.tick_params(axis="y", labelsize=8.5)
        def bracket(x1, x2, yy, txt):
            ax.plot([x1, x1, x2, x2], [yy, yy+0.03, yy+0.03, yy], lw=1, color="black")
            ax.text((x1+x2)/2, yy+0.04, txt, ha="center", va="bottom", fontsize=8)
        p_sh  = T.loc[T.comparison == "shown vs background", "perm_p"].values[0]
        p_sho = T.loc[T.comparison == "shown vs other miR-7", "perm_p"].values[0]
        fmt = lambda p: f"p={p:.4f}" if p >= 1e-4 else "p<0.0001"
        base = max(sh.rho.max(), mir.rho.max()) + 0.12
        bracket(XP[0], XP[2], base, fmt(p_sh)); bracket(XP[1], XP[2], base + 0.18, fmt(p_sho))
        ax.set_ylim(-1.12, base + 0.44)
        ax.set_title(f"{TISSUE_NAME.get(tag, tag)} ({tag})\n{A['ncl']} clusters", fontsize=10)
        ax.set_ylabel("Spearman ρ with Gul", fontsize=9.5)
    # one shared legend: group clouds + colour-coded genes
    grp = [Line2D([], [], marker="o", ls="none", mfc=META_GREY, mec="none", ms=6, label="Background"),
           Line2D([], [], marker="o", ls="none", mfc=COL_MIR7, mec="none", ms=6, label="miR-7 targets"),
           Line2D([], [], color="black", lw=1.8, label="group median")]
    gene_h = [Line2D([], [], marker=("o" if g not in NONTARGET else "D"), ls="none",
                     mfc=GENE_COLORS[g], mec="black", mew=0.6, ms=7,
                     label=(g if g not in NONTARGET else g + " (not a target)"))
              for g in GENE_ORDER]
    leg1 = fig.legend(handles=grp, frameon=False, ncol=3, loc="lower center",
                      bbox_to_anchor=(0.5, 0.135), fontsize=8.5)
    fig.add_artist(leg1)
    fig.legend(handles=gene_h, frameon=False, ncol=4, loc="lower center",
               bbox_to_anchor=(0.5, 0.0), fontsize=8.5,
               title="Shown genes  (circle = miR-7 target, diamond = not)", title_fontsize=8.5)
    fig.suptitle("Gul correlation: shown genes vs miR-7 target class vs background", fontsize=11)
    fig.tight_layout(rect=[0, 0.19, 1, 0.94], w_pad=1.0)
    fig.savefig(path, bbox_inches="tight")
    fig.savefig(path.replace(".png", ".svg"), bbox_inches="tight")
    plt.close(fig)

def make_main_fig(A, shown_list, tag, path):
    _style()
    res = A["res"]; bg = A["bg"]; mir = A["mir"]; sh = A["sh"]; T = A["T"]
    allrho = np.sort(res.rho.values)
    fig, axes = plt.subplots(1, 2, figsize=(11, 4.6), gridspec_kw=dict(width_ratios=[1.35, 1]))
    ax = axes[0]
    x, y = ecdf(bg.rho.values);  ax.plot(x, y, color=META_GREY, lw=1.8)
    x, y = ecdf(mir.rho.values); ax.plot(x, y, color=COL_MIR7, lw=1.8)
    for g in shown_list:
        if g not in sh.index: continue
        r = sh.loc[g, "rho"]; frac = (allrho < r).mean(); is7 = bool(sh.loc[g, "is_mir7"])
        ax.plot(r, frac, marker="o", ms=7, mfc=(COL_SHOWN if is7 else "white"),
                mec=COL_SHOWN, mew=1.6, zorder=5)
    ax.axvline(0, color="0.8", lw=0.8, ls="--")
    ax.set_xlabel("Spearman correlation with Gul (inclusive per-gene filter)")
    ax.set_ylabel("Cumulative fraction of genes")
    ax.set_title(f"{tag}: Gul-correlation of shown genes vs miR-7 targets vs background", fontsize=8)
    ln = [Line2D([], [], color=META_GREY, lw=1.8, label=f"Non-target background (n={len(bg)})"),
          Line2D([], [], color=COL_MIR7, lw=1.8, label=f"All miR-7 targets (n={len(mir)})")]
    mk = [Line2D([], [], marker="o", ls="none", mfc=COL_SHOWN, mec=COL_SHOWN, ms=7, label="Shown gene — miR-7 target"),
          Line2D([], [], marker="o", ls="none", mfc="white", mec=COL_SHOWN, mew=1.6, ms=7, label="Shown gene — not a target")]
    l1 = ax.legend(handles=ln, frameon=False, loc="upper left", fontsize=6.5); ax.add_artist(l1)
    ax.legend(handles=mk, frameon=False, loc="lower right", fontsize=6.5)
    ax = axes[1]
    groups = [("Background", bg.rho.values, META_GREY),
              ("miR-7\ntargets", mir.rho.values, COL_MIR7),
              ("shown", sh.rho.values, COL_SHOWN)]
    for i, (name, vals, c) in enumerate(groups):
        jit = (np.random.default_rng(i).random(len(vals)) - 0.5) * 0.32
        ax.scatter(np.full(len(vals), i) + jit, vals, s=(10 if i < 2 else 42),
                   alpha=(0.15 if i < 2 else 0.9), color=c,
                   edgecolor=("none" if i < 2 else "black"), linewidth=0.5, zorder=3)
        ax.plot([i-0.25, i+0.25], [np.median(vals)]*2, color="black", lw=2, zorder=4)
    ax.axhline(0, color="0.8", lw=0.8, ls="--")
    ax.set_xticks(range(3)); ax.set_xticklabels([g[0] for g in groups])
    ax.set_ylabel("Spearman correlation with Gul"); ax.set_title("Group medians (black bars)", fontsize=8)
    def bracket(x1, x2, yy, txt):
        ax.plot([x1, x1, x2, x2], [yy, yy+0.03, yy+0.03, yy], lw=1, color="black")
        ax.text((x1+x2)/2, yy+0.04, txt, ha="center", va="bottom", fontsize=6.5)
    ytop = max(sh.rho.max(), mir.rho.max())
    p_sh  = T.loc[T.comparison == "shown vs background", "perm_p"].values[0]
    p_sho = T.loc[T.comparison == "shown vs other miR-7", "perm_p"].values[0]
    bracket(0, 2, ytop+0.08, f"p={p_sh:.4f}"); bracket(1, 2, ytop+0.20, f"p={p_sho:.4f}")
    ax.margins(x=0.15)
    fig.tight_layout(); fig.savefig(path, bbox_inches="tight")
    fig.savefig(path.replace(".png", ".svg"), bbox_inches="tight")
    plt.close(fig)

def make_elbow_fig(A, shown_list, tag, path):
    _style()
    ncl = A["ncl"]
    fig, axes = plt.subplots(2, 4, figsize=(13, 6.4))
    for ax, g in zip(axes.ravel(), shown_list):
        if g not in A["sh"].index:
            ax.axis("off"); continue
        d = A["corr_inc"](g, detail=True); s = d["s"]; k = d["k"]; k_strict = knee_argmax(s) + 1
        x = np.arange(1, len(s)+1)
        ax.plot(x, s, color=META_GREY, lw=1.2); ax.plot(x[:k], s[:k], color=COL_SHOWN, lw=2.0)
        ax.axvline(k, color=COL_MIR7, lw=1.2); ax.axvline(k_strict, color="0.55", lw=1, ls=":")
        ax.scatter([k], [s[k-1]], color=COL_MIR7, s=26, zorder=5)
        is7 = bool(A["sh"].loc[g, "is_mir7"])
        ax.set_title(f"$\\it{{{g}}}$  {'(miR-7)' if is7 else '(not target)'}\n"
                     f"keep {k}/{ncl}   ρ={d['rho']:+.2f}", fontsize=7.5)
        ax.set_xlabel("cluster rank", fontsize=7); ax.set_ylabel("mean expr", fontsize=7)
        ax.text(0.96, 0.90, f"cut≥{s[k-1]:.2f}", transform=ax.transAxes, ha="right",
                fontsize=6.5, color=COL_MIR7)
    axes[0, 0].legend(handles=[Line2D([], [], color=COL_MIR7, lw=1.2, label="inclusive cut (used)"),
                               Line2D([], [], color="0.55", lw=1, ls=":", label="strict knee (rejected)")],
                      fontsize=6, frameon=False, loc="upper right")
    fig.suptitle(f"{tag} — per-gene inclusive filter: keep above the dead tail (red)", fontsize=9)
    fig.tight_layout(rect=[0, 0, 1, 0.97]); fig.savefig(path, bbox_inches="tight")
    fig.savefig(path.replace(".png", ".svg"), bbox_inches="tight")
    plt.close(fig)

def make_tissue_heatmap(piv, is7_map, path):
    _style()
    order = (piv.assign(m=piv.mean(1), t=[is7_map[g] for g in piv.index])
                .sort_values(["t", "m"], ascending=[False, False]).index.tolist())
    P = piv.loc[order]
    fig, ax = plt.subplots(figsize=(5.2, 4.4))
    im = ax.imshow(P.values, cmap="RdBu_r", vmin=-0.9, vmax=0.9, aspect="auto")
    ax.set_xticks(range(P.shape[1])); ax.set_xticklabels(P.columns)
    ax.set_yticks(range(len(P)))
    ax.set_yticklabels([f"$\\it{{{g}}}$" + ("" if is7_map[g] else "  (not target)") for g in P.index], fontsize=8)
    for i in range(len(P)):
        for j in range(P.shape[1]):
            v = P.values[i, j]
            if np.isnan(v): continue
            ax.text(j, i, f"{v:+.2f}", ha="center", va="center", fontsize=7.5,
                    color=("white" if abs(v) > 0.55 else "black"))
    cb = fig.colorbar(im, ax=ax, fraction=0.046, pad=0.04); cb.set_label("Spearman ρ with Gul", fontsize=8)
    ax.set_title("Gul correlation of shown genes across tissues", fontsize=9)
    fig.tight_layout(); fig.savefig(path, bbox_inches="tight")
    fig.savefig(path.replace(".png", ".svg"), bbox_inches="tight")
    plt.close(fig)

# ----------------------------------------------------------------------------
# miR-7 target list loader
# ----------------------------------------------------------------------------
REPO_MAX = 0.17          # repo above this marks a glial cluster (validated on OL's named glia)


def drop_glia(M, tag=""):
    """Remove glial clusters: repo >= REPO_MAX, plus OL's named ``G-`` clusters.

    The claim throughout is about GABAergic versus non-GABAergic *neurons*, so
    every correlation should be computed on neurons. Leaving glia in flips signs
    for glial-enriched genes: CG10444's OL rho with Gad1 is -0.285 with glia and
    +0.475 without.
    """
    if "repo" not in M.index:
        return M
    repo = M.loc["repo"]
    bad = (set(repo[repo >= REPO_MAX].index)
           | {c for c in M.columns if str(c).startswith("G-")})
    keep = [c for c in M.columns if c not in bad]
    if tag:
        print(f"[{tag}] {len(bad)} glial clusters dropped, {len(keep)} kept")
    return M[keep]


def load_mir7_symbols(path):
    """Read the TargetScanFly conserved-target TSV (GeneID<TAB>GeneSymbol)."""
    df = pd.read_csv(path, sep="\t")
    col = "GeneSymbol" if "GeneSymbol" in df.columns else df.columns[-1]
    syms = GN.canon_all(set(df[col].astype(str)))
    # a gene with no miR-7 site on its expressed transcript is not a target, so it
    # must not land in the "poorly conserved sites only" stratum either
    return syms - {g for g, v in GN.override_site_types().items() if v is None}

# ----------------------------------------------------------------------------
# Driver
# ----------------------------------------------------------------------------
def run(config, outdir=None, keep_glia=False):
    def O(name): return P.out(name, outdir)
    anchor = config["anchor"]
    mir7 = load_mir7_symbols(P.find(config["mir7_targets_tsv"]))

    results = {}
    long_rows = []
    for t in config["tissues"]:
        tag = t["tag"]
        M = GN.canon_index(pd.read_csv(P.find(t["matrix_csv"]), index_col=0))
        if not keep_glia:
            M = drop_glia(M, tag)
        assert anchor in M.index, f"{anchor} missing from {tag}"
        shown = t["shown"]
        target_ids = t["targets"]                 # subset of shown that are genuine targets
        mir7_syms = set(mir7) | set(t.get("mir7_extra", []))  # e.g. add ScsbetaG in CB
        A = analyze(tag, M, anchor, shown, target_ids, mir7_syms)
        results[tag] = A
        # per-tissue outputs
        A["res"].sort_values("rho", ascending=False).to_csv(O(f"gul_correlation_all_genes_{tag}.csv"))
        A["sh"].to_csv(O(f"gul_correlation_shown8_{tag}.csv"))
        A["T"].to_csv(O(f"gul_correlation_significance_tests_{tag}.csv"), index=False)
        make_main_fig(A, shown, tag, O(f"fig_gul_mir7_correlation_{tag}.png"))
        make_elbow_fig(A, shown, tag, O(f"fig_elbow_per_gene_{tag}.png"))
        for g in A["sh"].index:
            long_rows.append(dict(tissue=tag, gene=g,
                                  rho=A["sh"].loc[g, "rho"], n_clusters=int(A["sh"].loc[g, "n_clusters"]),
                                  is_mir7=bool(A["sh"].loc[g, "is_mir7"])))
        print(f"[{tag}] {M.shape[0]} genes x {M.shape[1]} clusters | "
              f"miR-7 present {len(A['mir7_in'])} | shown vs bg perm p="
              f"{A['T'].loc[A['T'].comparison=='shown vs background','perm_p'].values[0]:.4f} | "
              f"miR-7 class perm p="
              f"{A['T'].loc[A['T'].comparison=='miR-7 targets vs background','perm_p'].values[0]:.4f}")

    # cross-tissue summary
    combined = pd.DataFrame(long_rows)
    combined.to_csv(O("gul_correlation_shown_by_tissue_long.csv"), index=False)
    piv = combined.pivot_table(index="gene", columns="tissue", values="rho").round(3)
    tags = [t["tag"] for t in config["tissues"]]
    piv = piv[[c for c in tags if c in piv.columns]]
    piv.to_csv(O("gul_correlation_shown_by_tissue.csv"))
    is7_map = combined.groupby("gene").is_mir7.max().to_dict()
    make_tissue_heatmap(piv, is7_map, O("fig_shown_rho_by_tissue.png"))
    make_combined_fig(results, tags, O("fig_gul_mir7_correlation_3tissue.png"))

    tsum = []
    for tag in tags:
        T = results[tag]["T"].set_index("comparison")
        tsum.append(dict(tissue=tag,
            mir7_vs_bg_perm=T.loc["miR-7 targets vs background", "perm_p"],
            mir7_vs_bg_perm_mean=T.loc["miR-7 targets vs background", "perm_p_mean"],
            shown_vs_bg_perm=T.loc["shown vs background", "perm_p"],
            six_vs_bg_perm=T.loc["shown miR-7 targets vs background", "perm_p"],
            shown_med=T.loc["shown vs background", "med_fg"],
            bg_med=T.loc["shown vs background", "med_bg"]))
    pd.DataFrame(tsum).to_csv(O("gul_correlation_group_tests_by_tissue.csv"), index=False)
    print("\nCross-tissue rho:\n", piv.to_string())
    return results, piv

DEFAULT_CONFIG = {
    "anchor": "CG14989",
    "mir7_targets_tsv": "mir7_targetscanfly_conserved_targets.tsv",
    # symbols are canonicalised at load (genes.py), so one list serves all three
    "tissues": [
        {"tag": "OL",  "matrix_csv": "gene_by_cluster_meanexpr.csv",
         "shown": list(GN.SHOWN), "targets": list(GN.SHOWN_TARGETS)},
        {"tag": "VNC", "matrix_csv": "gene_by_cluster_meanexpr_VNC.csv",
         "shown": list(GN.SHOWN), "targets": list(GN.SHOWN_TARGETS)},
        {"tag": "CB",  "matrix_csv": "gene_by_cluster_meanexpr_CB.csv",
         "shown": list(GN.SHOWN), "targets": list(GN.SHOWN_TARGETS)},
    ],
}

if __name__ == "__main__":
    ap = argparse.ArgumentParser(description=__doc__, formatter_class=argparse.RawDescriptionHelpFormatter)
    ap.add_argument("--config", help="JSON config path; falls back to built-in DEFAULT_CONFIG")
    ap.add_argument("--outdir", default=None,
                    help="write everything here; default routes to results/{tables,figures}")
    ap.add_argument("--keep-glia", action="store_true",
                    help="do NOT drop glial clusters (default: excluded)")
    args = ap.parse_args()
    cfg = json.load(open(args.config)) if args.config else DEFAULT_CONFIG
    run(cfg, args.outdir, keep_glia=args.keep_glia)
```
