## supplemental tables and scripts for "Long non-coding RNA-3′UTR interactions coordinate GABA signaling in the brain": mirna_competition.py

```
#!/usr/bin/env python3
"""Correlation of conserved-8mer miR-7 targets tracks the number of competing
conserved miRNA families in their 3'UTR, not 3'UTR length per se.

The hypothesis: a short 3'UTR carrying a miR-7 site has few other regulatory
inputs, so miR-7 is a large share of its post-transcriptional control and the gene
responds to changes in miR-7 availability; a long UTR bearing the same 8mer is
subject to many other miRNAs, diluting any single input.

3'UTR length and competing-family count are collinear (rho ~ +0.90), separated
two ways: (1) partial correlations -- family count survives controlling for length,
length does not survive controlling for families; (2) within-length bands -- among
genes of similar UTR length, family count still predicts the correlation. The
response is computed by gul_duplex_model.response(), the same function the two-factor
figure uses, so the S13 panels cannot drift apart.

Flags:
  --anchor {Gad1,Gul}  gene the targets are correlated against (default Gad1).
                       Gad1 shares no variable with the sequence screen; anchoring
                       on Gul makes the figure argue partly from its own input.
  --keep-glia          retain glial clusters (default: excluded).

Inputs:
  targetscan/dmel_conserved_sites.tsv   TargetScanFly 7.2 Conserved_Family_Info,
                                        filtered to Species ID 7227
  gul_correlation_by_sitetype_pergene.csv
  3'UTR lengths derived from the FlyBase 3'UTR fasta

Usage:
    python3 mirna_competition.py
"""
import os, re, argparse
from collections import defaultdict
import numpy as np, pandas as pd
import paths as P
import genes as GN
import matplotlib
matplotlib.use("Agg")
import matplotlib.pyplot as plt
from matplotlib.lines import Line2D
from scipy.stats import spearmanr

TOP = [g for g in GN.TARGET_POOL if g != "VGAT"]
UTR_FASTA = P.find("dmel-all-three_prime_UTR-r6.58.fasta")
COL_TOP, COL_OTH, COL_TREND = "#C44E52", "#5B7C99", "#DD8452"
# symbols are canonicalised at load (genes.py), so the TargetScan tables and the
# three matrices agree on one name per gene; there is no SYN dict any more.
# target-site motifs; the base 5' of an 8mer / 7mer-m8 site faces miR-7 position 9,
# where a T can pair with the terminal A of Gul's TGGAAGACA repeats.
MOTIF = {"8mer": "GTCTTCCA", "7mer-m8": "GTCTTCC", "7mer-A1": "TCTTCCA"}


def site_positions(s, kind):
    if kind == "8mer":
        return [m.start() for m in re.finditer("GTCTTCCA", s)]
    if kind == "7mer-m8":
        return [m.start() for m in re.finditer("GTCTTCC", s)
                if s[m.start()+7:m.start()+8] != "A"]
    return [m.start() for m in re.finditer("TCTTCCA", s)
            if m.start() == 0 or s[m.start()-1] != "G"]


def sp(a, b): return spearmanr(a, b)[0]
def partial(x, y, z):
    rxy, rxz, ryz = sp(x, y), sp(x, z), sp(y, z)
    return (rxy - rxz*ryz) / np.sqrt(max((1-rxz**2)*(1-ryz**2), 1e-12))


def utr_lengths(sym2fb):
    """longest annotated 3'UTR per gene symbol, from the FlyBase fasta"""
    seqs = defaultdict(int); name = None; n = 0
    for line in open(UTR_FASTA):
        if line.startswith(">"):
            if name: seqs[name] = max(seqs[name], n)
            m = re.search(r"parent=(FBgn\d+)", line)
            name = m.group(1) if m else None; n = 0
        elif name: n += len(line.strip())
    if name: seqs[name] = max(seqs[name], n)
    return {s: seqs[fb] for s, fb in sym2fb.items() if fb in seqs and seqs[fb] > 0}


def build(datadir, anchor="Gul", exclude_glia=False):
    S = pd.read_csv(P.find("dmel_conserved_sites.tsv", datadir), sep="\t")
    S.columns = [c.strip() for c in S.columns]
    # n_fam counts the families COMPETING with miR-7, so miR-7 itself is excluded
    # (otherwise every count is +1, e.g. stol 31 rather than 30, sgll 1 rather than 0).
    # Because every gene in the 8mer analysis set carries a miR-7 site, that shift is
    # rank statistics -- rho, the partials, the two-factor model -- are unchanged.
    g = (S.groupby(["Gene ID", "Transcript ID"])
           .agg(n_sites=("miR Family", "size"))
           .join(S[S["miR Family"] != GN.MIR7_FAMILY]
                 .groupby(["Gene ID", "Transcript ID"])["miR Family"].nunique()
                 .rename("n_fam"))
           .fillna({"n_fam": 0}).astype({"n_fam": int})
           .reset_index().sort_values("n_sites", ascending=False)
           .drop_duplicates("Gene ID").set_index("Gene ID"))
    mir7 = S[S["miR Family"] == "GGAAGAC"].groupby("Gene ID").size()

    tgt = pd.read_csv(P.find("mir7_targetscanfly_conserved_targets.tsv", datadir), sep="\t")
    sym2fb = {GN.canon(s): fb for s, fb in zip(tgt.GeneSymbol.astype(str),
                                               tgt.GeneID.astype(str))}
    UL = utr_lengths(sym2fb)

    seqs = utr_seqs()
    # the UNFILTERED table: the >=25% retention filter is applied to the site-type
    # figure and group tests, but applying it here would cut the 8mer analysis set
    # from 33 genes to 27 and gut the two-factor model (see sitetype_stratification)
    D = pd.read_csv(P.find("gul_correlation_by_sitetype_pergene_unfiltered.csv", datadir))
    D["gene"] = D.gene.fillna("_").astype(str).map(GN.canon)

    # The cached table above is Gul-anchored with glia included. Under any other
    # specification the pivot supplies only the SITE-CLASS MEMBERSHIP (which genes
    # carry an 8mer, a 7mer-m8, a 7mer-A1) and the response is recomputed, so this
    # figure and the two-factor figure share one definition of rho.
    RR = None
    if anchor != "Gul" or exclude_glia:
        import gul_duplex_model as DM
        RR, _ = DM.response(anchor, exclude_glia, datadir)

    out = {}
    for kind in ["8mer", "7mer-m8", "7mer-A1"]:
        p = D[D.stratum == kind].pivot_table(index="gene", columns="tissue", values="rho")
        p = p[["OL", "VNC", "CB"]].dropna()
        if RR is not None:
            p = RR.reindex(p.index)[["OL", "VNC", "CB"]].dropna()
        rows = []
        for sym, r in p.iterrows():
            fb = sym2fb.get(sym)
            if fb not in g.index or sym not in UL: continue
            s = seqs.get(fb, "")
            pos = site_positions(s, kind)
            if not pos: continue
            flank = [s[i-1] for i in pos if i > 0]
            rows.append(dict(gene=sym, rho_OL=r.OL, rho_VNC=r.VNC, rho_CB=r.CB,
                             rho_mean=r.mean(), UTR3_len=UL[sym],
                             n_sites=int(g.loc[fb, "n_sites"]),
                             n_families=int(g.loc[fb, "n_fam"]),
                             mir7_sites=int(mir7.get(fb, 0)),
                             n_class_sites=len(pos),
                             flank5="".join(sorted(set(flank))),
                             hasT=("T" in flank)))
        F = pd.DataFrame(rows).set_index("gene")
        F["mir7_share"] = F.mir7_sites / F.n_sites
        out[kind] = F.sort_values("rho_mean", ascending=False)
    return out


def utr_seqs():
    seqs = defaultdict(str); name = None; buf = []
    for line in open(UTR_FASTA):
        if line.startswith(">"):
            if name:
                q = "".join(buf).upper()
                if len(q) > len(seqs[name]): seqs[name] = q
            m = re.search(r"parent=(FBgn\d+)", line)
            name = m.group(1) if m else None; buf = []
        elif name: buf.append(line.strip())
    if name:
        q = "".join(buf).upper()
        if len(q) > len(seqs[name]): seqs[name] = q
    return seqs


def sensitivity(F, cutoffs=range(4, 21)):
    """Is the T-adjacency effect dependent on where 'low competition' is drawn?"""
    from scipy.stats import mannwhitneyu
    rows = []
    for c in cutoffs:
        lo = F[F.n_families <= c]
        a = lo[lo.hasT].rho_mean; b = lo[~lo.hasT].rho_mean
        p = (mannwhitneyu(a, b, alternative="greater").pvalue
             if len(a) > 1 and len(b) > 1 else np.nan)
        rows.append(dict(cutoff=c, n_low=len(lo), n_T=len(a), n_noT=len(b),
                         med_T=(a.median() if len(a) else np.nan),
                         med_noT=(b.median() if len(b) else np.nan),
                         diff=(a.median()-b.median()) if len(a) and len(b) else np.nan,
                         p=p))
    return pd.DataFrame(rows)


def make_fig(F, path, anchor="Gul"):
    plt.rcParams.update({
        "figure.dpi": 110, "savefig.dpi": 200,
        "axes.spines.top": False, "axes.spines.right": False,
        "svg.fonttype": "none", "font.family": "sans-serif", "font.sans-serif": ["Arial"],
    })
    F = F.copy(); F["band"] = pd.qcut(F.UTR3_len, 3, labels=["short", "mid", "long"])
    short = F[F.band == "short"]
    fig, axes = plt.subplots(1, 3, figsize=(11.6, 4.3))

    def scat(ax, d, xcol, label_all=False):
        for gene, r in d.iterrows():
            top = gene in TOP
            ax.scatter([r[xcol]], [r.rho_mean], s=(70 if top else 34),
                       facecolor=(COL_TOP if top else COL_OTH), edgecolor="black",
                       linewidth=(1.1 if top else 0.5), zorder=(6 if top else 4))
            if top or label_all:
                ax.annotate(f"$\\it{{{gene}}}$", (r[xcol], r.rho_mean),
                            textcoords="offset points", xytext=(6, 4), fontsize=7,
                            color=(COL_TOP if top else "0.3"), zorder=8)
        ax.axhline(0, color="0.8", lw=0.9, ls="--", zorder=1)
        ax.set_ylabel(f"mean Spearman ρ with {anchor}", fontsize=9)
        ax.tick_params(labelsize=8); ax.set_ylim(-0.45, 0.82)

    # A -- UTR length
    ax = axes[0]; scat(ax, F, "UTR3_len")
    ax.set_xscale("log"); ax.set_xlabel("3′UTR length (nt)", fontsize=9.5)
    r, p = spearmanr(F.UTR3_len, F.rho_mean)
    ax.set_title("A  UTR length", fontsize=10, loc="left")
    ax.text(0.03, 0.04, f"ρ={r:+.3f} p={p:.3f}\npartial | families = "
                        f"{partial(F.rho_mean, F.UTR3_len, F.n_families):+.3f}",
            transform=ax.transAxes, fontsize=7.5, va="bottom", family="monospace")

    # B -- competing families
    # Linear x, not log: excluding miR-7 from the count puts sgll at zero, and log10(0)
    # is undefined. The counts span 0-31, so a linear axis loses nothing, and the
    # statistic quoted on the panel is rank-based and indifferent to the scale either way.
    ax = axes[1]; scat(ax, F, "n_families")
    ax.set_xlabel("competing conserved miRNA families in 3′UTR", fontsize=9.5)
    r, p = spearmanr(F.n_families, F.rho_mean)
    z = np.polyfit(F.n_families, F.rho_mean, 1)
    xs = np.linspace(F.n_families.min(), F.n_families.max(), 50)
    ax.plot(xs, np.polyval(z, xs), color=COL_TREND, lw=2, zorder=3)
    ax.set_title("B  competing miRNA families", fontsize=10, loc="left")
    ax.text(0.03, 0.04, f"ρ={r:+.3f} p={p:.3f}\npartial | UTR len = "
                        f"{partial(F.rho_mean, F.n_families, F.UTR3_len):+.3f}",
            transform=ax.transAxes, fontsize=7.5, va="bottom", family="monospace")

    # C -- within the short-UTR band
    ax = axes[2]; scat(ax, short, "n_families", label_all=True)
    ax.set_xlabel("competing families  (short-UTR genes only)", fontsize=9.5)
    r, p = spearmanr(short.n_families, short.rho_mean)
    ax.set_title(f"C  UTR-length-matched ({short.UTR3_len.min():.0f}–"
                 f"{short.UTR3_len.max():.0f} nt)", fontsize=10, loc="left")
    ax.text(0.03, 0.04, f"ρ={r:+.3f} p={p:.3f}  n={len(short)}",
            transform=ax.transAxes, fontsize=7.5, va="bottom", family="monospace")

    h = [Line2D([], [], marker="o", ls="none", mfc=COL_TOP, mec="black", mew=1.1, ms=8,
                label=", ".join(TOP)),
         Line2D([], [], marker="o", ls="none", mfc=COL_OTH, mec="black", mew=0.5, ms=7,
                label="other conserved-8mer miR-7 targets"),
         Line2D([], [], color=COL_TREND, lw=2, label="fit")]
    fig.legend(handles=h, frameon=False, ncol=3, loc="lower center",
               bbox_to_anchor=(0.5, -0.02), fontsize=8.5)
    fig.suptitle(f"{anchor} correlation tracks the number of competing miRNA inputs, "
                 "not 3′UTR length", fontsize=11)
    fig.tight_layout(rect=[0, 0.08, 1, 0.93], w_pad=1.3)
    fig.savefig(path, bbox_inches="tight")
    fig.savefig(path.replace(".png", ".svg"), bbox_inches="tight")
    plt.close(fig)


if __name__ == "__main__":
    from scipy.stats import mannwhitneyu
    ap = argparse.ArgumentParser(description=__doc__,
                                 formatter_class=argparse.RawDescriptionHelpFormatter)
    ap.add_argument("--outdir", default=None,
                    help="write everything here; default routes to results/{tables,figures}")
    ap.add_argument("--datadir", default=None, help="extra directory to search for inputs")
    ap.add_argument("--anchor", choices=["Gad1", "Gul"], default="Gad1",
                    help="correlate targets against this gene (default Gad1)"
                         "the manuscript uses Gad1)")
    ap.add_argument("--keep-glia", action="store_true",
                    help="retain glial clusters; the default drops repo-positive and named G- clusters, which is the manuscript specification")
    a = ap.parse_args()
    a.exclude_glia = not a.keep_glia
    O = lambda n: P.out(n, a.outdir)
    # same tagging rule as gul_duplex_model.py, so the default run still writes the
    # untagged gul_8mer_mirna_competition.csv that gul_duplex_model.build() reads
    tag = "" if (a.anchor == "Gad1" and a.exclude_glia) else \
          f"_{a.anchor.lower()}" + ("" if a.exclude_glia else "_glia")
    ALL = build(a.datadir, a.anchor, a.exclude_glia)
    F = ALL["8mer"]
    F.to_csv(O(f"gul_8mer_mirna_competition{tag}.csv"))
    make_fig(F, O(f"fig_mirna_competition{tag}.png"), a.anchor)

    print(f"n = {len(F)} conserved-8mer targets  "
          f"(ScsbetaG present: {'ScsbetaG' in F.index})\n")
    print(f"  rho ~ families            {sp(F.rho_mean, F.n_families):+.3f}")
    print(f"  rho ~ UTR length          {sp(F.rho_mean, F.UTR3_len):+.3f}")
    print(f"  families | UTR length     {partial(F.rho_mean, F.n_families, F.UTR3_len):+.3f}")
    print(f"  UTR length | families     {partial(F.rho_mean, F.UTR3_len, F.n_families):+.3f}")
    for lbl, d in [("excl Ssadh+CG32225", F.drop(["Ssadh", "CG32225"], errors="ignore")),
                   ("excl all 4 top", F.drop(TOP, errors="ignore"))]:
        r, p = spearmanr(d.n_families, d.rho_mean)
        print(f"  {lbl:<22} rho={r:+.3f} p={p:.3f} n={len(d)}")

    # ---- site-class summary ----
    rows = []
    for k, T in ALL.items():
        cr, cp = spearmanr(T.n_families, T.rho_mean)
        lo = T[T.n_families <= 10]; x = lo[lo.hasT].rho_mean; y = lo[~lo.hasT].rho_mean
        pv = (mannwhitneyu(x, y, alternative="greater").pvalue
              if len(x) > 1 and len(y) > 1 else np.nan)
        rows.append(dict(site_class=k, n=len(T), med_UTR=T.UTR3_len.median(),
                         med_families=T.n_families.median(), pct_T=100*T.hasT.mean(),
                         med_rho=T.rho_mean.median(), comp_rho=cr, comp_p=cp,
                         n_low=len(lo), lowT_med=(x.median() if len(x) else np.nan),
                         lowNoT_med=(y.median() if len(y) else np.nan),
                         n_T=len(x), n_noT=len(y), T_p=pv))
    SUM = pd.DataFrame(rows)
    SUM.to_csv(O(f"gul_sitetype_summary{tag}.csv"), index=False)
    print("\n=== site-class summary ===")
    print(SUM.to_string(index=False, float_format=lambda v: f"{v:.3f}"))

    # ---- sensitivity of the low-competition cutoff ----
    SEN = sensitivity(F)
    SEN.to_csv(O(f"gul_8mer_cutoff_sensitivity{tag}.csv"), index=False)
    print("\n=== cutoff sensitivity (8mer, T-adjacency within low competition) ===")
    print(f"{'cutoff':>7}{'n_low':>7}{'n_T':>5}{'n_noT':>7}{'med T':>9}{'med noT':>9}{'diff':>8}{'p':>8}")
    for _, r in SEN.iterrows():
        f = lambda v: "  --" if pd.isna(v) else f"{v:+.3f}"
        pp = "  --" if pd.isna(r.p) else f"{r.p:.3f}"
        star = " *" if (not pd.isna(r.p) and r.p < 0.05) else ""
        print(f"{int(r.cutoff):>7}{int(r.n_low):>7}{int(r.n_T):>5}{int(r.n_noT):>7}"
              f"{f(r.med_T):>9}{f(r.med_noT):>9}{f(r['diff']):>8}{pp:>8}{star}")
```
