## supplemental tables and scripts for "Long non-coding RNA-3′UTR interactions coordinate GABA signaling in the brain": paths.py

```
#!/usr/bin/env python3
"""Project paths.

Scripts refer to inputs and outputs by *filename*, not by directory, so the
folder layout can change without touching analysis code. `find()` searches the
data and results directories; `out()` routes a filename to results/tables or
results/figures by extension.

Layout (relative to the project root, i.e. this file's parent's parent):

    data/expression/   cluster-mean matrices
    data/targetscan/   TargetScanFly 7.2 target lists and conserved-site tables
    data/genome/       genome annotation (GTF, 3'UTR FASTA)
    results/tables/    every .csv an analysis writes
    results/figures/   every .png / .svg / .pdf an analysis writes

Derived tables are inputs to later scripts (e.g. mirna_competition.py reads
gul_correlation_by_sitetype_pergene.csv), so results/tables is on the search
path alongside data/.
"""
import os

ROOT    = os.path.dirname(os.path.dirname(os.path.abspath(__file__)))
DATA    = os.path.join(ROOT, "data")
EXPR    = os.path.join(DATA, "expression")
TSCAN   = os.path.join(DATA, "targetscan")
GENOME  = os.path.join(DATA, "genome")
RESULTS = os.path.join(ROOT, "results")
TABLES  = os.path.join(RESULTS, "tables")
FIGURES = os.path.join(RESULTS, "figures")

# order matters only for name collisions, of which there are currently none
SEARCH = [EXPR, TSCAN, GENOME, TABLES, FIGURES, DATA, ROOT]

FIG_EXT = (".png", ".svg", ".pdf")


def find(name, datadir=None):
    """Absolute path to an input file, located by name.

    `name` may include a subdirectory (``targetscan/dmel_conserved_sites.tsv``);
    the leading directories are tried first, then the basename alone. Passing
    `datadir` prepends a directory to the search, which is how the --datadir
    flag keeps working for out-of-tree copies of the data.
    """
    if os.path.isabs(name) and os.path.exists(name):
        return name
    dirs = ([datadir] if datadir else []) + SEARCH
    for d in dirs:
        for cand in (os.path.join(d, name), os.path.join(d, os.path.basename(name))):
            if os.path.exists(cand):
                return cand
    raise FileNotFoundError(
        f"{name!r} not found in {[os.path.relpath(d, ROOT) for d in dirs]}. "
        f"See {os.path.relpath(os.path.abspath(__file__), ROOT)} for the expected layout.")


def out(name, outdir=None):
    """Absolute path to write an output file to.

    With no `outdir`, .csv goes to results/tables and .png/.svg/.pdf to
    results/figures. An explicit `outdir` (the --outdir flag) overrides that and
    everything lands in one place, which is what the reproducibility runs want.
    """
    d = outdir or (FIGURES if name.lower().endswith(FIG_EXT) else TABLES)
    os.makedirs(d, exist_ok=True)
    return os.path.join(d, os.path.basename(name))
```
