## supplemental tables and scripts for "Long non-coding RNA-3′UTR interactions coordinate GABA signaling in the brain": rbp_gaba_screen.py

```
#!/usr/bin/env python3
"""Is any RNA-binding protein GABAergic-restricted, or specifically absent from
GABAergic neurons, anywhere in the adult CNS?

The negative result licenses searching for an RNA rather than a protein acting at
the VGAT miR-7 sites, so it is a stated threshold applied to a stated gene list
across all three CNS regions. Two readouts, either of which a reviewer could reject
alone:

  1. DETECTION (switch-like). A transcript is "GABAergic-restricted" if detected in
     >= 90% of GABAergic clusters and <= 20% of the others, "GABAergic-excluded" if
     the reverse. Gul and Gad1 are the only two transcripts of ~7,000-8,000 meeting
     the first criterion in any region; this asks whether any RBP joins them.
  2. LEVEL (fold change). Ranks every RBP by log2 mean(GABAergic) / mean(other) and
     reports where the most extreme sit relative to Gad1 and Gul.

GABAergic clusters are the Otsu cut on log Gad1 (the paper-wide definition); glia
excluded. The RBP list is an input, not hardcoded: pass --rbp-list FILE with one
gene per line as FlyBase symbols (any release; canonicalised) or FBgn IDs, blank
lines and lines starting with # ignored. A two-column TSV with a header is also
accepted (first column used).

Usage:
    python3 rbp_gaba_screen.py --rbp-list rbps.txt
    python3 rbp_gaba_screen.py --rbp-list rbps.txt --tissues OL
"""
import argparse, os
import numpy as np, pandas as pd
import matplotlib
matplotlib.use("Agg")
import matplotlib.pyplot as plt
from matplotlib.lines import Line2D

import paths as P
import genes as GN
import gul_mir7_correlation as G
from fig_switchlike_gaba import (gad1_threshold, GABA_CUT, OTHER_CUT,
                                 MIN_MEAN, MIN_P90, TISSUES)

PSEUDO = 0.01
#: Drawn for scale on both readouts -- the two transcripts that do occupy the corner.
#: Gul and Gad1 land within a few percent of each other on both axes (they are the same
#: expression pattern, which is the point), so each carries its own label offset and marker
#: size or they render as one dot with two labels stacked on it.
#:              label     colour     marker  (dx, dy) detection   line-label height
REFERENCE = {"CG14989": ("Gul",  "#C44E52", 116, (-12,  20), 0.99),
             "Gad1":    ("Gad1", "#333333",  46, (-12, -22), 0.84)}
#: display name -> colour, for the rank panel's tick labels
REFERENCE_COL = {v[0]: v[1] for v in REFERENCE.values()}

# NARRATIVE CONSTRAINT. This figure is Fig. S2, which appears *before* Gul is introduced --
# the point of the section is that no protein candidate exists, which is what motivates the
# BLAST search that finds Gul. So Gul is NOT labelled by default: it stays in the grey cloud
# of all transcripts like any other unnamed gene, and Gad1 alone is drawn, as the positive
# control proving the corner is reachable. Nothing is hidden by this -- Gul is plotted, just
# not named yet -- and the later "Gul is restricted to the GABAergic population" panel is
# this same plane with the corner finally labelled, which makes a clean callback.
# --reveal-anchor turns the label on for internal checking.
#
# Stronger still, and the reason BACKGROUND_DEFAULT is "none": the all-transcript cloud is
# not plotted either. The claim here is about RBPs, so RBPs and Gad1 are all the panel needs
# -- Gad1 alone proves the corner is reachable. Leaving the cloud out means Gul is not on
# this figure at all, rather than present-but-unlabelled, so there is no grey dot for a
# reviewer to find when the later all-transcript panel (fig_switchlike_gaba) reveals that
# Gul and Gad1 are the only two transcripts in that corner. That reveal is then a real
# result rather than a callback the reader could have pre-empted.
REVEAL_DEFAULT = False
BACKGROUND_DEFAULT = "none"          # "all" restores the full-transcriptome cloud

#: How row 2 draws the level readout.
#:
#: "rank" (default) is a ranked bar of the most GABAergic-enriched RBPs against Gad1.
#: "hist" is the original density histogram, kept because it shows both tails at once.
#:
#: A bar per named gene is preferred over the density histogram: it is one number per
#: RBP, and the labelled genes sit at the top of the distribution where a density curve
#: is ~0.01 tall, so a bar gives every label a mark to point at, and the comparison in
#: the text (best RBP 2.7x against Gad1's 17-37x)
#: becomes a length you can read off rather than a gap between a curve and a line.
LEVEL_STYLE_DEFAULT = "rank"
TOPN_LEVEL = 12

#: Set by --panel / --figsize. PANEL sends the suptitle, the footnote block and the legend
#: to the caption, prints the two shared axis labels once instead of three times, and cuts
#: each title to the region name -- the cluster counts, the Gad1 cut and the glial
#: exclusion move to the caption with everything else.
#:
#: It is what makes the figure fit a page. The full form is 12.19 x 7.48 in, nearly twice
#: A4's usable width, so it could only ever have gone on a page shrunk to ~56%, which
#: renders its 9 pt type at 5 pt. Whatever PANEL removes becomes a caption obligation:
#: record it in FIGURES.md at the same time or it is lost.
PANEL = False
FIGSIZE = None


def norm(sym):
    """Last-resort matching key: case- and punctuation-insensitive.

    Used only after identity matching has failed, for cases like bru-3 vs bru3 where the
    two files differ in punctuation alone and no shared identifier is available.
    """
    return str(sym).lower().replace("-", "").replace("_", "").replace(".", "")


def load_rbp_list(path, datadir=None):
    """RBP table: DataFrame indexed by FBgn with `symbol` and `CG` columns.

    Accepts the FBgn/symbol/CG TSV that build_rbp_list.py writes, or a bare one-per-line
    list of symbols or FBgns. Identity, not string similarity, is the primary key --
    symbols drift between FlyBase releases and the three matrices were built against
    different ones, whereas FBgn is stable and the CG annotation ID nearly so.
    """
    fp = path if os.path.exists(path) else P.find(path, datadir)
    rows = []
    with open(fp) as fh:
        for line in fh:
            s = line.strip()
            if not s or s.startswith("#"):
                continue
            f = [x.strip().strip('"') for x in s.split("\t")]
            if f[0] in ("FBgn", "gene", "symbol", "gene_symbol", "gene_id"):
                continue
            if f[0].startswith("FBgn"):
                rows.append((f[0], f[1] if len(f) > 1 else f[0], f[2] if len(f) > 2 else ""))
            else:                       # bare symbol list: no identifier available
                rows.append(("", f[0], f[0] if f[0].startswith("CG") else ""))
    R = pd.DataFrame(rows, columns=["FBgn", "symbol", "CG"]).drop_duplicates()
    n_id = int((R.FBgn != "").sum())
    print(f"RBP list: {len(R):,} genes ({n_id:,} with an FBgn, "
          f"{int((R.CG != '').sum()):,} with a CG)")
    return R


def resolve_index(index, R, datadir=None):
    """{matrix gene symbol: FBgn} for every matrix gene that can be identified.

    Resolution order, most to least authoritative:
      1. current FlyBase r6.68 symbol   (covers every gene the matrix names currently)
      2. CG annotation ID               (covers matrices that predate a gene being named)
      3. RBP-table symbol               (covers a gene the r6.68 table happens to miss)
    """
    cur = pd.read_csv(P.find("fbgn_symbol_r6.68.tsv", datadir), sep="\t")
    sym2fb = dict(zip(cur.symbol.astype(str), cur.FBgn.astype(str)))
    cg2fb = {cg: fb for fb, cg in zip(R.FBgn, R.CG) if cg and fb}
    rsym2fb = {sy: fb for fb, sy in zip(R.FBgn, R.symbol) if fb}
    out = {}
    for g in index:
        g = str(g)
        fb = sym2fb.get(g) or cg2fb.get(g) or rsym2fb.get(g)
        if fb:
            out[g] = fb
    return out


def match_rbps(index, R, datadir=None):
    """(matched genes, {gene: display symbol}, n_unresolved, ambiguous).

    A matrix gene is an RBP when its FBgn is in the list. This is what separates DCP1
    (FBgn0034921, decapping protein 1, a real RBP) from Dcp-1 (FBgn0010501, Death
    caspase-1, not one): they differ only in punctuation as symbols but are unrelated
    genes, and a symbol-normalising join scores the caspase as an RBP.

    Genes the identity path cannot resolve fall back to a normalised-symbol match against
    the list, accepted only when unambiguous.
    """
    idx = [str(g) for g in index]
    gene2fb = resolve_index(idx, R, datadir)
    want = set(R.FBgn) - {""}
    fb2sym = {fb: sy for fb, sy in zip(R.FBgn, R.symbol) if fb}

    matched, label = set(), {}
    for g, fb in gene2fb.items():
        if fb in want:
            matched.add(g)
            label[g] = fb2sym.get(fb, g)

    # normalised fallback, only for list entries no FBgn matched and only when unique
    hit_fb = {gene2fb[g] for g in matched}
    unclaimed = R[(~R.FBgn.isin(hit_fb)) | (R.FBgn == "")]
    by_key = {}
    for g in idx:
        if g not in matched:
            by_key.setdefault(norm(g), []).append(g)
    ambiguous = {}
    for _, r in unclaimed.iterrows():
        cand = by_key.get(norm(r.symbol), [])
        if len(cand) == 1:
            matched.add(cand[0]); label[cand[0]] = r.symbol
        elif len(cand) > 1:
            ambiguous[r.symbol] = cand
    if ambiguous:
        print("    ambiguous normalised keys, not matched: "
              + "; ".join(f"{k} -> {v}" for k, v in list(ambiguous.items())[:5]))
    return matched, label, len(idx) - len(gene2fb), ambiguous


def screen_one(csv, tag, R, datadir=None):
    """Detection and fold-change readouts for one region."""
    M = GN.canon_index(pd.read_csv(P.find(csv, datadir), index_col=0))
    # repo >= 0.17 plus OL's named G- clusters -- verified to give the identical cluster
    # set to fig_switchlike_gaba, gaba_derepression_class and the site-type panels, so
    # every figure in the paper is computed over the same 231 / 115 / 346 neuronal clusters
    N = G.drop_glia(M, tag)
    n_glia = M.shape[1] - N.shape[1]
    gad1 = N.loc[GN.GAD1]
    cut = gad1_threshold(gad1.values)
    gaba, other = gad1[gad1 >= cut].index, gad1[gad1 < cut].index

    p90 = N.quantile(.90, axis=1)
    thr = 0.1 * p90
    expressed = (N.mean(axis=1) > MIN_MEAN) & (p90 > MIN_P90)
    mu_hi, mu_lo = N[gaba].mean(axis=1), N[other].mean(axis=1)

    D = pd.DataFrame(dict(
        tissue=tag,
        det_gaba=N[gaba].gt(thr, axis=0).mean(axis=1),
        det_other=N[other].gt(thr, axis=0).mean(axis=1),
        lfc=np.log2((mu_hi + PSEUDO) / (mu_lo + PSEUDO)),
        mean_expr=N.mean(axis=1)))[expressed]

    matched, label, n_unres, _ = match_rbps(N.index, R, datadir)
    print(f"  [{tag}] {len(matched):,} of {len(R):,} listed RBPs present "
          f"({n_unres:,} of {N.shape[0]:,} matrix genes carry no resolvable identifier)")
    D = D.reset_index(names="gene")
    D["is_rbp"] = D.gene.isin(matched)
    D["label"] = [label.get(g, g) for g in D.gene]
    D["restricted"] = (D.det_gaba >= GABA_CUT) & (D.det_other <= OTHER_CUT)
    D["excluded"] = (D.det_other >= GABA_CUT) & (D.det_gaba <= OTHER_CUT)
    return D, cut, len(gaba), len(other), n_glia


def _level_rank(ax, d, rb, refs, c):
    """Row 2, rank style: the top RBPs by fold change, with Gad1 on the same axis.

    x is log2 fold change but ticked in fold change, because that is the unit the
    manuscript quotes. The bar for every named gene is visible by construction, which
    is the whole point of the change.
    """
    top = rb.nlargest(TOPN_LEVEL, "lfc").sort_values("lfc")
    y = np.arange(len(top), dtype=float)
    ax.barh(y, top.lfc.values, height=0.72, color="#4C72B0", zorder=3)
    names = list(top.label)

    gap = 1.2                     # blank row separating the reference genes from the RBPs
    for g, (name, col, _, _, _) in refs.items():
        if g not in set(d.gene): continue
        v = float(d[d.gene == g].iloc[0].lfc)
        yy = len(top) - 1 + gap
        ax.barh([yy], [v], height=0.72, color=col, zorder=3)
        ax.text(v - 0.12, yy, f"{2 ** v:.0f}×", va="center", ha="right",
                fontsize=8, color="white", fontweight="bold", zorder=5)
        y = np.append(y, yy); names.append(name)

    ax.set_yticks(y)
    ax.set_yticklabels(names, fontsize=7.6)
    for t, nm in zip(ax.get_yticklabels(), names):
        t.set_fontstyle("italic")
        t.set_color(REFERENCE_COL.get(nm, "0.25"))
    ax.tick_params(axis="y", length=0)
    ax.axvline(0, color="0.55", lw=0.8, zorder=2)

    # the best RBP, called out because it is the number the text quotes
    best = top.iloc[-1]
    ax.text(best.lfc + 0.10, len(top) - 1, f"{2 ** best.lfc:.1f}×", va="center",
            ha="left", fontsize=8, color="#2A4A6B", fontweight="bold")

    hi = max([float(d[d.gene == g].iloc[0].lfc) for g in refs if g in set(d.gene)]
             or [top.lfc.max()])
    ax.set_xlim(-0.25, hi * 1.16)
    # the footnote below sits inside the axes, so the axes must carry room for it. At full
    # width it cleared the lowest gene name on its own; at a third of that width the line
    # spans the whole panel and strikes through it, so panel mode both breaks the note in
    # two and opens a blank band beneath the last bar for it to sit in.
    ax.set_ylim(-2.2 if PANEL else -0.8, len(top) - 1 + gap + 0.9)
    ticks = [t for t in range(0, 7) if t <= hi * 1.16]
    ax.set_xticks(ticks)
    ax.set_xticklabels([f"{2 ** t:g}×" for t in ticks], fontsize=8)
    if not PANEL or c == 1:
        ax.set_xlabel("mean(GABAergic) / mean(other)", fontsize=9)
    # the range of what is NOT drawn, so it must exclude the drawn bars: quoting
    # top.lfc.iloc[-2] gave the second-highest bar on the panel and claimed it as the
    # ceiling of the genes the panel leaves out
    rest = rb.drop(top.index)
    sep = "\n" if PANEL else " "
    ax.text(0.985, 0.02,
            f"the other {len(rest):,} RBPs{sep}"
            f"span {2 ** rest.lfc.min():.1f}× to {2 ** rest.lfc.max():.1f}×",
            transform=ax.transAxes, ha="right", va="bottom", fontsize=7.2, color="0.45",
            linespacing=1.35)


def make_fig(frames, meta, path, reveal_anchor=REVEAL_DEFAULT,
             background=BACKGROUND_DEFAULT, level_style=LEVEL_STYLE_DEFAULT):
    refs = REFERENCE if reveal_anchor else {k: v for k, v in REFERENCE.items()
                                            if k != GN.ANCHOR}
    show_bg = background == "all"
    plt.rcParams.update({
        "figure.dpi": 110, "savefig.dpi": 200,
        "axes.spines.top": False, "axes.spines.right": False,
        "svg.fonttype": "none", "font.family": "sans-serif", "font.sans-serif": ["Arial"]})
    tags = list(frames)
    fig, axes = plt.subplots(2, len(tags), squeeze=False,
                             figsize=FIGSIZE or (3.9 * len(tags) + 0.6, 7.2))

    for c, tag in enumerate(tags):
        d = frames[tag]
        cut, n_gaba, n_other, n_glia = meta[tag]
        rb = d[d.is_rbp]

        # --- row 1: the detection plane, RBPs highlighted inside the full distribution
        ax = axes[0][c]
        # The corner is drawn to the axis limits, not to a number of its own: at 1.05
        # against a limit of 1.06 it stopped a point short of the right spine, leaving a
        # hairline of white between the shaded region and the frame. Same reason ylo is
        # shared with set_ylim below rather than written twice.
        xhi, ylo = 1.06, (-0.075 if PANEL else -0.03)
        ax.add_patch(plt.Rectangle((GABA_CUT, ylo), xhi - GABA_CUT, OTHER_CUT - ylo,
                                   color="0.92", zorder=0))
        if show_bg:
            ax.scatter(d.det_gaba, d.det_other, s=4, alpha=0.08, color="#8A8A8A",
                       edgecolor="none", rasterized=True, zorder=2)
        # isolated RBPs are the informative ones here -- the sparse lower-left tail is
        # where an RBP would have to sit to be GABAergic-restricted -- and at alpha 0.75
        # they read as faint. Same fix as fig_switchlike_gaba's cloud.
        ax.scatter(rb.det_gaba, rb.det_other, s=18, alpha=0.92, color="#3E6795",
                   edgecolor="none", zorder=4)
        ax.axvline(GABA_CUT, color="0.55", lw=0.8, ls=":", zorder=3)
        ax.axhline(OTHER_CUT, color="0.55", lw=0.8, ls=":", zorder=3)
        for g, (name, col, ms, off, _) in refs.items():
            if g not in set(d.gene): continue
            r = d[d.gene == g].iloc[0]
            # Gad1 sits at (1.0, 0.0), so its label hangs below the axis and lands on the
            # x tick labels once the panel is narrow. Point it up instead, into the shaded
            # corner, which is empty by construction -- that is the panel's whole result.
            # Only when Gul is not also drawn: the two coincide, so both labels go up and
            # collide. --reveal-anchor is internal checking, so it keeps the wide form.
            if PANEL and not reveal_anchor and g == GN.GAD1:
                off = (-9, 20)
            ax.scatter([r.det_gaba], [r.det_other], s=ms, facecolor=col,
                       edgecolor="white", linewidth=1.2, zorder=6)
            # style="italic" rather than mathtext: mathtext leaves the digit of Gad1
            # upright and shatters the label into one tspan per glyph, which Illustrator
            # cannot edit. The rank panel's tick labels already use set_fontstyle, so the
            # two halves of the figure now italicise the same gene the same way.
            ax.annotate(name, (r.det_gaba, r.det_other), style="italic",
                        textcoords="offset points", xytext=off, ha="right",
                        fontsize=9, color=col, fontweight="bold",
                        arrowprops=dict(arrowstyle="-", color=col, lw=0.7,
                                        shrinkA=0, shrinkB=4))
        n_in = int(rb.restricted.sum())
        ax.text(0.03, 0.97, f"{len(rb):,} RBPs expressed\n{n_in} in the corner",
                transform=ax.transAxes, va="top", fontsize=8, color="0.3")
        # Gad1 sits at det_other = 0.0, on the axis boundary, and marker size is in POINTS:
        # the 0.03 of bottom margin is wider than its radius at full size and narrower once
        # the panel shrinks, so the marker is sliced by the spine. Pad in panel mode, and
        # draw the corner rectangle to the same limit so it still reaches the edge.
        ax.set_xlim(-0.03, xhi); ax.set_ylim(ylo, 1.12)
        # the six axes share two labels; in a composite print each once, which also stops
        # the leftmost x label overhanging the figure and being clipped
        if not PANEL or c == len(tags) // 2:
            ax.set_xlabel("fraction of GABAergic clusters detected", fontsize=9)
        if c == 0: ax.set_ylabel("fraction of other clusters detected", fontsize=9.5)
        ax.set_title(f"{G.TISSUE_NAME.get(tag, tag)} ({tag})" if PANEL else
                     (f"{G.TISSUE_NAME.get(tag, tag)} ({tag})\n"
                      f"{n_gaba} GABAergic / {n_other} other neuronal clusters\n"
                      f"({n_glia} glial clusters excluded)"), fontsize=10)

        # --- row 2: the level readout, so the claim does not rest on one threshold
        ax = axes[1][c]
        if level_style == "rank":
            _level_rank(ax, d, rb, refs, c)
            continue
        if show_bg:
            ax.hist(d[~d.is_rbp].lfc.values, bins=60, color="#D9D9D9", edgecolor="none",
                    density=True, zorder=1)
        ax.hist(rb.lfc.values, bins=40, color="#4C72B0",
                alpha=0.6 if show_bg else 0.85, edgecolor="none", density=True, zorder=2)
        top = rb.nlargest(3, "lfc")
        ymax = ax.get_ylim()[1]
        for i, (_, r) in enumerate(top.iterrows()):
            ax.annotate(f"$\\it{{{r.label}}}$", (r.lfc, ymax * (0.30 + 0.14 * i)),
                        ha="center", fontsize=7.6, color="#2A4A6B",
                        arrowprops=dict(arrowstyle="-", color="#4C72B0", lw=0.7))
        for g, (name, col, _, _, h) in refs.items():
            if g not in set(d.gene): continue
            v = d[d.gene == g].iloc[0].lfc
            ax.axvline(v, color=col, lw=1.4, zorder=5)
            ax.annotate(f"$\\it{{{name}}}$ {2**v:.0f}×", (v, ymax * h), ha="right",
                        textcoords="offset points", xytext=(-3, 0), fontsize=8.4,
                        color=col, fontweight="bold")
        ax.set_xlabel("log$_2$ mean(GABAergic) / mean(other)", fontsize=9)
        if c == 0: ax.set_ylabel("density", fontsize=9.5)

    if PANEL:
        # Panel letters. The composites draw these in svg_layout, but S2 is generated whole
        # with no artwork of RL's to place beside it, so there is no layout script to hang
        # them on. Offset in POINTS from the first axes of each row, past the widest left
        # furniture in either row: the rotated y label in row 1, "alpha-PheRS" in row 2.
        for row, letter in ((0, "A"), (1, "B")):
            axes[row][0].annotate(letter, xy=(0, 1), xycoords="axes fraction",
                                  xytext=(-52, 14), textcoords="offset points",
                                  fontsize=12, fontweight="bold", va="top")
        # suptitle, footnote and legend go to the caption, the same convention every other
        # panel script follows, and it is what recovers the height the page needs
        fig.tight_layout(w_pad=1.6, h_pad=1.4)
        path = path.replace(".png", "_panel.png")
        fig.savefig(path, bbox_inches="tight")
        fig.savefig(path.replace(".png", ".svg"), bbox_inches="tight")
        plt.close(fig)
        return

    h1 = ([Line2D([], [], marker="o", ls="none", mfc="#8A8A8A", mec="none", ms=6,
                  label="all expressed transcripts")] if show_bg else [])
    h1 += [Line2D([], [], marker="o", ls="none", mfc="#4C72B0", mec="none", ms=7,
                  label="annotated RNA-binding proteins")]
    h1 += [Line2D([], [], marker="o", ls="none", mfc=REFERENCE[g][1], mec="white",
                  ms=7, label=f"$\\it{{{REFERENCE[g][0]}}}$") for g in refs]
    fig.legend(handles=h1, frameon=False, ncol=len(h1), loc="lower center",
               bbox_to_anchor=(0.5, -0.04), fontsize=8.6)
    fig.suptitle("No RNA-binding protein is restricted to, or excluded from, "
                 "GABAergic neurons", fontsize=12)
    fig.text(0.5, 0.012,
             "Top, detection breadth: shaded corner is detection in ≥90% of GABAergic "
             "clusters and ≤20% of others. Bottom, expression level, one bar per gene.\n"
             "Glial clusters are "
             "excluded throughout ($\\it{repo}$ ≥ 0.17 plus named glial clusters) and "
             "GABAergic clusters are defined by an Otsu split of the $\\it{Gad1}$\n"
             "distribution — the same neuronal cluster set and the same split used in "
             "every other figure.", ha="center", fontsize=7.6, color="0.35")
    fig.tight_layout(rect=[0, 0.06, 1, 0.94])
    fig.savefig(path, bbox_inches="tight")
    fig.savefig(path.replace(".png", ".svg"), bbox_inches="tight")
    plt.close(fig)


def run(rbp_list, tissues=None, outdir=None, datadir=None,
        reveal_anchor=REVEAL_DEFAULT, background=BACKGROUND_DEFAULT,
        level_style=LEVEL_STYLE_DEFAULT):
    def O(n): return P.out(n, outdir)
    R = load_rbp_list(rbp_list, datadir)
    todo = [(csv, tag) for csv, tag in TISSUES if not tissues or tag in tissues]
    frames, meta, rows = {}, {}, []

    def lfc_of(D, g):
        """log2FC for one gene, NaN if the expression filter dropped it."""
        s = D[D.gene == g].lfc
        return float(s.iloc[0]) if len(s) else float("nan")

    for csv, tag in todo:
        D, cut, n_gaba, n_other, n_glia = screen_one(csv, tag, R, datadir)
        frames[tag] = D
        meta[tag] = (cut, n_gaba, n_other, n_glia)
        rb = D[D.is_rbp]
        assert len(rb), f"[{tag}] no gene from the RBP list is expressed — check the list"
        rows.append(dict(tissue=tag, gad1_cut=cut, n_gaba=n_gaba, n_other=n_other,
                         n_glia_excluded=n_glia,
                         n_expressed=len(D), n_rbp_listed=len(R), n_rbp_expressed=len(rb),
                         n_rbp_restricted=int(rb.restricted.sum()),
                         n_rbp_excluded=int(rb.excluded.sum()),
                         n_all_restricted=int(D.restricted.sum()),
                         restricted_genes=",".join(sorted(D[D.restricted].gene)),
                         max_rbp_lfc=rb.lfc.max(), max_rbp_gene=rb.loc[rb.lfc.idxmax(), "label"],
                         min_rbp_lfc=rb.lfc.min(), min_rbp_gene=rb.loc[rb.lfc.idxmin(), "label"],
                         gad1_lfc=lfc_of(D, GN.GAD1), gul_lfc=lfc_of(D, GN.ANCHOR)))
        print(f"\n[{tag}] {len(D):,} expressed transcripts, {len(rb):,} of them RBPs; "
              f"{n_gaba} GABAergic / {n_other} other clusters")
        print(f"  in the GABAergic-restricted corner: {int(D.restricted.sum())} transcripts "
              f"({', '.join(sorted(D[D.restricted].gene)) or 'none'}), "
              f"of which RBPs: {int(rb.restricted.sum())}")
        print(f"  GABAergic-excluded corner: {int(rb.excluded.sum())} RBPs")
        print(f"  most GABAergic-biased RBPs by level:")
        for _, r in rb.nlargest(5, "lfc").iterrows():
            print(f"    {r.label:<14} log2FC {r.lfc:+.2f} ({2**r.lfc:.2f}x)  "
                  f"detected {100*r.det_gaba:.0f}% GABAergic / {100*r.det_other:.0f}% other")
        g1, gu = lfc_of(D, GN.GAD1), lfc_of(D, GN.ANCHOR)
        print(f"    -- for scale: Gad1 {g1:+.2f} ({2**g1:.1f}x), Gul {gu:+.2f} ({2**gu:.1f}x)")

    A = pd.concat(frames.values(), ignore_index=True)
    S = pd.DataFrame(rows)
    A[A.is_rbp].to_csv(O("rbp_gaba_screen_pergene.csv"), index=False)
    S.to_csv(O("rbp_gaba_screen_summary.csv"), index=False)
    make_fig(frames, meta, O("fig_rbp_gaba_screen.png"), reveal_anchor, background,
             level_style)

    print("\n=== summary ===")
    print(S[["tissue", "n_expressed", "n_rbp_expressed", "n_rbp_restricted",
             "n_rbp_excluded", "max_rbp_gene", "max_rbp_lfc", "gad1_lfc"]]
          .to_string(index=False, float_format=lambda v: f"{v:.2f}"))
    hit = S.n_rbp_restricted.sum() + S.n_rbp_excluded.sum()
    print(f"\n{'NO' if hit == 0 else hit} RBP{'' if hit == 1 else 's'} meet either "
          f"criterion in any region.")

    # RBPs that come closest, pooled across regions -- the honest near-miss list
    rb = A[A.is_rbp].copy()
    rb["margin"] = rb.det_gaba - rb.det_other
    print("\nclosest approaches to the restricted corner (any region):")
    print(rb.nlargest(8, "margin")[["tissue", "label", "det_gaba", "det_other", "lfc"]]
            .to_string(index=False, float_format=lambda v: f"{v:.2f}"))
    return A, S


if __name__ == "__main__":
    ap = argparse.ArgumentParser(description=__doc__,
                                 formatter_class=argparse.RawDescriptionHelpFormatter)
    ap.add_argument("--rbp-list", required=True,
                    help="file of RBP gene symbols or FBgn IDs, one per line")
    ap.add_argument("--tissues", nargs="+", default=None, help="subset of OL VNC CB")
    ap.add_argument("--outdir", default=None)
    ap.add_argument("--datadir", default=None)
    ap.add_argument("--reveal-anchor", action="store_true",
                    help="label Gul in the figure; off by default because Fig. S2 runs "
                         "before Gul is introduced (see REVEAL_DEFAULT)")
    ap.add_argument("--background", choices=["none", "all"], default=BACKGROUND_DEFAULT,
                    help="'all' adds the full-transcriptome cloud behind the RBPs; off by "
                         "default so Gul is absent from this figure entirely")
    ap.add_argument("--level-style", choices=["rank", "hist"],
                    default=LEVEL_STYLE_DEFAULT,
                    help="row 2: 'rank' bars the top RBPs against Gad1 (default), "
                         "'hist' restores the original density histogram")
    ap.add_argument("--panel", action="store_true",
                    help="strip the suptitle, footnote and legend to the caption and "
                         "print the shared axis labels once; writes *_panel.svg")
    ap.add_argument("--figsize", nargs=2, type=float, default=None, metavar=("W", "H"),
                    help="figure size in inches; generate a panel at the size it will "
                         "be placed at, or its type is scaled down with it")
    a = ap.parse_args()
    FIGSIZE, PANEL = (tuple(a.figsize) if a.figsize else None), a.panel
    run(a.rbp_list, a.tissues, a.outdir, a.datadir, a.reveal_anchor, a.background,
        a.level_style)
```
