## supplemental tables and scripts for "Long non-coding RNA-3′UTR interactions coordinate GABA signaling in the brain": sitetype_stratification.py

```
#!/usr/bin/env python3
"""
Does Gul-correlation scale with miR-7 site strength?

Stratifies every gene's Spearman rho with Gul by its best conserved miR-7 site
type (8mer > 7mer-m8 > 7mer-A1), plus a "poorly conserved sites only" class,
against the non-target background -- independently in OL, VNC and CB.

Motivation: the pooled miR-7 target class is not elevated (Fig. S12). Two
explanations were possible: (a) the 549-gene list is diluted by weak/poorly
conserved predictions, so a real effect is washed out; or (b) there is no
site-strength effect. This script distinguishes them.

IMPORTANT -- the shown genes are EXCLUDED from the strata in the primary
analysis. VGAT (3 conserved 8mer sites), sgll and ScsbetaG are all 8mer targets and
all in the shown set; leaving them in makes the 8mer stratum test circular. Both
versions are computed so the difference is visible.

Inputs (all already in this folder):
  TargetScan7.2__miR-7-5p.predicted_targets.txt  per-gene conserved site counts
  mir7_targetscanfly_conserved_targets.tsv       the broad 549-gene list
  gene_by_cluster_meanexpr*.csv                  the cluster-mean matrices

Usage:
    python sitetype_stratification.py [--outdir .]
"""
import os, sys, argparse
import numpy as np, pandas as pd
import paths as P
import genes as GN
import matplotlib
matplotlib.use("Agg")
import matplotlib.pyplot as plt
from matplotlib.lines import Line2D
from scipy.stats import mannwhitneyu

import gul_mir7_correlation as G

PRED_TSV = "TargetScan7.2__miR-7-5p.predicted_targets.txt"
STRATA = ["background", "poorly-cons-only", "7mer-A1", "7mer-m8", "8mer"]
LABEL  = {"background": "Bg", "poorly-cons-only": "poorly\ncons.",
          "7mer-A1": "7mer\nA1", "7mer-m8": "7mer\nm8", "8mer": "8mer"}
COLOR  = {"background": "#8A8A8A", "poorly-cons-only": "#C6DBEF",
          "7mer-A1": "#9ECAE1", "7mer-m8": "#4C72B0", "8mer": "#08306B"}
LEGEND = {"background": "Non-target background",
          "poorly-cons-only": "Poorly conserved sites only",
          "7mer-A1": "conserved 7mer-A1 site", "7mer-m8": "conserved 7mer-m8 site",
          "8mer": "conserved 8mer site"}


def load_site_types(datadir=None):
    """gene symbol -> best conserved site type (8mer > 7mer-m8 > 7mer-A1).

    Source is TargetScan's per-gene `predicted_targets` export (183 genes), **not** the
    per-site `dmel_conserved_sites.tsv` (549 genes). That looks like the narrower choice
    and is the right one.

    The two files differ by PCT -- TargetScan's probability of *preferential* conservation.
    A site can appear in the conserved-site file with PCT = 0, meaning it is present in the
    alignment but conserved no better than background expectation. Of the 366 genes in the
    per-site file but not in this export, 266 have best-site PCT = 0 and 70 have no PCT at
    all; every one of the 151 genes with PCT > 0.5 is in the export. So "poorly conserved
    sites only" is a fair name for what is dropped, and the exclusion is a conservation
    criterion rather than an arbitrary subset.

    This also explains something that looks wrong on the panels: conserved 8mers outnumber
    7mer-m8s, where raw site counts run 2:1 the other way. Selection is the reason -- 53%
    of 8mer sites are preferentially conserved against 22-23% of 7mer sites -- so requiring
    conservation inverts the ratio (8mer:7mer-m8 goes 0.52 -> 1.19 at PCT > 0.5).

    Building the strata from the per-site file instead was tried and reverted the same day:
    it adds 13 genes to the two-factor model set, none with rho > +0.2, and halves its R2
    from 0.36 to 0.22. They are sites with no evidence of selection.
    """
    T = pd.read_csv(P.find(PRED_TSV, datadir), sep="\t")
    def best(r):
        if r["Conserved 8mer sites"] > 0:    return "8mer"
        if r["Conserved 7mer-m8 sites"] > 0: return "7mer-m8"
        if r["Conserved 7mer-A1 sites"] > 0: return "7mer-A1"
        return None
    # keyed on canonical symbols: CB uses newer FlyBase names than OL/VNC, and
    # 30 CB targets were falling into the background stratum without this
    site = {GN.canon(g): b for g, b in zip(T["Target gene"].astype(str),
                                           T.apply(best, axis=1))}
    # genes whose TargetScan representative transcript is not the expressed one
    site.update(GN.override_site_types(datadir))
    return site, T


#: minimum share of clusters a gene must be retained in by the dead-tail filter.
#: Below this a Spearman on a handful of points reaches +0.78 on noise — see the
#: docstring of the same constant in sitetype_stratification_gad1.py.
MIN_RETAIN = 0.25


def collect(cfg, datadir=None, keep_glia=False, min_retain=None):
    """Per-gene rho + stratum for every tissue.

    Returns every gene, with ``n_clusters`` and ``n_total`` so the caller can
    apply the retention filter. It is applied for the figure and the group tests
    but deliberately NOT for the downstream gene set: see the note in __main__.
    """
    site, _ = load_site_types(datadir)
    broad = G.load_mir7_symbols(P.find(cfg["mir7_targets_tsv"], datadir))
    frames = []
    for t in cfg["tissues"]:
        M = GN.canon_index(pd.read_csv(P.find(t["matrix_csv"], datadir), index_col=0))
        if not keep_glia:
            M = G.drop_glia(M, t["tag"])
        A = G.analyze(t["tag"], M, cfg["anchor"], t["shown"], t["targets"],
                      set(broad) | set(t.get("mir7_extra", [])))
        res = A["res"]; extra = set(t.get("mir7_extra", []))
        def stratum(g):
            s = site.get(g)
            if s: return s
            return "poorly-cons-only" if (g in broad or g in extra) else "background"
        d = pd.DataFrame(dict(
            tissue=t["tag"], gene=res.index, rho=res.rho.values,
            mean_expr=res.mean_expr.values, n_clusters=res.n_clusters.values,
            n_total=M.shape[1],
            stratum=[stratum(g) for g in res.index],
            is_shown=[g in set(t["shown"]) for g in res.index]))
        frames.append(d)
        print(f"[{t['tag']}] {len(d)} genes stratified over {M.shape[1]} clusters")
    return pd.concat(frames, ignore_index=True)


def apply_retention(D, min_retain=MIN_RETAIN):
    """Drop genes the dead-tail filter kept in < min_retain of clusters."""
    if not min_retain:
        return D
    keep = D.n_clusters >= min_retain * D.n_total
    for tag, d in D.groupby("tissue"):
        n = int((~(d.n_clusters >= min_retain * d.n_total)).sum())
        print(f"[{tag}] {n} genes retained in < {min_retain:.0%} of clusters dropped")
    return D[keep]


def group_tests(D):
    rows = []
    for tag in D.tissue.unique():
        for excl in (True, False):
            d = D[(D.tissue == tag) & (~D.is_shown if excl else slice(None))] \
                if excl else D[D.tissue == tag]
            bg = d[d.stratum == "background"].rho.values
            for s in STRATA:
                v = d[d.stratum == s].rho.values
                if s == "background":
                    rows.append(dict(tissue=tag, excl_shown=excl, stratum=s, n=len(v),
                                     med_rho=np.median(v), med_bg=np.median(bg),
                                     mean_rho=np.mean(v), mean_bg=np.mean(bg),
                                     mwu_p=np.nan, perm_p=np.nan,
                                     perm_p_mean=np.nan)); continue
                if len(v) < 3: continue
                _, pp = G.perm_test(v, bg)                      # medians
                _, pm = G.perm_test(v, bg, stat="mean")          # tail-sensitive
                rows.append(dict(tissue=tag, excl_shown=excl, stratum=s, n=len(v),
                                 med_rho=np.median(v), med_bg=np.median(bg),
                                 mean_rho=np.mean(v), mean_bg=np.mean(bg),
                                 mwu_p=mannwhitneyu(v, bg, alternative="greater").pvalue,
                                 perm_p=pp, perm_p_mean=pm))
    return pd.DataFrame(rows)


def make_fig(D, T, order, path):
    plt.rcParams.update({
        "figure.dpi": 110, "savefig.dpi": 200,
        "axes.spines.top": False, "axes.spines.right": False,
        "svg.fonttype": "none", "font.family": "sans-serif", "font.sans-serif": ["Arial"],
    })
    S = T[T.excl_shown == True]
    fig, axes = plt.subplots(1, 3, figsize=(8.6, 4.6))
    for ax, tag in zip(np.atleast_1d(axes), order):
        d = D[(D.tissue == tag) & (~D.is_shown)]
        st = S[S.tissue == tag].set_index("stratum")
        ax.axhline(np.median(d[d.stratum == "background"].rho.values),
                   color="0.45", lw=0.9, ls=":", zorder=2)
        for i, s in enumerate(STRATA):
            v = d[d.stratum == s].rho.values
            if not len(v): continue
            big = (s == "background")
            jit = (np.random.default_rng(i).random(len(v)) - 0.5) * (0.34 if big else 0.30)
            ax.scatter(np.full(len(v), i) + jit, v, s=(7 if big else 16),
                       alpha=(0.13 if big else 0.55), color=COLOR[s],
                       edgecolor="none", zorder=3, rasterized=True)
            ax.plot([i-0.30, i+0.30], [np.median(v)]*2, color="black", lw=2.0, zorder=5)
            ax.text(i, 1.30, f"n={len(v)}", ha="center", va="bottom", fontsize=7, color="0.3")
            if s != "background":
                ax.text(i, 1.13, f"p={st.loc[s, 'perm_p']:.2f}", ha="center",
                        va="bottom", fontsize=7.5)
        ax.axhline(0, color="0.85", lw=0.8, ls="--", zorder=1)
        ax.set_xticks(range(len(STRATA)))
        ax.set_xticklabels([LABEL[s] for s in STRATA], fontsize=8.5)
        ax.set_xlim(-0.6, len(STRATA) - 0.4); ax.set_ylim(-1.1, 1.5)
        ax.tick_params(axis="y", labelsize=8.5)
        ax.set_title(f"{G.TISSUE_NAME.get(tag, tag)} ({tag})", fontsize=10.5)
        ax.set_ylabel("Spearman ρ with Gul", fontsize=9.5)
    handles = [Line2D([], [], marker="o", ls="none", mfc=COLOR[s], mec="none", ms=7,
                      label=LEGEND[s]) for s in STRATA]
    handles += [Line2D([], [], color="black", lw=2.0, label="group median"),
                Line2D([], [], color="0.45", lw=0.9, ls=":", label="background median")]
    fig.legend(handles=handles, frameon=False, ncol=4, loc="lower center",
               bbox_to_anchor=(0.5, 0.005), fontsize=8.5)
    fig.suptitle("Gul correlation does not scale with miR-7 site strength "
                 "(shown genes excluded)", fontsize=11)
    fig.tight_layout(rect=[0, 0.10, 1, 0.94], w_pad=1.0)
    fig.savefig(path, bbox_inches="tight")
    fig.savefig(path.replace(".png", ".svg"), bbox_inches="tight")
    plt.close(fig)


def make_expr_fig(D, path, stratum="8mer"):
    """rho vs mean expression for one site-type stratum, background behind.

    Shows that the shown genes' high rho is not a by-product of expression level:
    they sit at ordinary abundance but are vertical outliers among site-matched peers.
    """
    from scipy.stats import spearmanr
    plt.rcParams.update({
        "figure.dpi": 110, "savefig.dpi": 200,
        "axes.spines.top": False, "axes.spines.right": False,
        "svg.fonttype": "none", "font.family": "sans-serif", "font.sans-serif": ["Arial"],
    })
    col_s = COLOR[stratum]; col_shown = "#C44E52"
    order = ["OL", "VNC", "CB"]
    fig, axes = plt.subplots(1, 3, figsize=(9.6, 3.9))
    for ax, tag in zip(np.atleast_1d(axes), order):
        d = D[D.tissue == tag]
        bg = d[d.stratum == "background"]; S = d[d.stratum == stratum]
        bulk, shown = S[~S.is_shown], S[S.is_shown]
        ax.scatter(bg.mean_expr, bg.rho, s=5, alpha=0.10, color=COLOR["background"],
                   edgecolor="none", zorder=2, rasterized=True)
        ax.scatter(bulk.mean_expr, bulk.rho, s=30, alpha=0.85, color=col_s,
                   edgecolor="black", linewidth=0.4, zorder=4)
        offs = [(7, 5), (7, -11), (-8, 6), (-8, -12)]
        for k, (_, r) in enumerate(shown.sort_values("rho", ascending=False).iterrows()):
            ax.scatter([r.mean_expr], [r.rho], s=64, color=col_shown,
                       edgecolor="black", linewidth=0.8, zorder=6)
            dx, dy = offs[k % len(offs)]
            ax.annotate(f"$\\it{{{r.gene}}}$", (r.mean_expr, r.rho),
                        textcoords="offset points", xytext=(dx, dy), fontsize=8,
                        color=col_shown, zorder=7, ha=("left" if dx > 0 else "right"))
        re_, pe_ = spearmanr(S.mean_expr, S.rho)
        ax.text(0.03, 0.04, f"{stratum}: ρ(expr,ρ) = {re_:+.2f}\np = {pe_:.2f}   n = {len(S)}",
                transform=ax.transAxes, fontsize=7.5, va="bottom", ha="left")
        ax.axhline(0, color="0.75", lw=0.9, ls="--", zorder=1)
        ax.axhline(np.median(bg.rho), color="0.45", lw=0.9, ls=":", zorder=1)
        ax.set_xscale("log")
        ax.set_xlabel("Mean expression across clusters", fontsize=9)
        ax.set_ylabel("Spearman ρ with Gul", fontsize=9)
        ax.set_ylim(-1.05, 1.05); ax.tick_params(labelsize=8)
        ax.set_title(f"{G.TISSUE_NAME.get(tag, tag)} ({tag})", fontsize=10.5)
    handles = [Line2D([], [], marker="o", ls="none", mfc=COLOR["background"], mec="none",
                      ms=6, label="Non-target background"),
               Line2D([], [], marker="o", ls="none", mfc=col_s, mec="black", mew=0.4,
                      ms=7, label=f"conserved {stratum} target"),
               Line2D([], [], marker="o", ls="none", mfc=col_shown, mec="black", mew=0.8,
                      ms=8, label=f"{stratum} target among shown genes"),
               Line2D([], [], color="0.45", lw=0.9, ls=":", label="background median ρ")]
    fig.legend(handles=handles, frameon=False, ncol=4, loc="lower center",
               bbox_to_anchor=(0.5, -0.02), fontsize=8.5)
    fig.suptitle(f"{stratum} miR-7 targets: Gul correlation vs expression level", fontsize=11)
    fig.tight_layout(rect=[0, 0.10, 1, 0.93], w_pad=1.2)
    fig.savefig(path, bbox_inches="tight")
    fig.savefig(path.replace(".png", ".svg"), bbox_inches="tight")
    plt.close(fig)


if __name__ == "__main__":
    ap = argparse.ArgumentParser(description=__doc__,
                                 formatter_class=argparse.RawDescriptionHelpFormatter)
    ap.add_argument("--outdir", default=None,
                    help="write everything here; default routes to results/{tables,figures}")
    ap.add_argument("--datadir", default=None, help="extra directory to search for inputs")
    ap.add_argument("--keep-glia", action="store_true",
                    help="do NOT drop glial clusters (default: excluded)")
    ap.add_argument("--min-retain", type=float, default=MIN_RETAIN,
                    help="drop genes retained in less than this share of clusters "
                         f"(default {MIN_RETAIN}); 0 disables")
    args = ap.parse_args()
    cfg = dict(G.DEFAULT_CONFIG)
    O = lambda n: P.out(n, args.outdir)

    # Two tables on purpose. The retention filter is right for the figure and the
    # group tests, where dropping thin genes removes a 5-cluster artefact from the
    # top of the 8mer stratum and costs nothing against a background of thousands.
    # It is wrong for the downstream 8mer gene set, where it takes n from 33 to 27
    # and power is the binding constraint, so mirna_competition.py and
    # gul_duplex_model.py read the *unfiltered* table.
    D_all = collect(cfg, args.datadir, keep_glia=args.keep_glia)
    D = apply_retention(D_all, args.min_retain)
    T = group_tests(D)
    D_all.to_csv(O("gul_correlation_by_sitetype_pergene_unfiltered.csv"), index=False)
    D.to_csv(O("gul_correlation_by_sitetype_pergene.csv"), index=False)
    T.to_csv(O("gul_correlation_by_sitetype_tests.csv"), index=False)
    make_fig(D, T, [t["tag"] for t in cfg["tissues"]],
             O("fig_gul_correlation_by_sitetype.png"))
    make_expr_fig(D, O("fig_gul_correlation_8mer_vs_expr.png"), stratum="8mer")

    print("\n=== primary (shown genes excluded): median rho and perm p vs background ===")
    print(T[T.excl_shown].pivot_table(index="stratum", columns="tissue",
                                      values=["med_rho", "perm_p"])
          .reindex(STRATA).to_string(float_format=lambda x: f"{x:.3f}"))
```
