## supplemental tables and scripts for "Long non-coding RNA-3′UTR interactions coordinate GABA signaling in the brain": sitetype_stratification_gad1.py

```
#!/usr/bin/env python3
"""
Does correlation with Gad1 or with Gul scale with miR-7 site strength?

Companion to sitetype_stratification.py, with two deliberate differences:

  1. The anchor is selectable with --anchor. **Gad1** asks the question in the
     GABAergic-identity frame; **Gul** asks it about the regulator itself. The two
     are 0.88-0.93 collinear and their genome-wide rankings agree at 0.91-0.96, so
     the figures look nearly identical — the choice governs what each one may be
     used to claim, not what it shows.

  2. Genes whose only miR-7 sites are **poorly conserved are dropped entirely**,
     not folded into the background. The background here is genes with no
     annotated miR-7 site of any kind, so the contrast is site vs no site rather
     than site vs (no site + weak site). "Poorly conserved" means best-site
     PCT = 0 -- present in the alignment, conserved no better than chance; see
     load_site_types in sitetype_stratification.py for the evidence.

CIRCULARITY, which differs by anchor:
  --anchor Gad1  no shown gene was selected on Gad1 correlation, so labelling any
                 of them is safe. VGAT is the natural positive control.
  --anchor Gul   VGAT, sgll, ScsbetaG, and stol came from a BLAST search for
                 Gul complementarity and are independent of this ranking, but
                 **Ssadh and CG32225 were found by ranking Gul co-variation among
                 8mer targets** — this figure is the screen that produced them.
                 Label them as the screen's output, never as its confirmation.
Either way the exclude-shown control is computed and printed.

Inputs (all already in this folder):
  TargetScan7.2__miR-7-5p.predicted_targets.txt  per-gene conserved site counts
  mir7_targetscanfly_conserved_targets.tsv       the broad 549-gene list
  gene_by_cluster_meanexpr*.csv                  the cluster-mean matrices

Usage:
    python3 sitetype_stratification_gad1.py          # writes to results/{tables,figures}

Outputs are named `gad1_...` / `fig_gad1_...` under --anchor Gad1 and `gulanchor_...` /
`fig_gulanchor_...` under --anchor Gul. The Gul-anchored files are *not* called `gul_...`:
sitetype_stratification.py owns that namespace and writes a different five-stratum table
there (see PREFIX).
"""
import os, argparse
import numpy as np, pandas as pd
import paths as P
import genes as GN
import matplotlib
matplotlib.use("Agg")
import matplotlib.pyplot as plt
from matplotlib.lines import Line2D
from scipy.stats import mannwhitneyu

import gul_mir7_correlation as G
from sitetype_stratification import load_site_types

# anchor is selectable: Gad1 (GABAergic identity) or Gul itself. They are 0.88-0.93
# collinear, so the two rankings are near-identical (rank correlation 0.91-0.96); the
# choice matters for what the figure *claims*, not for what it shows.
ANCHORS = {"Gad1": ("Gad1", "Gad1"), "Gul": ("CG14989", "Gul")}
ANCHOR, ANCHOR_LABEL = ANCHORS["Gad1"]      # overridden by --anchor

#: Output-name prefix per anchor -- deliberately NOT the anchor's own name for Gul.
#: sitetype_stratification.py already owns `gul_correlation_by_sitetype_*`, and its
#: version of those files is a different analysis: five strata, keeping the
#: poorly-conserved class that this script drops (difference 2 above), plus an
#: `n_total` column. The Gul-anchored run must NOT be named `gul_`, or it collides with
#: sitetype_stratification.py, which writes a different table under that prefix.
PREFIX = {"Gad1": "gad1", "Gul": "gulanchor"}

# Which genes get a label. Symbols are canonicalised at load (genes.py), so a
# gene carries one name in every panel regardless of which FlyBase release the
# tissue's matrix was built against.
LABEL_SETS = {
    "vgat":  {"VGAT": "VGAT"},
    "panel": {g: g for g in GN.PANEL},
}
GENE_COLORS = dict(GN.COLORS)

#: minimum share of clusters a gene must be retained in by the dead-tail filter
MIN_RETAIN = 0.25

# poorly-conserved-only genes are dropped, not merged into the background
STRATA = ["background", "7mer-A1", "7mer-m8", "8mer"]
LABEL  = {"background": "Bg", "7mer-A1": "7mer\nA1",
          "7mer-m8": "7mer\nm8", "8mer": "8mer"}
COLOR  = {"background": "#8A8A8A", "7mer-A1": "#9ECAE1",
          "7mer-m8": "#4C72B0", "8mer": "#08306B"}
LEGEND = {"background": "No annotated miR-7 site",
          "7mer-A1": "conserved 7mer-A1 site",
          "7mer-m8": "conserved 7mer-m8 site",
          "8mer": "conserved 8mer site"}
COL_LABEL = "#C44E52"


def collect(cfg, datadir=None, keep_glia=False, min_retain=MIN_RETAIN):
    """Per-gene rho with Gad1 + stratum for every tissue.

    Genes carrying only poorly conserved miR-7 sites are assigned stratum
    ``drop`` and removed before any test or plot.

    Genes retained in fewer than ``min_retain`` of clusters by the per-gene
    dead-tail filter are also dropped. Without this the top of the 8mer stratum
    is a five-cluster artefact rather than a result: Brd in OL and E(spl)m5-HLH
    in CB both reach rho = +0.78 on 5 of 231 and 5 of 346 clusters, at mean
    expression ~1e-5 and p = 0.12, and both plot above VGAT. The same threshold
    is what the genome-wide ranking in the text uses.
    """
    site, _ = load_site_types(datadir)
    broad = G.load_mir7_symbols(P.find(cfg["mir7_targets_tsv"], datadir))
    frames = []
    for t in cfg["tissues"]:
        M = GN.canon_index(pd.read_csv(P.find(t["matrix_csv"], datadir), index_col=0))
        if not keep_glia:
            M = G.drop_glia(M, t["tag"])
        A = G.analyze(t["tag"], M, ANCHOR, t["shown"], t["targets"],
                      set(broad) | set(t.get("mir7_extra", [])))
        res = A["res"]; extra = set(t.get("mir7_extra", []))

        def stratum(g):
            s = site.get(g)
            if s: return s
            return "drop" if (g in broad or g in extra) else "background"

        d = pd.DataFrame(dict(
            tissue=t["tag"], gene=res.index, rho=res.rho.values,
            mean_expr=res.mean_expr.values, n_clusters=res.n_clusters.values,
            stratum=[stratum(g) for g in res.index],
            is_shown=[g in set(t["shown"]) for g in res.index]))
        n_drop = int((d.stratum == "drop").sum())
        d = d[d.stratum != "drop"]
        cut = min_retain * M.shape[1]
        n_thin = int((d.n_clusters < cut).sum())
        d = d[d.n_clusters >= cut]
        frames.append(d)
        print(f"[{t['tag']}] {len(d)} genes stratified "
              f"({n_drop} poorly-conserved-only dropped, "
              f"{n_thin} retained in < {min_retain:.0%} of {M.shape[1]} clusters dropped)")
    return pd.concat(frames, ignore_index=True)


def group_tests(D):
    rows = []
    for tag in D.tissue.unique():
        for excl in (False, True):
            d = D[D.tissue == tag]
            if excl: d = d[~d.is_shown]
            bg = d[d.stratum == "background"].rho.values
            for s in STRATA:
                v = d[d.stratum == s].rho.values
                if s == "background":
                    rows.append(dict(tissue=tag, excl_shown=excl, stratum=s, n=len(v),
                                     med_rho=np.median(v), med_bg=np.median(bg),
                                     mean_rho=np.mean(v), mean_bg=np.mean(bg),
                                     mwu_p=np.nan, perm_p=np.nan,
                                     perm_p_mean=np.nan)); continue
                if len(v) < 3: continue
                _, pp = G.perm_test(v, bg)                      # medians
                _, pm = G.perm_test(v, bg, stat="mean")          # tail-sensitive
                rows.append(dict(tissue=tag, excl_shown=excl, stratum=s, n=len(v),
                                 med_rho=np.median(v), med_bg=np.median(bg),
                                 mean_rho=np.mean(v), mean_bg=np.mean(bg),
                                 mwu_p=mannwhitneyu(v, bg, alternative="greater").pvalue,
                                 perm_p=pp, perm_p_mean=pm))
    return pd.DataFrame(rows)


def _declutter(ys, gap, lo, hi):
    """Nudge label positions apart, preserving order, staying inside [lo, hi]."""
    idx = np.argsort(ys)
    y0 = np.sort(np.asarray(ys, float))
    y = y0.copy()
    for i in range(1, len(y)):                       # push up to open the gaps
        y[i] = max(y[i], y[i-1] + gap)
    y -= (y.mean() - y0.mean())                      # re-centre on the points
    if y[-1] > hi: y -= y[-1] - hi                   # then keep it on the axes
    if y[0] < lo:  y += lo - y[0]
    out = np.empty_like(y); out[idx] = y
    return out


def make_fig(D, T, order, path, labels=None, figsize=None, panel=False):
    """labels: {matrix symbol: display name}; each gets a coloured marker and a
    leader line to a name in the right-hand margin.

    figsize overrides the default, for placing this as a panel in a composite.
    panel strips the suptitle and the legend, which belong in the caption -- the
    same convention fig_gad1_enrichment and fig_class_vs_six follow. Between them
    that is about 0.9 in of height recovered."""
    labels = labels or LABEL_SETS["vgat"]
    plt.rcParams.update({
        "figure.dpi": 110, "savefig.dpi": 200,
        "axes.spines.top": False, "axes.spines.right": False,
        "svg.fonttype": "none", "font.family": "sans-serif", "font.sans-serif": ["Arial"],
    })
    S = T[T.excl_shown == False]          # all genes retained; see the circularity note
    n = len(order)
    wide = len(set(labels.values())) > 1  # room for a label margin
    fig, axes = plt.subplots(1, n, figsize=figsize or (
        ((4.2 if wide else 3.4) if n == 1 else (10.2 if wide else 8.0)), 4.6))
    x_lab = len(STRATA) - 0.55            # where the label column sits
    for ax, tag in zip(np.atleast_1d(axes), order):
        d = D[D.tissue == tag]
        st = S[S.tissue == tag].set_index("stratum")
        ax.axhline(np.median(d[d.stratum == "background"].rho.values),
                   color="0.45", lw=0.9, ls=":", zorder=2)
        marks = []                          # (x, y, display name) for the label pass
        for i, s in enumerate(STRATA):
            v = d[d.stratum == s]
            if not len(v): continue
            big = (s == "background")
            jit = (np.random.default_rng(i).random(len(v)) - 0.5) * (0.34 if big else 0.30)
            ax.scatter(np.full(len(v), i) + jit, v.rho.values, s=(7 if big else 16),
                       alpha=(0.13 if big else 0.55), color=COLOR[s],
                       edgecolor="none", zorder=3, rasterized=True)
            ax.plot([i-0.30, i+0.30], [np.median(v.rho)]*2, color="black", lw=2.0, zorder=5)
            ax.text(i, 1.30, f"n={len(v)}", ha="center", va="bottom", fontsize=7, color="0.3")
            if s != "background":
                ax.text(i, 1.13, f"p={st.loc[s, 'perm_p']:.2f}", ha="center",
                        va="bottom", fontsize=7.5)
            # highlight the labelled genes where they actually fall
            for k, g in enumerate(v.gene.values):
                if g not in labels: continue
                name = labels[g]
                # always on the gene's own jittered x, so the highlight covers its
                # cloud point instead of appearing beside it. Offsetting the marker
                # (previously i + 0.30 whenever there was a single label) drew every
                # labelled gene twice: once dark in the stratum, once coloured to the
                # right, at the same rho.
                x = i + jit[k]
                y = float(v.rho.values[k])
                ax.scatter([x], [y], s=46, color=GENE_COLORS.get(name, COL_LABEL),
                           edgecolor="black", linewidth=0.7, zorder=6, clip_on=False)
                marks.append((x, y, name))

        if wide:                            # leader lines out to a right-hand column
            marks.sort(key=lambda m: -m[1])
            ys = _declutter([m[1] for m in marks], 0.115, -1.05, 1.05)
            for (x, y, name), yl in zip(marks, ys):
                c = GENE_COLORS.get(name, COL_LABEL)
                ax.annotate(f"$\\it{{{name}}}$", (x, y), xycoords="data",
                            xytext=(x_lab + 0.62, yl), textcoords="data",
                            fontsize=8.5, color=c, ha="left", va="center", zorder=7,
                            annotation_clip=False,
                            arrowprops=dict(arrowstyle="-", color=c, lw=0.7,
                                            shrinkA=2, shrinkB=3,
                                            connectionstyle="arc3,rad=0"))
        else:
            for x, y, name in marks:
                ax.annotate(f"$\\it{{{name}}}$", (x, y), textcoords="offset points",
                            xytext=(7, -3), fontsize=8.5, color=COL_LABEL,
                            ha="left", va="center", zorder=7, annotation_clip=False)

        ax.axhline(0, color="0.85", lw=0.8, ls="--", zorder=1)
        ax.set_xticks(range(len(STRATA)))
        ax.set_xticklabels([LABEL[s] for s in STRATA], fontsize=8.5)
        ax.set_xlim(-0.6, len(STRATA) - 0.25 + (1.15 if wide else 0)); ax.set_ylim(-1.1, 1.5)
        ax.tick_params(axis="y", labelsize=8.5)
        ax.spines["bottom"].set_bounds(-0.6, len(STRATA) - 0.25)   # not into the margin
        if n > 1:   # single panel names its tissue in the suptitle instead
            ax.set_title(f"{G.TISSUE_NAME.get(tag, tag)} ({tag})", fontsize=10.5)
        ax.set_ylabel(f"Spearman ρ with {ANCHOR_LABEL}", fontsize=9.5)
    handles = [Line2D([], [], marker="o", ls="none", mfc=COLOR[s], mec="none", ms=7,
                      label=LEGEND[s]) for s in STRATA]
    if not wide:   # a single highlighted gene needs a key; a labelled panel does not
        handles += [Line2D([], [], marker="o", ls="none", mfc=COL_LABEL, mec="black",
                           mew=0.7, ms=8, label=list(labels.values())[0])]
    handles += [Line2D([], [], color="black", lw=2.0, label="group median"),
                Line2D([], [], color="0.45", lw=0.9, ls=":", label="background median")]
    title = f"{ANCHOR_LABEL} correlation by miR-7 site strength"
    if n == 1:
        tag = order[0]
        title += f"\n{G.TISSUE_NAME.get(tag, tag)} ({tag})"
    if not panel:
        fig.legend(handles=handles, frameon=False, ncol=(2 if n == 1 else 4),
                   loc="lower center", bbox_to_anchor=(0.5, 0.0), fontsize=8.5,
                   handletextpad=0.5, columnspacing=1.2, labelspacing=0.4)
        fig.suptitle(title, fontsize=11)
    fig.tight_layout(rect=[0, (0.15 if n == 1 else 0.12), 1, (0.90 if n == 1 else 0.94)],
                     w_pad=1.0)
    fig.savefig(path, bbox_inches="tight")
    fig.savefig(path.replace(".png", ".svg"), bbox_inches="tight")
    plt.close(fig)


if __name__ == "__main__":
    ap = argparse.ArgumentParser(description=__doc__,
                                 formatter_class=argparse.RawDescriptionHelpFormatter)
    ap.add_argument("--outdir", default=None,
                    help="write everything here; default routes to results/{tables,figures}")
    ap.add_argument("--datadir", default=None, help="extra directory to search for inputs")
    ap.add_argument("--keep-glia", action="store_true",
                    help="do NOT drop glial clusters (default: excluded)")
    ap.add_argument("--min-retain", type=float, default=MIN_RETAIN,
                    help="drop genes retained in less than this share of clusters "
                         f"(default {MIN_RETAIN}); 0 disables")
    ap.add_argument("--tissues", default="OL,VNC,CB",
                    help="comma-separated subset, e.g. 'OL' for a single panel")
    ap.add_argument("--labels", default="vgat", choices=sorted(LABEL_SETS),
                    help="which genes to label: 'vgat' or the 7-gene 'panel'")
    ap.add_argument("--anchor", default="Gad1", choices=sorted(ANCHORS),
                    help="correlate against Gad1 (identity) or Gul (the regulator)")
    ap.add_argument("--panel", action="store_true",
                    help="strip the title and legend for a composite; they belong in "
                         "the caption, and it recovers about 0.9 in of height")
    ap.add_argument("--figsize", nargs=2, type=float, default=None,
                    metavar=("W", "H"), help="override the figure size, in inches")
    args = ap.parse_args()
    ANCHOR, ANCHOR_LABEL = ANCHORS[args.anchor]
    prefix = PREFIX[args.anchor]        # see PREFIX: not ANCHOR_LABEL.lower()
    keep = [s.strip() for s in args.tissues.split(",") if s.strip()]
    suffix = "" if keep == ["OL", "VNC", "CB"] else "_" + "".join(keep)
    if args.labels != "vgat":
        suffix = f"_{args.labels}" + suffix

    cfg = dict(G.DEFAULT_CONFIG)
    O = lambda n: P.out(n, args.outdir)
    cfg["tissues"] = [t for t in cfg["tissues"] if t["tag"] in keep]
    assert cfg["tissues"], f"no tissue matched {keep}"

    labels = LABEL_SETS[args.labels]
    D = collect(cfg, args.datadir, keep_glia=args.keep_glia, min_retain=args.min_retain)
    T = group_tests(D)
    D.to_csv(O(f"{prefix}_correlation_by_sitetype_pergene{suffix}.csv"), index=False)
    T.to_csv(O(f"{prefix}_correlation_by_sitetype_tests{suffix}.csv"), index=False)
    make_fig(D, T, [t["tag"] for t in cfg["tissues"]],
             O(f"fig_{prefix}_correlation_by_sitetype{suffix}"
               # NOT "_panel": --labels panel already claims that suffix, and using
               # it here silently overwrote the 7-gene-labelled figure with a
               # title-stripped one
               + ("_bare" if args.panel else "") + ".png"),
             labels=labels, figsize=args.figsize, panel=args.panel)

    for excl, name in [(False, "all genes (plotted)"), (True, "shown genes excluded (control)")]:
        print(f"\n=== {name}: median rho with {ANCHOR_LABEL} and perm p vs background ===")
        print(T[T.excl_shown == excl].pivot_table(index="stratum", columns="tissue",
                                                  values=["med_rho", "perm_p", "n"])
              .reindex(STRATA).to_string(float_format=lambda x: f"{x:.3f}"))

    L = D[D.gene.isin(labels)].copy()
    L["label"] = [labels[g] for g in L.gene]
    print(f"\nlabelled genes, rho with {ANCHOR_LABEL}:")
    print(L.pivot_table(index=["label", "stratum"], columns="tissue", values="rho")
          .round(3).to_string())
    missing = set(labels.values()) - set(L.label)
    if missing:
        print(f"  NOT FOUND in the matrices: {sorted(missing)}")
```
