## supplemental tables and scripts for "Long non-coding RNA-3′UTR interactions coordinate GABA signaling in the brain": paths.py

```
#!/usr/bin/env python3
"""Input files for the miR-7:VGAT duplex analyses, resolved beside this script.

Every path is relative to this file, so the folder runs wherever it is unpacked.
`data/` holds two FASTA files:

    VGAT_3UTR_mir7_region.fa   the 205 nt VGAT 3'UTR fragment carrying the three
                               miR-7 sites; the sequence run_cofold.py and
                               thermo.py fold against
    mir7_align_vgat.fa         the 24 nt site windows and reverse-complemented
                               miR-7 used as RNAcofold input by run_cofold.py
"""
import os

HERE = os.path.dirname(os.path.abspath(__file__))
DATA = os.path.join(HERE, "data")

UTR_FA = os.path.join(DATA, "VGAT_3UTR_mir7_region.fa")
INPUTS = os.path.join(DATA, "mir7_align_vgat.fa")


def require(path):
    """Return `path`, or exit with a message naming what is missing and where."""
    if not os.path.exists(path):
        raise SystemExit(
            "missing input: %s\n"
            "Expected it in %s\n"
            "The three FASTA inputs ship with these scripts; if data/ is empty the "
            "folder was not copied whole." % (os.path.basename(path), DATA))
    return path
```
