## supplemental tables and scripts for "Long non-coding RNA-3′UTR interactions coordinate GABA signaling in the brain": run_cofold.py

```
#!/usr/bin/env python3
"""RNAcofold on the three VGAT miR-7 sites, using the 24-nt site windows."""

import RNA

import paths as P

INPUTS = P.require(P.INPUTS)
UTR_FA = P.require(P.UTR_FA)
MIR = "UGGAAGACUAGUGAUUUUGUUGU"
WCONLY = {("A", "U"), ("U", "A"), ("G", "C"), ("C", "G")}


def read_fa(path):
    recs, name, seq = [], None, []
    for line in open(path):
        line = line.rstrip("\n")
        if line.startswith(">"):
            if name:
                recs.append((name, "".join(seq)))
            name, seq = line[1:], []
        elif line.strip():
            seq.append(line.strip())
    if name:
        recs.append((name, "".join(seq)))
    return recs


U = "".join(l.strip() for l in open(UTR_FA) if not l.startswith(">")).upper()
inputs = {n.split("/")[0]: s.upper() for n, s in read_fa(INPUTS)}


def draw(tgt, prs):
    """prs: {target_pos: mir_pos}, 1-based. Returns three aligned lines."""
    ts = sorted(prs)
    top, mid, bot = [], [], []
    lt = tgt[:ts[0] - 1]
    lm = "".join(MIR[m - 1] for m in range(len(MIR), prs[ts[0]], -1))
    w = max(len(lt), len(lm))
    top.append(lt.rjust(w)); bot.append(lm.rjust(w)); mid.append(" " * w)
    for k, t in enumerate(ts):
        m = prs[t]
        if k:
            pt, pm = ts[k - 1], prs[ts[k - 1]]
            gt = tgt[pt:t - 1]
            gm = "".join(MIR[x - 1] for x in range(pm - 1, m, -1))
            g = max(len(gt), len(gm))
            top.append(gt.ljust(g, "-")); bot.append(gm.ljust(g, "-"))
            mid.append(" " * g)
        top.append(tgt[t - 1]); bot.append(MIR[m - 1])
        mid.append("|" if (tgt[t - 1], MIR[m - 1]) in WCONLY else ":")
    rt = tgt[ts[-1]:]
    rm = "".join(MIR[m - 1] for m in range(prs[ts[-1]] - 1, 0, -1))
    w = max(len(rt), len(rm))
    top.append(rt.ljust(w)); bot.append(rm.ljust(w)); mid.append(" " * w)
    return "".join(top), "".join(mid), "".join(bot)


print("=" * 74)
print("RNAcofold — VGAT site + dme-miR-7-5p, 37 C, Turner 2004, dangles=2")
print("=" * 74)
print("Inputs are the 24-nt windows from mir7_align_vgat.fa\n")

rows = []
for site in ("vgat-mir7-2", "vgat-mir7-3", "vgat-mir7-4"):
    tgt = inputs[site].replace("T", "U")
    n = len(tgt)
    utr_lo = U.find(inputs[site]) + 1

    st, e_ab = RNA.fold_compound(tgt + "&" + MIR).mfe_dimer()
    _, e_a = RNA.fold(tgt)                 # target monomer
    _, e_b = RNA.fold(MIR)                 # miR-7 monomer
    e_bind = e_ab - e_a - e_b

    stack, inter, intra = [], {}, []
    for i, c in enumerate(st, 1):
        if c == "(":
            stack.append(i)
        elif c == ")":
            t = stack.pop()
            if t <= n < i:
                inter[t] = i - n
            else:
                intra.append((t, i))

    rows.append((site, e_ab, e_a, e_b, e_bind, len(inter)))
    print(f"--- {site}   (VGAT 3'UTR {utr_lo}-{utr_lo+n-1}, {n} nt)")
    print(f"    {tgt}&{MIR}")
    print(f"    {st[:n]}&{st[n:]}")
    print()
    print(f"    dG(AB) dimer            = {e_ab:7.2f} kcal/mol")
    print(f"    dG(A)  VGAT alone       = {e_a:7.2f}")
    print(f"    dG(B)  miR-7 alone      = {e_b:7.2f}")
    print(f"    dG(binding) = AB-A-B    = {e_bind:7.2f}")
    print(f"    intermolecular pairs    = {len(inter)}")
    print(f"    intramolecular pairs    = {len(intra)}  "
          f"{'(none - nothing competes with the duplex)' if not intra else intra}")
    print(f"    miR-7 positions paired  = {sorted(inter.values())}")
    print()
    t, m, b = draw(tgt, inter)
    print(f"      VGAT  5'-{t}-3'")
    print(f"               {m}")
    print(f"      miR-7 3'-{b}-5'")
    print()

print("=" * 74)
print("Summary")
print("=" * 74)
print(f"   {'site':14s}{'dG(AB)':>9s}{'dG(A)':>8s}{'dG(B)':>8s}{'dG(bind)':>10s}{'pairs':>7s}")
for s, ab, a, b, bind, np_ in rows:
    print(f"   {s:14s}{ab:9.2f}{a:8.2f}{b:8.2f}{bind:10.2f}{np_:7d}")
print()
print("   ranking by dG(AB)     : " +
      " > ".join(r[0] for r in sorted(rows, key=lambda r: r[1])))
print("   ranking by dG(binding): " +
      " > ".join(r[0] for r in sorted(rows, key=lambda r: r[4])))
```
