## supplemental tables and scripts for "Long non-coding RNA-3′UTR interactions coordinate GABA signaling in the brain": thermo.py

```
#!/usr/bin/env python3
"""Thermodynamic miR-7:VGAT duplex analysis with ViennaRNA (RNAduplex equivalent)."""

import RNA

import paths as P

UTR_FA = P.require(P.UTR_FA)
MIR = "UGGAAGACUAGUGAUUUUGUUGU"          # dme-miR-7-5p 5'->3'
SITES = [("vgat-miR7-2", 42), ("vgat-miR7-3", 168), ("vgat-miR7-4", 191)]


def utr_rna():
    s = "".join(l.strip() for l in open(UTR_FA) if not l.startswith(">"))
    return s.upper().replace("T", "U")


def gapless_pairs(t, m3to5):
    wc = {("A", "U"), ("U", "A"), ("G", "C"), ("C", "G")}
    gu = {("G", "U"), ("U", "G")}
    w = sum(1 for a, b in zip(t, m3to5) if (a, b) in wc)
    g = sum(1 for a, b in zip(t, m3to5) if (a, b) in gu)
    return w, g


utr = utr_rna()
print("=" * 74)
print("1. RNAduplex (RNA.duplexfold): MFE intermolecular structure per site")
print("=" * 74)
print("   Turner 2004 params. Bulges/loops allowed but ENERGY-scored, not")
print("   identity-scored. G:U wobbles are real pairs here.\n")

rows = []
for name, a1 in SITES:
    win = utr[a1 - 23:a1]
    d = RNA.duplexfold(win, MIR)
    wc, gu = gapless_pairs(win, MIR[::-1])
    rows.append((name, a1, win, d, wc, gu))
    print(f"--- {name}   UTR {a1-22}-{a1}")
    print(f"    target 5'-{win}-3'")
    print(f"    miR-7  5'-{MIR}-3'")
    print(f"    MFE structure : {d.structure}")
    print(f"    dG            : {d.energy:.2f} kcal/mol")
    print(f"    duplex spans  : target ..{d.i}   miRNA {d.j}..")
    print(f"    my gapless count: {wc} WC + {gu} G:U = {wc+gu} bp")
    print()

print("=" * 74)
print("2. Ranking by dG vs ranking by base-pair count")
print("=" * 74)
by_dg = sorted(rows, key=lambda r: r[3].energy)
by_bp = sorted(rows, key=lambda r: -(r[4] + r[5]))
print(f"   {'by dG (most stable first)':<34}{'by bp count (most pairs first)'}")
for a, b in zip(by_dg, by_bp):
    print(f"   {a[0]} {a[3].energy:7.2f} kcal/mol       "
          f"{b[0]} {b[4]+b[5]:2d} bp")
print()

print("=" * 74)
print("3. Does the MFE structure use bulges, or is it the gapless duplex?")
print("=" * 74)
for name, a1, win, d, wc, gu in rows:
    t_struct, m_struct = d.structure.split("&")
    # count paired positions on each side
    tp, mp = t_struct.count("("), m_struct.count(")")
    # contiguity: a gapless seed-anchored duplex has no internal '.' runs
    core = t_struct.strip(".")
    internal_dots = core.count(".")
    print(f"   {name:14s} paired={tp:2d}  internal unpaired in target arm={internal_dots:2d}  "
          f"{'BULGED' if internal_dots else 'contiguous'}")
print()

print("=" * 74)
print("4. Suboptimal duplexes within 2 kcal/mol (alternative registers?)")
print("=" * 74)
for name, a1 in SITES:
    win = utr[a1 - 23:a1]
    subs = RNA.duplex_subopt(win, MIR, 200, 0)
    seen = []
    for s in subs:
        seen.append((s.energy, s.structure))
    seen = sorted(set(seen))[:4]
    print(f"   {name}:")
    for e, st in seen:
        print(f"      {e:7.2f}  {st}")
    print()

print("=" * 74)
print("5. Scan the WHOLE UTR: where does miR-7 bind best, unconstrained?")
print("=" * 74)
best = RNA.duplexfold(utr, MIR)
print(f"   best duplex anywhere in the 205-nt UTR:")
print(f"      dG = {best.energy:.2f} kcal/mol   ends at UTR position {best.i}")
print(f"      structure {best.structure}")
print()
for name, a1 in SITES:
    print(f"      ({name} A1 sits at UTR {a1})")
```
