## supplemental tables and scripts for "Long non-coding RNA-3′UTR interactions coordinate GABA signaling in the brain": analyze.py

```
#!/usr/bin/env python3
"""ORF analysis of FBgn0035495 (CG14989): RF (short, suspected non-coding) vs RE (long, used ORF)."""
import os

HERE = os.path.dirname(os.path.abspath(__file__))
D    = os.path.join(HERE, "data")      # inputs
OUT  = os.path.join(HERE, "results")   # outputs

def read_fasta(path):
    seq = []
    with open(path) as fh:
        for line in fh:
            if line.startswith(">"):
                continue
            seq.append(line.strip())
    return "".join(seq).upper()

re_cdna = read_fasta(os.path.join(D, "re_cdna.fa"))
re_cds  = read_fasta(os.path.join(D, "re_cds.fa"))
rf_cdna = read_fasta(os.path.join(D, "rf_cdna.fa"))
rf_cds  = read_fasta(os.path.join(D, "rf_cds.fa"))

STOPS = {"TAA", "TAG", "TGA"}

def find_orfs(seq, min_aa=1):
    """Return ATG->stop ORFs (0-based start, end-exclusive of stop). Only complete ORFs (with stop)."""
    orfs = []
    n = len(seq)
    for frame in range(3):
        i = frame
        while i + 3 <= n:
            if seq[i:i+3] == "ATG":
                j = i
                while j + 3 <= n:
                    cod = seq[j:j+3]
                    if cod in STOPS:
                        aa = (j - i) // 3           # aa excluding stop
                        if aa >= min_aa:
                            orfs.append({"frame": frame, "start": i, "stop_end": j+3,
                                         "cds_end": j, "aa": aa, "nt": j+3-i})
                        i = j  # move past; next ATG search continues after this codon
                        break
                    j += 3
                else:
                    # no stop until end (incomplete ORF)
                    pass
            i += 3
    return orfs

def cds_location(cdna, cds):
    idx = cdna.find(cds)
    return idx  # 0-based start of CDS in cDNA, -1 if not found

# --- RE: locate used CDS in cDNA ---
re_cds_start = cds_location(re_cdna, re_cds)
print("=== RE (FBtr0308363, long isoform) ===")
print(f"cDNA length: {len(re_cdna)} nt")
print(f"CDS: {len(re_cds)} nt -> {len(re_cds)//3 - 1} aa + stop")
print(f"CDS start in cDNA (0-based): {re_cds_start}  (1-based: {re_cds_start+1})")
print(f"  5'UTR: {re_cds_start} nt | CDS: {len(re_cds)} nt | 3'UTR: {len(re_cdna)-re_cds_start-len(re_cds)} nt")
print(f"  ATG at cDNA {re_cds_start+1}, stop ends at cDNA {re_cds_start+len(re_cds)}")

# --- RF: locate annotated CDS and enumerate all ORFs ---
rf_cds_start = cds_location(rf_cdna, rf_cds)
print("\n=== RF (FBtr0345137, short/suspected non-coding) ===")
print(f"cDNA length: {len(rf_cdna)} nt")
print(f"Annotated CDS: {len(rf_cds)} nt -> {len(rf_cds)//3 - 1} aa + stop")
print(f"Annotated CDS start in cDNA (0-based): {rf_cds_start} (1-based: {rf_cds_start+1}), frame {rf_cds_start%3}")
print(f"  5'UTR (leader): {rf_cds_start} nt | CDS: {len(rf_cds)} nt | 3'UTR: {len(rf_cdna)-rf_cds_start-len(rf_cds)} nt")

orfs = find_orfs(rf_cdna, min_aa=1)
orfs.sort(key=lambda o: -o["aa"])
print(f"\nAll complete ATG->stop ORFs in RF cDNA (both counts): total={len(orfs)}")
print(f"ORFs >= 20 aa: {sum(1 for o in orfs if o['aa']>=20)}")
print(f"ORFs >= 50 aa: {sum(1 for o in orfs if o['aa']>=50)}")
print("\nTop ORFs by length:")
print(f"{'aa':>5} {'nt':>5} {'frame':>5} {'start(1b)':>9} {'stopEnd(1b)':>11}  note")
for o in orfs[:12]:
    note = ""
    if o["start"] == rf_cds_start:
        note = "<-- FlyBase-annotated 171-aa ORF"
    print(f"{o['aa']:>5} {o['nt']:>5} {o['frame']:>5} {o['start']+1:>9} {o['stop_end']:>11}  {note}")

# upstream ATGs / uORFs before the annotated start
u = [o for o in orfs if o["start"] < rf_cds_start]
print(f"\nComplete ORFs entirely/partly starting upstream of annotated ATG: {len(u)}")
# count ATGs upstream of annotated start
n_upstream_atg = sum(1 for i in range(rf_cds_start) if rf_cdna[i:i+3]=="ATG")
print(f"ATG codons upstream of annotated start (potential to initiate first): {n_upstream_atg}")

# Does RF contain RE's start-codon region (the real N-terminus)?
re_nterm = re_cds[:60]
print(f"\nDoes RF cDNA contain RE's ORF 5' region (first 60 nt of RE CDS)? {'YES' if re_nterm in rf_cdna else 'NO'}")

# Save a machine-readable summary for the plotting script
import json
summary = {
    "re": {"cdna_len": len(re_cdna), "cds_start": re_cds_start, "cds_len": len(re_cds), "aa": len(re_cds)//3-1},
    "rf": {"cdna_len": len(rf_cdna), "ann_cds_start": rf_cds_start, "ann_cds_len": len(rf_cds), "ann_aa": len(rf_cds)//3-1,
           "orfs": orfs},
}
with open(os.path.join(D, "summary.json"), "w") as fh:
    json.dump(summary, fh)
print("\nSaved summary.json")
```
