## supplemental tables and scripts for "Long non-coding RNA-3′UTR interactions coordinate GABA signaling in the brain": cons.py

```
#!/usr/bin/env python3
"""Fetch phyloP124way / phastCons124way (dm6, 124 insects) means over key CG14989 segments."""
import json, urllib.request

def fetch(track, chrom, start1, end1):
    # Ensembl 1-based inclusive -> UCSC 0-based half-open
    url = (f"https://api.genome.ucsc.edu/getData/track?genome=dm6;track={track};"
           f"chrom={chrom};start={start1-1};end={end1}")
    with urllib.request.urlopen(url, timeout=60) as r:
        d = json.load(r)
    vals = []
    data = d.get(track, [])
    if isinstance(data, dict):
        data = sum(data.values(), []) if all(isinstance(v, list) for v in data.values()) else [data]
    for item in data:
        if isinstance(item, dict) and "value" in item:
            span = item.get("end", item["start"]+1) - item["start"]
            vals.extend([item["value"]]*max(1, span))
    return vals

segments = [
    ("RE-unique N-term CDS (has real ATG; absent in RF)", 4088967, 4089130),
    ("RF 5' leader (before TTG; = RE coding seq)",         4089131, 4089191),
    ("Shared coding exon (both, RE frame)",                4090273, 4090661),
    ("3'UTR control (both, past stop)",                    4090800, 4091302),
]
chrom = "chr3L"
print(f"{'segment':52} {'len':>4} {'phyloP mean':>11} {'phastCons mean':>14}")
print("-"*86)
for name, s, e in segments:
    pp = fetch("phyloP124way", chrom, s, e)
    pc = fetch("phastCons124way", chrom, s, e)
    mpp = sum(pp)/len(pp) if pp else float('nan')
    mpc = sum(pc)/len(pc) if pc else float('nan')
    print(f"{name:52} {e-s+1:>4} {mpp:>11.3f} {mpc:>14.3f}")
print("\nphyloP: per-base conservation (higher=more constrained; ~>1 constrained). phastCons: P(conserved) 0-1.")
```
