## supplemental tables and scripts for "Long non-coding RNA-3′UTR interactions coordinate GABA signaling in the brain": kozak.py

```
#!/usr/bin/env python3
"""Kozak / translation-initiation context of candidate start codons.
Drosophila consensus (Cavener 1987): (C/A) A A (A/C) ATG ;  key positions -3 (purine, A preferred) and +4 (G)."""
import os
HERE = os.path.dirname(os.path.abspath(__file__))
D    = os.path.join(HERE, "data")      # inputs
OUT  = os.path.join(HERE, "results")   # outputs
def read_fasta(p): return "".join(l.strip() for l in open(p) if not l.startswith(">")).upper()
re_cdna = read_fasta(f"{D}/re_cdna.fa"); re_cds = read_fasta(f"{D}/re_cds.fa")
rf_cdna = read_fasta(f"{D}/rf_cdna.fa"); rf_cds = read_fasta(f"{D}/rf_cds.fa")

def context(seq, start0):
    """Return (-6..-1, codon, +4) around a start codon at 0-based start0."""
    up = seq[max(0,start0-6):start0]
    up = up.rjust(6, ".")
    codon = seq[start0:start0+3]
    plus4 = seq[start0+3] if start0+3 < len(seq) else "."
    m3 = seq[start0-3] if start0-3 >= 0 else "."
    return up, codon, plus4, m3

def classify(m3, plus4):
    purine = m3 in "AG"
    g4 = plus4 == "G"
    if purine and g4: return "STRONG (purine -3 + G +4)"
    if purine or g4:  return "moderate (" + ("purine -3" if purine else "G +4") + " only)"
    return "WEAK (no purine -3, no G +4)"

def cavener(up, m3):
    # Drosophila consensus (C/A)AA(A/C) at -4..-1 ; report simple match count
    # up = positions -6,-5,-4,-3,-2,-1
    p = {-4: up[2], -3: up[3], -2: up[4], -1: up[5]}
    ok = 0
    ok += p[-4] in "CA"; ok += p[-3] in "A";  # Cavener strongest: -3 A ; but A/C tolerated
    ok += p[-2] in "A";  ok += p[-1] in "AC"
    return f"{ok}/4 Cavener positions (-4 {p[-4]}, -3 {p[-3]}, -2 {p[-2]}, -1 {p[-1]})"

re_atg = re_cdna.find(re_cds)
rf_ttg = rf_cdna.find(rf_cds)

candidates = [
    ("RE  used ATG (246 aa)",            re_cdna, re_atg),
    ("RF  annotated start (171 aa)",     rf_cdna, rf_ttg),
    ("RF  ATG frame+2 (115 aa, nt230)",  rf_cdna, 229),
    ("RF  ATG frame+1 (101 aa, nt250)",  rf_cdna, 249),
    ("RF  ATG frame+3 (65 aa,  nt399)",  rf_cdna, 398),
    ("RF  ATG frame+3 (44 aa,  nt150)",  rf_cdna, 149),
]

print(f"{'start codon':38} {'context (-6..-1 | codon | +4)':30} {'-3':>3} {'+4':>3}  strength / Cavener")
print("-"*118)
for name, seq, s0 in candidates:
    up, codon, plus4, m3 = context(seq, s0)
    ctx = f"{up} {codon} {plus4}"
    if codon == "ATG":
        cls = classify(m3, plus4) + "  |  " + cavener(up, m3)
    else:
        cls = f"NOT AUG (start = {codon}) -> no functional Kozak; near-cognate initiation only"
    print(f"{name:38} {ctx:30} {m3:>3} {plus4:>3}  {cls}")
```
