## supplemental tables and scripts for "Long non-coding RNA-3′UTR interactions coordinate GABA signaling in the brain": startcons.py

```
#!/usr/bin/env python3
"""Conservation OF THE START CODONS THEMSELVES.
(a) map each candidate start to genomic coords, (b) per-base phyloP/phastCons at those 3 bases,
(c) check ortholog protein alignment at RE Met1 and at the residue matching RF's TTG start."""
import os, json, urllib.request

def get(url):
    req = urllib.request.Request(url, headers={"Content-Type": "application/json"})
    with urllib.request.urlopen(req, timeout=90) as r:
        return json.load(r)

def read_fasta(p):
    return "".join(l.strip() for l in open(p) if not l.startswith(">")).upper()

HERE = os.path.dirname(os.path.abspath(__file__))
D    = os.path.join(HERE, "data")
re_cdna = read_fasta(f"{D}/re_cdna.fa"); re_cds = read_fasta(f"{D}/re_cds.fa")
rf_cdna = read_fasta(f"{D}/rf_cdna.fa"); rf_cds = read_fasta(f"{D}/rf_cds.fa")

# ---- exon structures (dm6, + strand) ----
RE_EX = [(4087581,4087704),(4088960,4089214),(4090039,4090142),(4090273,4092706)]
RF_EX = [(4089131,4089214),(4090039,4090142),(4090273,4091302)]

def cdna_to_genomic(exons, pos1):
    """1-based cDNA position -> 1-based genomic (plus strand)."""
    rem = pos1
    for s,e in exons:
        L = e-s+1
        if rem <= L: return s+rem-1
        rem -= L
    raise ValueError("out of range")

cands = [
    ("RE used ATG (246 aa)",           RE_EX, re_cdna.find(re_cds)+1,  "ATG"),
    ("RF annotated start (171 aa)",    RF_EX, rf_cdna.find(rf_cds)+1,  "TTG"),
    ("RF ATG 115 aa (frame+2)",        RF_EX, 230, "ATG"),
    ("RF ATG 101 aa (frame+1)",        RF_EX, 250, "ATG"),
    ("RF ATG  65 aa (frame+3)",        RF_EX, 399, "ATG"),
    ("RF ATG  44 aa (frame+3)",        RF_EX, 150, "ATG"),
    ("RF first ATG (nt114, 7 aa)",     RF_EX, 114, "ATG"),
]

def ucsc_vals(track, start1, end1):
    url = (f"https://api.genome.ucsc.edu/getData/track?genome=dm6;track={track};"
           f"chrom=chr3L;start={start1-1};end={end1}")
    d = get(url); out=[]
    data = d.get(track, [])
    if isinstance(data, dict):
        data = sum((v for v in data.values() if isinstance(v,list)), [])
    for it in data:
        if isinstance(it,dict) and "value" in it:
            span = it.get("end", it["start"]+1)-it["start"]
            out.extend([it["value"]]*max(1,span))
    return out

print("=== (a,b) Genomic position and per-base conservation of each candidate start codon ===")
print(f"{'candidate':32} {'genomic (dm6)':24} {'codon':6} {'phyloP per base':26} {'mean':>6} {'phastCons':>10}")
print("-"*112)
rows=[]
for name, ex, cpos, expect in cands:
    g = cdna_to_genomic(ex, cpos)
    pp = ucsc_vals("phyloP124way", g, g+2)
    pc = ucsc_vals("phastCons124way", g, g+2)
    seq = (re_cdna if ex is RE_EX else rf_cdna)[cpos-1:cpos+2]
    mean = sum(pp)/len(pp) if pp else float('nan')
    mpc  = sum(pc)/len(pc) if pc else float('nan')
    pps = " ".join(f"{v:6.2f}" for v in pp)
    print(f"{name:32} chr3L:{g}-{g+2:<10} {seq:6} {pps:26} {mean:6.2f} {mpc:10.2f}")
    rows.append({"name":name,"genomic":g,"codon":seq,"phyloP":pp,"mean":mean,"phastCons":mpc})
json.dump(rows, open(f"{D}/startcons.json","w"))

# ---- (c) ortholog protein alignment at Met1 and at RE residue 76 ----
print("\n=== (c) Ortholog protein alignment: is RE Met1 conserved? what aligns to RF's TTG start (RE res.76)? ===")
h = get("https://rest.ensembl.org/homology/id/drosophila_melanogaster/FBgn0035495"
        "?content-type=application/json;type=orthologues;sequence=protein;aligned=1")
homs = h["data"][0]["homologies"]
print(f"orthologues returned: {len(homs)}")

met1 = {"M":0,"other":0}
res76 = {}
detail=[]
for hh in homs:
    src = hh["source"]["align_seq"]; tgt = hh["target"]["align_seq"]
    sp  = hh["target"]["species"]
    # walk source alignment to find ungapped source residue index 1 and 76
    idx=0; c1=c76=None
    for i,ch in enumerate(src):
        if ch!="-":
            idx+=1
            if idx==1: c1=i
            if idx==76: c76=i; break
    if c1 is None: continue
    t1  = tgt[c1]
    t76 = tgt[c76] if c76 is not None else "?"
    if t1=="M": met1["M"]+=1
    else: met1["other"]+=1
    res76[t76]=res76.get(t76,0)+1
    detail.append((sp,t1,t76))

n=len(detail)
print(f"\nRE Met1 (the used start): M in {met1['M']}/{n} orthologues "
      f"({100*met1['M']/n:.0f}%) -> {'CONSERVED' if met1['M']/n>0.8 else 'not well conserved'}")
print(f"\nResidue aligning to RF's TTG start (RE residue 76), across {n} orthologues:")
for aa,c in sorted(res76.items(), key=lambda kv:-kv[1]):
    lab = " <- Met (would allow AUG start)" if aa=="M" else ""
    print(f"   {aa}: {c:>3} ({100*c/n:.0f}%){lab}")
print("\nsample (species, res1, res76):")
for sp,a,b in detail[:12]:
    print(f"   {sp:52} {a}  {b}")
```
