## supplemental tables and scripts for "Long non-coding RNA-3′UTR interactions coordinate GABA signaling in the brain": utr_background.py

```
#!/usr/bin/env python3
"""How unusual is a 3'UTR as conserved as the CDS?
Empirical background: sample dm6 genes, compute mean phyloP over CDS vs 3'UTR, compare to CG14989."""
import json, os, random, urllib.request
from concurrent.futures import ThreadPoolExecutor

def get(url, tries=3):
    for t in range(tries):
        try:
            with urllib.request.urlopen(url, timeout=120) as r:
                return json.load(r)
        except Exception as e:
            if t == tries-1: raise
    return None

TRACK = "ncbiRefSeqCurated"

def genes_in(chrom, start, end):
    d = get(f"https://api.genome.ucsc.edu/getData/track?genome=dm6;track={TRACK};"
            f"chrom={chrom};start={start};end={end}")
    items = d.get(TRACK, [])
    if isinstance(items, dict):
        items = sum((v for v in items.values() if isinstance(v, list)), [])
    return items

def phylop_map(chrom, start0, end0):
    """dict genomic0 -> value"""
    d = get(f"https://api.genome.ucsc.edu/getData/track?genome=dm6;track=phyloP124way;"
            f"chrom={chrom};start={start0};end={end0}")
    data = d.get("phyloP124way", [])
    if isinstance(data, dict):
        data = sum((v for v in data.values() if isinstance(v, list)), [])
    m = {}
    for it in data:
        if isinstance(it, dict) and "value" in it:
            s = it["start"]; e = it.get("end", s+1)
            for p in range(s, e): m[p] = it["value"]
    return m

def exon_blocks(g):
    starts = [int(x) for x in str(g["exonStarts"]).strip(",").split(",") if x]
    ends   = [int(x) for x in str(g["exonEnds"]).strip(",").split(",") if x]
    return list(zip(starts, ends))

def analyse(g):
    try:
        chrom = g["chrom"]; cs = g["cdsStart"]; ce = g["cdsEnd"]
        if ce - cs < 300: return None
        blocks = exon_blocks(g)
        strand = g["strand"]
        cds_pos, utr3_pos = [], []
        for bs, be in blocks:
            for p in range(bs, be):
                if cs <= p < ce: cds_pos.append(p)
                elif (strand == "+" and p >= ce) or (strand == "-" and p < cs): utr3_pos.append(p)
        if len(cds_pos) < 300 or len(utr3_pos) < 200: return None
        span_s = min(min(cds_pos), min(utr3_pos)); span_e = max(max(cds_pos), max(utr3_pos))+1
        if span_e - span_s > 60000: return None
        pm = phylop_map(chrom, span_s, span_e)
        cv = [pm[p] for p in cds_pos if p in pm]
        uv = [pm[p] for p in utr3_pos if p in pm]
        if len(cv) < 300 or len(uv) < 200: return None
        return {"name": g.get("name2") or g.get("name"), "cds": sum(cv)/len(cv), "utr3": sum(uv)/len(uv),
                "n_cds": len(cv), "n_utr": len(uv)}
    except Exception:
        return None

random.seed(11)
print("collecting genes ...")
cands = []
for chrom, s, e in [("chr3L", 1_000_000, 6_000_000), ("chr2L", 1_000_000, 6_000_000),
                    ("chr2R", 10_000_000, 15_000_000), ("chr3R", 10_000_000, 15_000_000),
                    ("chrX", 1_000_000, 6_000_000)]:
    try: cands += genes_in(chrom, s, e)
    except Exception as ex: print("skip", chrom, ex)
# unique by name2
seen = set(); uniq = []
for g in cands:
    k = g.get("name2") or g.get("name")
    if k and k not in seen:
        seen.add(k); uniq.append(g)
random.shuffle(uniq)
print(f"candidate genes: {len(uniq)}; sampling ...")

res = []
with ThreadPoolExecutor(max_workers=6) as ex:
    for r in ex.map(analyse, uniq[:220]):
        if r: res.append(r)
        if len(res) >= 120: break

print(f"\nGenes analysed: {len(res)}")
ratio = [r for r in res if r["utr3"] >= r["cds"]]
print(f"3'UTR mean phyloP >= CDS mean phyloP: {len(ratio)}/{len(res)} = {100*len(ratio)/len(res):.1f}%")
import statistics as st
print(f"median CDS  phyloP: {st.median([r['cds'] for r in res]):.2f}")
print(f"median 3UTR phyloP: {st.median([r['utr3'] for r in res]):.2f}")
d = [r["utr3"]-r["cds"] for r in res]
print(f"median (3'UTR - CDS): {st.median(d):+.2f}")

CG_CDS, CG_UTR = 2.937, 2.194     # measured earlier for CG14989
print(f"\nCG14989: CDS {CG_CDS:.2f}, 3'UTR {CG_UTR:.2f}, diff {CG_UTR-CG_CDS:+.2f}")
pct = 100*sum(1 for x in d if x <= (CG_UTR-CG_CDS))/len(d)
print(f"CG14989's (3'UTR - CDS) sits at the {pct:.0f}th percentile of the background distribution")
above = 100*sum(1 for r in res if r['utr3'] >= CG_UTR)/len(res)
print(f"Genes whose 3'UTR is as conserved as CG14989's ({CG_UTR:.2f}): {above:.0f}%")
OUT = os.path.join(os.path.dirname(os.path.abspath(__file__)), "results")
os.makedirs(OUT, exist_ok=True)
json.dump(res, open(os.path.join(OUT, "utr_bg.json"), "w"))
```
