## supplemental tables and scripts for "Long non-coding RNA-3′UTR interactions coordinate GABA signaling in the brain": failure_rate.py

```
#!/usr/bin/env python3
"""SENSOR failure rate: one self-contained script, from masks to the reported table.

WHAT IT COMPUTES
    For each imaged region, the fraction of GABA-positive (Cyto+) neurons whose SENSOR
    signal fails to rise above the SENSOR level of the off cells immediately around them.

    Quoted against a 25% null: the bar is the 75th percentile of that region's OWN Cyto-
    cells, so a cell with no enrichment whatsoever clears it 25% of the time, in every
    region. The EXCESS over 25% is the signal; the rate is meaningless quoted alone.

THE FIVE STEPS, in order
    1. CLASSIFY    every nucleus as Cyto+ / Cyto- / borderline from the GABA channel alone.
    2. BASELINE    give every cell a LOCAL reference: the median SENSOR of its K nearest
                   Cyto- neighbours, in microns.  rel = sensor / local reference.
    3. Z-EDGE      drop cells too close to the first/last imaged plane (truncated nuclei).
    4. BAR         per region, the 75th percentile of the surviving Cyto- cells' rel.
    5. RATE        per region, the fraction of Cyto+ cells with rel <= that bar, + 95% CI.

    Steps 1-2 run on the WHOLE stack; steps 3-5 on the analysed region. That order is
    deliberate: the local reference should be built from every available off cell, not
    only those inside an arbitrary box.

    Nothing in steps 1 or 3 consults the SENSOR channel. The classification and the
    exclusions cannot see the measurement, so they cannot be tuned to it.

INPUTS
    manuscript/failure_rate_inputs.csv   one row per region: image and mask paths, voxel
                                         size, the analysed box, and the per-stack
                                         parameters that are NOT shared (see below).
    manuscript/patch_exclusions.csv      GABA mask objects confirmed BY EYE to be stain
                                         patches / debris rather than cells, one row per
                                         stack. This is the one human-judgement input;
                                         it was scored on the GABA channel, blind to SENSOR.

OUTPUTS
    manuscript/failure_rate.csv          the reported table (one row per region)
    manuscript/failure_rate_cells.csv    every scored cell, with rel, its local reference,
                                         and why it was excluded if it was. Every number in
                                         the table above is recomputable from this file.

USAGE
    python manuscript/failure_rate.py                 # write both outputs
    python manuscript/failure_rate.py --check         # also verify against the archived tables
    python manuscript/failure_rate.py --stacks i347s1 i348s2

REQUIREMENTS
    python >= 3.9, numpy, pandas, scipy, tifffile.  No project code is imported: this
    script does not depend on anything else in the repository.

UPSTREAM
    Nucleus / GABA / glia masks come from Cellpose-SAM (cellpose 4.0.9, do_3D, min_size 100,
    max_size_fraction 0.02) as run by the segmentation workflow; the per-channel parameters
    are recorded per region in the inputs file. Segmentation is NOT re-run here -- this
    script starts from the masks and the per-cell quantification they produced.
"""
from __future__ import annotations

import argparse
import sys
from pathlib import Path

import numpy as np
import pandas as pd
import tifffile
from scipy import ndimage as ndi
from scipy.spatial import cKDTree

HERE = Path(__file__).resolve().parent

# =============================================================================================
# PARAMETERS.  Every one of these is shared by all regions; anything that differs between
# regions is a COLUMN in the inputs file instead, so that no per-stack special case is hidden
# in the code.  The justification for each value is the comment beside it.
# =============================================================================================

POS_FRAC = 0.60
# Fraction of a NUCLEUS that must lie inside the GABA mask to call the cell Cyto+.
# The segmentation default is 0.10, which also admits nuclei that merely abut a GABA soma or
# touch one of its processes: one GABA object then yields several Cyto+ cells, most of them
# marginal, which dilutes the positive group. 0.60 requires the nucleus to sit inside the
# soma. Chosen on the GABA channel; the rank statistics are monotonic in this cut and so
# cannot be used to pick it.

BORDERLINE_OFFSET_DEFAULT = 5.0
# A mask-negative nucleus whose own nuclear GABA exceeds round(background) + offset is called
# BORDERLINE and excluded from both groups rather than counted as a negative. Grey levels, on
# the 8-bit-equivalent scale. Per-stack override in the inputs file (column
# `borderline_offset`): the offset tracks the antibody, not the analysis -- see i284s2.

DAPI_FLOOR_FRAC = 0.35
# EMPTY-MASK EXCLUSION, relative to each stack's own median nuclear DAPI. A minority of
# nucleus masks contain no nucleus: they are dim in DAPI (~0.22x the normal median), dim in
# GABA, have zero overlap with either marker mask, near-zero SENSOR max AND standard deviation
# (so not even noise structure), normal volume, and sit ~1.5x further from the field centre --
# masks drawn on empty space at the tissue margin. Applied on DAPI, the channel the masks were
# segmented from ("this nucleus mask has no nucleus in it"), never on SENSOR.

K = 10
# Number of nearest Cyto- cells forming a cell's local reference. Tuned on the NULL ONLY:
# Cyto- cells scored leave-one-out against their own K nearest Cyto-, where a perfect
# reference returns 1 plus noise. The spread of that null is minimised at K = 4-10 in every
# stack; K = 50 is 10-15% worse (null MAD 0.173 at K=10 vs 0.198 at K=50 in i347s1, median
# reach 6.9 vs 11.0 um). K = 10 rather than the numerical optimum K = 4 because the median of
# four cells is fragile. There is NO distance cutoff -- the neighbourhood is defined by count,
# and the distance it had to reach is reported per cell (`reach_um`) rather than used to
# filter. A reach limit was tested and rejected: it removes real cells in regions where off
# cells are genuinely sparse.

REF_FLOOR = 2.0
# Minimum SENSOR (8-bit-equivalent grey levels) for a Cyto- cell to ENTER the reference pool.
# Cells with essentially no SENSOR are not "off cells" but out-of-domain -- outside the
# sensor's expression territory, or outside the tissue. Left in the pool they drag the local
# reference toward zero wherever they cluster, and a ratio against a near-zero denominator
# explodes (115 objects had a reference below 2 with median SENSOR 2.2 but median rel 5.9,
# i.e. no signal scoring as strong enrichment). Excluded from the REFERENCE only: such cells
# are still scored themselves.

ZEDGE = 4
# Exclude cells whose centroid lies within this many z SLICES of the stack's first or last
# imaged plane. A nucleus intersecting the boundary is TRUNCATED: its SENSOR mean comes from a
# clipped sliver while its local reference is built from whole neighbours, so rel is pushed
# down for that cell alone. Systematic, not anecdotal -- median nucleus volume in the outer
# two slices is 12-15% below the interior in EVERY region. Applied to BOTH groups, with the
# bar recomputed from the surviving Cyto- cells: filtering only the failures would manufacture
# the result. It costs 7-19% of the counted cells, so every n is smaller and every confidence
# interval wider than in an uncorrected analysis. Every corrected rate falls inside its own
# uncorrected 95% CI. Set to 0 to disable.

BAR_PCTILE = 75
# The bar, as a percentile of each region's OWN Cyto- rel distribution. A percentile of the
# local background rather than a fixed rel fixes the FALSE-POSITIVE RATE at 25% everywhere: a
# fixed bar is not equally hard in every region, because the backgrounds differ in spread.
# 75 rather than 95 because the bar then also lands at nearly the same ABSOLUTE value in every
# region (rel 1.05-1.12), so the regions are held to nearly the same standard on both readings;
# at 95 the bars diverge (1.23 to 1.50) and part of any difference between regions would be the
# stricter bar rather than dimmer cells.

Z_CRIT = 1.96      # 95% Wilson score interval

INPUTS = HERE / "failure_rate_inputs.csv"
PATCHES = HERE / "patch_exclusions.csv"


# =============================================================================================
# STEP 1 -- CLASSIFY
# =============================================================================================

def classify(cfg: pd.Series, patch_labels: set[int]) -> pd.DataFrame:
    """One row per nucleus: group, SENSOR, and the values every call was made from.

    Cyto+       >= POS_FRAC of the nucleus lies inside the (patch-cleaned) GABA mask
    borderline  mask-negative, but nuclear GABA > round(background) + offset   -> excluded
    Cyto-       mask-negative, and nuclear GABA <= that cut

    Glia-positive nuclei are dropped first where a glia counterstain exists; empty masks are
    dropped on DAPI. Grey levels are put on a common 8-bit-equivalent scale by the divisors in
    the inputs file, because the borderline cut is an ABSOLUTE level and the acquisitions are
    not all the same bit depth.
    """
    nuc = tifffile.imread(cfg.nucleus_mask)
    gaba_mask = tifffile.imread(cfg.gaba_mask)
    dapi = tifffile.imread(cfg.dapi_image).astype(np.float32) / cfg.dapi_divisor
    gaba = tifffile.imread(cfg.gaba_image).astype(np.float32) / cfg.gaba_divisor
    quant = pd.read_csv(cfg.quantification)

    # Confirmed stain patches are removed from the GABA mask BEFORE anything is measured, so
    # Cyto+ status is decided by the cleaned mask rather than corrected afterwards.
    clean = (gaba_mask > 0)
    if patch_labels:
        clean &= ~np.isin(gaba_mask, sorted(patch_labels))

    labels = np.arange(1, int(nuc.max()) + 1)
    nuclear_gaba = ndi.mean(gaba, nuc, labels)                      # per-nucleus GABA
    nuclear_dapi = ndi.mean(dapi, nuc, labels)                      # per-nucleus DAPI
    overlap = ndi.mean(clean.astype(np.float32), nuc, labels)       # fraction inside GABA mask

    # Background is the GABA level OUTSIDE every nucleus mask, read per stack from the image.
    background = float(np.median(gaba[nuc == 0]))
    cut = round(background) + float(cfg.borderline_offset)

    q = quant.set_index("cell_label").reindex(labels)
    sensor = q.ch4_mean_intensity.values / cfg.sensor_divisor
    if cfg.has_glia:
        glia = q.nuclear_marker_positive.astype(bool).values
    else:
        glia = np.zeros(len(labels), bool)      # no counterstain: the exclusion cannot be made
    empty = nuclear_dapi < DAPI_FLOOR_FRAC * np.median(nuclear_dapi)

    group = np.where(overlap >= POS_FRAC, "Cyto+",
                     np.where(nuclear_gaba <= cut, "Cyto-", "borderline"))

    out = pd.DataFrame(dict(
        stack=cfg.stack, cell_label=labels, group=group,
        sensor=sensor, nuclear_gaba=nuclear_gaba, gaba_overlap_frac=overlap.round(4),
        excluded=np.where(glia, "glia", np.where(empty, "empty_mask", "")),
    ))
    out.attrs["background"] = background
    out.attrs["cut"] = cut
    return out


# =============================================================================================
# STEP 2 -- LOCAL BASELINE
# =============================================================================================

def local_reference(query_um, pool_um, pool_sensor, drop_self, k=None):
    """Median SENSOR of the K nearest reference cells, and the distance that reach required.

    Distances are EUCLIDEAN IN MICRONS between mask centroids, not in voxels: z is 2-4x
    coarser than xy in these acquisitions, so a voxel-space metric would draw a neighbourhood
    several times taller in z than wide in xy.

    `k` overrides the module-level K. It exists for local_baseline.py's K sweep, which rescores
    the NULL at other neighbourhood sizes; the analysis itself always runs at K.
    """
    K_ = K if k is None else k
    tree = cKDTree(pool_um)
    k = min(K_ + 1 if drop_self else K_, len(pool_um))
    dist, idx = tree.query(query_um, k=k)
    idx, dist = np.atleast_2d(idx), np.atleast_2d(dist)
    if drop_self:
        idx, dist = idx[:, 1:], dist[:, 1:]     # the nearest hit is the cell itself
    vals = pool_sensor[idx]
    return np.median(vals, axis=1), dist[:, -1]


def add_local_baseline(cells: pd.DataFrame, cfg: pd.Series) -> pd.DataFrame:
    """Attach local_ref, rel and reach_um using every off cell in the WHOLE stack."""
    quant = pd.read_csv(cfg.quantification)[
        ["cell_label", "centroid_x", "centroid_y", "centroid_z"]]
    cells = cells.merge(quant, on="cell_label", how="left", validate="one_to_one")
    if cells[["centroid_x", "centroid_y", "centroid_z"]].isna().any().any():
        raise SystemExit(f"{cfg.stack}: nuclei missing from the quantification table")

    um = np.column_stack([cells.centroid_x * cfg.pixel_um,
                          cells.centroid_y * cfg.pixel_um,
                          cells.centroid_z * cfg.z_step_um])

    # The reference pool: Cyto- cells that survived every exclusion and clear REF_FLOOR.
    pool = ((cells.group == "Cyto-") & (cells.excluded == "") & (cells.sensor >= REF_FLOOR)).values
    if pool.sum() <= K:
        raise SystemExit(f"{cfg.stack}: only {pool.sum()} reference cells, need > {K}")
    pool_um, pool_sensor = um[pool], cells.sensor.values[pool]

    ref, reach = local_reference(um, pool_um, pool_sensor, drop_self=False)
    # A reference cell must not anchor on itself: rescore the pool leave-one-out.
    ref_loo, reach_loo = local_reference(pool_um, pool_um, pool_sensor, drop_self=True)
    ref[pool], reach[pool] = ref_loo, reach_loo

    cells["local_ref"] = ref.round(3)
    cells["rel"] = (cells.sensor / ref).round(4)
    cells["reach_um"] = reach.round(2)
    cells["in_reference_pool"] = pool
    return cells


# =============================================================================================
# STEPS 3-5 -- REGION, Z-EDGE, BAR, RATE
# =============================================================================================

def wilson(k: int, n: int) -> tuple[float, float]:
    """95% Wilson score interval, in percent. Correct at the small counts and rates here,
    where the normal approximation is not (it can put the lower bound below zero)."""
    if n == 0:
        return float("nan"), float("nan")
    p, d = k / n, 1 + Z_CRIT ** 2 / n
    centre = (p + Z_CRIT ** 2 / (2 * n)) / d
    half = Z_CRIT * np.sqrt(p * (1 - p) / n + Z_CRIT ** 2 / (4 * n ** 2)) / d
    return 100 * (centre - half), 100 * (centre + half)


def select_region(cells: pd.DataFrame, cfg: pd.Series) -> pd.DataFrame:
    """Mark which cells the rate is computed from, and why each of the others is out.

    The z extent comes from the WHOLE stack, not from the box: the analysed region is an xy
    box, so the stack's own first and last planes are the truncation boundary wherever it sits.
    """
    inside = ((cells.centroid_x >= cfg.x_min) & (cells.centroid_x <= cfg.x_max) &
              (cells.centroid_y >= cfg.y_min) & (cells.centroid_y <= cfg.y_max))
    z_lo, z_hi = cells.centroid_z.min(), cells.centroid_z.max()
    z_ok = np.minimum(cells.centroid_z - z_lo, z_hi - cells.centroid_z) >= ZEDGE

    reason = np.where(cells.excluded != "", cells.excluded,
              np.where(cells.group == "borderline", "borderline",
              np.where(~inside, "outside_region",
              np.where(~z_ok, "z_edge", ""))))
    # "scored" rather than an empty string, so the reason column survives a CSV round-trip
    # as a value instead of coming back as a missing one.
    cells["excluded"] = np.where(reason == "", "scored", reason)
    cells["scored"] = reason == ""
    return cells


def rate_for(cells: pd.DataFrame, cfg: pd.Series) -> dict:
    scored = cells[cells.scored]
    neg = scored[scored.group == "Cyto-"]
    pos = scored[scored.group == "Cyto+"]
    bar = float(np.percentile(neg.rel.values, BAR_PCTILE))
    fail = pos.rel <= bar
    n, k = len(pos), int(fail.sum())
    lo, hi = wilson(k, n)
    return dict(
        stack=cfg.stack, n_cytopos=n, n_cytoneg=len(neg),
        bar_rel=round(bar, 4), n_failed=k,
        failure_rate_pct=round(100 * k / n, 1) if n else float("nan"),
        ci_lo_pct=round(lo, 1), ci_hi_pct=round(hi, 1),
        null_rate_pct=100 - BAR_PCTILE,
        median_rel_cytopos=round(float(pos.rel.median()), 2),
        reach_um_p50=round(float(np.median(scored.reach_um)), 1),
        reach_um_p95=round(float(np.percentile(scored.reach_um, 95)), 1),
        gaba_background=cells.attrs.get("background"), borderline_cut=cells.attrs.get("cut"),
        n_dropped_zedge=int((cells.excluded == "z_edge").sum()),
        n_dropped_glia=int((cells.excluded == "glia").sum()),
        n_dropped_empty=int((cells.excluded == "empty_mask").sum()),
        n_borderline=int((cells.excluded == "borderline").sum()),
    )


# =============================================================================================
# DRIVER
# =============================================================================================

def read_patches() -> dict:
    if not PATCHES.exists():
        return {}
    tab = pd.read_csv(PATCHES).fillna({"labels": ""})
    return {r.stack: {int(v) for v in str(r.labels).split()} for r in tab.itertuples()}


def main(argv=None):
    ap = argparse.ArgumentParser(description=__doc__,
                                 formatter_class=argparse.RawDescriptionHelpFormatter)
    ap.add_argument("--stacks", nargs="+", help="restrict to these regions")
    ap.add_argument("--check", action="store_true",
                    help="compare the result against the archived per-cell tables")
    args = ap.parse_args(argv)

    cfgs = pd.read_csv(INPUTS)
    cfgs["borderline_offset"] = cfgs["borderline_offset"].fillna(BORDERLINE_OFFSET_DEFAULT)
    if args.stacks:
        missing = set(args.stacks) - set(cfgs["stack"])
        if missing:
            raise SystemExit(f"not in {INPUTS.name}: {sorted(missing)}")
        cfgs = cfgs[cfgs["stack"].isin(args.stacks)]

    patches = read_patches()
    rows, per_cell = [], []
    for cfg in cfgs.itertuples(index=False):
        cfg = pd.Series(cfg._asdict())
        cells = classify(cfg, patches.get(cfg.stack, set()))
        attrs = cells.attrs
        cells = add_local_baseline(cells, cfg)
        cells.attrs = attrs
        cells = select_region(cells, cfg)
        row = rate_for(cells, cfg)
        rows.append(row)
        per_cell.append(cells)
        print(f"{cfg.stack}: {row['n_failed']}/{row['n_cytopos']} Cyto+ at or below the bar "
              f"(rel <= {row['bar_rel']}) = {row['failure_rate_pct']}% "
              f"[{row['ci_lo_pct']}-{row['ci_hi_pct']}], null {row['null_rate_pct']}%")

    table = pd.DataFrame(rows).sort_values("failure_rate_pct")
    cells = pd.concat(per_cell, ignore_index=True)
    table.to_csv(HERE / "failure_rate.csv", index=False)
    cells.to_csv(HERE / "failure_rate_cells.csv", index=False)
    print(f"\n{table.to_string(index=False)}")
    print(f"\nwrote {HERE/'failure_rate.csv'} and {HERE/'failure_rate_cells.csv'}")

    if args.check:
        return check(cells, table)
    return 0


def check(cells: pd.DataFrame, table: pd.DataFrame) -> int:
    """Verify against the archived per-cell tables from the original pipeline.

    Not a unit test: it re-derives the published numbers from the original pipeline's own
    intermediate files and requires them to agree cell by cell.
    """
    root = HERE.parent
    ref = pd.concat([pd.read_csv(p) for p in sorted(
        root.glob("analysis/sensor/local_baseline_nuclei*.csv")) if "_ov" not in p.name])
    ref = ref.drop_duplicates(["stack", "cell_label"])
    ok = True
    for stack, mine in cells.groupby("stack"):
        theirs = ref[ref["stack"] == stack]
        if theirs.empty:
            print(f"  {stack}: no archived table, skipped")
            continue
        m = mine.merge(theirs, on=["stack", "cell_label"], suffixes=("", "_ref"))
        if len(m) != len(theirs):
            print(f"  {stack}: FAIL {len(theirs)} archived cells, {len(m)} matched")
            ok = False
            continue
        bad_group = (m.group != m.group_ref).sum()
        drel = (m.rel - m.rel_ref).abs()
        worst = float(drel.max())
        status = "ok" if bad_group == 0 and worst < 1e-3 else "FAIL"
        ok &= status == "ok"
        print(f"  {stack}: {status}  n={len(m)}  group mismatches={bad_group}  "
              f"max |d rel|={worst:.2e}")
    print("CHECK PASSED" if ok else "CHECK FAILED")
    return 0 if ok else 1


if __name__ == "__main__":
    sys.exit(main())
```
