## supplemental tables and scripts for "Long non-coding RNA-3′UTR interactions coordinate GABA signaling in the brain": local_baseline.py

```
#!/usr/bin/env python3
"""Local baseline analysis: score every cell's SENSOR against the off cells around it.

WHY A LOCAL BASELINE
    SENSOR expression varies smoothly across the tissue, so one stack-wide reference mixes
    territories with different floors: the same cell reads as bright or as ordinary depending
    only on where the reference was drawn from. The gradient is measured here and reported per
    region (`gradient_p90_p10`, the ratio of the 90th to the 10th percentile of the local
    reference) rather than assumed.

    So each cell is referenced to the cells beside it: local_ref = the median SENSOR of its
    K nearest Cyto- (off) cells, and rel = sensor / local_ref. rel = 1 means a cell is
    indistinguishable from the off cells immediately around it.

WHAT IT COMPUTES
    1. CLASSIFY   every nucleus as Cyto+ / Cyto- / borderline from the GABA channel alone.
    2. BASELINE   local_ref, rel, and the distance the reference had to reach, per cell.
    3. DIAGNOSE   per region: the reach the neighbourhood required, how far the reference
                  varies across the field, and the effect size computed BOTH ways -- on raw
                  SENSOR (r_global) and on rel (r_local) -- so the local reference can be seen
                  to sharpen rather than manufacture the separation.
    4. SWEEP      (--k-sweep) the evidence for K, scored on the NULL ONLY.

THE NULL, AND HOW K WAS CHOSEN
    Cyto- cells are scored leave-one-out against their own K nearest Cyto-. A perfect local
    reference returns rel = 1 plus noise for them, so the SPREAD of that null -- reported as
    its median absolute deviation -- measures how well the neighbourhood tracks the tissue:
    too large a K reaches across the gradient, too small a K takes the median of too few
    cells. The Cyto+ group plays no part in the choice, and `--k-sweep` reprints that evidence
    rather than asking the reader to take K = 10 on trust.

INPUTS
    manuscript/failure_rate_inputs.csv   one row per region: image and mask paths, voxel
                                         size, the analysed box, and the per-stack parameters
                                         that are NOT shared between regions.
    manuscript/patch_exclusions.csv      GABA mask objects confirmed BY EYE to be stain
                                         patches / debris rather than cells.

OUTPUTS
    manuscript/local_baseline_cells.csv    every nucleus: group, sensor, local_ref, rel,
                                           reach_um, its percentile among its own neighbours,
                                           and its exclusion reason if it was not scored.
    manuscript/local_baseline_summary.csv  the per-region table.
    manuscript/local_baseline_k_sweep.csv  only with --k-sweep: the null spread against K.

USAGE
    python manuscript/local_baseline.py                 # write both outputs
    python manuscript/local_baseline.py --k-sweep       # also write the K evidence
    python manuscript/local_baseline.py --stacks i347s1 i348s2

REQUIREMENTS
    python >= 3.9, numpy, pandas, scipy, tifffile.  No project code outside this directory is
    imported. Classification, the local-reference routine and the shared parameters live in
    failure_rate.py, which sits beside this file, so all three manuscript scripts score one
    set of cells with one implementation rather than three.
"""
from __future__ import annotations

import argparse
import sys
from pathlib import Path

import numpy as np
import pandas as pd
from scipy.spatial import cKDTree
from scipy.stats import mannwhitneyu

from failure_rate import (INPUTS, K, REF_FLOOR, BORDERLINE_OFFSET_DEFAULT,
                          add_local_baseline, classify, local_reference,
                          read_patches, select_region)

HERE = Path(__file__).resolve().parent

K_SWEEP = (4, 6, 10, 15, 25, 50)
# The K values swept by --k-sweep. K = 10 is the analysis value (defined, with its
# justification, in failure_rate.py); 50 is the value this analysis originally used before the
# null was examined, kept in the sweep so that the change is auditable rather than asserted.


def rank_biserial(pos: np.ndarray, neg: np.ndarray) -> float:
    """2*(U / (n_pos * n_neg)) - 1: a linear rescaling of Mann-Whitney U, so (r + 1) / 2 is
    the probability that a random Cyto+ cell exceeds a random Cyto- one."""
    if len(pos) == 0 or len(neg) == 0:
        return float("nan")
    U = mannwhitneyu(pos, neg, alternative="two-sided").statistic
    return 2 * (U / (len(pos) * len(neg))) - 1


def local_percentile(cells: pd.DataFrame, cfg: pd.Series) -> np.ndarray:
    """Percentile of each cell's SENSOR among its K nearest reference cells.

    The rank counterpart of `rel`: it says the same thing without a ratio, and unlike a ratio
    it cannot be inflated by a small denominator. Reported per cell; no rule uses it.
    """
    um = np.column_stack([cells.centroid_x * cfg.pixel_um,
                          cells.centroid_y * cfg.pixel_um,
                          cells.centroid_z * cfg.z_step_um])
    pool = cells.in_reference_pool.values
    pool_um, pool_sensor = um[pool], cells.sensor.values[pool]
    idx = np.atleast_2d(cKDTree(pool_um).query(um, k=min(K, len(pool_um)))[1])
    return 100.0 * (pool_sensor[idx] < cells.sensor.values[:, None]).mean(axis=1)


def summarise(cells: pd.DataFrame, cfg: pd.Series) -> dict:
    """One row per region. The scored set is the same one the failure rate uses: inside the
    analysed box, not z-truncated, not glia, not an empty mask, not borderline."""
    scored = cells[cells.scored]
    pos = scored[scored.group == "Cyto+"]
    neg = scored[scored.group == "Cyto-"]
    # Every reported quantity is over the SCORED cells, i.e. the cropped region alone -- the
    # same set the figures show. (The reference POOL behind local_ref is still drawn from the
    # whole stack, deliberately: an off cell just outside the box is still the right neighbour
    # for a cell just inside it. What is reported, and what is scored, is the region.)
    ref = scored.local_ref.values
    return dict(
        stack=cfg.stack, experiment=cfg.experiment, K=K,
        n_cytopos=len(pos), n_cytoneg=len(neg),
        n_reference_pool=int(cells.in_reference_pool.sum()),
        n_below_ref_floor=int(((cells.group == "Cyto-") & (cells.excluded == "scored")
                               & (cells.sensor < REF_FLOOR)).sum()),
        ref_floor=REF_FLOOR,
        # How far "local" actually had to reach. Reported, never used to filter: a reach limit
        # was tested and rejected, because it deletes real cells wherever off cells are sparse.
        reach_um_p50=round(float(np.median(scored.reach_um)), 1),
        reach_um_p95=round(float(np.percentile(scored.reach_um, 95)), 1),
        reach_um_max=round(float(scored.reach_um.max()), 1),
        # The gradient the local reference exists to absorb, across the analysed region.
        local_ref_p10=round(float(np.percentile(ref, 10)), 2),
        local_ref_p90=round(float(np.percentile(ref, 90)), 2),
        gradient_p90_p10=round(float(np.percentile(ref, 90) / np.percentile(ref, 10)), 2),
        # The same effect size before and after the local reference is applied.
        r_global=round(rank_biserial(pos.sensor.values, neg.sensor.values), 3),
        r_local=round(rank_biserial(pos.rel.values, neg.rel.values), 3),
        median_rel_cytopos=round(float(pos.rel.median()), 3),
        # The null: Cyto- cells scored leave-one-out against their own neighbours. A centre at
        # ~1 with a small spread means the reference tracks the tissue. This is the quantity
        # K was chosen on, and it involves no Cyto+ cell. Taken over every scored Cyto- cell;
        # --k-sweep recomputes it over the reference pool alone, so the two differ by the few
        # scored negatives that sit below REF_FLOOR and are therefore not in the pool.
        null_median_rel=round(float(neg.rel.median()), 3),
        null_mad_rel=round(float((neg.rel - neg.rel.median()).abs().median()), 4),
    )


def null_mad(cells: pd.DataFrame, cfg: pd.Series, k: int) -> dict:
    """Re-score the reference pool leave-one-out at a different K and report the null spread.

    Only Cyto- cells are involved, so nothing in this sweep can be tuned to the result.
    """
    um = np.column_stack([cells.centroid_x * cfg.pixel_um,
                          cells.centroid_y * cfg.pixel_um,
                          cells.centroid_z * cfg.z_step_um])
    pool = cells.in_reference_pool.values & cells.scored.values
    if pool.sum() <= k:
        return dict(stack=cfg.stack, K=k, n=int(pool.sum()))
    pool_um, pool_sensor = um[pool], cells.sensor.values[pool]
    ref, reach = local_reference(pool_um, pool_um, pool_sensor, drop_self=True, k=k)
    rel = pool_sensor / ref
    return dict(stack=cfg.stack, K=k, n=int(pool.sum()),
                null_median_rel=round(float(np.median(rel)), 4),
                null_mad_rel=round(float(np.median(np.abs(rel - np.median(rel)))), 4),
                reach_um_p50=round(float(np.median(reach)), 1))


def main(argv=None):
    ap = argparse.ArgumentParser(description=__doc__,
                                 formatter_class=argparse.RawDescriptionHelpFormatter)
    ap.add_argument("--stacks", nargs="+", help="restrict to these regions")
    ap.add_argument("--k-sweep", action="store_true",
                    help="also write the null spread against K, the evidence for K = %d" % K)
    args = ap.parse_args(argv)

    cfgs = pd.read_csv(INPUTS)
    cfgs["borderline_offset"] = cfgs["borderline_offset"].fillna(BORDERLINE_OFFSET_DEFAULT)
    if args.stacks:
        missing = set(args.stacks) - set(cfgs["stack"])
        if missing:
            raise SystemExit(f"not in {INPUTS.name}: {sorted(missing)}")
        cfgs = cfgs[cfgs["stack"].isin(args.stacks)]

    patches = read_patches()
    rows, per_cell, sweep = [], [], []
    for cfg in cfgs.itertuples(index=False):
        cfg = pd.Series(cfg._asdict())
        cells = classify(cfg, patches.get(cfg.stack, set()))
        cells = add_local_baseline(cells, cfg)      # whole stack: every off cell is available
        cells = select_region(cells, cfg)           # analysed box + z-edge exclusion
        cells["local_pct"] = local_percentile(cells, cfg).round(1)
        row = summarise(cells, cfg)
        rows.append(row)
        per_cell.append(cells)
        if args.k_sweep:
            sweep += [null_mad(cells, cfg, k) for k in K_SWEEP]
        print(f"{cfg.stack}: local reference varies {row['gradient_p90_p10']}x across the "
              f"field, reach {row['reach_um_p50']} um (p95 {row['reach_um_p95']});  "
              f"r {row['r_global']} global -> {row['r_local']} local;  "
              f"null {row['null_median_rel']} +/- {row['null_mad_rel']}")

    table = pd.DataFrame(rows).sort_values("stack")
    cells = pd.concat(per_cell, ignore_index=True)
    table.to_csv(HERE / "local_baseline_summary.csv", index=False)
    cells.to_csv(HERE / "local_baseline_cells.csv", index=False)
    pd.set_option("display.width", 220)
    print(f"\n{table.to_string(index=False)}")
    print(f"\nwrote {HERE/'local_baseline_summary.csv'} and {HERE/'local_baseline_cells.csv'}")

    if args.k_sweep:
        sw = pd.DataFrame(sweep)
        sw.to_csv(HERE / "local_baseline_k_sweep.csv", index=False)
        print("\n=== null spread (MAD of rel) against K, Cyto- cells only, leave-one-out ===")
        print(sw.pivot(index="stack", columns="K", values="null_mad_rel").to_string())
        print(f"\nwrote {HERE/'local_baseline_k_sweep.csv'}")
    return 0


if __name__ == "__main__":
    sys.exit(main())
```
