## supplemental tables and scripts for "Long non-coding RNA-3′UTR interactions coordinate GABA signaling in the brain": quantify_cells.py

```
#!/usr/bin/env python3
"""Per-cell quantification and statistics: from the segmentation masks to the reported
Cyto+ vs Cyto- SENSOR comparison.

WHAT IT COMPUTES
    One row per nucleus -- its SENSOR level, its nuclear GABA level, how much of it lies
    inside the GABA mask, its size and position -- and, per imaged region, the statistics
    quoted for the comparison between GABA-positive (Cyto+) and GABA-negative (Cyto-) cells:
    group medians, the fold change between them, a two-sided Mann-Whitney U test, and the
    rank-biserial correlation r with its probability-of-superiority reading.

    r (equivalently, the probability that a randomly chosen Cyto+ cell is brighter than a
    randomly chosen Cyto- cell) is the effect size that carries the claim. The p value is
    reported for completeness only: at these n, significance is not in question and it is
    not evidence of a large difference.

THE STEPS, in order
    1. QUANTIFY   per-nucleus SENSOR, GABA, DAPI, GABA-mask overlap, volume and centroid.
    2. CLASSIFY   every nucleus as Cyto+ / Cyto- / borderline from the GABA channel alone.
    3. SELECT     restrict to the analysed region and drop z-truncated nuclei.
    4. COMPARE    per region, Cyto+ vs Cyto- SENSOR: medians, fold change, Mann-Whitney, r.

    Steps 1-2 run on the WHOLE stack; steps 3-4 on the analysed region, and the reported
    table is the analysed region ALONE -- the same box the figures are drawn from
    (`x_min`..`y_max`, echoed into the output table so it declares its own scope). It is not
    a whole-field measurement: in some stacks the field of view extends past the optic lobe,
    and the discrete bright cell clusters outside it inflate r without saying anything about
    the tissue under study. Whole-field values are deliberately NOT reported here; the
    per-cell output marks every cell it excluded and why, so any other scope is recomputable
    from it, but nothing quoted mixes the two.

    Nothing in steps 1-3 consults the SENSOR channel. The classification and the exclusions
    cannot see the measurement, so they cannot be tuned to it.

INPUTS
    manuscript/failure_rate_inputs.csv   one row per region: image and mask paths, voxel
                                         size, the analysed box, and the per-stack
                                         parameters that are NOT shared between regions.
    manuscript/patch_exclusions.csv      GABA mask objects confirmed BY EYE to be stain
                                         patches / debris rather than cells. This is the one
                                         human-judgement input; it was scored on the GABA
                                         channel, blind to SENSOR.

OUTPUTS
    manuscript/quantification_cells.csv  every nucleus, with its measurements, its group,
                                         and its exclusion reason if it was not scored.
    manuscript/quantification_stats.csv  the reported per-region table.

USAGE
    python manuscript/quantify_cells.py                  # write both outputs
    python manuscript/quantify_cells.py --stacks i347s1 i348s2

REQUIREMENTS
    python >= 3.9, numpy, pandas, scipy, tifffile.  No project code outside this directory
    is imported. The classification and the shared parameters live in failure_rate.py, which
    sits beside this file: the two scripts classify cells with one implementation rather than
    two, so the statistics below and the failure rate are computed on the same cells.

UPSTREAM
    Nucleus / GABA / glia masks come from Cellpose-SAM (cellpose 4.0.9, do_3D, min_size 100,
    max_size_fraction 0.02) as run by the segmentation workflow; the per-channel parameters
    are recorded per region in the inputs file. Segmentation is NOT re-run here.
"""
from __future__ import annotations

import argparse
import sys
from pathlib import Path

import numpy as np
import pandas as pd
from scipy.stats import mannwhitneyu

from failure_rate import (INPUTS, BORDERLINE_OFFSET_DEFAULT,
                          classify, read_patches, select_region)

HERE = Path(__file__).resolve().parent

# The parameters this script needs are the classification's, and they are defined once, in
# failure_rate.py: POS_FRAC, BORDERLINE_OFFSET_DEFAULT, DAPI_FLOOR_FRAC, ZEDGE. Their
# justifications are the comments beside them there. The only choice made here is which cells
# the statistics are computed over, and that is the same scored set the failure rate uses:
# inside the analysed region, not z-truncated, not glia, not an empty mask, not borderline.


# =============================================================================================
# STEP 1 -- PER-CELL QUANTIFICATION
# =============================================================================================

def measure(cells: pd.DataFrame, cfg: pd.Series) -> pd.DataFrame:
    """Attach size and position to the classified nuclei, in physical units.

    Volume is converted to cubic microns and the centroid to microns because the voxel is
    anisotropic and not the same size in every acquisition: 0.14-0.20 um in xy against a
    0.5 or 1.0 um z step. Voxel counts are not comparable between regions; cubic microns are.
    """
    quant = pd.read_csv(cfg.quantification)[
        ["cell_label", "centroid_x", "centroid_y", "centroid_z", "volume_voxels",
         "ch4_max_intensity", "ch4_std_intensity"]]
    cells = cells.merge(quant, on="cell_label", how="left", validate="one_to_one")
    if cells[["centroid_x", "centroid_y", "centroid_z"]].isna().any().any():
        raise SystemExit(f"{cfg.stack}: nuclei missing from the quantification table")

    voxel_um3 = cfg.pixel_um ** 2 * cfg.z_step_um
    cells["volume_um3"] = (cells.volume_voxels * voxel_um3).round(2)
    cells["x_um"] = (cells.centroid_x * cfg.pixel_um).round(2)
    cells["y_um"] = (cells.centroid_y * cfg.pixel_um).round(2)
    cells["z_um"] = (cells.centroid_z * cfg.z_step_um).round(2)
    # SENSOR max and std are on the raw scale in the pipeline table; put them on the same
    # 8-bit-equivalent scale as `sensor`, or a uint16 stack's values are 16x the others'.
    cells["sensor_max"] = (cells.ch4_max_intensity / cfg.sensor_divisor).round(3)
    cells["sensor_std"] = (cells.ch4_std_intensity / cfg.sensor_divisor).round(3)
    return cells.drop(columns=["ch4_max_intensity", "ch4_std_intensity"])


# =============================================================================================
# STEP 4 -- THE COMPARISON
# =============================================================================================

def compare(pos: np.ndarray, neg: np.ndarray) -> dict:
    """Cyto+ vs Cyto- SENSOR: medians, fold change, Mann-Whitney U, rank-biserial r.

    r = 2*(U / (n_pos * n_neg)) - 1, the rank-biserial correlation. It is a linear rescaling
    of U, so it needs no distributional assumption, and (r + 1) / 2 is the probability that a
    randomly drawn Cyto+ cell exceeds a randomly drawn Cyto- cell -- the form quoted in the
    text, because it is readable without knowing what a rank statistic is.

    The medians are reported rather than means: both distributions are right-skewed, and the
    Cyto+ mean is set by a handful of very bright cells.
    """
    if len(pos) == 0 or len(neg) == 0:
        return dict(n_cytopos=len(pos), n_cytoneg=len(neg))
    U = mannwhitneyu(pos, neg, alternative="two-sided")
    r = 2 * (U.statistic / (len(pos) * len(neg))) - 1
    med_pos, med_neg = float(np.median(pos)), float(np.median(neg))
    return dict(
        n_cytopos=len(pos), n_cytoneg=len(neg),
        median_sensor_cytopos=round(med_pos, 2), median_sensor_cytoneg=round(med_neg, 2),
        fold_change=round(med_pos / med_neg, 3) if med_neg else float("nan"),
        mannwhitney_u=float(U.statistic), p_value=float(U.pvalue),
        rank_biserial_r=round(r, 3), prob_superiority_pct=round(100 * (r + 1) / 2, 1),
    )


def stats_for(cells: pd.DataFrame, cfg: pd.Series) -> dict:
    """One row per region, computed over the SCORED cells only -- i.e. inside the analysed
    box, not z-truncated, not glia, not an empty mask, not borderline. That is the same set
    the figures show and the same set the failure rate is computed on."""
    scored = cells[cells.scored]
    sensor = lambda d, g: d.loc[d.group == g, "sensor"].values          # noqa: E731

    row = dict(stack=cfg.stack, experiment=cfg.experiment,
               x_min=cfg.x_min, x_max=cfg.x_max, y_min=cfg.y_min, y_max=cfg.y_max)
    row.update(compare(sensor(scored, "Cyto+"), sensor(scored, "Cyto-")))
    n_pos, n_neg = row["n_cytopos"], row["n_cytoneg"]
    row.update(
        cytopos_pct_of_scored=round(100 * n_pos / (n_pos + n_neg), 1) if n_pos + n_neg else float("nan"),
        median_nuclear_gaba_cytopos=round(float(scored.loc[scored.group == "Cyto+", "nuclear_gaba"].median()), 2),
        median_nuclear_gaba_cytoneg=round(float(scored.loc[scored.group == "Cyto-", "nuclear_gaba"].median()), 2),
        median_volume_um3_cytopos=round(float(scored.loc[scored.group == "Cyto+", "volume_um3"].median()), 1),
        median_volume_um3_cytoneg=round(float(scored.loc[scored.group == "Cyto-", "volume_um3"].median()), 1),
        gaba_background=cells.attrs.get("background"), borderline_cut=cells.attrs.get("cut"),
        n_nuclei_total=len(cells),
        n_borderline=int((cells.excluded == "borderline").sum()),
        n_dropped_glia=int((cells.excluded == "glia").sum()),
        n_dropped_empty=int((cells.excluded == "empty_mask").sum()),
        n_dropped_zedge=int((cells.excluded == "z_edge").sum()),
        n_outside_region=int((cells.excluded == "outside_region").sum()),
    )
    return row


# =============================================================================================
# DRIVER
# =============================================================================================

def main(argv=None):
    ap = argparse.ArgumentParser(description=__doc__,
                                 formatter_class=argparse.RawDescriptionHelpFormatter)
    ap.add_argument("--stacks", nargs="+", help="restrict to these regions")
    args = ap.parse_args(argv)

    cfgs = pd.read_csv(INPUTS)
    cfgs["borderline_offset"] = cfgs["borderline_offset"].fillna(BORDERLINE_OFFSET_DEFAULT)
    if args.stacks:
        missing = set(args.stacks) - set(cfgs["stack"])
        if missing:
            raise SystemExit(f"not in {INPUTS.name}: {sorted(missing)}")
        cfgs = cfgs[cfgs["stack"].isin(args.stacks)]

    patches = read_patches()
    rows, per_cell = [], []
    for cfg in cfgs.itertuples(index=False):
        cfg = pd.Series(cfg._asdict())
        cells = classify(cfg, patches.get(cfg.stack, set()))
        attrs = cells.attrs
        cells = measure(cells, cfg)
        cells = select_region(cells, cfg)       # analysed box + z-edge exclusion
        cells.attrs = attrs
        row = stats_for(cells, cfg)
        rows.append(row)
        per_cell.append(cells)
        print(f"{cfg.stack}: Cyto+ {row['n_cytopos']} (median SENSOR "
              f"{row.get('median_sensor_cytopos')}) vs Cyto- {row['n_cytoneg']} "
              f"({row.get('median_sensor_cytoneg')})  fold {row.get('fold_change')}  "
              f"r={row.get('rank_biserial_r')}  P(sup)={row.get('prob_superiority_pct')}%")

    table = pd.DataFrame(rows).sort_values("stack")
    cells = pd.concat(per_cell, ignore_index=True)
    table.to_csv(HERE / "quantification_stats.csv", index=False)
    cells.to_csv(HERE / "quantification_cells.csv", index=False)
    cols = ["stack", "n_cytopos", "n_cytoneg", "cytopos_pct_of_scored",
            "median_sensor_cytopos", "median_sensor_cytoneg", "fold_change",
            "rank_biserial_r", "prob_superiority_pct", "p_value"]
    pd.set_option("display.width", 200)
    print(f"\n{table[cols].to_string(index=False)}")
    print(f"\nwrote {HERE/'quantification_stats.csv'} and {HERE/'quantification_cells.csv'}")
    return 0


if __name__ == "__main__":
    sys.exit(main())
```
