## supplemental tables and scripts for "Long non-coding RNA-3′UTR interactions coordinate GABA signaling in the brain": README.md

# SENSOR analysis — reproduction

Everything needed to recompute the reported per-cell quantification, the Cyto+ vs Cyto−
statistics, the local baseline analysis and the failure rates, starting from the segmentation
masks.

    python quantify_cells.py          # per-cell quantification + the reported statistics
    python local_baseline.py          # the local baseline, its diagnostics and its null
    python local_baseline.py --k-sweep    # + the evidence that fixed K
    python failure_rate.py            # the failure-rate table
    python failure_rate.py --check    # additionally verifies against the archived tables

Each script may be restricted with `--stacks i347s1 i348s2`.

`python >= 3.9`, `numpy`, `pandas`, `scipy`, `tifffile`. No code outside this directory is
imported. Each script takes ~1–2 min for the eight regions at a measured peak of ~2.5 GB
(whole 3-D masks and images are held in memory one region at a time).

The three scripts share one implementation of the classification and of the local reference,
which lives in `failure_rate.py` and is imported by the other two, so all three score the same
cells by the same rules; every shared parameter is defined once, in the block at the top of
that file, next to the reasoning that fixed it.

## Files

| file | role |
|---|---|
| `quantify_cells.py` | per-cell measurement → classify → region → Cyto+ vs Cyto− statistics |
| `local_baseline.py` | classify → local baseline → reach / gradient / null diagnostics, K sweep |
| `failure_rate.py` | the whole procedure: classify → local baseline → z-edge → bar → rate |
| `failure_rate_inputs.csv` | one row per analysed region: image/mask paths, voxel size, the analysed box, and every parameter that is **not** shared between regions |
| `patch_exclusions.csv` | GABA mask objects confirmed by eye to be stain patches or debris rather than cells |
| `quantification_cells.csv` | output: **the per-cell SENSOR quantification table** — one row per nucleus |
| `quantification_stats.csv` | output: the reported per-region Cyto+ vs Cyto− statistics |
| `local_baseline_cells.csv` | output: one row per nucleus with its local reference, `rel`, reach and local percentile |
| `local_baseline_summary.csv` | output: per-region reach, gradient, r before/after the local reference, and the null |
| `local_baseline_k_sweep.csv` | output (`--k-sweep`): the null spread against K |
| `failure_rate.csv` | output: the reported failure-rate table |
| `failure_rate_cells.csv` | output: every nucleus, with its group, local reference, `rel`, and its exclusion reason if it was not scored |

Shared parameters live in a single block at the top of `failure_rate.py`, each with the
reasoning that fixed it. Anything that varies between regions is a column of the inputs file
rather than a branch in the code, so no per-stack special case is hidden.

## The per-cell table (`quantification_cells.csv`)

One row per nucleus mask, every region, **including the cells that were not scored** — the
`excluded` column says why each one was left out, so any subset in the paper can be rebuilt
from this file alone.

| column | meaning |
|---|---|
| `stack` | imaged region |
| `cell_label` | label of the nucleus in that region's nucleus mask (the join key to the masks) |
| `group` | `Cyto+` / `Cyto-` / `borderline`, from the GABA channel alone |
| `sensor` | **mean SENSOR intensity over the nucleus**, 8-bit-equivalent grey levels |
| `sensor_max`, `sensor_std` | maximum and standard deviation of SENSOR over the same voxels |
| `nuclear_gaba` | mean GABA intensity over the nucleus, same scale (this is what the borderline cut is applied to) |
| `gaba_overlap_frac` | fraction of the nucleus lying inside the patch-cleaned GABA mask (≥ 0.60 → Cyto+) |
| `volume_voxels`, `volume_um3` | nucleus size, in voxels and in cubic microns |
| `centroid_x/y/z` | centroid in voxels; `x_um`, `y_um`, `z_um` are the same point in microns |
| `excluded` | `scored`, or the reason: `glia`, `empty_mask`, `borderline`, `outside_region`, `z_edge` |
| `scored` | `True` for the cells the reported statistics are computed from |

Intensities are put on a common 8-bit-equivalent scale by the divisors in the inputs file: the
acquisitions are not all the same bit depth, and the borderline cut is an absolute grey level.

`local_baseline_cells.csv` carries the same identifying columns plus `local_ref`, `rel`,
`reach_um`, `local_pct` and `in_reference_pool`, so the two per-cell tables join on
(`stack`, `cell_label`).

## The statistics (`quantification_stats.csv`)

Per region: group sizes and medians, the fold change between them, a two-sided Mann–Whitney U
test, and the rank-biserial correlation `rank_biserial_r` with its readable form
`prob_superiority_pct` — the probability that a randomly chosen Cyto+ cell is brighter than a
randomly chosen Cyto− cell. The effect size carries the claim; at these n the p value is not
in question and is reported only for completeness.

**The table is the cropped region alone.** Statistics are computed over the **scored** cells
only — inside the analysed box, not z-truncated, not glia, not an empty mask, not borderline —
which is the same set the figures are drawn from and the same set the failure rate uses, so the
figure, this table and `failure_rate.csv` all describe the same cells. The box is echoed into
the table (`x_min`, `x_max`, `y_min`, `y_max`) so it declares its own scope, and matches
`cropped/regions.csv` for the canonical regions.

No whole-field statistics are reported. In some regions the field of view extends past the
optic lobe, and the discrete bright clusters outside it inflate r without saying anything about
the tissue under study. `quantification_cells.csv` still lists every nucleus with its
`excluded` reason (`outside_region` for cells outside the box), so a different scope is
recomputable — but nothing quoted mixes the two.

## The local baseline (`local_baseline_summary.csv`)

Each cell's reference is the median SENSOR of its K = 10 nearest `Cyto-` cells in microns;
`rel = sensor / local_ref`. The table reports, per region:

- `gradient_p90_p10` — how far the local reference varies across the analysed region
  (1.42–1.75x here). This is the reason a stack-wide reference is not used: the variation is
  comparable in size to the enrichment being tested.
- `reach_um_p50/p95/max` — the distance the neighbourhood had to span. Reported, never used to
  filter: a reach limit was tested and rejected, because it deletes real cells wherever off
  cells are genuinely sparse.
- `r_global` vs `r_local` — the same rank-biserial effect size on raw SENSOR and on `rel`. The
  local reference **increases** it in every region (e.g. 0.42 → 0.74 in i339s1), i.e. it removes
  a nuisance gradient rather than manufacturing the separation.
- `null_median_rel`, `null_mad_rel` — `Cyto-` cells scored leave-one-out against their own
  neighbours. The null sits at 0.99–1.00 in all eight regions, which is the check that the
  reference tracks the tissue.

Every column of this table is over the **scored** cells, i.e. the cropped region alone, to match
the figures and `quantification_stats.csv`. The reference *pool* behind `local_ref` is still
drawn from the whole stack on purpose: an off cell just outside the box is still the right
neighbour for a cell just inside it.

`--k-sweep` writes that null spread against K = 4, 6, 10, 15, 25, 50 (`Cyto-` cells only, so
nothing in the choice can see the measurement). It is minimised at K = 4–10 in every region and
K = 50 is worse in every region; K = 10 is used, rather than the numerical optimum K = 4,
because the median of four cells is fragile.

## The failure rate

For each region, the fraction of GABA-positive (Cyto+) neurons whose SENSOR signal fails to
rise above the SENSOR level of the off cells immediately around them.

1. **Classify** every nucleus from the GABA channel alone. `Cyto+` = at least 60% of the
   nucleus lies inside a (patch-cleaned) GABA mask object; `borderline` = mask-negative but
   nuclear GABA above `round(background) + offset`, excluded from both groups; `Cyto-` =
   mask-negative and at or below that cut. Glia-positive nuclei are dropped first where the
   counterstain exists, and DAPI-empty masks are dropped.
2. **Local baseline.** Each cell's reference is the median SENSOR of its **K = 10 nearest
   `Cyto-` cells**, by Euclidean distance between mask centroids **in microns**
   (`x·pixel, y·pixel, z·z_step`). `rel = sensor / reference`. `Cyto-` cells are scored
   leave-one-out. Cells below `REF_FLOOR = 2` grey levels are excluded from the reference pool
   (they are out-of-domain, not off cells) but are still scored themselves.
   **There is no distance cutoff:** the neighbourhood is defined by count. The distance the
   reference had to reach is reported per cell as `reach_um` — median 3.3–5.8 µm, 95th
   percentile 5.4–20.3 µm — rather than used to filter. A reach limit was tested and rejected,
   because it removes real cells wherever off cells are genuinely sparse.
3. **z-edge exclusion.** Cells within 4 z slices of the stack's first or last imaged plane are
   dropped from **both** groups: such nuclei are truncated, so their SENSOR mean comes from a
   clipped sliver while their reference is built from whole neighbours.
4. **Bar.** The 75th percentile of the surviving `Cyto-` cells' `rel`, computed **per region**.
5. **Rate.** Fraction of `Cyto+` cells at or below that bar, with a 95% Wilson interval.

Steps 1–2 use the whole stack; steps 3–5 the analysed region. Nothing in steps 1 or 3 reads
the SENSOR channel, so neither the classification nor the exclusions can be tuned to the
measurement.

**Quote the rate against its null.** Because the bar is a percentile of each region's own
background, a cell with no enrichment clears it 25% of the time by construction. The excess
over 25% is the result; the rate alone is not interpretable.

## Caveats carried by the inputs, not hidden in the code

- **i284s2 used a different GABA antibody** (segmentation `cellprob −3.0`, borderline offset
  59 from the p90 rule on its own negatives). Its Cyto+ *fraction* is not comparable to the
  other regions. Its *failure rate* is, because it is scored against its own background.
- **i339s1/s2 have no glia counterstain**, so the glia exclusion cannot be applied there. Its
  measured cost elsewhere is |Δr| ≤ 0.043 and ≤ 1.2 percentage points of Cyto+.
- **`patch_exclusions.csv` is a human judgement.** Patches were identified on the GABA channel
  and confirmed by eye, blind to SENSOR; the review images are the source for those calls. No
  automatic rule selects which objects are dropped.
- **Segmentation is not re-run here.** The script starts from the Cellpose-SAM masks
  (cellpose 4.0.9, `do_3D`, `min_size 100`, `max_size_fraction 0.02`) and the per-cell
  quantification derived from them; per-channel parameters are recorded per region in the
  inputs file.

## Verification

`--check` re-derives every number from the original analysis pipeline's own intermediate
tables and requires cell-by-cell agreement. As of the last run: **75,380 nuclei across eight
regions, zero group mismatches, `rel` agreeing to 5×10⁻⁴** (the archived tables store `rel`
rounded to three decimals), and all eight rates identical to the reported values.

---

## Note for this copy

The three per-cell output tables (`quantification_cells.csv`, `local_baseline_cells.csv`,
`failure_rate_cells.csv`, ~30 MB together) are **not duplicated here**. They are Supplementary
Table S11, which ships with the manuscript. `outputs/` keeps the four summary tables, which are
the numbers reported in the paper.

`failure_rate.py --check` verifies against the original pipeline's intermediate tables by globbing
`../analysis/sensor/local_baseline_nuclei*.csv`. Those live in the working tree this folder was
staged from, not here, so `--check` only runs there. Everything else runs from the segmentation
masks, whose paths and per-region parameters are recorded in `failure_rate_inputs.csv`.
