## supplemental tables and scripts for "Long non-coding RNA-3′UTR interactions coordinate GABA signaling in the brain": README.txt

Analysis code and supplementary tables
Long non-coding RNA-3'UTR interactions coordinate GABA signaling in the brain

Provided for peer review. The citable version of the code will be deposited in a public
repository with a DOI (listed in the Key Resources Table and Data-and-code-availability
statement of the manuscript).

Contents
  analysis_code/            Analysis and figure-generation scripts, grouped by module
                            (blast, correlation, duplex, orf, sensor). SUBMISSION_CODE_TABLE.xlsx
                            lists every script and what it produces. Small derived inputs each
                            module needs are under its data/ or outputs/ subfolder.
  supplementary_tables/     Supplementary Tables S1-S11 (see SUPPLEMENTARY_TABLES_INDEX for a
                            one-line description of each; every table also has its own About sheet).

Not included
  Large upstream inputs (single-cell expression matrices, raw imaging) are too large for this
  bundle and will be deposited in a data repository; scripts read them by their documented paths.
